# Excessive Censoring Degrades Individual-Specific Cortical Parcellations and Personalized TMS Targets

**DOI:** 10.64898/2026.03.09.710457

**Authors:** Trevor Wei Kiat Tan, Ru Kong, Aihuiping Xue, Jingwen Cheng, Bjorn Burgher, Luca Cocchi, Shan H. Siddiqi, Thomas E. Nichols, Amanda F. Mejia, Phern-Chern Tor, B. T. Thomas Yeo

**Affiliations:** Centre for Sleep and Cognition & Centre for Translational MR Research, Yong Loo Lin School of Medicine, National University of Singapore, Singapore, Singapore; Department of Electrical and Computer Engineering, National University of Singapore, Singapore, Singapore; Department of Medicine, Healthy Longevity Translational Research Programme, Human Potential Translational Research Programme & Institute for Digital Medicine (WisDM), Yong Loo Lin School of Medicine, National University of Singapore, Singapore, Singapore; N.1 Institute for Health, National University of Singapore, Singapore, Singapore; Integrative Sciences and Engineering Programme (ISEP), National University of Singapore, Singapore, Singapore; Clinical Brain Networks Group, QIMR Berghofer Medical Research Institute, Brisbane, Queensland, Australia; School of Biomedical Sciences, Faculty of Health, Medicine and Behavioural Sciences, University of Queensland, Brisbane, Queensland, Australia; Department of Psychiatry, Brigham & Women’s Hospital, Harvard Medical School, Boston, MA, USA; Center for Brain Circuit Therapeutics, Brigham & Women’s Hospital, Harvard Medical School, Boston, MA, USA; Big Data Institute, Li Ka Shing Centre for Health Information and Discovery, Nuffield Department of Population Health, University of Oxford, Oxford, UK; Centre for Integrative Neuroimaging (OxCIN), FMRIB, Nuffield Department of Clinical Neurosciences, University of Oxford, Oxford, UK; Department of Statistics, Indiana University, Bloomington, IN, USA; Department of Psychological Medicine, National University Hospital, Singapore, Singapore; Martinos Center for Biomedical Imaging, Massachusetts General Hospital, Charlestown, MA, USA

## Abstract

Head motion systematically biases functional connectivity (FC) estimates in resting-state functional MRI (rs-fMRI). A common mitigation strategy is to censor high-motion volumes and discard high-motion runs. However, overly stringent censoring risks discarding signal alongside noise, potentially degrading FC estimates. Here, we test the efficacy of various censoring strategies on individual-specific cortical parcellations and personalized transcranial magnetic stimulation (TMS) target selection. Using precision-fMRI datasets comprising 50 individuals, we define individualized “ground-truth” references from ≥1 hour of low-motion data per participant. We then simulate 10-min or 20-min rs-fMRI sessions with varying motion levels from the remaining data, yielding final samples of 22 and 19 participants, respectively. Higher motion produces parcellations and TMS targets that deviate further from the ground-truth references. However, at any given motion level, lenient censoring produces higher quality parcellations and personalized TMS targets than strict censoring. The improvement is comparable to doubling scan duration from 20 to 40 min under strict censoring. With personalized connectome-guided TMS, a common dilemma is whether to re-scan patients with only high-motion runs. A mixed-motion session with one low-motion run and one high-motion run may often be considered usable after discarding the high-motion run and strict censoring. We find that lenient censoring of high-motion-only sessions yields TMS targets comparable to – or even better than – those derived from strictly censored mixed-motion sessions. Therefore, within the motion range and parcellation/TMS targeting frameworks evaluated here, patients may not need to be re-scanned solely because all runs exceed strict censoring criteria.

## 1. INTRODUCTION

Head motion is a prominent challenge in resting-state functional MRI (rs-fMRI). Motion is known to systematically bias estimates of functional connectivity, and even submillimeter movements have been shown to introduce widespread artifacts (Power et al., 2012; Satterthwaite et al., 2012; Van Dijk et al., 2012; Yan et al., 2013). To mitigate these effects, many studies employ motion correction procedures such as censoring (i.e., removal) of high-motion volumes and exclusion of high-motion rs-fMRI runs (Marek et al., 2022; Lynch et al., 2024; Ooi et al., 2025).

However, emerging evidence suggests that censoring introduces its own challenges. Excluding high-motion rs-fMRI runs – for example, those falling below a minimum usable duration after censoring – can lead to the complete removal of some participants, thereby reducing sample representativeness (Nebel et al., 2022; Dhamala et al., 2025). In large population studies such as the Adolescent Brain Cognitive Development (ABCD) study, this practice disproportionately excludes male, non-White children and those from lower socioeconomic backgrounds, who tend to exhibit higher in-scanner motion (Chen et al., 2022; Cosgrove et al., 2022; Peverill et al., 2025).

Furthermore, motion-contaminated fMRI volumes are not necessarily devoid of signal. Because head motion exists on a continuum, stringent censoring may discard volumes with relatively low noise and relatively meaningful signal, potentially degrading functional connectivity (FC) estimates (Phạm et al., 2023). To evaluate this trade-off, Phạm and colleagues leveraged the observation that sufficiently long, artifact-free acquisitions converge on a stable estimate of an individual’s functional architecture (Laumann et al., 2015; Braga & Buckner, 2017; Gordon et al., 2017; Du et al., 2024), which can serve as a silver-standard ground-truth. Using this benchmark, they demonstrated that stringent censoring can indeed worsen FC estimates (Phạm et al., 2023). However, the implication of the worsened FC estimates for further downstream applications remains unclear.

In this study, we investigated how censoring and exclusion of high-motion runs affect the downstream applications of individual-specific cortical parcellations (Kong et al., 2019) and personalized transcranial magnetic stimulation (TMS) target selection (Fox et al., 2013; Kong et al., 2026a). Individual-specific brain parcellations have uncovered fundamental organizational principles of the human brain (Gordon et al., 2023; Du et al., 2024) and brain disorders (Laumann et al., 2021; Lynch et al., 2024; Vaghi et al., 2025). In parallel, using rs-fMRI to derive personalized stimulation targets for treatment-resistant psychiatric conditions has emerged as one of the most impactful clinical applications of resting-state functional connectivity (Cole et al., 2020; Oathes et al., 2024; DeSouza et al., 2025; Taylor et al., 2026; Kong et al., 2026b).

To assess the impact of censoring and high-motion run exclusion, we leveraged precision fMRI datasets with hours of data per participant (Zuo et al., 2014; Choe et al., 2015; Gorgolewski et al., 2015; Poldrack et al., 2015; Gordon et al., 2017; Noble et al., 2017; O’Connor et al., 2017; Newbold et al., 2020; Allen et al., 2022). Following the strategy of Phạm et al. (2023), we defined individual-specific “ground-truth” references using aggressively censored, high-quality rs-fMRI sessions of sufficient scan duration. These references comprised parcellations estimated with the multi-session hierarchical Bayesian model (MS-HBM; Kong et al., 2019) and personalized TMS targets estimated using the tree-based MS-HBM (Kong et al., 2026a), cone (Fox et al., 2013), and cluster (Cash et al., 2021b) algorithms.

We then simulated fMRI runs with varying motion levels from each participant’s remaining data and applied different censoring strategies. For clarity, censoring was performed based on framewise displacement (FD) and DVARS, where D refers to temporal derivative of timecourses and VARS refers to root mean squared variance over voxels (Power et al., 2012). In the main analyses, FD was computed after respiratory pseudo-motion filtering (Fair et al., 2020; Gratton et al., 2020). “Strict censoring” refers to censoring frames with FD > 0.08 mm or DVARS > 50, whereas “lenient censoring” refers to censoring frames with FD > 0.5 mm or DVARS > 100. In sensitivity analyses using root-mean-square framewise displacement (FDrms; Jenkinson et al., 2002) without respiratory pseudo-motion filtering, the corresponding strict and lenient censoring criteria were FDrms > 0.2 mm or DVARS > 50, and FDrms > 0.5 mm or DVARS > 100, respectively. Finally, individual-specific parcellations and TMS targets were estimated from the censored simulated fMRI, and compared with corresponding ground-truth parcellations and targets.

Within the range of motion that could be simulated from these precision fMRI datasets, lenient censoring without excluding high-motion runs consistently produced the highest-quality parcellations and personalized TMS targets. These within-individual findings complement prior across-participant studies advocating strict motion exclusion (Power et al., 2012; Satterthwaite et al., 2012; Van Dijk et al., 2012; Yan et al., 2013), suggesting that less stringent censoring may improve the accuracy of individualized parcellations and TMS targets.

## 2. METHODS

### 2.1. Overview

An overview of the study design is illustrated in Figure 1. For each individual with hours of rs-fMRI, we first extracted the cleanest fMRI sessions (of sufficiently long duration) to estimate “ground-truth” parcellations and TMS targets. The remaining non-ground-truth fMRI sessions were used to generate simulated fMRI runs with varying degrees of motion. We then applied different censoring strategies to the simulated fMRI runs. Finally, individual-specific parcellations and TMS targets were estimated from the censored simulated fMRI and compared with ground-truth parcellations and targets.

**Figure 1.**
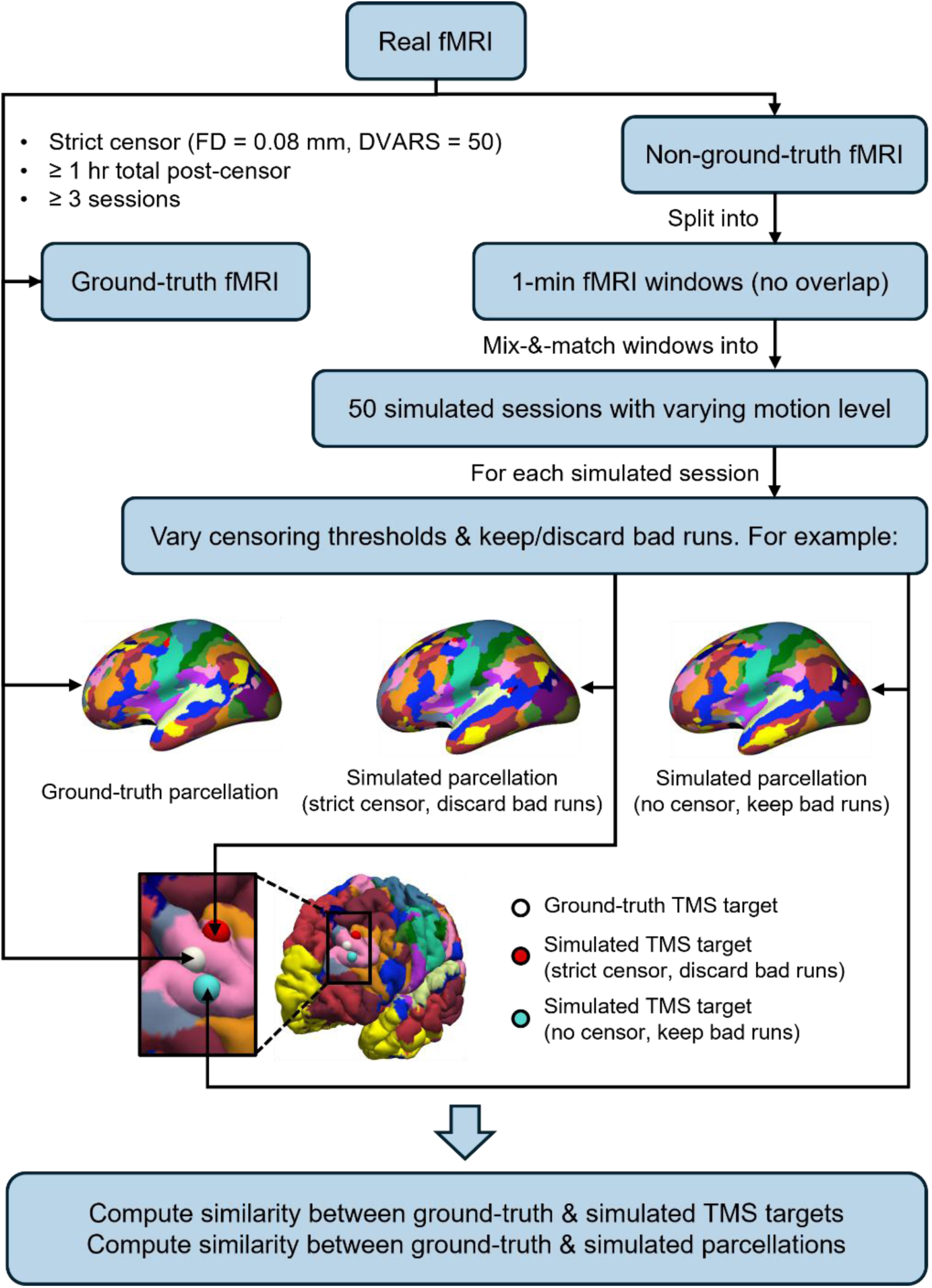
Study design. For each participant, real fMRI was divided into ground-truth and non-ground-truth data. The ground-truth fMRI comprised high-quality data obtained through strict censoring. Each participant was required to have at least 1 hour of post-censored data across a minimum of three sessions. The clean data were then used to generate an individual-specific ground-truth parcellation and ground-truth TMS target. The remaining non-ground-truth fMRI was divided into 1-minute, non-overlapping windows, which were then recombined to create 50 simulated sessions spanning a range of motion levels. For each simulated session, different censoring strategies were applied, such as strict censoring with exclusion of high-motion runs or no censoring with retention of all runs. Parcellations and TMS targets were then estimated and compared with the ground-truth. Similarity between parcellations was quantified using the Dice coefficient. TMS targets were assessed based on Euclidean distance from the ground-truth target.

### 2.2. Datasets

We considered 50 densely sampled participants from eight datasets: the Healthy Brain Network Serial Scanning Initiative (HBNSSI) dataset (O’Connor et al., 2017), the Institute of Psychology, Chinese Academy of Sciences (IPCAS6) dataset (Zuo et al., 2014; Gorgolewski et al., 2015), the Kirby Weekly (Kirby) dataset (Choe et al., 2015), the Midnight Scan Club (MSC) dataset (Gordon et al., 2017), the MyConnectome dataset (Poldrack et al., 2015), the Natural Scenes Dataset (NSD) (Allen et al., 2022), the Cast-induced Plasticity (Newbold) dataset (Newbold et al., 2020), and the Yale Test-Retest (YaleTestRetest) dataset (Noble et al., 2017). Each dataset comprised structural MRI data and densely sampled resting-state fMRI data of healthy adults. Participant characteristics are reported in Supplementary Table 1, following the Identifying Social factors that Stratify Health Opportunities and Outcomes (ISSHOOs) Set A reporting recommendations (Karran et al., 2025), to the extent permitted by the source datasets. Across the 50 considered participants, ages ranged from 19 to 56 years (mean = 32.0 ± 8.7), with 52% female. Informed consent was obtained from all participants in the original studies from which the data were drawn. The use of de-identified data for the current study is approved by the National University of Singapore (NUS) institutional review board (IRB).

The HBNSSI dataset comprised 13 participants (ages 18-45; 61.6% female), each with 14 sessions. All imaging data were collected on a Siemens Avanto 1.5T scanner. Each session comprised a 10-minute rs-fMRI run (TR = 1450 ms; multiband factor 3; voxel resolution = 2.46 × 2.46 × 2.5 mm). The anatomical T1 scans were acquired with the DESPOT1 sequence (TR = 5.2 ms; voxel resolution = 1.72 × 1.72 × 1.8 mm). Further details can be found elsewhere (O’Connor et al., 2017).

The IPCAS6 dataset comprised two participants (ages 21-25; 50% female), each with 15 sessions. All imaging data were collected on a Siemens 3T Tim Trio scanner. Each session comprised three 10-minute rs-fMRI runs (TR = 2500 ms; isotropic voxel resolution = 3.5 mm). The anatomical T1 scans were acquired with the MPRAGE sequence (TR = 1900 ms; isotropic voxel resolution = 1 mm). Further details can be found elsewhere (Zuo et al., 2014; Gorgolewski et al., 2015).

The Kirby dataset comprised a participant (age 40 at initial scan; male), with 158 sessions. All imaging data were collected on a Philips 3T Achieva scanner. Each session comprised a 6-minute, 40-second rs-fMRI run (TR = 2000 ms; isotropic voxel resolution = 3 mm). The anatomical T1 scans were acquired with the MPRAGE sequence (TR = 6.7 ms; voxel resolution = 1 × 1 × 1.2 mm). Further details can be found elsewhere (Choe et al., 2015).

The MSC dataset comprised 10 participants (ages 24-34; 50% female), each with 12 sessions. All imaging data were collected on a Siemens 3T Trio scanner. Each session comprised a 30-minute rs-fMRI run (TR = 2200 ms; isotropic voxel resolution = 4 mm). Participant MSC08 was excluded due to drowsiness during the scans. The anatomical T1 scans were acquired using the following parameters: TR = 2400 ms; isotropic voxel resolution = 0.8 mm. Further details can be found elsewhere (Gordon et al., 2017).

The MyConnectome dataset comprised a single participant (age 45 at initial scan; male). All imaging data were collected on a Siemens 3T Skyra scanner. The participant had 109 sessions. After excluding sessions with differing rs-fMRI acquisition parameters or scanners, 80 sessions remained. Each remaining session comprised a 10-minute rs-fMRI run (TR = 1160 ms; multiband factor 4; voxel resolution = 2.4 × 2.4 × 2 mm). The anatomical T1 scans were acquired with the MPRAGE sequence (TR = 2400 ms; isotropic voxel resolution = 0.8 mm). Further details can be found elsewhere (Poldrack et al., 2015).

The NSD dataset comprised eight participants (ages 19-32; 75% female), each with either 10 or 18 sessions. The functional imaging data were collected on a Siemens Magnetom 7T scanner. Each session comprised two 5-minute rs-fMRI runs (TR = 1600 ms; multiband factor 3; isotropic voxel resolution = 1.8 mm). The anatomical T1 scans were acquired using a Siemens Prisma 3T scanner with the MPRAGE sequence (TR = 2400 ms; isotropic voxel resolution = 0.8 mm). Further details can be found elsewhere (Allen et al., 2022).

The Newbold dataset comprised 3 participants (ages 25-35; 33.3% female). Imaging was done on a Siemens Trio 3T scanner for one participant and a Siemens Prisma 3T scanner for the other two. Only sessions prior to the cast intervention were included in this study, leaving each participant with 10, 12, or 14 sessions. Each session comprised a 30-minute rs-fMRI run. The acquisition parameters for the Siemens Trio 3T scanner were TR = 2200 ms and isotropic voxel resolution = 4 mm. The acquisition parameters for the Siemens Prisma 3T scanner were TR = 1100 ms, multiband factor 4, and isotropic voxel resolution = 2.6 mm. The anatomical T1 scans were acquired with the MPRAGE sequence for both scanners (TR = 2400 ms; isotropic voxel resolution = 0.8 mm). Further details can be found elsewhere (Newbold et al., 2020).

The YaleTestRetest dataset comprised 12 participants (ages 27-56; 50% female), each with four sessions. All imaging data were collected on two identically-configured Siemens Tim Trio 3T scanners. Each session comprised six 6-minute rs-fMRI runs (TR = 1000 ms; multiband factor 5; isotropic voxel resolution = 2 mm). The anatomical T1 scans were acquired with the MPRAGE sequence (TR = 2400 ms; isotropic voxel resolution = 1 mm). Further details can be found elsewhere (Noble et al., 2017).

### 2.3. Selection of ground-truth runs

The goal of this study is to evaluate how different censoring criteria influence the fidelity of individual-specific parcellations and TMS targets. Ideally, we would like to compare parcellations and TMS targets estimated under different censoring criteria with individual-level ground-truths. However, gold-standard ground-truths do not exist. However, previous studies have shown that extended, artifact-free acquisitions converge toward a stable representation of an individual’s functional architecture (Laumann et al., 2015; Braga & Buckner, 2017; Gordon et al., 2017; Du et al., 2024). Therefore, these converged estimates provide a reasonable benchmark for evaluating different censoring criteria.

In other words, when sufficiently long, clean data are available for an individual, the parcellations and TMS targets derived from these data can serve as a silver-standard ground-truth. To operationalize this ground-truth, a recent study has suggested that 30-min scans might be sufficient to estimate individualized brain networks (Lynch et al., 2024), so here we (conservatively) required at least 1 hour of post-censored rs-fMRI for defining the ground-truths. Furthermore, there is significant within-individual inter-session variability in resting-state functional connectivity (Mueller et al., 2013; Laumann et al., 2015). Since our goal is to estimate trait-level networks and TMS targets, we required the ground-truth to be estimated from at least three sessions of data.

We first identified ground-truth runs by running all rs-fMRI scans through an initial preprocessing stage to evaluate motion characteristics. This stage included removal of the first ten frames, slice timing correction, motion correction and respiratory pseudo-motion filtering. Framewise displacement (FD) and DVARS (D referring to temporal derivative of timecourses, VARS referring to root mean squared variance over voxels) were computed (Power et al., 2012). Frames with FD > 0.08 mm or DVARS > 50 were flagged as outliers. We note that 0.08 mm after pseudo-motion filtering has been recommended as a strict censoring threshold (Gratton et al., 2020). One frame before and two frames after the outlier frames were also censored (Power et al., 2012). Short uncensored segments < 5 frames were excluded.

Within each participant, runs were ranked by FD and DVARS to prioritize the cleanest data. More specifically, FD and DVARS were z-scored for all frames. The z-scoring was performed using the mean and standard deviation computed from all frames from all runs. For each run, the z-scored FD and DVARS were averaged across all frames to yield a run-specific summary outlier score. Post-censored segments of runs were then accumulated in order of increasing motion until the total post-censored scan time ≥ 1 hour, and the selected runs spanned ≥ 3 distinct sessions. These runs were selected as the ground-truth runs.

Participants with insufficient low-motion data and sessions for defining the ground-truth were excluded from the study. As a result, we ended up with 22 (out of 50) participants after this step.

### 2.4. Preprocessing of ground-truth runs

Of the commonly used preprocessing pipelines, the combination of censoring and global signal regression (GSR) has emerged as one of the most effective approaches for mitigating motion-related artifacts (Ciric et al., 2017; Parkes et al., 2018). In addition, for historical reasons, most connectome-guided brain stimulation studies employed global signal regression (Fox et al., 2013; Cole et al., 2020; Cash et al., 2021a; Siddiqi et al., 2021). As such, the preprocessing strategy in this study will include both censoring and GSR.

For all datasets, FreeSurfer was run on the T1 data (Fischl, 2012). The resting-state fMRI was processed with the following steps. (1) First ten frames were removed to account for magnetization stabilization (Supplementary Figures 1 to 3). (2) Slice timing correction was applied. (3) Motion correction was performed. (4) Respiratory pseudo-motion filtering was applied to the motion traces (Fair et al., 2020; Gratton et al., 2020). More specifically, for scans with fast TR < 1600 ms (HBNSSI, MyConnectome, NSD, Newbold, and YaleTestRetest), a bandstop filter of 0.2-0.33 Hz was applied to the motion traces (Fair et al., 2020; Supplementary Figures 4 and 5) based on the typical respiratory rate of a non-respiratory compromised healthy adult (Flenady et al., 2017; Natarajan et al., 2021; Chourpiliadis & Bhardwaj, 2025). The bandstop filter was implemented as a second-order IIR notch filter, applied via zero-phase forward-backward filtering (Fair et al., 2020). For scans with slower TR ≥ 1600 ms (IPCAS6, Kirby, MSC, and Newbold), a low-pass filter of 0.1 Hz was applied to the motion traces (Gratton et al., 2020). (5) Frames with FD > 0.08 mm or DVARS > 50 were marked as outliers. One frame before and two frames after each outlier frame were also censored. Uncensored segments of data shorter than five contiguous frames were also censored. (6) Alignment with structural image using boundary-based registration (Greve & Fischl, 2009) was performed. (7) Global, white matter, and ventricular signals, along with six motion parameters and their temporal derivatives, were regressed from the fMRI data. Regression coefficients were estimated from uncensored frames. (8) Censored frames were interpolated using the Lomb-Scargle periodogram (Power et al., 2014), which estimates sinusoidal components from the uncensored, unevenly sampled time series. (9) The data underwent bandpass filtering (0.009-0.08 Hz), implemented by ordinary least-squares regression of sine and cosine regressors corresponding to frequencies outside the passband. This regression formulation is conceptually aligned with the spectral interpolation step, as both procedures rely on sinusoidal representations of the time series: Lomb–Scargle estimates sinusoidal components in the presence of missing/censored samples, whereas the subsequent regression step projects out components whose frequencies fall outside the desired passband. (10) Finally, the data were projected onto the FreeSurfer fsaverage6 surface space and smoothed using a 6 mm full-width at half-maximum kernel. (11) Alignment between the structural image and MNI152 was computed using ANTs registration (Avants et al., 2011).

### 2.5. Generation of “ground-truth” parcellations and TMS targets

Uncensored frames from the ground-truth pre-processed fMRI in fsaverage6 surface space were used to estimate individual-specific cortical parcellations based on the multi-session hierarchical Bayesian model (MS-HBM; Kong et al, 2019). More specifically, we used the pretrained 17-network MS-HBM model from Kong et al. (2019), whose group-level priors were derived from the Human Connectome Project and defined on the fsaverage6 surface. For each participant, the resulting “ground-truth” parcellation was the individual-specific 17-network MS-HBM parcellation estimated from that participant’s ≥1 hour of low-motion data. The same pretrained 17-network MS-HBM model and priors were later used to estimate parcellations from the simulated sessions, ensuring that ground-truth and simulated parcellations differed only in the fMRI data used for estimation (Section 2.7).

We used the 17-network MS-HBM model because it is the parcellation model used by the tree-based personalized TMS targeting algorithm analyzed here (Kong et al., 2026a). This algorithm operates on networks defined at the 17-network MS-HBM granularity, including the dorsal attention and salience/ventral attention networks for depression targeting. These networks are relevant because depression TMS targeting aims to identify dorsolateral prefrontal cortex locations anticorrelated with the subgenual anterior cingulate cortex, and attentional networks are known to be anticorrelated with the default network (Fox et al., 2005; Fox et al., 2006).

Individual-specific MS-HBM parcellations were mapped back to each participant’s native volume space, where they served as input to the tree-based TMS depression targeting algorithm (Kong et al., 2026a). In prior work, tree-based targets compared favorably with cone (Fox et al., 2013) and cluster (Cash et al., 2021b) in terms of reliability, scalp proximity, and functional connectivity to the subgenual anterior cingulate cortex in new out-of-sample MRI sessions from the same individuals (Kong et al., 2026a). More generally, by accounting for both within-individual and between-individual variability in resting-state functional connectivity, MS-HBM can generate cortical parcellations from 10-min scans with equivalent quality to other approaches using 50 mins of fMRI (Kong et al., 2019). MS-HBM network topography also aligns with task-based fMRI activations (Du et al., 2024), predicts cognition and mental health (Kong et al., 2019), and is heritable (Anderson et al., 2021).

### 2.6. Simulated runs generation and preprocessing

Non-ground-truth fMRI runs were processed using the same procedure as the ground-truth runs (Section 2.4), except that censoring was not performed. Instead, censoring was only performed after generating the simulated rs-fMRI runs. More specifically, to evaluate how different censoring thresholds perform under varying motion levels (as measured by FD and DVARS), we generated simulated rs-fMRI runs that differed in motion contamination.

After processing the non-ground-truth fMRI runs, each run was z-scored to have zero mean and unit variance. Each run was then divided into non-overlapping 1-minute windows, which served as building blocks for the simulated runs. For a given individual, a simulated run was generated by randomly sampling ten unique windows from the non-ground-truth runs and concatenated to construct a 10-minute simulated run.

Each simulated run was assigned a motion level category based on the proportion of frames that would have been censored if we used strict censoring thresholds of FD = 0.08 mm and DVARS = 50, consistent with the censoring thresholds for the ground-truth runs. We emphasize that these thresholds were only used to quantify the motion level, and not actually used for censoring (at this stage). Four motion categories were defined: 0-25%, 25-50%, 50-75%, and 75-100% censored frames.

To ensure that our results were robust across different scan durations, we simulated fMRI sessions with two 10-min runs and fMRI sessions with one 10-min fMRI run. More specifically, a simulated fMRI session for an individual was created by pairing two random 10-min simulated runs, with no shared windows between the two runs. Because each simulated run belonged to one of four motion categories, 10 unique motion-level pairs (motion bins) were possible (Figure 2). For each participant and each motion bin, we attempted to generate 50 simulated sessions. However, many participants have low motion, so not all participants were able to contribute to all motion bins (Figure 2; Supplementary Figure 6A).

**Figure 2.**
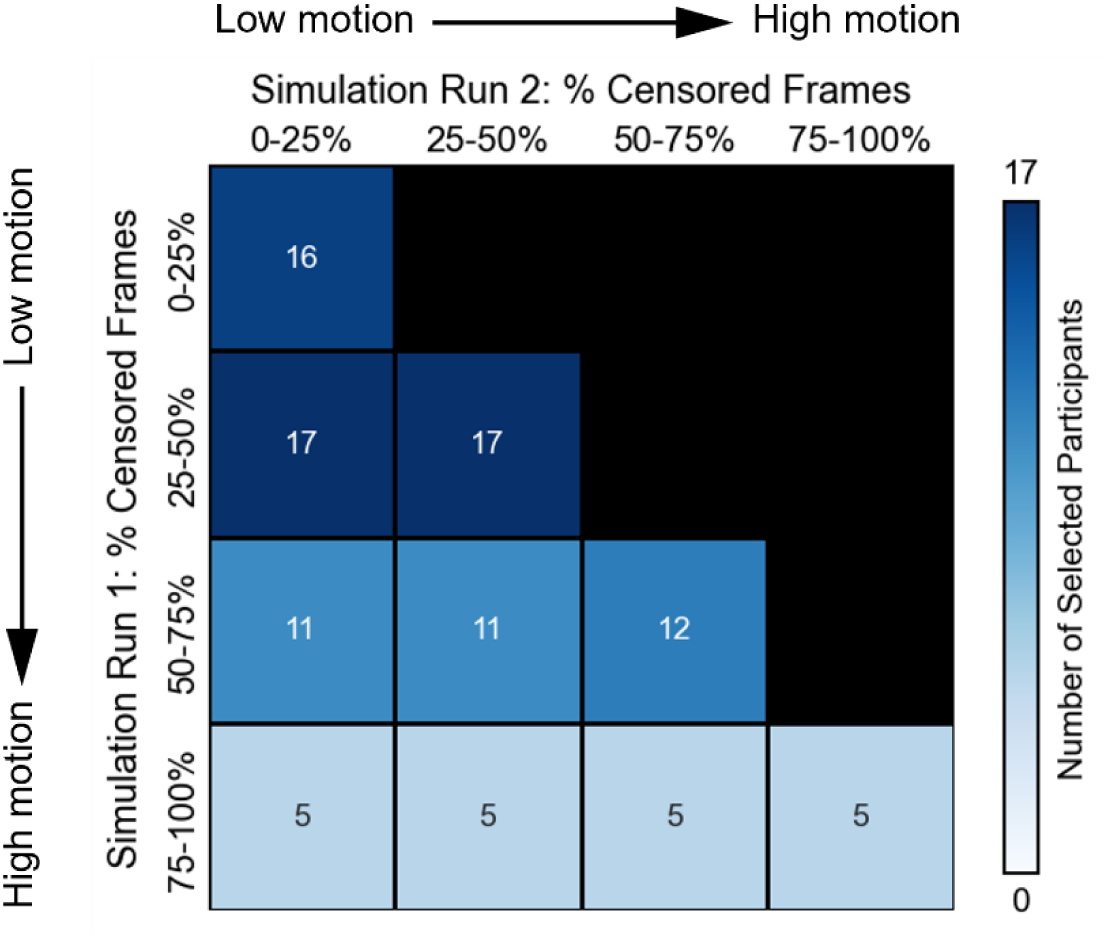
Distribution of participants (N = 19) across motion bins for the two-run FD setup. The number within each bin and the intensity of the blue shading indicate the number of participants. The number of participants who contributed simulated fMRI sessions to all ten bins is four.

Three (out of 22) participants were excluded because they could not provide 50 simulated sessions for any motion bin, resulting in a final set of 19 participants in the case of the two-run sessions. Across these 19 participants, ages ranged from 25 to 46 years (mean = 31.1 ± 5.7), with 31.6% female (Supplementary Table 1). The one-run session retained all 22 participants, whose ages ranged from 24 to 46 years (mean = 30.5 ± 5.6), with 36.4% female (Supplementary Table 1).

We note that some 1-min windows might appear in multiple simulated sessions, which meant that the sessions were not independent. This statistical dependency will be handled by linear mixed effects modeling (see Section 2.9). In the case of fMRI sessions with one 10-min run, all 22 participants were able to contribute enough simulated sessions for at least one motion bin (Supplementary Figure 7A).

### 2.7. Censoring and discarding high-motion runs

After constructing the simulated runs (Section 2.6), different combination of FD and DVARS thresholds were applied to these runs before parcellation and TMS target computation. FD thresholds of 0.08, 0.10, 0.15, 0.20, 0.50 (and none), as well as DVARS thresholds of 50, 55, 60, 65, 100 (and none) were empirically selected to yield approximately uniform increments in the proportion of censored frames. The relationship between censoring thresholds and the percentage of censored frames is shown in Supplementary Figure 8.

Many studies exclude high-motion runs (Wang et al., 2021; Chen et al., 2022; Dhamala et al., 2024; Feng et al., 2024). Here, we classified simulated runs with less than 5 minutes of data after censoring as “high-motion”. We compared scenarios in which high-motion runs were retained versus excluded.

The uncensored frames of retained simulated runs were then used to estimate individual-specific parcellations and TMS depression targets, using the same 17-network MS-HBM model (Kong et al., 2019) and the same tree-based targeting algorithm (Kong et al., 2026a) as applied to the ground-truth runs (Section 2.5). A higher Dice coefficient between parcellations from simulated runs and ground-truth parcellations indicates better censoring strategy. On the other hand, a smaller distance between TMS targets from simulated runs and ground-truth TMS targets indicates better censoring strategy.

### 2.8. Benchmarking censoring effects against scan duration

To provide an interpretable benchmark for the magnitude of the censoring effects, we quantified how much parcellation and TMS target quality improved when the total amount of data was doubled. Specifically, we compared 20-min and 40-min sessions under strict censoring without discarding high-motion runs. Both conditions followed the simulated-run paradigm described in Section 2.6 – each 20-min session comprised two 10-min simulated runs, whereas each 40-min session comprised four 10-min simulated runs. As in the main analyses, parcellations were estimated using the 17-network MS-HBM model, and TMS depression targets were estimated using the tree-based MS-HBM targeting algorithm.

Because each simulated 10-min run was constructed from unique 1-min windows, the 40-min condition required participants to have sufficient available data to construct four non-overlapping 10-min simulated runs within the same motion bin. This requirement restricted the analysis to seven participants: one from IPCAS6 (sub-0026044), one from Kirby (sub-01), three from MSC (sub-MSC07, sub-MSC09, sub-MSC10), one from MyConnectome (sub-01), and one from Newbold (sub-cast2). The analysis was performed across four motion bins, defined by the percentage of frames that would be censored under strict censoring: 0–25%, 25–50%, 50–75%, and 75–100%. For each simulated session, all runs were drawn from the same motion bin.

For each motion bin, we computed the change in parcellation Dice coefficient and TMS target Euclidean distance with respect to the ground-truth parcellation and TMS target, respectively, when scan duration was doubled from 20 to 40 min. We then averaged these changes across the four motion bins to obtain an empirical benchmark for the effect of acquiring twice as much data under strict censoring.

### 2.9. Additional analyses

#### 2.9.1. Alternative TMS algorithms

To evaluate the generalizability of our findings, we also considered the cone algorithm (Fox et al., 2013), and cluster algorithm (Cash et al., 2021b) to derive personalized TMS depression targets, as well as using the tree-based algorithm to derive a personalized TMS anxiety target (Kong et al., 2026a). We note that both cone and cluster algorithms were implemented in the native volumetric space of each individual.

#### 2.9.2. FDrms metric

Some studies use a root mean square variant of FD, FDrms (Jenkinson et al., 2002) instead of FD, and many studies do not perform respiratory pseudo-motion filtering. Therefore, control analyses were repeated using FDrms (instead of FD) with no respiratory pseudo-motion filtering.

Ground-truth runs were defined using the same selection procedure as the FD analyses (Section 2.3), but using FDrms instead of FD and without respiratory pseudo-motion filtering. FDrms threshold was set to be 0.2 mm, while the DVARS threshold remained the same, i.e., 50.

Preprocessing (Section 2.4) and generation of simulated runs (Section 2.6) were also the same except that FDrms was utilized in place of FD, and no pseudo-motion filtering was performed. When exploring censoring thresholds, we considered FDrms thresholds of 0.2, 0.25, 0.3, 0.35, and 0.5 mm, which led to similar number of censored frames as the FD thresholds (Supplementary Figure 8). DVARS thresholds remained the same: 50, 55, 60, 65, and 100.

In these analyses, we ended up with 19 (out of 50) participants for the one-run analyses (Supplementary Figure 7B) and 17 participants for the two-run analyses (Supplementary Figure 6B). Because the FD and FDrms metrics retain partially overlapping but non-identical participants, 23 unique participants contributed to at least one analysis across all FD and FDrms setups (Supplementary Table 1).

#### 2.9.3. Simultaneous nuisance regression

Performing nuisance regression and bandpass filtering as separate sequential steps can, in principle, re-introduce variance that the earlier step was meant to remove; this concern can be addressed by combining them into a single simultaneous regression (Lindquist et al., 2019). To assess whether our main conclusions depended on this preprocessing implementation, we performed a sensitivity analysis in which nuisance regression and bandpass filtering were implemented as a single simultaneous regression. Specifically, instead of applying bandpass filtering as a separate step after nuisance regression, we included the nuisance regressors together with sine and cosine regressors corresponding to frequencies outside the passband in a single regression model, and used the residuals for subsequent analyses. Both the ground-truth and non-ground-truth data were re-processed for this analysis.

#### 2.9.4. Transformed metrics

The Dice coefficient is bounded between 0 and 1, and Euclidean distance is bounded below by 0. In contrast, Gaussian linear mixed-effects models assume approximately normally distributed conditional errors, whose support extends over the real line. We therefore performed sensitivity analyses in which the linear mixed-effects models were refit using transformed outcome variables: a logit transformation for Dice coefficients and a natural logarithm transformation for Euclidean distances.

#### 2.9.5. Mean Hausdorff distance

The Dice coefficient is an overlap-based metric and can be influenced by parcel size, since a given boundary displacement will typically have a larger proportional effect on smaller parcels. We therefore performed a complementary sensitivity analysis using a boundary-based metric, which we refer to as symmetric mean Hausdorff distance (Yeo et al., 2010; Taha & Hanbury, 2015). For each pair of corresponding networks in the estimated and ground-truth parcellations, we first identified the boundary vertices of each network on the cortical surface. We then computed the geodesic distance from each boundary vertex in the simulated parcellation to the nearest boundary vertex of the corresponding network in the ground-truth parcellation, and averaged these distances. The same procedure was repeated in the reverse direction, from ground-truth to estimated boundaries. The two directional averages were then averaged to obtain a symmetric mean Hausdorff distance for each network. Finally, these network-level distances were averaged across networks to obtain a single parcellation-level mean Hausdorff distance. Smaller values indicate closer agreement with the ground-truth parcellation. We repeated the parcellation analyses using this mean Hausdorff distance metric in place of the Dice coefficient.

#### 2.9.6. Continuous runs

The simulated-run paradigm described in Section 2.6 constructs runs by concatenating 1-min windows sampled from different real runs, which reduced temporal autocorrelation and might lead to denser information than continuous runs. To evaluate whether our findings depended on this simulated-run construction, we performed a control analysis using continuous 10-min runs instead of concatenated simulated runs.

Because the simulated-run paradigm was originally introduced to overcome the limited availability of naturally occurring high-motion continuous data, this control analysis was necessarily restricted to a smaller subset of participants. To be included, a participant was required to have at least one continuous 10-min low-motion run and at least one continuous 10-min high-motion run. Runs longer than 10 min were truncated to 10 min, whereas runs shorter than 10 min were excluded. This yielded six participants: four from the MSC dataset (sub-MSC03, sub-MSC07, sub-MSC09, sub-MSC10) and two from the Newbold dataset (sub-cast2, sub-cast3).

Two continuous-run analyses were performed. First, we compared strict censoring with high-motion run exclusion against no censoring in mixed-motion sessions, where each session comprised one high-motion run and one low-motion run. For each participant, each 10-min high-motion run was paired with a randomly selected 10-min low-motion run to construct a two-run mixed-motion session. Thus, if a participant had three high-motion runs and five low-motion runs, three mixed-motion sessions were created for that participant. Under the strict-censoring condition, the high-motion run was discarded and the target or parcellation was estimated from the remaining low-motion run after censoring. Under the no-censoring condition, both runs were retained without censoring.

Second, we compared strict, lenient, and no censoring across a range of FD and DVARS thresholds without discarding high-motion runs. This analysis included the mixed-motion sessions created for the first analysis, as well as high-motion sessions comprising two high-motion runs. To construct high-motion sessions, each participant’s high-motion runs were randomly paired to form two-run sessions. Thus, if a participant had five high-motion runs, two high-motion sessions were created, and the remaining unpaired run was discarded. These continuous-run analyses were then processed using the same parcellation and TMS targeting procedures as in the main simulated-run analyses.

### 2.10. Statistical analyses

For analyses in which the same participants contributed data to both conditions being compared – such as (1) comparing two censoring strategies within a given motion bin, or (2) comparing between two motion bins – we first averaged the Dice coefficients (for parcellations) or Euclidean distances (for TMS targets) across the 50 simulated sessions for each participant and condition. These participant-level averages served as observations for the paired-sample t-tests. Thus, for each comparison, the degrees of freedom were equal to N-1, where N was the number of participants contributing data to both conditions.

To assess overall effects across all motion bins, we note that not all participants contributed data to every bin, resulting in an unbalanced repeated-measures structure. Furthermore, we note that for a given participant, simulated data from different motion bins might overlap, so the data are not independent across motion bins. These issues were addressed with linear mixed-effects models (LMEs) fitted using restricted maximum likelihood (REML) with the pymer4 package (Jolly, 2018), which in turn calls lme4 version 1.1-37 for model fitting (Bates et al., 2015). Optimization was performed using lme4’s default optimizer (nloptwrap), which internally calls the NLOPT_LN_BOBYQA algorithm (Johnson, 2007; Powell, 2009) and lmerTest version 3.1-3 (Kuznetsova et al., 2017) for significance testing.

Consistent with the paired-sample t-tests, Dice coefficients (or Euclidean distances) were first averaged across the 50 simulated sessions for each participant and condition, and these participant-level averages were used as observations for the model. P-values for the predictor of interest were obtained from two-sided t-tests using the Satterthwaite approximation for degrees of freedom with the lmerTest package. Two different LME models were used depending on the comparison: (1) When comparing different censoring strategies across all motion bins, the censoring strategy (e.g., strict vs. none) served as the primary predictor of interest, with the percentage of frames censored in each run and the number of surviving runs included as covariates. Participant identity was modelled as a random intercept, and motion bin was included as a nested random intercept within participant to account for the hierarchical structure of the data. When comparing two censoring strategies (e.g., strict vs none), the p-value for the censoring-strategy coefficient was used, with statistical significance assessed at p < 0.05. When comparing more than two censoring strategies (e.g., strict vs lenient vs none), censoring strategy was modelled as a categorical predictor (with strict censoring as the reference level), and pairwise comparisons between strategies were computed via linear contrasts on the fixed-effects coefficients. P-values were FDR-corrected (Benjamini-Hochberg) across pairwise comparisons, with q < 0.05 as the significance threshold. (2) When examining how Dice coefficients or Euclidean distances varied across motion bins, the percentage of censored frames averaged across both runs was the primary predictor of interest (also tested at p < 0.05), with participant identity modelled as a random intercept to account for repeated observations. In contrast to the LME model used for comparing censoring strategies, this model did not include a nested random intercept for the motion bin. The reason is that each participant contributed only a single observation per motion bin, so there was no within-bin variance to estimate, making a nested motion bin-level random effect non-identifiable.

## 3. RESULTS

### 3.1. Parcellation and TMS target quality decreases with higher motion and strict censoring

As a reminder, parcellation quality refers to the similarity between estimated parcellations and ground-truth parcellations. A higher Dice coefficient indicates higher parcellation quality. In the case of TMS, target quality is defined as the Euclidean distance between estimated and ground-truth targets. A smaller Euclidean distance indicates better targeting quality.

As a sanity check, we found that parcellation quality decreased with higher motion, considering for the moment only strict censoring (p = 1.2e-32; Figure 3). Similar results were obtained for personalized depression targets generated by the tree algorithm (Kong et al., 2026a), i.e., TMS target quality decreased with higher motion using strict censoring (p = 4e-8; Figure 4). Here strict censoring refers to FD threshold of 0.08 mm (after pseudo-motion filtering) and DVARS threshold of 50. For this analysis, high-motion runs were not removed, to allow for comparison with low-motion bins.

**Figure 3.**
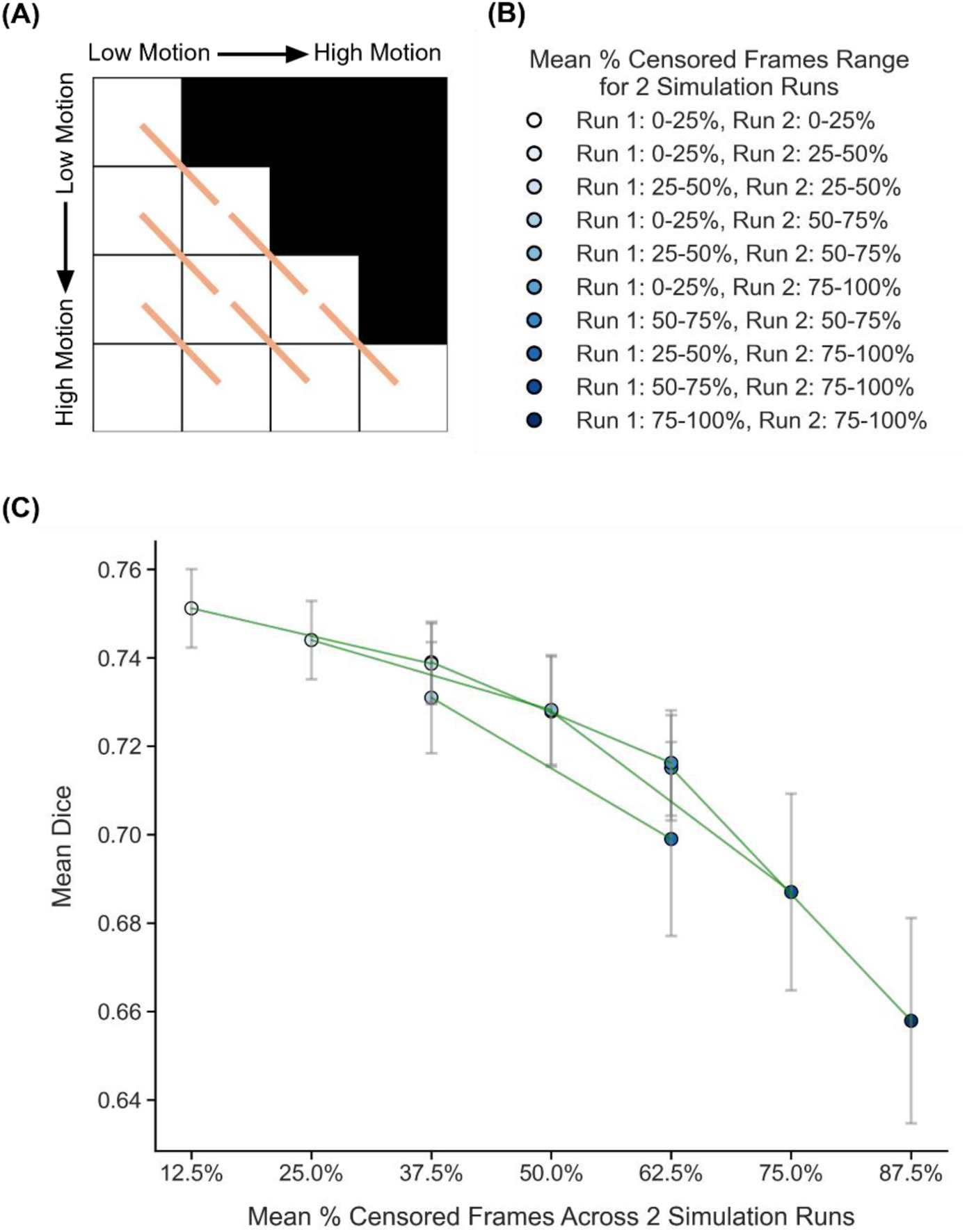
Comparison of parcellation quality (Dice) across diagonally adjacent motion bins under strict censoring, for the two-run FD setup. Here strict censoring refers to FD threshold of 0.08 mm (after pseudo-motion filtering) and DVARS threshold of 50. High-motion runs were not discarded to allow comparison with low-motion runs. Higher Dice indicates better quality. (A) Schematic of comparisons across diagonally adjacent motion bins. Each orange line indicates a single comparison. (B) Legend indicating the percentage of censored frames in each of the two simulated runs from each motion bin in panel C. (C) Relationship between parcellation quality (Dice) and motion. Each circle indicates the mean Dice coefficient across simulated runs of all participants in a motion bin. Each line indicates a single comparison between two adjacent motion bins. The error bars indicate the standard error of the mean across participants. P-values were computed using the two-sided paired-sampled t-test using only participants common to both motion bins. An overall statistical test combining all comparisons was conducted using a linear mixed effects model (p = 1.2e-32).

**Figure 4.**
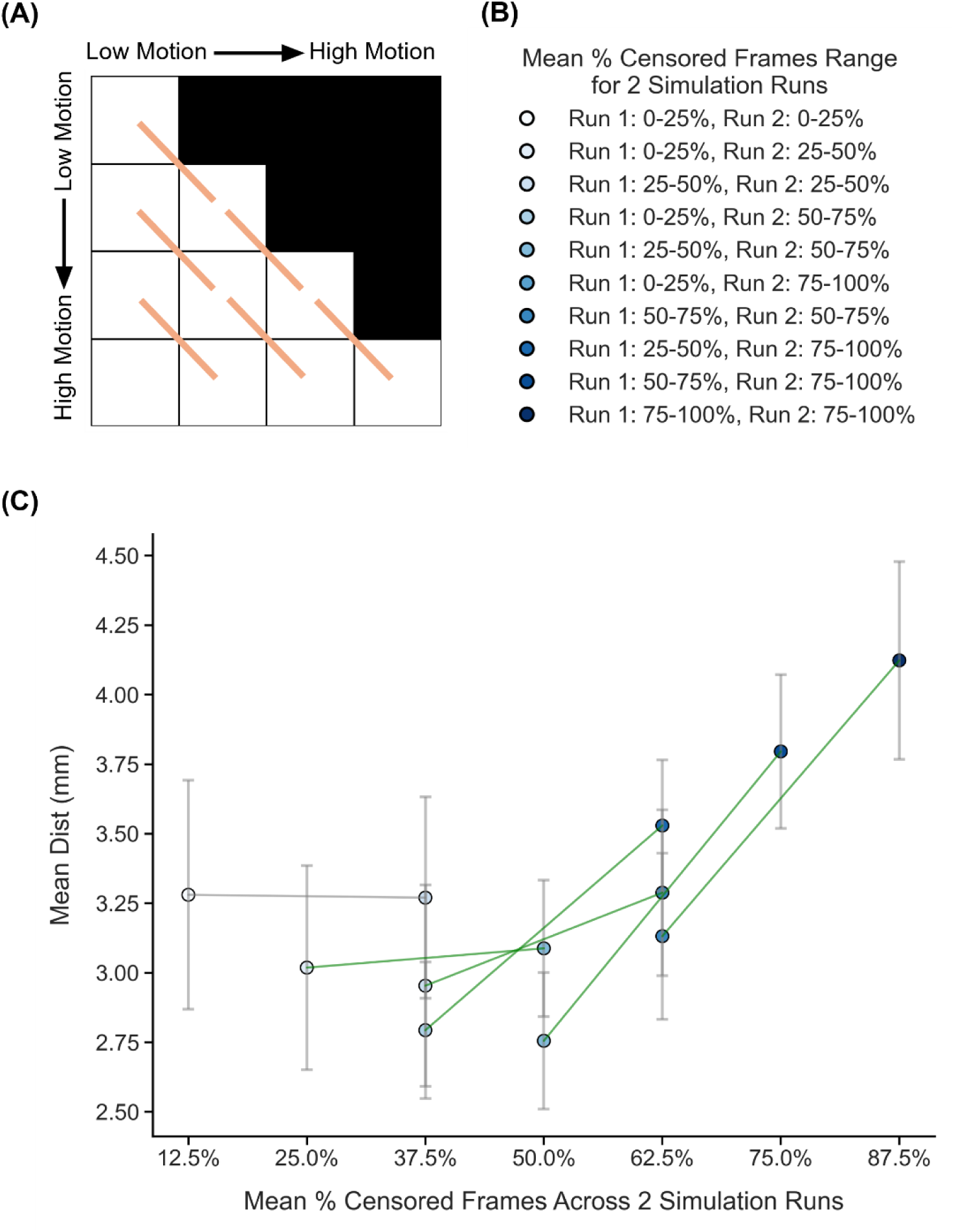
Comparison of TMS depression target quality (Euclidean distance from ground-truth targets) across diagonally adjacent motion bins under strict censoring, for the two-run FD setup. Here strict censoring refers to FD threshold of 0.08 mm (after pseudo-motion filtering) and DVARS threshold of 50. High-motion runs were not discarded to allow comparison with low-motion runs. Smaller Euclidean distance to ground-truth target indicates better quality. TMS depression targets were estimated using the tree-algorithm (Kong et al., 2026a). (A) Schematic of comparisons across diagonally adjacent motion bins. Each orange line indicates a single comparison. (B) Legend indicating the percentage of censored frames in each of the two simulated runs from each motion bin in panel C. (C) Relationship between TMS depression target quality (Euclidean distance) and motion. Each circle indicates the mean Euclidean distance across simulated runs of all participants in a motion bin. Each line indicates a single comparison between two adjacent motion bins. The error bars indicate the standard error of the mean across participants. P-values were computed using the two-sided paired-sampled t-test using only participants common to both motion bins. An overall statistical test combining all comparisons was conducted using a linear mixed effects model (p = 4e-8).

### 3.2. No censoring yields better parcellations and TMS targets than strict censoring

In the previous section, we showed that parcellation and TMS targeting quality decreased with higher motion when strict censoring was applied. Here we compared two extreme approaches – (1) strict censoring (FD = 0.08 mm and DVARS = 50) with high-motion runs removed versus (2) no censoring – to determine which approach produced parcellations and TMS targets more similar to the ground-truth.

Figure 5A shows the parcellation quality (Dice) of the two censoring approaches across 10 motion bins. In three motion bins, no comparison could be made because both fMRI runs were excluded under strict censoring. For the remaining seven bins, no censoring yielded comparable or higher parcellation quality than strict censoring. Pooling results across the seven motion bins, we observed that no censoring yielded statistically better parcellation quality than strict censoring (p = 1.4e-5; Figure 5B). The magnitude of this difference was comparable to the improvement in Dice coefficient achieved by doubling scan duration from 20 minutes to 40 minutes under strict censoring without discarding high-motion runs (brown dashed line in Figure 5B). Differences between estimated parcellations and ground-truth parcellations were mostly present near network boundaries (Figures 5C and 5D).

**Figure 5.**
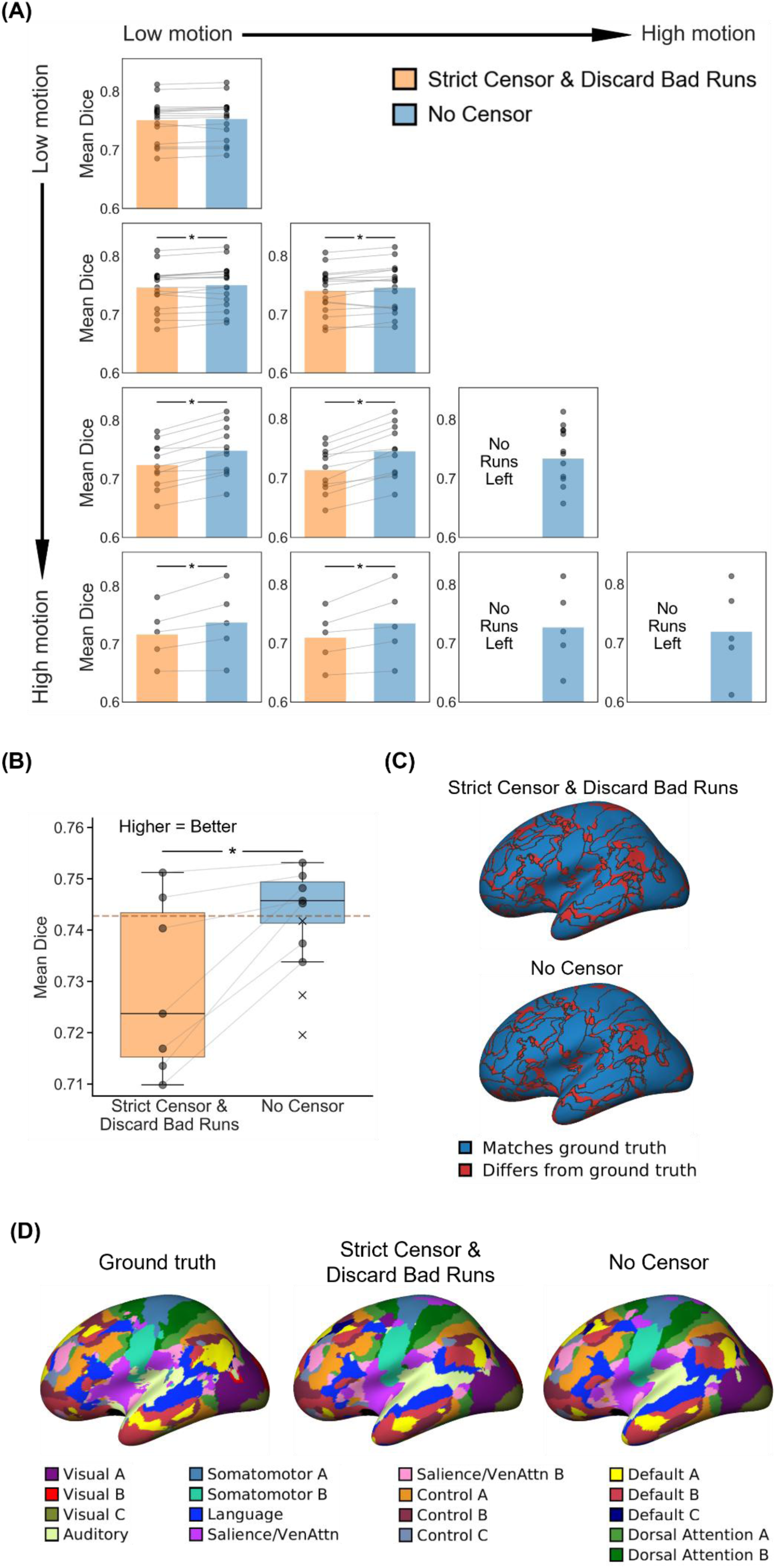
Comparison of parcellation quality (Dice) between strict censoring and no censoring, for the two-run FD setup. Here, strict censoring refers to a FD threshold of 0.08 mm (after pseudo-motion filtering) and a DVARS threshold of 50. High-motion runs were discarded. Higher Dice indicates better quality. (A) Parcellation quality for the two censoring approaches across 10 motion bins. Each cell in the grid indicates a motion bin. Each dot represents the mean Dice coefficient across 50 simulated sessions of a participant in that motion bin. P-values were computed using the two-sided paired-sampled t-test. “*” indicates statistical significance after multiple comparisons correction with FDR q < 0.05. Under strict censoring, no runs remained for the bottom-right three motion bins because all fMRI runs were discarded under this criterion. (B) Parcellation quality across all motion bins for both censoring strategies. There are seven pairs of dots, corresponding to the seven motion bins where comparisons could be made between strict censoring and no censoring. The brown dashed line indicates the improvement in Dice coefficient (0.019) achieved by doubling scan duration from 20 minutes to 40 minutes under strict censoring without discarding high-motion runs (Supplementary Figure 9). This improvement (0.019) is added to the median of strict censoring boxplot, resulting in the brown dashed line. P-value was computed using a linear mixed effects model (p = 1.4e-5). “*” indicates statistical significance after multiple comparisons correction with FDR q < 0.05. There are three “×” corresponding to the Dice coefficient of the three bottom-right bins under the no censoring condition. The × were not included in the boxplots or statistical comparisons between the two censoring strategies. For each box plot, the horizontal line indicates the median. The bottom and top edges of the box indicate the 25th and 75th percentiles, respectively. The outliers are defined as data points beyond 1.5 times the interquartile range. The whiskers extend to the most extreme data points not considered outliers. (C) Discrepancy with respect to the ground-truth parcellation in a representative participant from the highest motion bin, where the strict censoring had results. Blue regions indicate agreement with ground-truth parcellation, while red regions indicate differences. (D) Parcellations of the representative participant from panel C: ground-truth parcellation (left), simulated parcellation under strict censoring (middle), and simulated parcellation under no censoring (right).

To allow for comparison in the three highest-motion bins, we performed a supplementary analysis in which high-motion runs were not discarded under strict censoring. No censoring again outperformed strict censoring (Supplementary Figure 11). Overall, these results suggest that no censoring yielded higher agreement with the ground-truth parcellation than strict censoring.

Similar conclusions were obtained for TMS depression targets (Kong et al., 2026a). Figure 6A shows the TMS targeting quality (Euclidean distance) of the two censoring approaches across 10 motion bins. In three motion bins, no comparison could be made because both fMRI runs were excluded under strict censoring. For the remaining seven bins, similar to the parcellation results, no censoring yielded comparable or better TMS targeting quality than strict censoring. Pooling results across the seven motion bins, we observed that no censoring yielded statistically better TMS targeting than strict censoring (p = 8.2e-4; Figure 6B). The magnitude of this difference was comparable to the improvement in Euclidean distance achieved by doubling scan duration from 20 minutes to 40 minutes under strict censoring without discarding high-motion runs (brown dashed line in Figure 6B). Figure 6C illustrates the targets for a representative participant.

**Figure 6.**
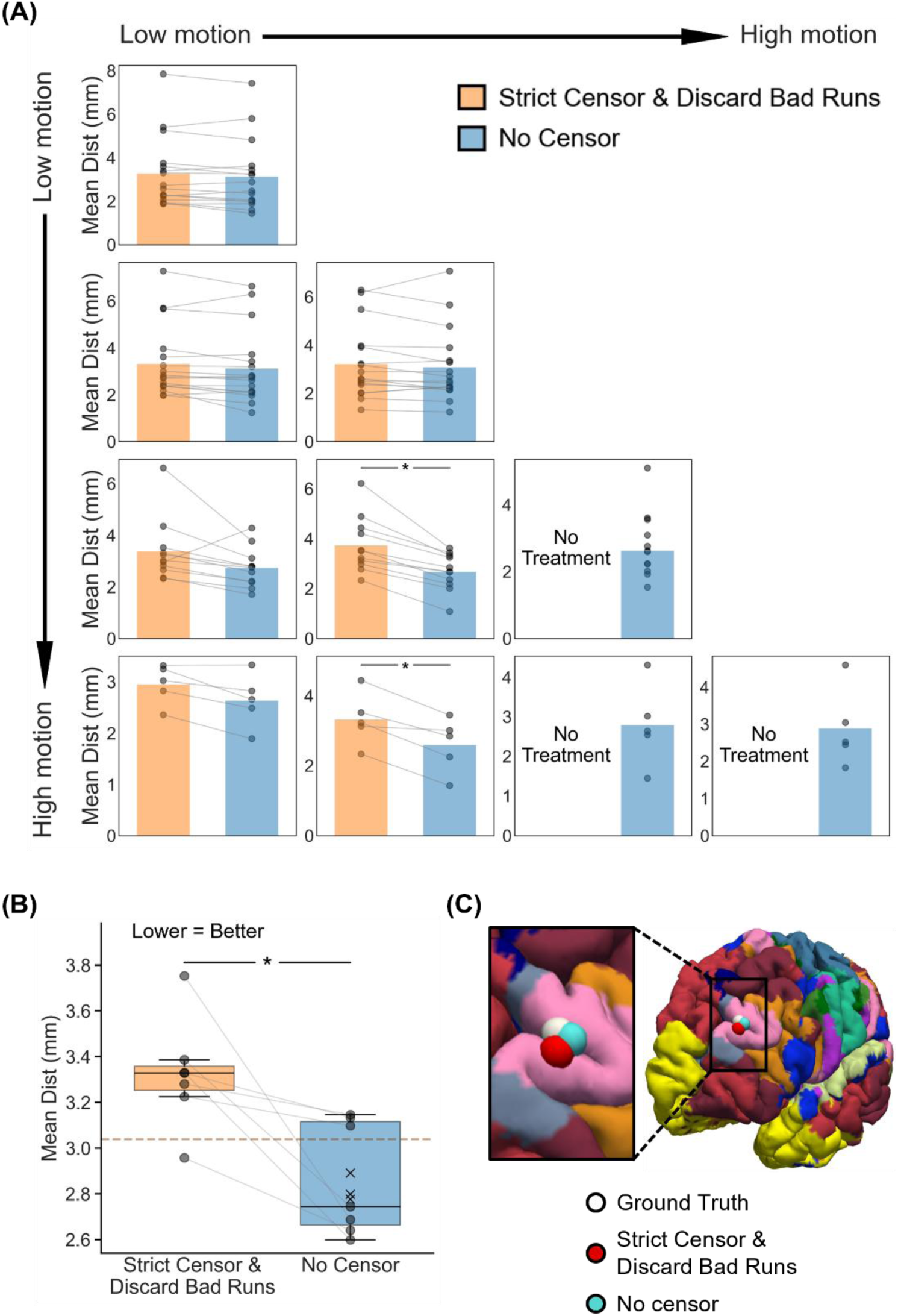
Comparison of TMS depression targeting quality (Euclidean distance to ground-truth depression targets) between strict censoring and no censoring, for the two-run FD setup. Here strict censoring refers to FD threshold of 0.08 mm (after pseudo-motion filtering) and DVARS threshold of 50. High-motion runs were discarded. Smaller Euclidean distance indicates better quality. TMS depression targets were estimated using the tree-based algorithm (Kong et al., 2026a). (A) TMS depression targeting quality (Euclidean distance) for the two censoring approaches across 10 motion bins. Each cell in the grid indicates a motion bin. Each dot represents the mean Euclidean distance across 50 simulated sessions of a participant in that motion bin. P-values were computed using the two-sided paired-sampled t-test. “*” indicates statistical significance after multiple comparisons correction with FDR q < 0.05. Under strict censoring, no runs remained for the bottom-right three motion bins because all fMRI runs were discarded under this criterion. (B) TMS depression targeting quality across all motion bins for both censoring strategies. There are seven pairs of dots, corresponding to the seven motion bins where comparisons could be made between strict censoring and no censoring. The brown dashed line indicates the improvement in Euclidean distance (0.29 mm) achieved by doubling scan duration from 20 minutes to 40 minutes under strict censoring without discarding high-motion runs (Supplementary Figure 10). This improvement (0.29 mm) is subtracted from the median of strict censoring boxplot, resulting in the brown dashed line. P-value was computed using a linear mixed effects model (p = 8.2e-4). “*” indicates statistical significance after multiple comparisons correction with FDR q < 0.05. There are three “×” corresponding to the mean Euclidean distance of the 3 bottom-right bins under the no censoring condition. The × were not included in the boxplots or statistical comparisons between the two censoring strategies. For each box plot, the horizontal line indicates the median. The bottom and top edges of the box indicate the 25th and 75th percentiles, respectively. The outliers are defined as data points beyond 1.5 times the interquartile range. The whiskers extend to the most extreme data points not considered outliers. (C) Visualization of TMS depression targets in a representative participant from the highest motion bin where the strict censoring had results.

To allow for comparison in the three highest-motion bins, we performed a supplementary analysis in which high motion runs were not discarded under strict censoring. No censoring again outperformed strict censoring (Supplementary Figure 17). Overall, these results suggest that no censoring yielded higher agreement with the ground-truth target than strict censoring.

### 3.3. No censoring in high-motion runs is better than strict censoring in mixed-motion runs

In the previous section, we showed that within a given motion bin, avoiding censoring produced better parcellations and TMS targets than strict censoring. However, this did not guarantee that parcellations from two high-motion runs without censoring were adequate for downstream analyses. On the other hand, when a participant has one low-motion and one high-motion run (mixed-motion runs), researchers often discard the high-motion run and analyze only the low-motion run after strict censoring (Power et al., 2014; Liégeois et al., 2019; Phạm et al., 2023; Lamouroux et al., 2025).

Therefore, we compared parcellations derived from two high-motion runs without censoring to those derived from mixed-motion runs subjected to strict censoring (FD = 0.08 mm, DVARS = 50) with the high-motion run discarded (Figure 7A). If the uncensored high-motion parcellations matched or exceeded the quality of the strictly censored mixed-motion parcellations, this would indicate that the high-motion data were adequate for downstream use.

**Figure 7.**
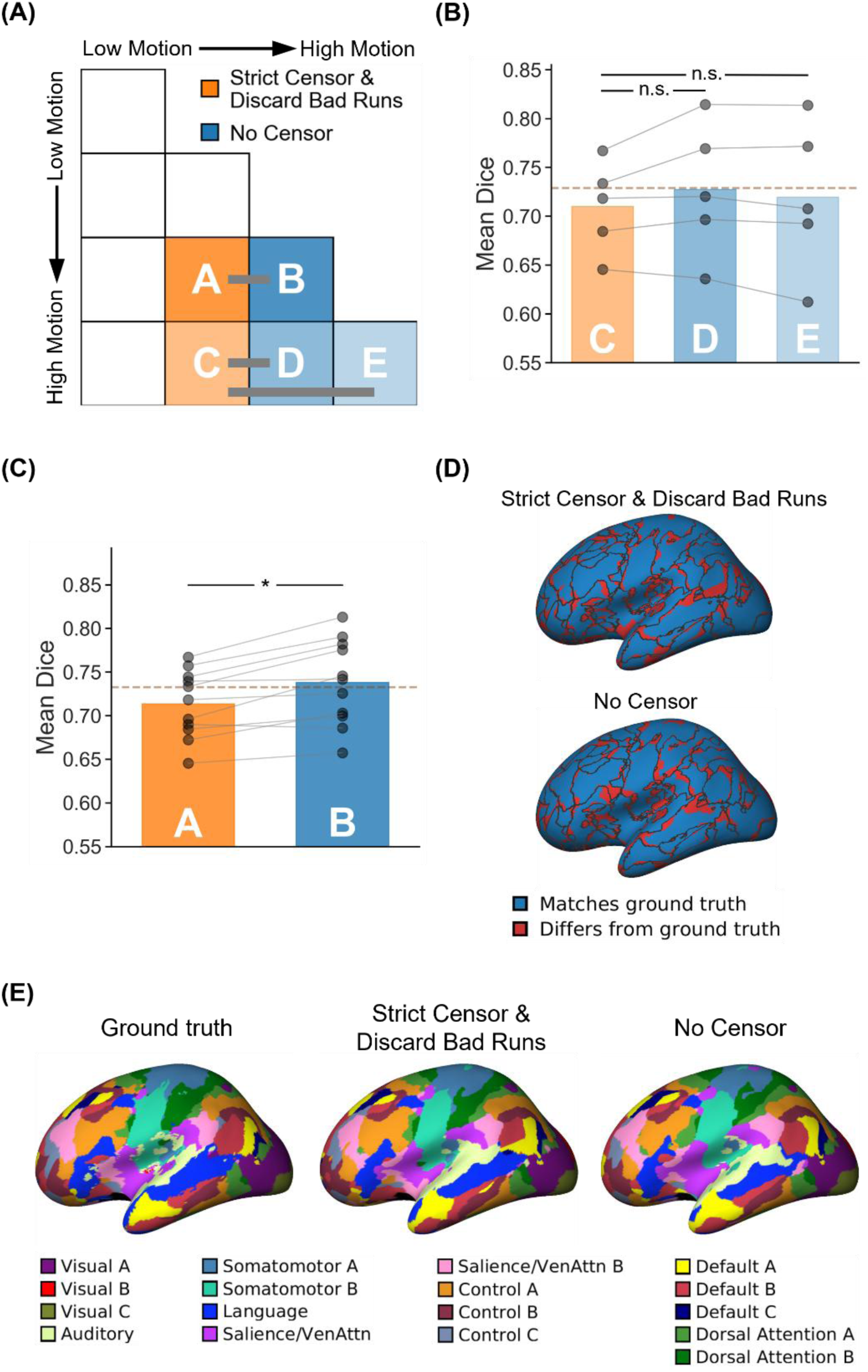
Parcellation quality (Dice) for two high-motion runs with no censoring vs mixed-motion runs with strict censoring. Here mixed-motion runs refer to sessions with one high-motion and one low-motion run, so the high-motion run will be discarded under strict censoring. Strict censoring refers to FD threshold of 0.08 mm (after pseudo-motion filtering) and DVARS threshold of 50. Higher Dice indicates better quality. (A) Schematic illustrating the comparison. Motion bins A and B were compared, while motion bins C, D and E were compared. (B) Comparison of motion bins C vs D, and C vs E. (C) Comparison of motion bins A vs B. For both panels (B) and (C), each dot represents the mean Dice coefficient across 50 simulated sessions for a participant in that motion bin. The brown dashed lines indicate the improvement in Dice coefficient (0.019) achieved by doubling scan duration from 20 minutes to 40 minutes under strict censoring without discarding high-motion runs (Supplementary Figure 9). This improvement (0.019) is added to the mean of the strict censoring bar plots, resulting in the brown dashed lines. P-values were computed using the two-sided paired-sampled t-test. Each statistical test was performed using only participants common to both motion bins. “*” indicates statistical significance after multiple comparisons correction with FDR q < 0.05. (D) Discrepancy with respect to the ground-truth parcellation in a representative participant from motion bins A (with strict censoring) and B (with no censoring). Blue regions indicate agreement with ground-truth parcellation, while red regions indicate differences. (E) Parcellations of the representative participant from panel D: ground-truth parcellation (left), simulated parcellation under strict censoring for motion bin A (middle), and simulated parcellation under no censoring for motion bin B (right). Parcellations from two high-motion runs without censoring were equivalent (panel B) or superior (panel C) to those from strictly censored mixed-motion runs.

Intriguingly, we found that parcellations from two high-motion runs without censoring were equivalent (Figure 7B) or superior (Figure 7C) to those from strictly censored mixed-motion runs. The magnitude of this difference was comparable to the improvement in Dice coefficient achieved by doubling scan duration from 20 minutes to 40 minutes under strict censoring without discarding high-motion runs (brown dashed lines in Figures 7B and 7C). Figure 7D illustrates network label agreement with the ground-truth parcellation under strict censoring versus no censoring for a representative participant. Figure 7E juxtaposes the ground-truth parcellation, parcellation estimated from mixed-motion runs with strict censoring, and parcellation estimated from two high-motion runs with no censoring in the same representative participant.

Similar to the parcellation results, we found that TMS targets from two high-motion runs without censoring were superior to those from strictly censored mixed-motion runs (Figure 8). The magnitude of this difference was comparable to the improvement in Euclidean distance achieved by doubling scan duration from 20 minutes to 40 minutes under strict censoring without discarding high-motion runs (brown dashed lines in Figures 8B and 8C). These results suggest that TMS targets from two high-motion runs without censoring might be of sufficient quality for personalized connectivity-guided TMS for major depressive disorder.

**Figure 8.**
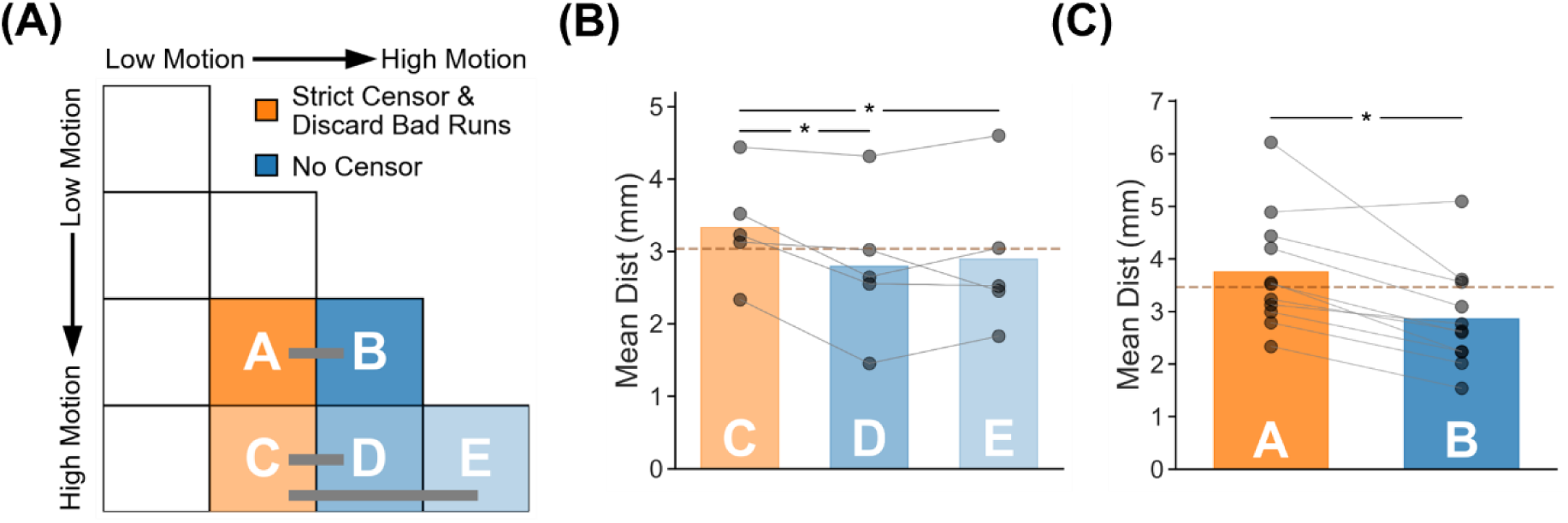
TMS depression target quality (Euclidean distance to ground-truth targets) for two high-motion runs with no censoring vs mixed-motion runs with strict censoring. Here mixed-motion runs refer to sessions with one high-motion and one low-motion run, so the high-motion run will be discarded under strict censoring. Strict censoring refers to FD threshold of 0.08 mm (after pseudo-motion filtering) and DVARS threshold of 50. Lower Euclidean distance indicates better quality. TMS depression targets were estimated using the tree-algorithm (Kong et al., 2026a). (A) Schematic illustrating the comparison. Motion bins A and B were compared, while motion bins C, D and E were compared. (B) Comparison of motion bins C vs D, and C vs E. (C) Comparison of motion bins A vs B. For both panels (B) and (C), each dot represents the mean Euclidean distance across 50 simulated sessions for a participant in that motion bin. The brown dashed lines indicate the improvement in Euclidean distance (0.29 mm) achieved by doubling scan duration from 20 minutes to 40 minutes under strict censoring without discarding high-motion runs (Supplementary Figure 10). This improvement (0.29 mm) is subtracted from the means of the strict censoring bar plots, resulting in the brown dashed lines. P-values were computed using the two-sided paired-sampled t-test. Each statistical test was performed using only participants common to both motion bins. “*” indicates statistical significance after multiple comparisons correction with FDR q < 0.05.

### 3.4. Lenient censoring yields close-to-optimal parcellations & TMS targets

The previous sections suggest that no censoring is better than strict censoring for estimating cortical parcellations and TMS targets. However, perhaps an intermediate level of censoring is better than no censoring and strict censoring. Here, we examined the effect of varying censoring thresholds without discarding high-motion runs. Specifically, we varied FD and DVARS thresholds across a range of values: FD = 0.08, 0.1, 0.15, 0.2, 0.5 or none, and DVARS = 50, 55, 60, 65, 100 or none. This resulted in 36 combinations of censoring parameters.

Figure 9A shows the parcellation quality (Dice coefficient) for the ten motion bins. For each motion bin, the Dice coefficient is plotted as a function of FD and DVARS thresholds. Brighter color indicates better parcellation quality. A striking observation was that for all motion bins, parcellation quality improves with lower FD threshold. For very high motion bin (bottom right), some intermediate level of FD threshold might be better. On the other hand, the effect of DVARS thresholds was less salient.

**Figure 9.**
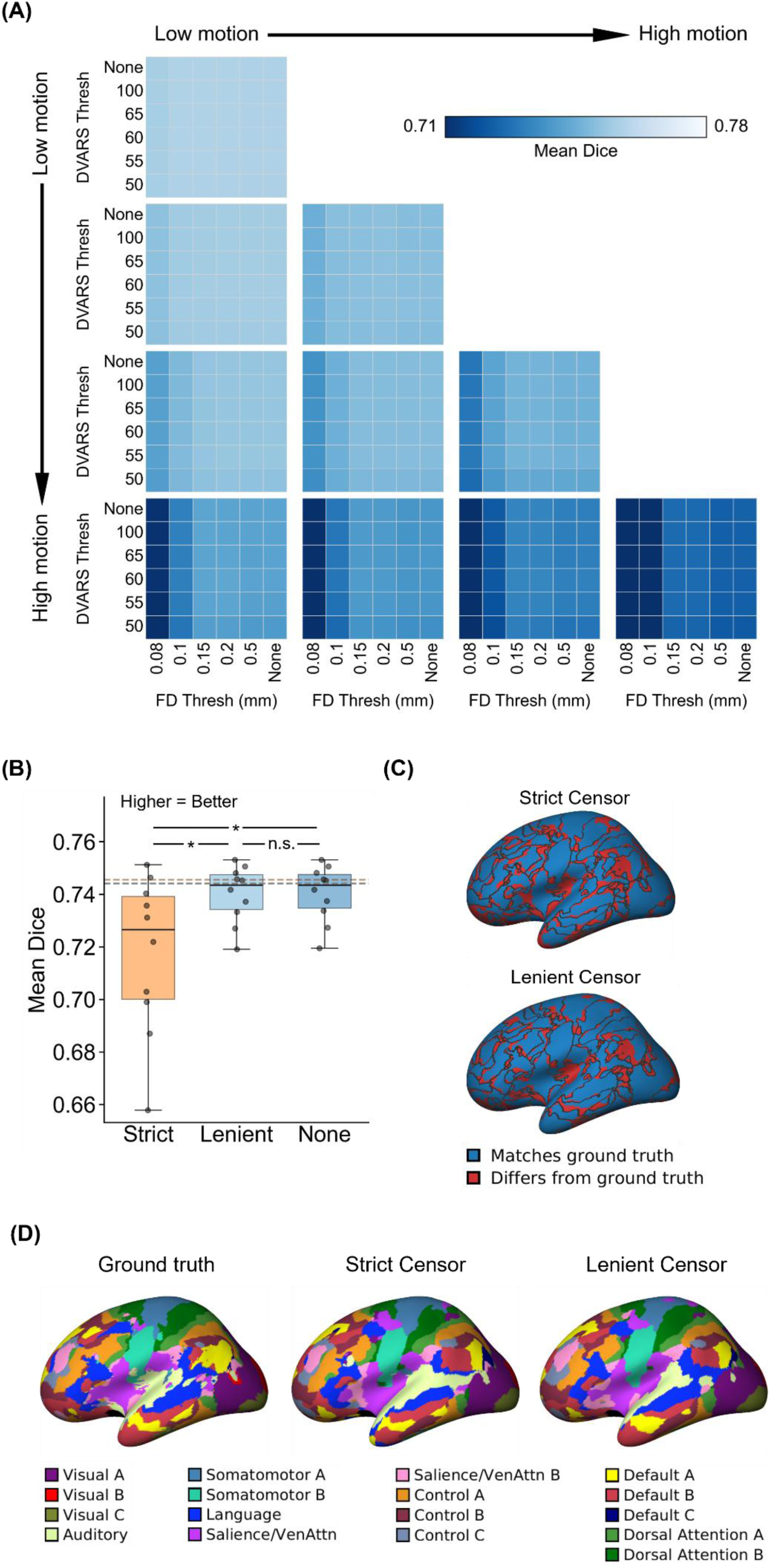
Comparison of parcellation quality (Dice) across varying FD and DVARS thresholds for the two-run FD setup. For this analysis, high-motion runs were not discarded. Higher Dice indicates better quality. (A) Parcellation quality for different combinations of FD and DVARS thresholds for 10 motion bins. Each cell in the grid indicates a motion bin. For each motion bin, there are 36 small cells corresponding to different FD and DVARS thresholds. (B) Parcellation quality of strict censoring, lenient censoring and no censoring. Here strict censoring refers to FD threshold of 0.08 mm (after pseudo-motion filtering) and DVARS threshold of 50. Lenient censoring refers to FD threshold of 0.5 mm and DVARS threshold of 100. Each dot represents the Dice coefficient for a motion bin. The brown dashed line indicates the improvement in Dice coefficient (0.019) achieved by doubling scan duration from 20 minutes to 40 minutes under strict censoring without discarding high-motion runs (Supplementary Figure 9). This improvement (0.019) is added to the median of strict censoring boxplot, resulting in the brown dashed line. The gray dashed line is the “noise ceiling”, achieved by picking the best combination of FD and DVARS threshold for each motion bin, and computing the median across motion bins. P-values were computed using a linear mixed effects model (p = 9.3e-23 for strict vs. none, p = 1.5e-22 for strict vs. lenient, p = 0.96 for lenient vs. none). “*” indicates statistical significance after multiple comparisons correction with FDR q < 0.05 and “n.s.” indicates not significant after FDR correction. For each box plot, the horizontal line indicates the median across all motion bins. The bottom and top edges of the box indicate the 25th and 75th percentiles, respectively. The outliers are defined as data points beyond 1.5 times the interquartile range. The whiskers extend to the most extreme data points not considered outliers. (C) Discrepancy with respect to the ground-truth parcellation in a representative participant from the highest motion bin. Blue regions indicate agreement with ground-truth parcellation, while red regions indicate differences. (D) Parcellations of the representative participant from panel C: ground-truth parcellation (left), simulated parcellation under strict censoring (middle), and simulated parcellation under lenient censoring (right).

To determine whether a single combination of FD and DVARS thresholds can lead to close-to-optimal result, Figure 9B shows the parcellation quality (Dice) for three censoring regimes: strict censoring (FD = 0.08 mm, DVARS = 50), lenient censoring (FD = 0.5 mm, DVARS = 100), and no censoring. Each dot in the plot represents one of ten motion bins. Both lenient censoring and no censoring exhibited equivalent parcellation quality and were statistically better than strict censoring. The magnitude of this difference was comparable to the improvement in Dice coefficient achieved by doubling scan duration from 20 minutes to 40 minutes under strict censoring without discarding high-motion runs (brown dashed line in Figure 9B).

The gray dashed line in Figure 9B is the “noise ceiling”, achieved by picking the best combination of FD and DVARS threshold for each motion bin. Therefore, lenient censoring yielded parcellations close in quality to the “noise ceiling” across the motion bins examined here. No censoring also achieved parcellation quality close to the “noise ceiling”.

Similar conclusions were obtained with TMS targeting (Figure 10). For most motion bins, TMS targeting quality improved with lower FD threshold. On the other hand, the effect of DVARS thresholds was again less salient. Both lenient censoring and no censoring exhibited equivalent TMS targeting quality, and were statistically better than strict censoring. The magnitude of this difference was comparable to the improvement in Euclidean distance achieved by doubling scan duration from 20 minutes to 40 minutes under strict censoring without discarding high-motion runs (brown dashed line in Figure 10B).

**Figure 10.**
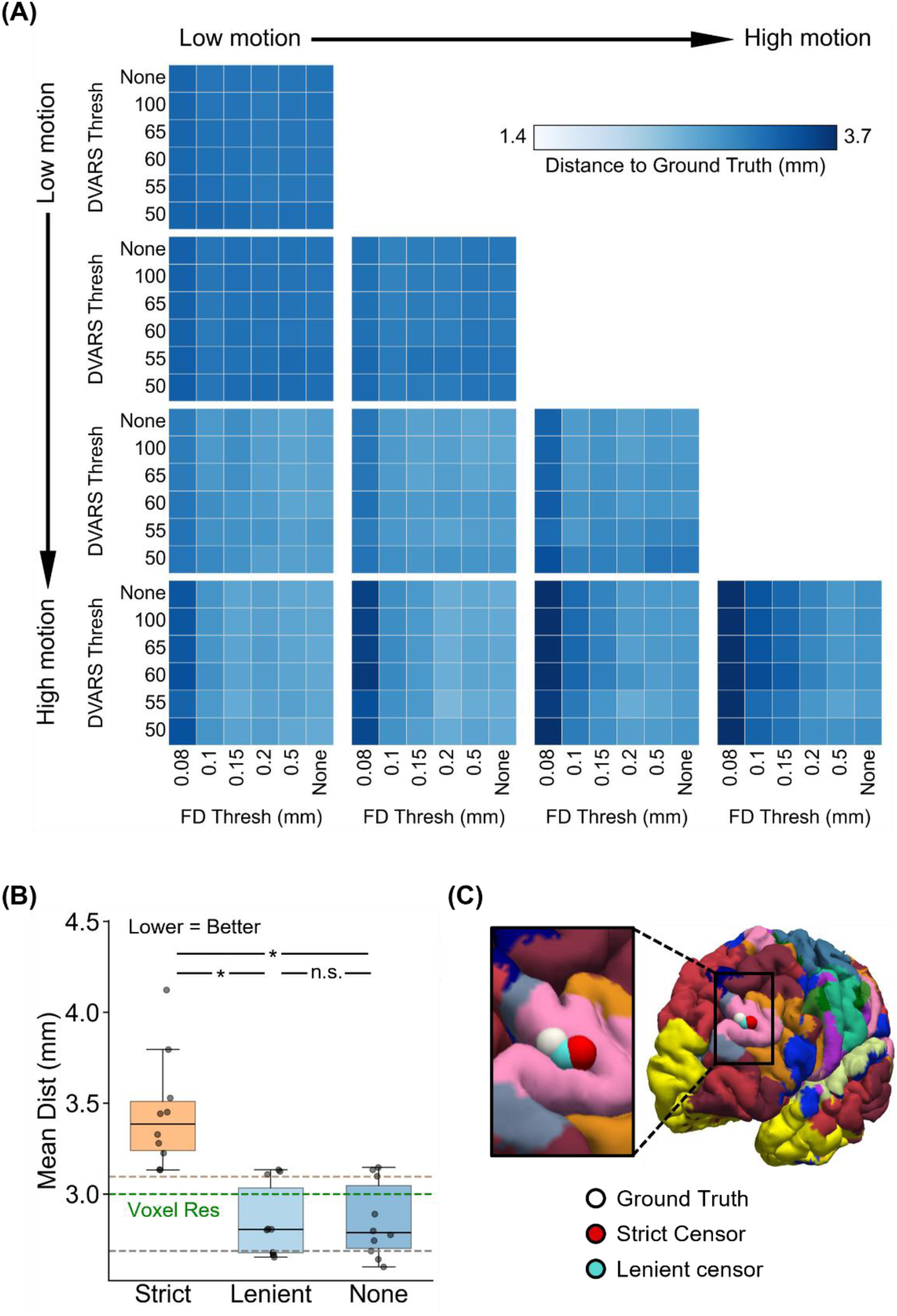
Comparison of TMS depression target quality (Euclidean distance from ground-truth targets) across varying FD and DVARS thresholds for the two-run FD setup. For this analysis, high-motion runs were not discarded. Lower Euclidean distance indicates better quality. TMS depression targets were estimated using the tree-algorithm (Kong et al., 2026a). (A) TMS depression target quality (Euclidean distance) for different combinations of FD and DVARS thresholds for 10 motion bins. Each cell in the grid indicates a motion bin. For each motion bin, there are 36 small cells corresponding to different FD and DVARS thresholds. (B) TMS depression target quality for strict censoring, lenient censoring and no censoring. Here strict censoring refers to FD threshold of 0.08 mm (after pseudo-motion filtering) and DVARS threshold of 50. Lenient censoring refers to FD threshold of 0.5 mm and DVARS threshold of 100. Each dot represents the mean Euclidean distance for a motion bin. The brown dashed line indicates the improvement in Euclidean distance (0.29 mm) achieved by doubling scan duration from 20 minutes to 40 minutes under strict censoring without discarding high-motion runs (Supplementary Figure 10). This improvement (0.29 mm) is subtracted from the median of strict censoring boxplot, resulting in the brown dashed line. The gray dashed line is the “noise ceiling”, achieved by picking the best combination of FD and DVARS threshold for each motion bin, and computing the median across motion bins. P-values were computed using a linear mixed effects model (p = 5.9e-20 for strict vs. none, p = 1.6e-20 for strict vs. lenient, p = 0.89 for lenient vs. none). “*” indicates statistical significance after multiple comparisons correction with FDR q < 0.05 and “n.s.” indicates not significant after FDR correction. For each box plot, the horizontal line indicates the median across all motion bins. The bottom and top edges of the box indicate the 25th and 75th percentiles, respectively. The outliers are defined as data points beyond 1.5 times the interquartile range. The whiskers extend to the most extreme data points not considered outliers. (C) Visualization of TMS depression targets in a representative participant from the highest motion bin.

Lenient censoring yielded TMS targets close in quality to the “noise ceiling” (gray dash line in Figure 10B). Interestingly, the Euclidean distances of lenient censoring TMS targets to ground-truth targets were similar to the median voxel resolution of the fMRI data (green dashed line in Figure 10B).

Overall, this suggests that lenient censoring is appropriate across the motion bins examined here for both cortical parcellations and personalized TMS targeting. Of course, no censoring also achieved parcellation and TMS target quality close to the “noise ceiling”. However, we do not recommend no censoring in case future participants have higher motion than the sample used in the current study (see Discussion).

### 3.5. Supplementary analyses

The previous sections focused on fMRI sessions with two runs. Results from analyzing an alternative one-run sessions were consistent with the previous findings from two-run sessions. In particular, parcellation quality decreased with higher motion and strict censoring (Supplementary Figure 12). No censoring yielded better parcellations than strict censoring (Supplementary Figures 13 and 14). No censoring in high-motion sessions yielded comparable parcellations to strict censoring in low-motion sessions (Supplementary Figure 15). Finally, lenient censoring yielded close-to-optimal parcellations (Supplementary Figure 16).

In the case of TMS depression targeting, similar conclusions were also obtained with one-run fMRI sessions (Supplementary Figures 18 to 22), cone TMS targeting algorithm (Fox et al., 2013; Supplementary Figures 23 to 27) and cluster TMS targeting algorithm (Cash et al., 2021b; Supplementary Figures 28 to 32). Similar conclusions were also obtained with FDrms (instead of FD) for both two-runs (Supplementary Figures 33 to 37) and one-run (Supplementary Figures 38 to 42) scenarios.

For both cortical parcellations and TMS depression targeting, conclusions were consistent when nuisance regression and bandpass filtering were implemented as a single simultaneous regression rather than as sequential processing steps (Supplementary Figures 43–52). For both cortical parcellations and TMS depression targeting, conclusions were also consistent when statistical tests were performed on logit-transformed Dice coefficients and natural-log-transformed Euclidean distances (Supplementary Figures 53–62). Parcellation results were also consistent when parcellation quality was quantified using the mean Hausdorff distance rather than Dice coefficient (Supplementary Figures 63–67).

In analyses using continuous runs rather than simulated runs, both parcellation and TMS depression targeting results were consistent with the main findings (Supplementary Figures 68 and 69).

### 3.6. Lenient censoring yields close-to-optimal TMS anxiety targets

The tree-based algorithm can also be used to derive personalized TMS anxiety targets (Kong et al., 2026a). Under the two-run FD scenario, similar to the TMS depression targets, we found that TMS anxiety target quality decreased with higher motion and strict censoring (Supplementary Figure 70). No censoring yielded better TMS anxiety targets than strict censoring (Supplementary Figures 71 and 72). No censoring in high-motion sessions yields comparable TMS anxiety targets to strict censoring in mixed-motion sessions (Supplementary Figure 73). Finally, lenient censoring yielded close-to-optimal TMS anxiety targets (Supplementary Figure 74). Similar conclusions were obtained with two-run FDrms setup (Supplementary Figures 75 to 79).

## 4. DISCUSSION

In this study, we found that the quality of individual-specific parcellations and personalized TMS targets decreased with higher motion levels and with strict motion censoring. At any given motion level, no censoring yielded better parcellations and TMS targets than strict censoring. No censoring in high-motion sessions produced parcellations and TMS targets that were comparable to, or better than, those obtained by strict censoring and discarding high-motion runs in mixed-motion sessions. Finally, lenient censoring without discarding high-motion runs yielded close-to-optimal parcellations and TMS targets. The magnitude of these censoring effects was comparable to the benefit of doubling scan duration from 20 to 40 min under strict censoring. Overall, within the parcellation and TMS targeting frameworks evaluated here, these findings suggest that lenient censoring better preserves individual-specific information than strict censoring.

### 4.1. Implications for clinical practice in personalized TMS

Beyond research settings, our findings have practical implications for personalized TMS as individualized connectivity-guided targeting becomes increasingly adopted (DeSouza et al., 2025; Hearne et al., 2025). Current clinical protocols have not converged on a single censoring threshold, with reported thresholds ranging from FD = 0.2 mm (Siddiqi et al., 2019a; Conelea et al., 2024) to 0.3 mm (Balderston et al., 2020) and 0.5 mm (Hearne et al., 2025; Burgher et al., 2026), while some protocols specify no threshold (Zhang et al., 2026). In one case study, the censoring threshold was loosened from 0.2 mm to 0.5 mm because the patient’s FD noise floor was substantially higher than typically reported (Siddiqi et al., 2019b). Existing studies and protocols also provide limited guidance on whether high-motion runs should be retained or discarded, and censoring thresholds have generally been selected without empirical evidence regarding their impact on personalized TMS targets. Our study addresses this gap by directly evaluating how censoring thresholds and high-motion run retention affect target quality.

Real-time motion monitoring, mock-scanner acclimatization, and acquisition of additional data are valuable strategies for reducing motion during scanning (Dosenbach et al., 2017). However, these approaches are complementary to the preprocessing question addressed here. In practice, scan time may be limited by fixed scheduling, patient tolerability, cost, and scanner availability. Moreover, even when motion-reduction strategies are used, analysts must still decide how to process already-acquired data when motion remains present.

Consider a patient with a mixed-motion fMRI session: one run exhibits high motion and the other low motion. Under strict censoring practices, the high-motion run may be discarded, and the personalized TMS target would be derived only from the low-motion run after aggressive censoring. Our results suggest that applying lenient censoring while retaining the high-motion run can yield targets that are comparable to, or better than, those obtained after discarding the high-motion run and applying strict censoring.

Similarly, if a patient’s fMRI session contains only high-motion runs, strict censoring may leave insufficient usable data and trigger repeat scanning, incurring additional cost, inconvenience, and delayed treatment for the patient. Within the motion range examined here, our results suggest that processing these high-motion runs with lenient censoring – without discarding them – produces personalized TMS targets that are comparable to, or even better than, those derived from strictly censored low-motion data in the mixed-motion session under strict censoring and discarding of high-motion runs. Accordingly, the patient might not require repeat scanning; rather, the existing data can be effectively leveraged through lenient censoring.

### 4.2. Interpreting TMS target distances

In the current study, TMS target quality was quantified as the Euclidean distance to a within-individual “ground-truth” target estimated from a large quantity (≥1 hour) of low-motion data. This ground-truth target is an operational reference, rather than a proven therapeutic optimum for the individual. For the tree-based MS-HBM algorithm, differences between strict and lenient/no censoring were on the order of 1 mm. These differences are smaller than the average fMRI voxel size across the cohort (∼3.0 mm), smaller than the average voxel size among participants contributing to the highest-motion bins (∼3.6 mm), smaller than the spatial uncertainty of neuronavigated TMS delivery (Nieminen et al., 2022), and far below the approximate centimeter scale over which the TMS electric field remains relatively strong (Deng et al., 2013).

However, TMS-induced direct neural activation can be more focal, on the order of millimeters (Aberra et al., 2020). Consistent with this, single-unit recordings during single-pulse TMS in macaque cortex found that stimulation modulated spiking within a cortical region less than 2 mm in diameter (Romero et al., 2019). This focality is why TMS can distinguish the representations of individual fingers in motor cortex, although this degree of specificity depends on cortical folding and is not attainable everywhere (Numssen et al., 2023). Whether the focal region of direct activation or the broader electric-field exposure is the more relevant scale for therapeutic effects remains unresolved. Nevertheless, the approximately 1-mm differences observed for the tree-based MS-HBM algorithm are unlikely to represent clinically meaningful shifts in stimulation site.

Instead, the clinical relevance of the tree-based MS-HBM results lies primarily in data retention: strict censoring can discard high-motion data and potentially require re-scanning, whereas lenient censoring preserved more data while yielding comparable or even better targets. By contrast, supplementary analyses using the cluster and cone algorithms showed larger strict-versus-lenient target differences in the highest-motion bin, approximately 6.4 mm and 13.5 mm, respectively. These distances exceed the typical voxel size in the highest-motion bin (∼3.6 mm) and approach the scale at which target differences may plausibly matter. As one clinical benchmark, a recent randomized trial found that personalized FC targets produced better antidepressant outcomes than scalp-based Beam-F3 targeting in treatment-resistant depression (Taylor et al., 2026). Personalized FC targets differ from Beam-F3 targets by ∼15 mm (Figure 2D in Kong et al., 2026b). Thus, for some targeting algorithms, censoring choice may affect the selected target at a spatial scale that could plausibly be clinically relevant. Ultimately, however, whether censoring-related differences in target location translate into differences in therapeutic response will need to be tested in prospective clinical studies.

### 4.3. Session variability and broader precision neuroimaging applications

A related issue is whether single-session rs-fMRI can yield clinically meaningful targets despite session-to-session variability in FC. Although FC estimates vary across sessions, inter-individual differences in functional network organization generally dominate across-session variability (Gratton et al., 2018). In personalized TMS, prior studies have similarly found that within-individual variability in FC-derived target location is generally smaller than between-individual variability in targets (Cash et al., 2021b; Kong et al., 2026a). Most notably, a recent randomized clinical trial found that individualized FC targets derived from single-session rs-fMRI produced greater antidepressant effects than scalp-based Beam-F3 targeting (Taylor et al., 2026). Thus, despite session-to-session variability, individualized connectivity-based targets derived from single-session rs-fMRI can be clinically meaningful.

For presurgical functional mapping, intraoperative electrocortical stimulation remains the clinical gold standard, while task-fMRI and rs-fMRI can provide non-invasive adjunctive information during preoperative planning (Mitchell et al., 2013; Kumar et al., 2024). Current consensus recommendations for presurgical rs-fMRI specify a minimum acquisition duration of 6 min for mapping motor, language, and visual areas (Kumar et al., 2024), but this should be viewed as a pragmatic minimum rather than evidence that such short acquisitions are optimal for individualized functional localization. Indeed, the broader precision-fMRI literature shows that longer scan durations improve the reliability and predictive utility of individual-level FC estimates (Laumann et al., 2015; Gordon et al., 2017; Gratton et al., 2018; Ooi et al., 2025). Thus, very short acquisitions (e.g., 6 min) may yield unstable individual-level maps, which could limit the clinical utility of rs-fMRI for high-stakes neurosurgical applications. Although we did not explicitly evaluate presurgical functional mapping, our findings suggest that strict censoring might further reduce the stability of individualized maps by discarding usable data. Whether lenient censoring improves presurgical rs-fMRI mapping should be tested directly in future studies.

### 4.4. Reconciling with literature advocating strict censoring

How can our findings be reconciled with prior studies arguing that strict censoring is necessary to remove motion-related artifacts (Power et al., 2014; Gratton et al., 2020; Weiler et al., 2022)? One key distinction is that many earlier studies aim to eliminate across-individual correlations between functional connectivity (FC) and head motion measures, commonly referred to as QC–FC correlations (Power et al., 2014; Ciric et al., 2017; Parkes et al., 2018; Li et al., 2019). However, across-individual QC–FC correlations do not distinguish between trait-level and state-level contributions of motion to FC. Notably, previous work has shown that high-motion individuals exhibit systematic FC differences relative to low-motion individuals even during fMRI sessions in which their motion is low (Zeng et al., 2014). Therefore, standard QC-FC may partially reflect participants’ tendency to move, rather than residual motion artifacts per se.

Consistent with this interpretation, a recent study decomposed standard QC–FC into between-individual and within-individual components using repeated-measures data from the Human Connectome Project (Mejia et al., 2026). The within-individual component might capture FC related motion state and motion artifact, while the between-individual component might capture stable individual differences in FC associated with individuals’ tendency to move. The study found that standard QC–FC was dominated by the between-individual component, suggesting that a significant portion of the signal targeted by strict censoring reflects stable individual-level differences rather than transient motion artifact. Importantly, censoring reduced the within-individual component, consistent with removal of motion artifact, but also reduced the between-individual component, suggesting removal of stable individual-level FC information (Mejia et al., 2026).

This distinction helps explain why strict censoring can reduce standard QC–FC while degrading individual-specific estimates. When the goal is to estimate an individual’s cortical parcellation or personalized TMS target, aggressive censoring may discard data that contribute to that individual’s stable FC profile, thereby worsening agreement with the within-individual ground-truth. Our findings should therefore not be interpreted as contradicting the existence of motion artifacts, but rather as showing that minimizing standard QC–FC is not necessarily the same as optimizing individual-specific FC estimates for precision neuroimaging applications.

### 4.5. Scope of generalization

An important caveat is that our study was conducted in healthy adult participants undergoing precision fMRI, who typically exhibit relatively low motion on average. Nevertheless, the extended scan duration allowed us to simulate fMRI runs with comparatively high motion. Across our simulations, maximum frame-level FD (after pseudo-motion filtering) was 1.14 mm, maximum frame-level FDrms was 1.19 mm, and maximum frame-level DVARS was 133.4 (Supplementary Figures 80–83). On the other hand, when averaging motion metrics (FD, FDrms and DVARS) across the whole run, the maximum run-level FD (after pseudo-motion filtering), FDrms and DVARS were 0.16 mm, 0.24 mm and 46.3 respectively (Supplementary Figures 80–83). Accordingly, our findings should be applied with caution to participants whose motion exceeds these observed ranges. For this reason, we advocate lenient censoring over no censoring, because participant motion may surpass the maximum levels present in our data.

Another consideration is that our parcellation analyses used the 17-network MS-HBM model. MS-HBM combines individual-specific functional connectivity with group-level priors within a Bayesian framework, allowing stable individualized parcellations to be estimated from relatively modest scan durations (Kong et al., 2019). This regularization is a strength of the method, rather than a limitation: in held-out data, MS-HBM parcellations exhibit better resting-state functional homogeneity and task-state functional inhomogeneity than alternative approaches across scan durations ranging from 10 to 50 min (Kong et al., 2019). We used the 17-network MS-HBM model because it is the parcellation model used by the tree-based personalized TMS targeting algorithm evaluated here (Kong et al., 2026a).

Nevertheless, because MS-HBM explicitly balances individual-specific data with population-informed priors, strict censoring could in principle affect parcellation quality by reducing the amount of usable individual-specific data and thereby altering the relative influence of the observed FC data and the group-level prior. We therefore scope our parcellation findings to the 17-network MS-HBM model evaluated here, and whether similar results would be obtained with finer-grained or non-MS-HBM parcellation methods remains an important question for future work.

However, the broader evidence suggests that the detrimental effect of aggressive censoring is unlikely to be unique to MS-HBM. First, Mejia et al. (2026) showed that aggressive censoring increases FC estimation error relative to a within-individual reference, indicating that the loss of individual-specific information occurs at the level of FC estimation itself, before any parcellation algorithm is applied. Second, our supplementary TMS analyses using the cone (Supplementary Figures 23–27) and cluster (Supplementary Figures 28–32) algorithms yielded conclusions consistent with the main TMS analyses, even though these algorithms do not rely on MS-HBM parcellations, Bayesian group-level priors, or the tree-based targeting framework.

Finally, the MS-HBM parcellation analyses required projecting functional data to fsaverage6, where the pretrained group-level priors are defined, before mapping individual-specific parcellations back to native space for TMS targeting (Kong et al., 2019; Kong et al., 2026a). This projection may introduce some smoothing or loss of individual-specific spatial detail. However, this processing step was applied identically to the ground-truth and simulated-session estimates, so it is unlikely to explain the observed differences between censoring strategies. Thus, while the parcellation findings should be interpreted within the scope of the 17-network MS-HBM model, the TMS targeting findings are not an artifact of a single recent algorithmic choice, and the broader evidence suggests that aggressive censoring can degrade individual-specific FC estimates in a way that is not specific to MS-HBM.

### 4.6. Additional preprocessing and simulation considerations

Another limitation is that we focused on censoring decisions within a fixed nuisance-regression and global-signal-regression preprocessing framework. This framework is of particular relevance because most personalized connectome-guided TMS studies have used nuisance regression with GSR rather than ICA-FIX (Fox et al., 2013; Cole et al., 2020; Cash et al., 2021a; Siddiqi et al., 2021; Taylor et al., 2026). ICA-based denoising approaches, including FIX and related component-based methods, provide an important alternative to framewise censoring because they can partition variance within frames rather than discarding entire frames (Griffanti et al., 2014; Salimi-Khorshidi et al., 2014; Glasser et al., 2018). Directly comparing these approaches is beyond the scope of the present study because our evaluation depends on an operational ground truth derived under a specific preprocessing pipeline. Evaluating ICA-FIX against a nuisance-regression-derived ground truth would partly test whether ICA-FIX reproduces the output of a different denoising pipeline, whereas constructing separate pipeline-specific ground truths would not provide a common reference for directly comparing ICA-FIX with nuisance regression. We leave such comparisons to future work.

A final consideration is that our simulated-run paradigm concatenated 1-min windows sampled from different real runs, which may reduce temporal autocorrelation relative to continuous acquisitions and thereby increase the stability of FC estimates. However, because all censoring strategies were applied to the same simulated runs within each individual, any reduction in autocorrelation should affect all censoring conditions similarly. Moreover, a control analysis using continuous 10-min runs yielded results consistent with the main analyses: lenient censoring and no censoring produced parcellations and TMS depression targets that were comparable to, or better than, those obtained with strict censoring (Supplementary Figures 68 and 69). Thus, while the simulated-run paradigm may produce more stable estimates than continuous runs of the same scan duration, it is unlikely to explain the relative advantage of lenient censoring observed in this study.

## 5. CONCLUSION

In this study, we leveraged precision-fMRI datasets to construct individualized “ground-truths” for cortical parcellations and personalized TMS targets. We found that lenient censoring and retaining high-motion runs produced cortical parcellations and brain stimulation targets with comparable or higher quality than those derived from stringent censoring and exclusion of high-motion runs. Overall, these findings provide practical guidance for optimizing personalized cortical parcellations and TMS targets in both research and clinical settings.

## DATA & CODE AVAILABILITY STATEMENT

This study utilized publicly available data from HBNSSI (https://fcon_1000.projects.nitrc.org/indi/hbn_ssi/), IPCAS6 (https://fcon_1000.projects.nitrc.org/indi/CoRR/html/ipcas_6.html), Kirby (https://www.nitrc.org/projects/kirbyweekly), MSC (https://openneuro.org/datasets/ds000224/versions/1.0.4), MyConnectome (https://openneuro.org/datasets/ds000031/versions/2.0.2), NSD (https://naturalscenesdataset.org/), Newbold (https://openneuro.org/datasets/ds002766/versions/3.0.0), and YaleTestRetest (https://fcon_1000.projects.nitrc.org/indi/retro/yale_trt.html). Data can be accessed via data use agreements. Preprocessing code is based on the CBIG preprocessing pipeline: https://github.com/ThomasYeoLab/CBIG/tree/master/stable_projects/preprocessing/CBIG_fMRI_Preproc2016. Analysis code specific to this study can be found on GitHub: GITHUB_LINK.

## AUTHOR CONTRIBUTIONS

TWKT, RK, BTTY conceptualized the study and designed the methodology. TWKT carried out the analyses. TWKT, TEN, BTTY designed and/or executed statistical analyses across datasets. RK, AX, JC reviewed the code used in the study. TWKT and BTTY wrote the manuscript and created the figures. All authors contributed to project direction via discussion. All authors edited the manuscript.

## ACKNOWLEDGEMENTS

Our research is supported by the NUS Yong Loo Lin School of Medicine (NUHSRO/2020/124/TMR/LOA), the Singapore National Medical Research Council (NMRC) LCG (OFLCG19May-0035), NMRC CTG-IIT (CTGIIT23jan-0001), NMRC OF-IRG (OFIRG24jan-0006; OFIRG24jul-0049), NMRC STaR (STaR20nov-0003), Singapore Ministry of Health (MOH) Centre Grant (CG21APR1009), the United States National Institutes of Health (R01MH133334, 2R01MH120080 & R01AG083919) and the Singapore National Research Foundation (NRF) Investigatorship (NRFI10-2024-0014). LC is supported by the Australian NHMRC (grant 2027597). Any opinions, findings and conclusions or recommendations expressed in this material are those of the authors and do not reflect the views of the funders. Data for the MyConnectome project were obtained from the OpenNeuro database (ds000031). Data for the MSC dataset was obtained from the OpenNeuro database. Its accession number is ds000224. Collection of the NSD dataset was supported by NSF IIS-1822683 and NSF IIS-1822929.

