## Supplementary Material for "Excessive Censoring Degrades Individual-Specific Cortical Parcellations and Personalized TMS Targets"

### Supplementary Results

#### **Methods section**

| **DS** | **PID** | **Age** | **Sex** | **GI** | **Res.** | **Race/Cult.** | **Edu.** | **Fin.** | **Wrk.** | **Excl.** | **2FD** | **1FD** | **2FDrms** | **1FDrms** |
| --- | --- | --- | --- | --- | --- | --- | --- | --- | --- | --- | --- | --- | --- | --- |
| HBNSSI | sub-0031121 | 24 | M | - | - | - | - | - | - | NA | NA | ✓ | NA | ✓ |
| HBNSSI | sub-0031122 | 33 | F | - | - | - | - | - | - | IGT | NA | NA | NA | NA |
| HBNSSI | sub-0031123 | 27 | M | - | - | - | - | - | - | NA | ✓ | ✓ | NA | NA |
| HBNSSI | sub-0031124 | 21 | F | - | - | - | - | - | - | IGT | NA | NA | NA | NA |
| HBNSSI | sub-0031125 | 25 | M | - | - | - | - | - | - | NA | ✓ | ✓ | ✓ | ✓ |
| HBNSSI | sub-0031126 | 37 | F | - | - | - | - | - | - | NA | ✓ | ✓ | ✓ | ✓ |
| HBNSSI | sub-0031127 | 26 | M | - | - | - | - | - | - | NA | ✓ | ✓ | ✓ | ✓ |
| HBNSSI | sub-0031128 | 33 | F | - | - | - | - | - | - | IGT | NA | NA | NA | NA |
| HBNSSI | sub-0031129 | 42 | M | - | - | - | - | - | - | IGT | NA | NA | NA | NA |
| HBNSSI | sub-0031130 | 33 | F | - | - | - | - | - | - | NA | ✓ | ✓ | ✓ | ✓ |
| HBNSSI | sub-0031131 | 36 | F | - | - | - | - | - | - | IGT | NA | NA | NA | NA |
| HBNSSI | sub-0031132 | 34 | F | - | - | - | - | - | - | IGT | NA | NA | NA | NA |
| HBNSSI | sub-0031133 | 23 | M | - | - | - | - | - | - | IGT | NA | NA | NA | NA |
| IPCAS6 | sub-0026044 | 25 | M | - | - | - | - | - | - | NA | ✓ | ✓ | ✓ | ✓ |
| IPCAS6 | sub-0026045 | 21 | F | - | - | - | - | - | - | IGT | NA | NA | NA | NA |
| Kirby | sub-01 | 40 | M | - | - | - | - | - | - | NA | ✓ | ✓ | ✓ | ✓ |
| MSC | sub-MSC01 | 34 | M | - | - | - | Doc. | - | - | NA | ✓ | ✓ | ✓ | ✓ |
| MSC | sub-MSC02 | 34 | M | - | - | - | Doc. | - | - | NA | ✓ | ✓ | ✓ | ✓ |
| MSC | sub-MSC03 | 29 | F | - | - | - | Mast. | - | - | NA | ✓ | ✓ | ✓ | ✓ |
| MSC | sub-MSC04 | 28 | F | - | - | - | Bach. | - | - | IGT | NA | NA | NA | NA |
| MSC | sub-MSC05 | 27 | M | - | - | - | Bach. | - | - | NA | ✓ | ✓ | ✓ | ✓ |
| MSC | sub-MSC06 | 24 | F | - | - | - | Bach. | - | - | IGT | NA | NA | NA | NA |
| MSC | sub-MSC07 | 31 | F | - | - | - | Mast. | - | - | NA | ✓ | ✓ | ✓ | ✓ |
| MSC | sub-MSC08 | 27 | F | - | - | - | Prof. | - | - | Drow. | NA | NA | NA | NA |
| MSC | sub-MSC09 | 26 | M | - | - | - | Prof. | - | - | NA | ✓ | ✓ | ✓ | ✓ |
| MSC | sub-MSC10 | 31 | M | - | - | - | Prof. | - | - | NA | ✓ | ✓ | ✓ | ✓ |
| MC | sub-01 | 46 | M | - | - | Cauc. | - | - | - | NA | ✓ | ✓ | NA | NA |
| NB | sub-cast1 | 35 | M | - | - | - | - | - | - | NA | ✓ | ✓ | NA | NA |
| NB | sub-cast2 | 25 | F | - | - | - | - | - | - | NA | ✓ | ✓ | ✓ | ✓ |
| NB | sub-cast3 | 27 | M | - | - | - | - | - | - | NA | ✓ | ✓ | ✓ | ✓ |
| NSD | sub-01 | 30 | M | - | - | - | - | - | - | IGT | NA | NA | NA | NA |
| NSD | sub-02 | 28 | F | - | - | - | - | - | - | NA | NA | ✓ | NA | NA |
| NSD | sub-03 | 29 | F | - | - | - | - | - | - | NA | NA | ✓ | NA | ✓ |
| NSD | sub-04 | 27 | F | - | - | - | - | - | - | IGT | NA | NA | NA | NA |
| NSD | sub-05 | 32 | F | - | - | - | - | - | - | NA | ✓ | ✓ | ✓ | ✓ |
| NSD | sub-06 | 23 | M | - | - | - | - | - | - | IGT | NA | NA | NA | NA |
| NSD | sub-07 | 24 | F | - | - | - | - | - | - | IGT | NA | NA | NA | NA |
| NSD | sub-08 | 19 | F | - | - | - | - | - | - | IGT | NA | NA | NA | NA |
| YTR | sub-032401 | 28 | M | - | - | - | - | - | - | IGT | NA | NA | NA | NA |
| YTR | sub-032402 | 52 | F | - | - | - | - | - | - | IGT | NA | NA | NA | NA |
| YTR | sub-032403 | 33 | F | - | - | - | - | - | - | IGT | NA | NA | NA | NA |
| YTR | sub-032404 | 54 | M | - | - | - | - | - | - | IGT | NA | NA | NA | NA |
| YTR | sub-032405 | 56 | F | - | - | - | - | - | - | IGT | NA | NA | NA | NA |
| YTR | sub-032406 | 41 | F | - | - | - | - | - | - | IGT | NA | NA | NA | NA |
| YTR | sub-032407 | 53 | F | - | - | - | - | - | - | IGT | NA | NA | NA | NA |
| YTR | sub-032408 | 35 | M | - | - | - | - | - | - | IGT | NA | NA | NA | NA |
| YTR | sub-032409 | 38 | F | - | - | - | - | - | - | IGT | NA | NA | NA | NA |
| YTR | sub-032410 | 27 | M | - | - | - | - | - | - | NA | NA | NA | ✓ | ✓ |
| YTR | sub-032411 | 38 | M | - | - | - | - | - | - | IGT | NA | NA | NA | NA |
| YTR | sub-032412 | 28 | M | - | - | - | - | - | - | IGT | NA | NA | NA | NA |

##### **Supplementary Table 1.** Participant characteristics and inclusion status for the 50 densely sampled participants considered for this study. Demographics are reported following the Identifying Social factors that Stratify Health Opportunities and Outcomes (ISSHOOs) Set A reporting recommendations (Karran et al., 2025), to the extent permitted by the source datasets. Key: DS=dataset; PID=participant ID; GI=gender identity; Res.=place of residence; Race/Cult.=race, ethnicity and/or cultural identity; Edu.=education; Fin.=financial position; Wrk.=work status; Excl.=exclusion reason. '-'=not reported; NA=not applicable; IGT=insufficient data for ground-truth; Drow.=drowsiness; Doc.=Doctorate; Mast.=Master's; Bach.=Bachelor's; Prof.=Professional; MC=MyConnectome; NB=Newbold; YTR=YaleTestRetest; 2FD=two runs FD setup; 1FD=one run FD setup; 2FDrms=two runs FDrms setup; 1FDrms= one run FDrms setup. '✓'=contributed to setup. The exclusion reason column indicates why a participant was excluded from all analyses; "NA" indicates participants included in at least one analysis setup. Because the FD and FDrms analyses retain partially overlapping but non-identical participants (owing to differing motion distributions under the two metrics), 23 participants are marked "NA" in total, though no single analysis uses all 23. The two-run FD analyses used 19 participants, the one-run FD analyses used 22, the two-run FDrms analyses used 17, and the one-run FDrms analyses used 19.

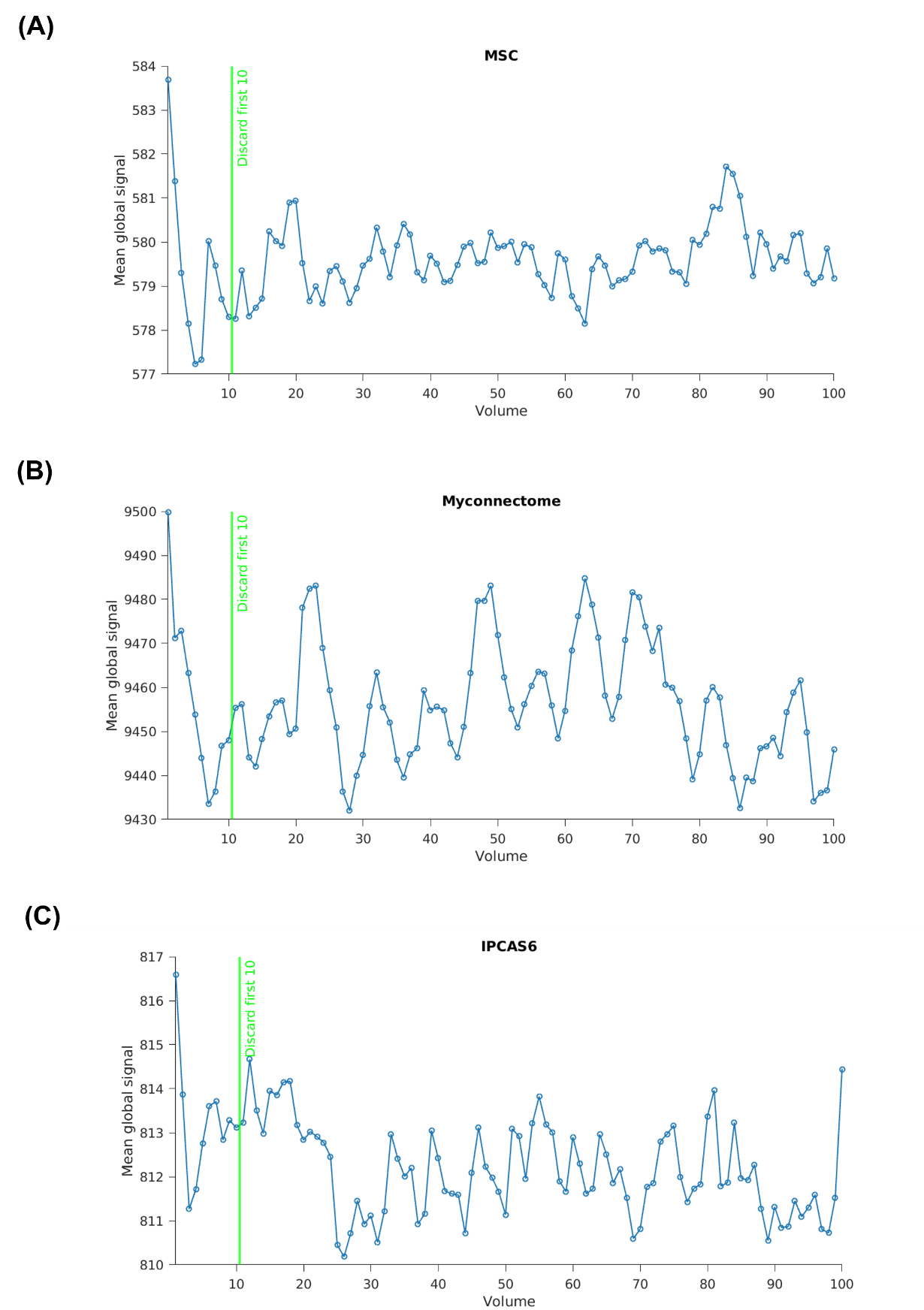

##### **Supplementary Figure 1.** Global signal of a representative single run from a single participant in each of the (A) MSC, (B) MyConnectome, and (C) IPCAS6 datasets. Each circle represents the global signal of a single volume, computed as the average BOLD signal across all in-brain voxels before any preprocessing. The green vertical line marks the cutoff at volume 10 used in this study. Volumes before this line were discarded.

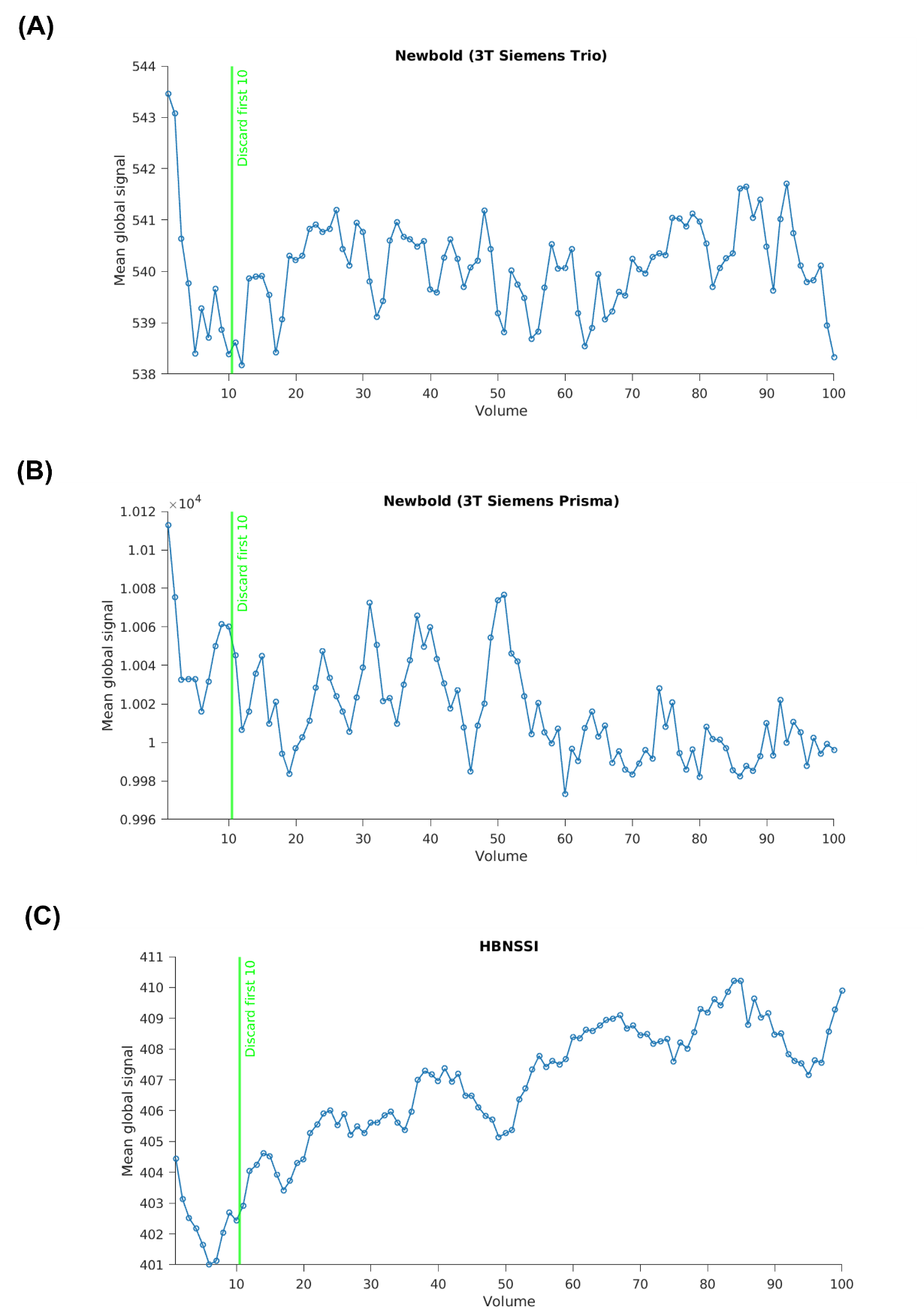

##### **Supplementary Figure 2.** Global signal of a representative single run from a single participant in each of the (A) Newbold (3T Siemens Trio scanner), (B) Newbold (3T Siemens Prisma scanner), and (C) HBNSSI datasets. Each circle represents the global signal of a single volume, computed as the average BOLD signal across all in-brain voxels before any preprocessing. The green vertical line marks the cutoff at volume 10 used in this study. Volumes before this line were discarded.

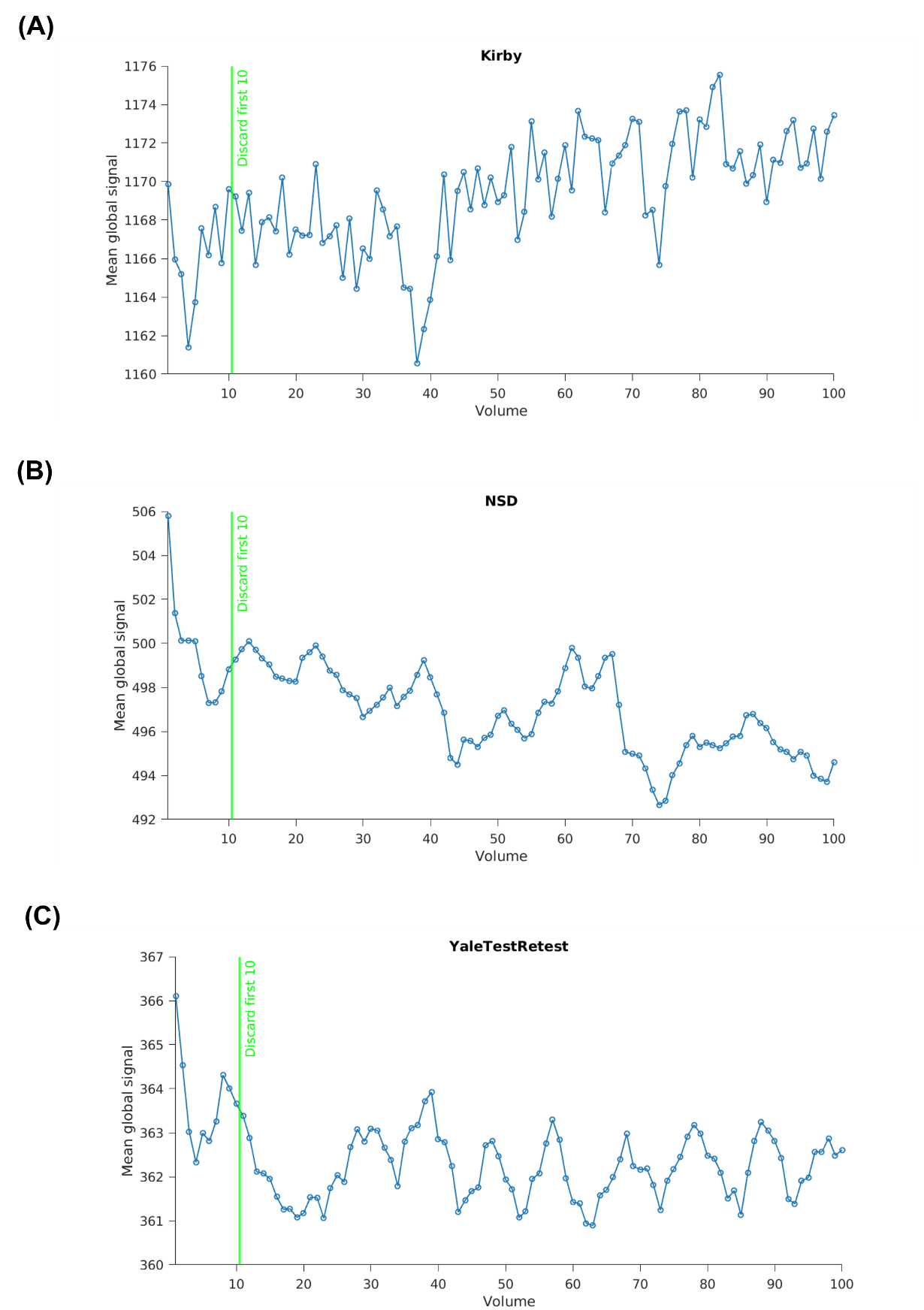

##### **Supplementary Figure 3.** Global signal of a representative single run from a single participant in each of the (A) Kirby, (B) NSD, and (C) YaleTestRetest datasets. Each circle represents the global signal of a single volume, computed as the average BOLD signal across all in-brain voxels before any preprocessing. The green vertical line marks the cutoff at volume 10 used in this study. Volumes before this line were discarded.

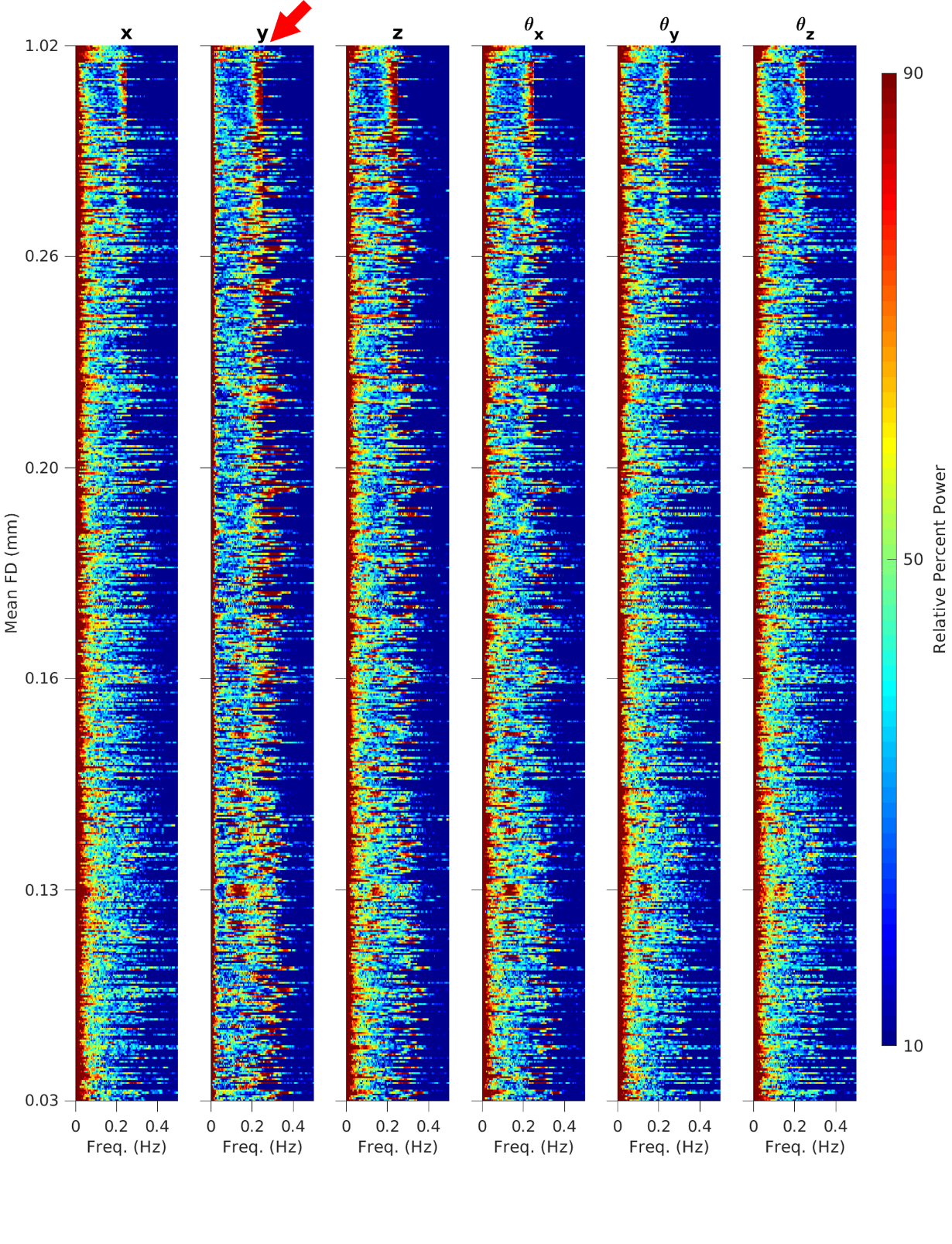

##### **Supplementary Figure 4.** Power spectra of motion parameters across all runs from all participants in all datasets, ordered by mean framewise displacement (FD) with the lowest-motion runs at the bottom. Each column corresponds to one of the six motion parameters: three translations (x, y, z) and three rotations ($\theta_{x},\theta_{y},\theta_{z}$). Within each column, each row is a single run, and the colormap indicates the percentile of Z-scored log-power across runs, as a function of frequency. To allow for visual comparisons across a wide range of motion time-series power, the spectra were first computed using multitaper spectral estimation, represented logarithmically (i.e., in dB), and then Z-scored across frequencies within each run, following Fair et al. (2020). The red arrow on the y-translation column highlights the characteristic frequency of the respiratory artifact (~0.2–0.33 Hz).

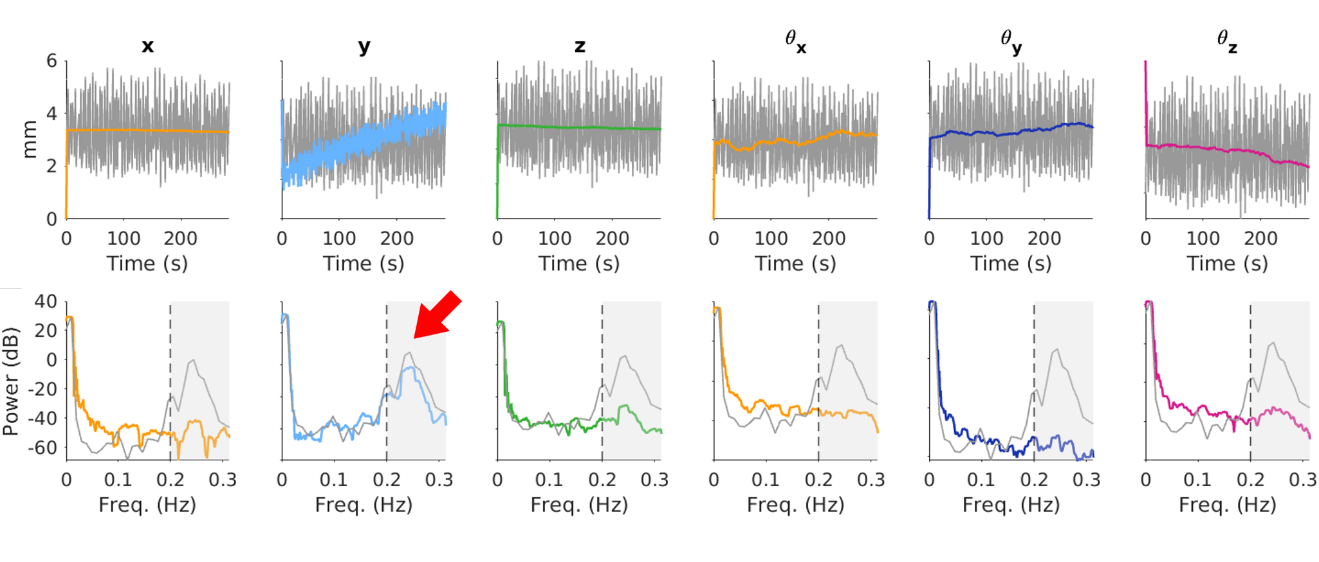

##### **Supplementary Figure 5.** Motion parameter traces and power spectra from a representative single run of a single participant in the NSD dataset (the only dataset in this study with simultaneously acquired respiratory belt recordings), following Fair et al. (2020). Each column corresponds to one of the six motion parameters: three translations (x, y, z) and three rotations ($\theta_{x},\theta_{y},\theta_{z}$). (Top row) Full motion (color) and respiratory (gray) traces across the entire run. (Bottom row) Power spectra of the motion (color) and respiratory (gray) belt traces. The shaded region (0.2–0.33 Hz) indicates the bandstop range applied to the motion traces in this study. The red arrow on the y-translation column highlights the characteristic frequency of the respiratory artifact (~0.2–0.33 Hz); observe strong correspondence between the power spectra of motion and respiratory traces in this frequency range.

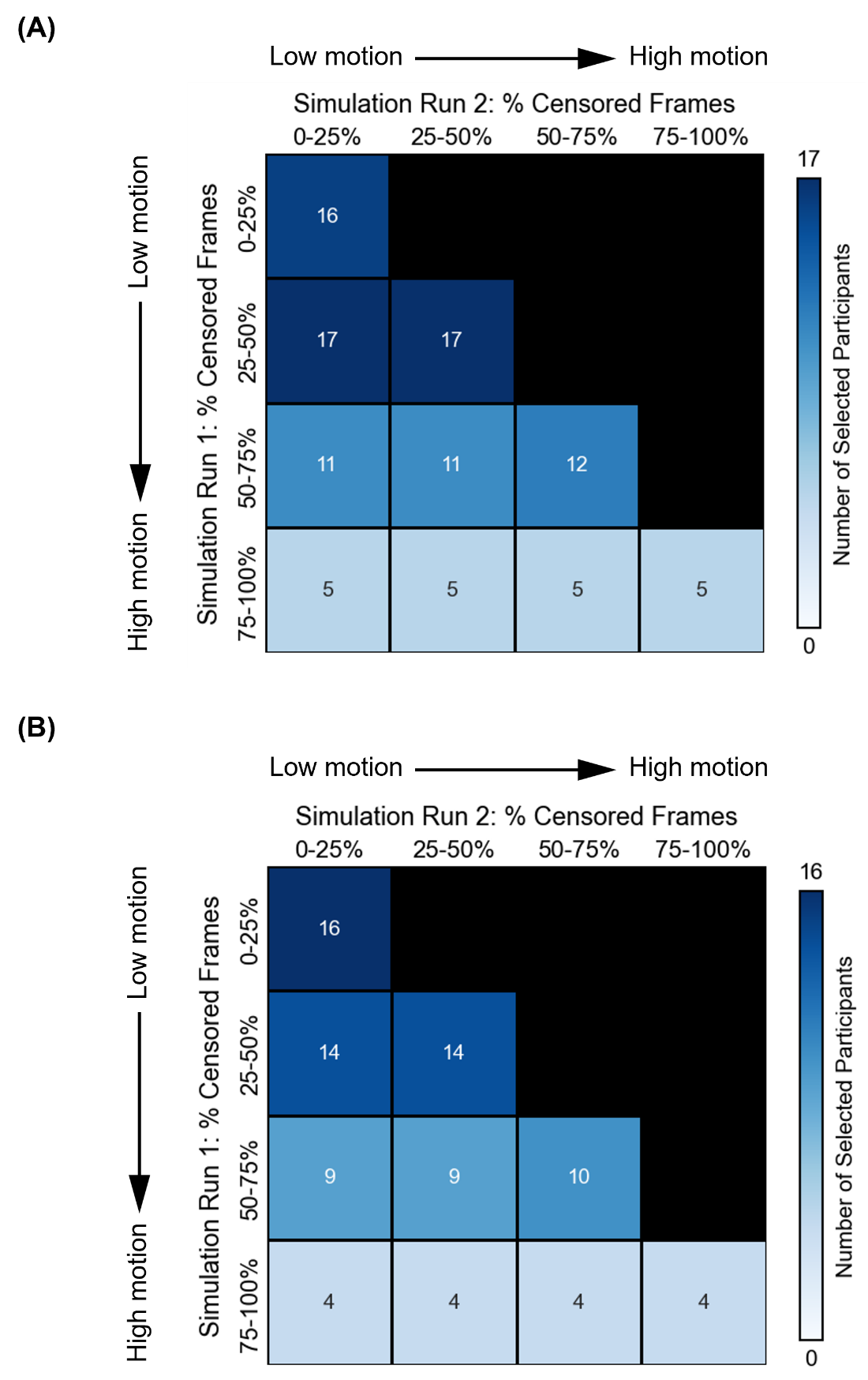

##### **Supplementary Figure 6.** Participant distribution across motion bins for the two-run FD and FDrms setups. (A) Distribution for the two-run FD setup, comprising 19 unique participants across all motion bins. (B) Distribution for the two-run FDrms setup, comprising 17 unique participants across all motion bins. The number within each bin and the intensity of the blue shading indicate the number of participants.

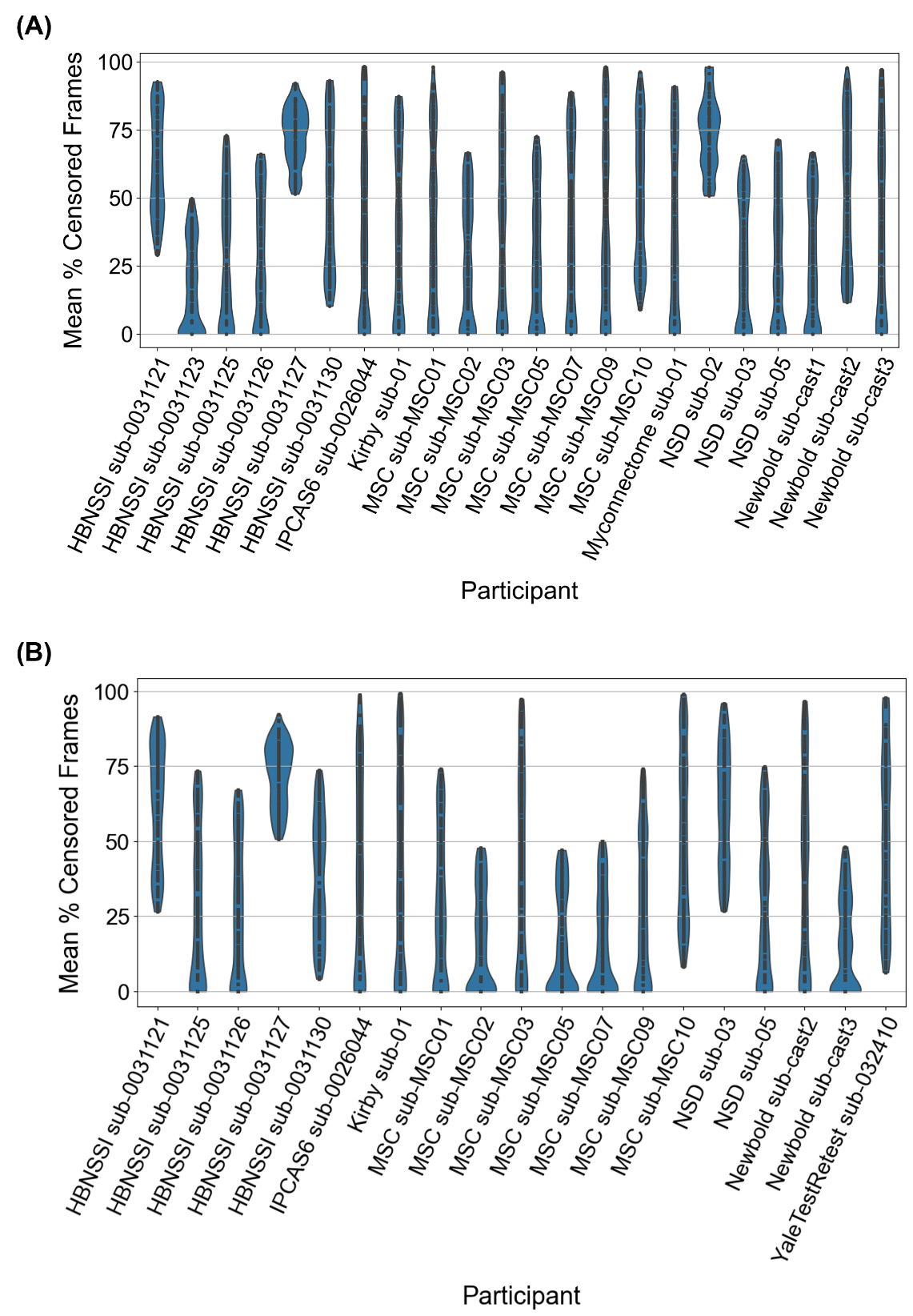

##### **Supplementary Figure 7.** Participant distribution across motion bins for the one-run FD and FDrms setups. (A) Distribution for the one-run FD setup, comprising 22 unique participants across all motion bins. (B) Distribution for the one-run FDrms setup, comprising 19 unique participants across all motion bins. The y-axis is divided into four intervals, with each interval representing a motion bin. Each violin indicates a participant. Each dot in each violin plot represents a simulated run.

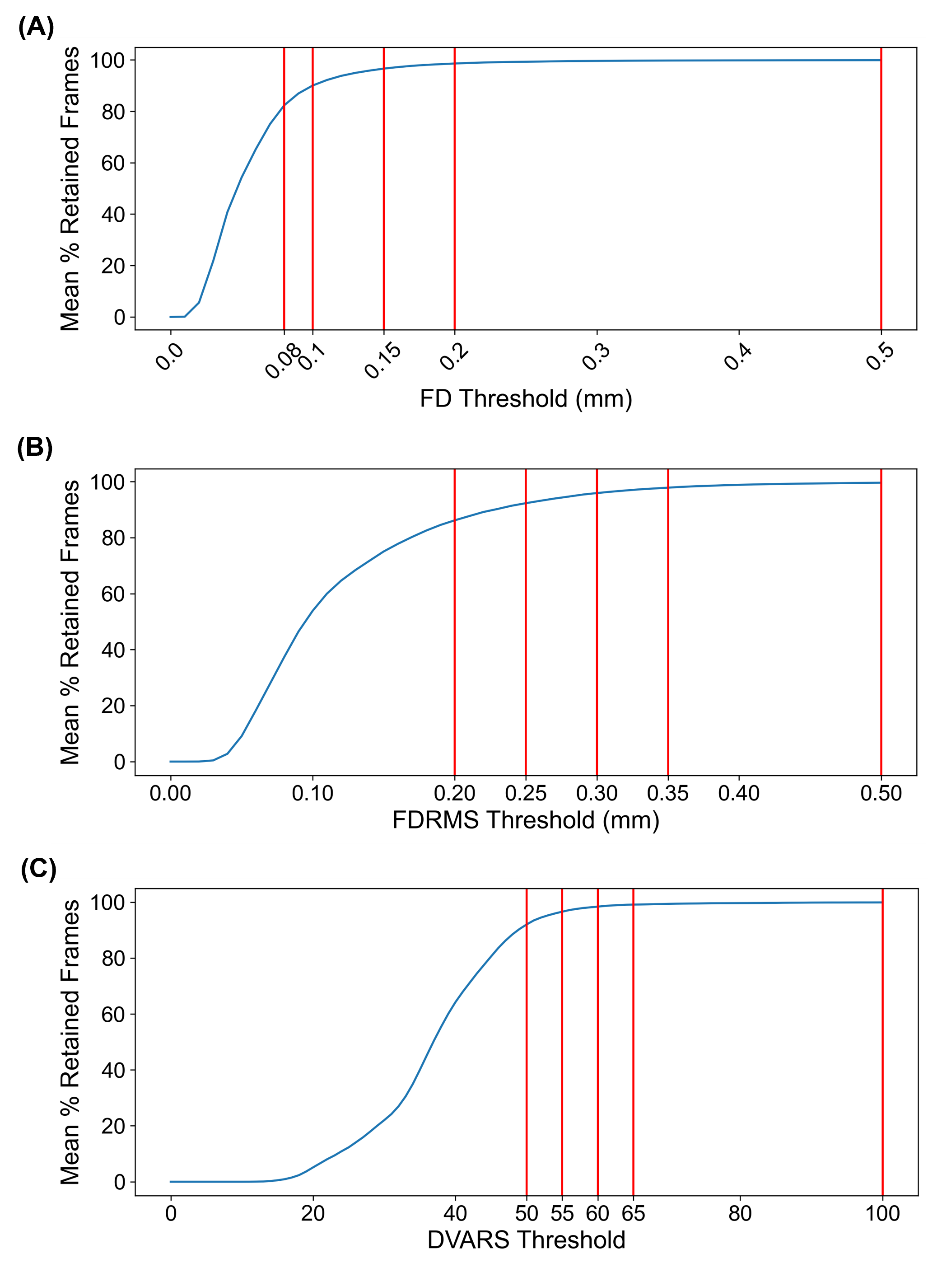

##### **Supplementary Figure 8.** Effect of varying (A) FD, (B) FDrms, and (C) DVARS thresholds on the mean percentage of retained frames in the original fMRI runs. Red vertical lines denote the thresholds explored in the current study.

#### **20 minutes vs. 40 minutes benchmark FD setup for parcellations**

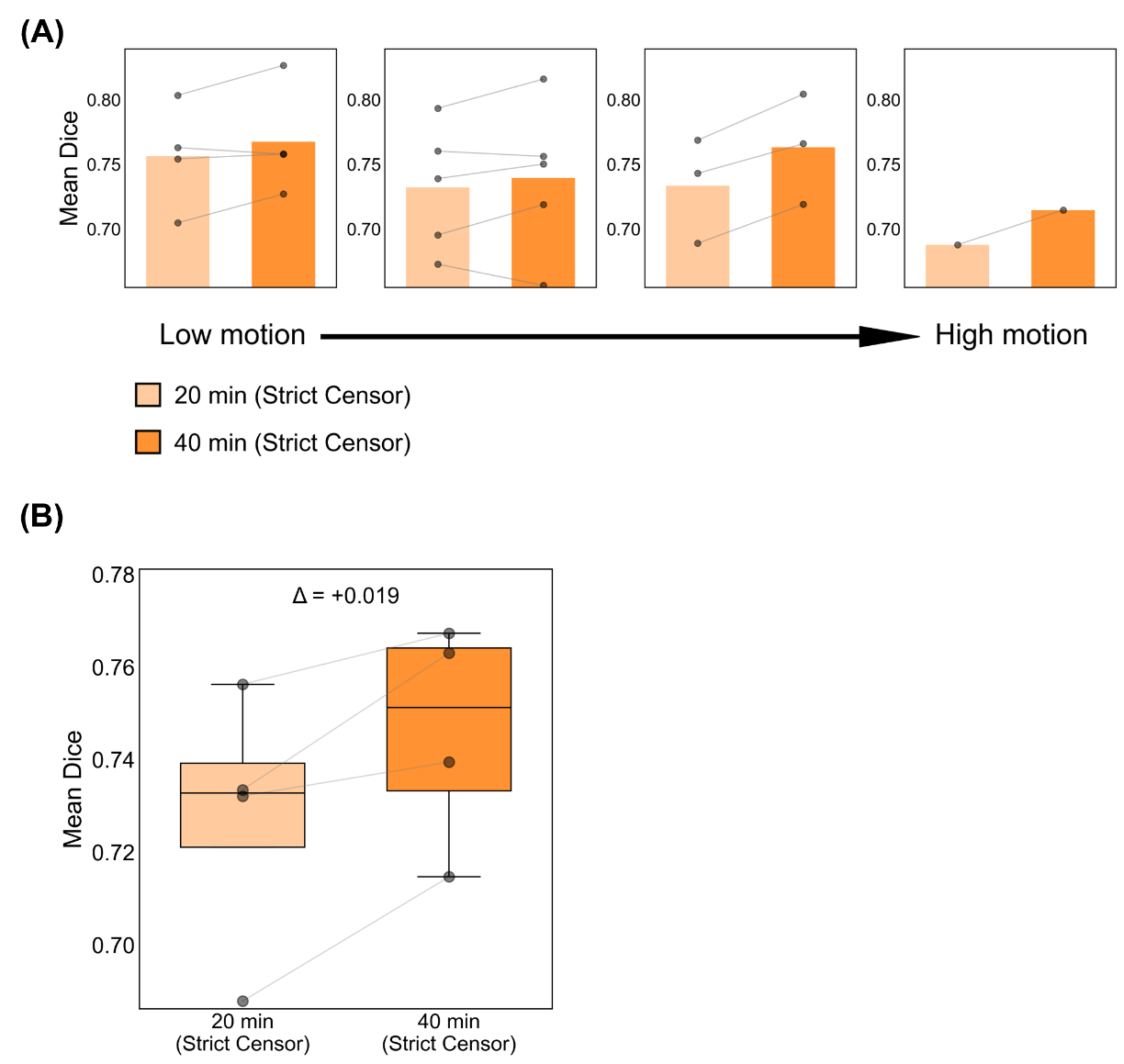

##### **Supplementary Figure 9.** Comparison of parcellation quality (Dice) between 20 minutes and 40 minutes of rs-fMRI data per session under strict censoring for the FD setup. Here, strict censoring refers to a FD threshold of 0.08 mm (after pseudo-motion filtering) and a DVARS threshold of 50. For this analysis, high-motion runs were not discarded. Higher Dice indicates better quality. (A) Parcellation quality for the 20-minute and 40-minute conditions across four motion bins. Each cell indicates a motion bin. Each dot represents the mean Dice coefficient across 50 simulated sessions of a participant in that motion bin. (B) Parcellation quality across all four motion bins for both durations. There are four pairs of dots, corresponding to the four motion bins where comparisons could be made between the two durations. Δ indicates the mean improvement in Dice from the 20-minute to the 40-minute condition across the four motion bins. For each box plot, the horizontal line indicates the median across motion bins. The bottom and top edges of the box indicate the 25th and 75th percentiles, respectively. The outliers are defined as data points beyond 1.5 times the interquartile range. The whiskers extend to the most extreme data points not considered outliers.

#### **20 minutes vs. 40 minutes benchmark FD setup for tree-based (Kong2026) TMS depression targets**

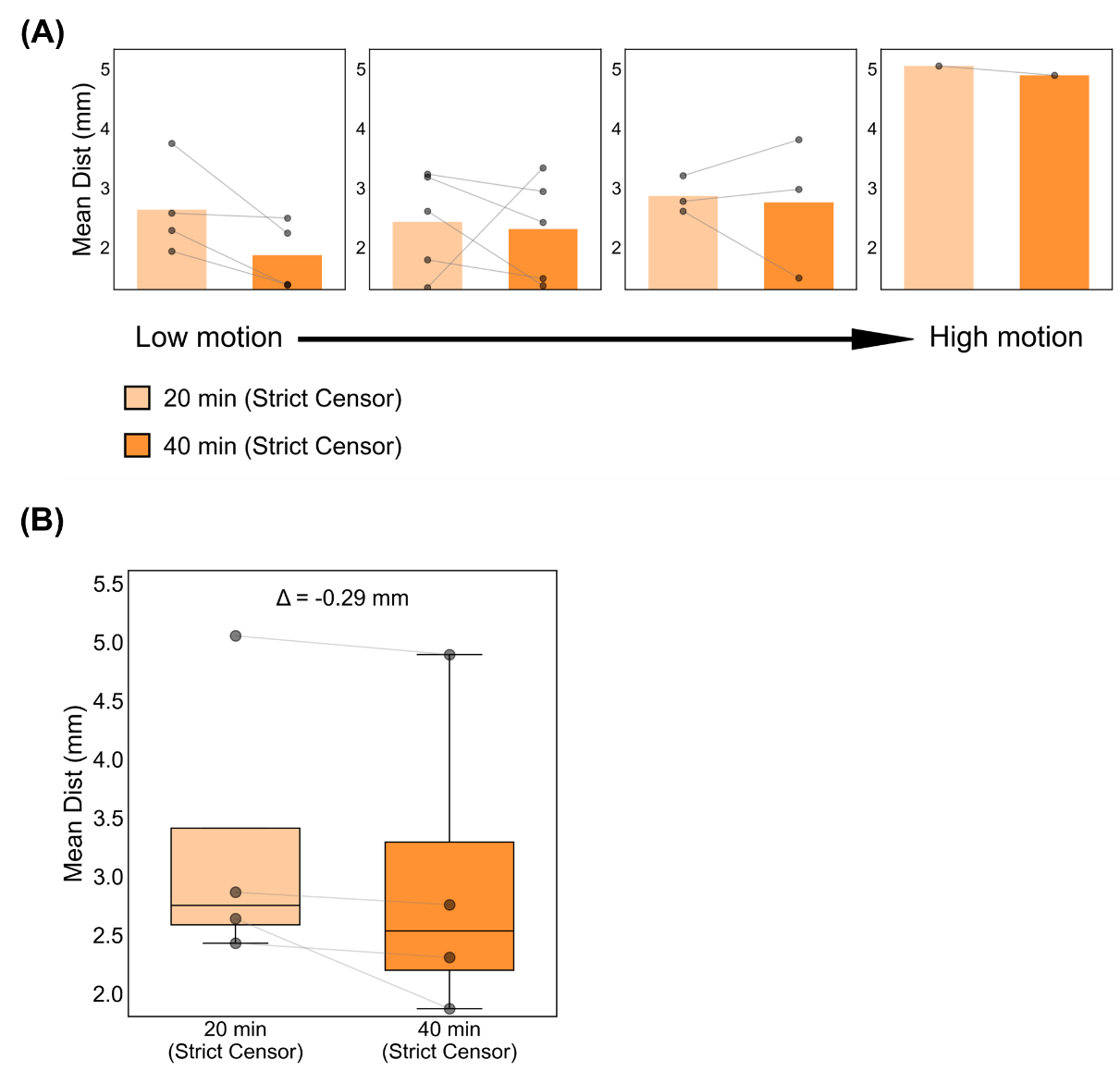

##### **Supplementary Figure 10.** Comparison of TMS depression target quality (Euclidean distance to ground-truth depression targets) between 20 minutes and 40 minutes of rs-fMRI data per session under strict censoring for the FD setup. Here, strict censoring refers to a FD threshold of 0.08 mm (after pseudo-motion filtering) and a DVARS threshold of 50. For this analysis, high-motion runs were not discarded. Smaller Euclidean distance indicates better quality. TMS depression targets were estimated using the tree-algorithm (Kong et al., 2026a). (A) TMS depression target quality for the 20-minute and 40-minute conditions across four motion bins. Each cell indicates a motion bin. Each dot represents the mean Euclidean distance across 50 simulated sessions of a participant in that motion bin. (B) TMS depression target quality across all four motion bins for both durations. There are four pairs of dots, corresponding to the four motion bins where comparisons could be made between the two durations. Δ indicates the mean improvement in Euclidean distance from the 20-minute to the 40-minute condition across the four motion bins. For each box plot, the horizontal line indicates the median across motion bins. The bottom and top edges of the box indicate the 25th and 75th percentiles, respectively. The outliers are defined as data points beyond 1.5 times the interquartile range. The whiskers extend to the most extreme data points not considered outliers.

#### **Two runs FD for parcellations**

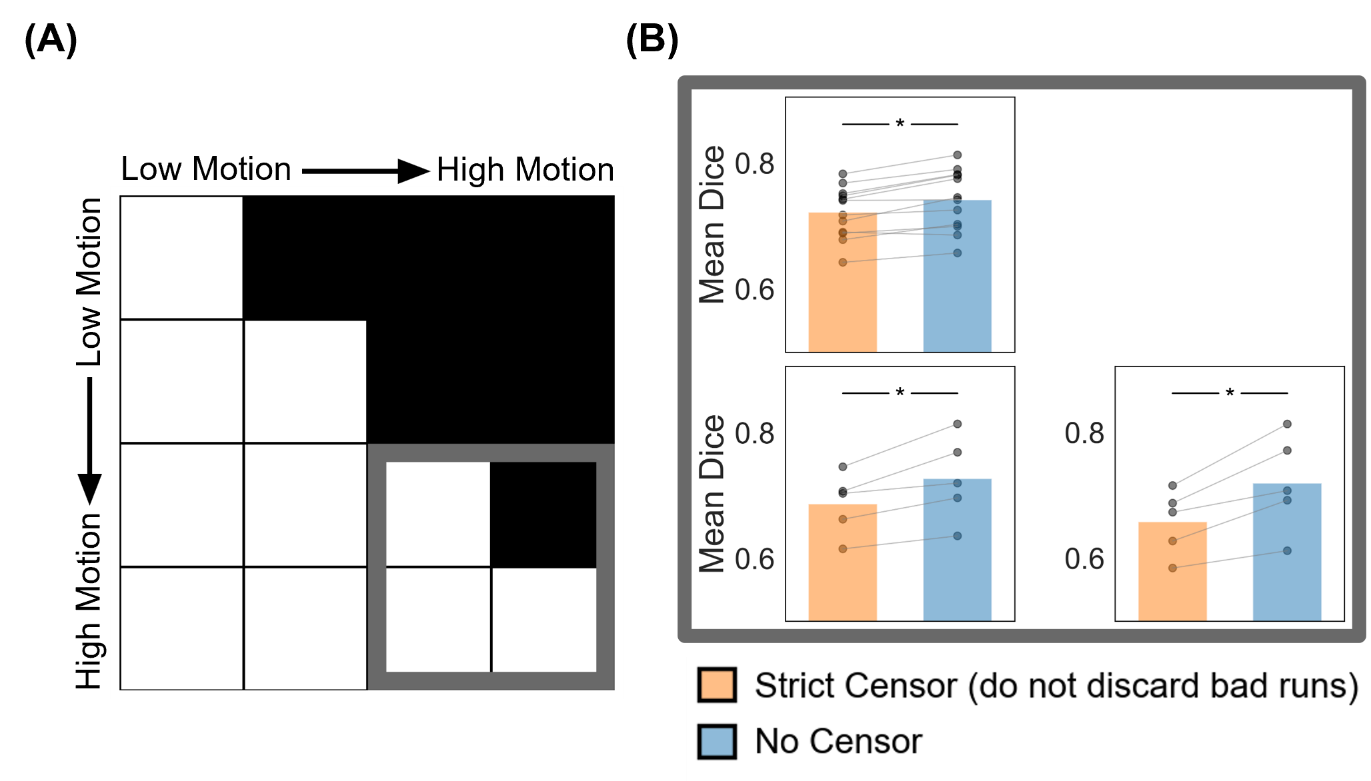

##### **Supplementary Figure 11.** Comparison of parcellation quality (Dice) between strict censoring and no censoring for the high-motion bins in the two-run FD setup. Here strict censoring refers to FD threshold of 0.08 mm (after pseudo-motion filtering) and DVARS threshold of 50. High-motion runs were not discarded for strict censoring, so a comparison could be made with no censoring (A) Schematic highlighting the three high-motion bins with thick gray outlines. (B) Parcellation quality (Dice) for the two censoring approaches across the 3 high-motion bins. Each cell in the grid indicates a motion bin. Each dot represents the mean Dice coefficient across 50 simulated sessions for a participant in that motion bin. P-values were computed using the two-sided paired-sampled t-test. "*" indicates statistical significance after multiple comparisons correction with FDR q < 0.05.

#### **One run FD for parcellations**

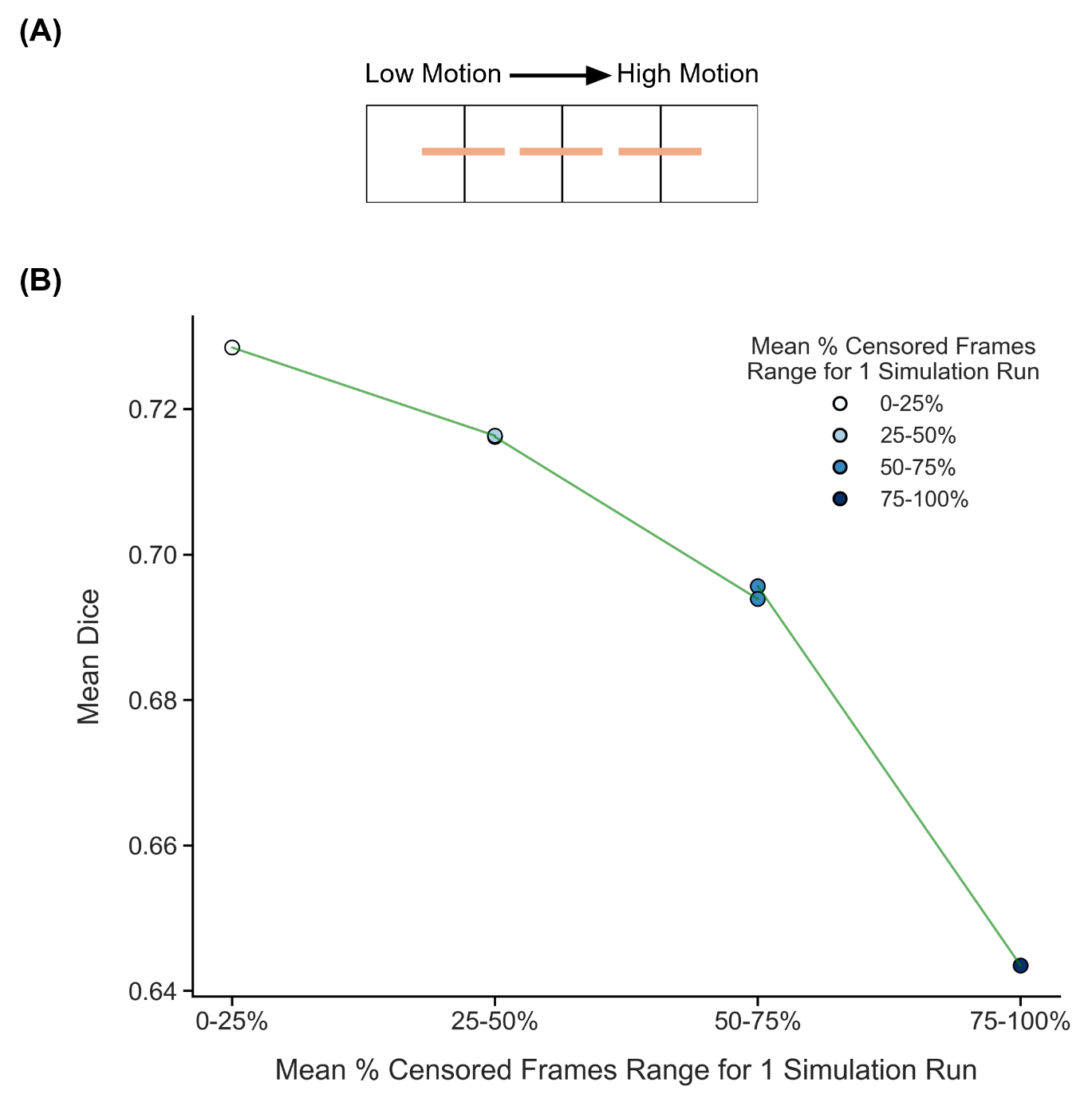

##### **Supplementary Figure 12.** Comparison of parcellation quality (Dice) across adjacent motion bins under strict censoring, for the one-run FD setup. Here strict censoring refers to FD threshold of 0.08 mm (after pseudo-motion filtering) and DVARS threshold of 50. Here, high-motion runs were not discarded to allow comparison with low-motion runs. Higher Dice indicates better quality. (A) Schematic of comparisons across adjacent motion bins. Each orange line indicates a single comparison. (B) Relationship between parcellation quality (Dice) and motion. Each circle indicates the mean Dice coefficient across simulation runs of all participants in a motion bin. Each line indicates a single comparison between two adjacent motion bins. P-values were computed using the two-sided paired-sampled t-test using only participants common to both motion bins. An overall statistical test across all comparisons was conducted using a linear mixed effects model (p = 1.9e-22).

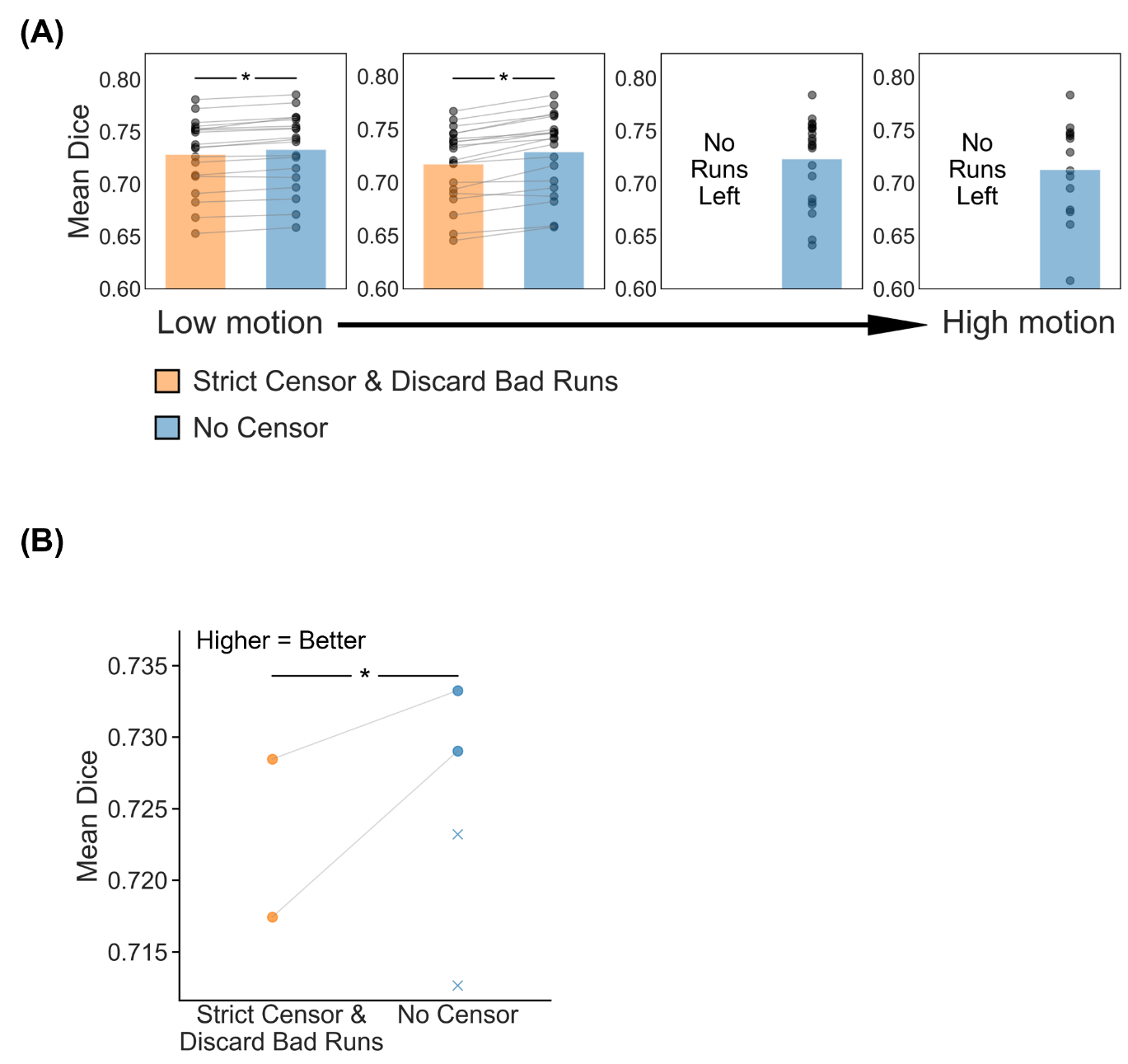

##### **Supplementary Figure 13.** Comparison of parcellation quality (Dice) between strict censoring and no censoring, for the one-run FD setup. Here strict censoring refers to FD threshold of 0.08 mm (after pseudo-motion filtering) and DVARS threshold of 50. High-motion runs were discarded. Higher Dice indicates better quality. (A) Parcellation quality for the two censoring approaches across 4 motion bins. Each cell indicates a motion bin. Each dot represents the mean Dice coefficient across 50 simulated sessions of a participant in that motion bin. P-values were computed using the two-sided paired-sampled t-test. "*" indicates statistical significance after multiple comparisons correction with FDR q < 0.05. Under strict censoring, no runs remained for the high-motion bins because all fMRI runs were discarded under this criterion. (B) Parcellation quality across all motion bins for both censoring strategies. There are two pairs of dots, corresponding to the two motion bins where comparisons could be made between strict censoring and no censoring. P-value was computed using a linear mixed effects model (p = 2.6e-10). "*" indicates statistical significance after multiple comparisons correction with FDR q < 0.05. There are two “×” corresponding to the Dice coefficient of the high-motion bins under the no censoring condition.

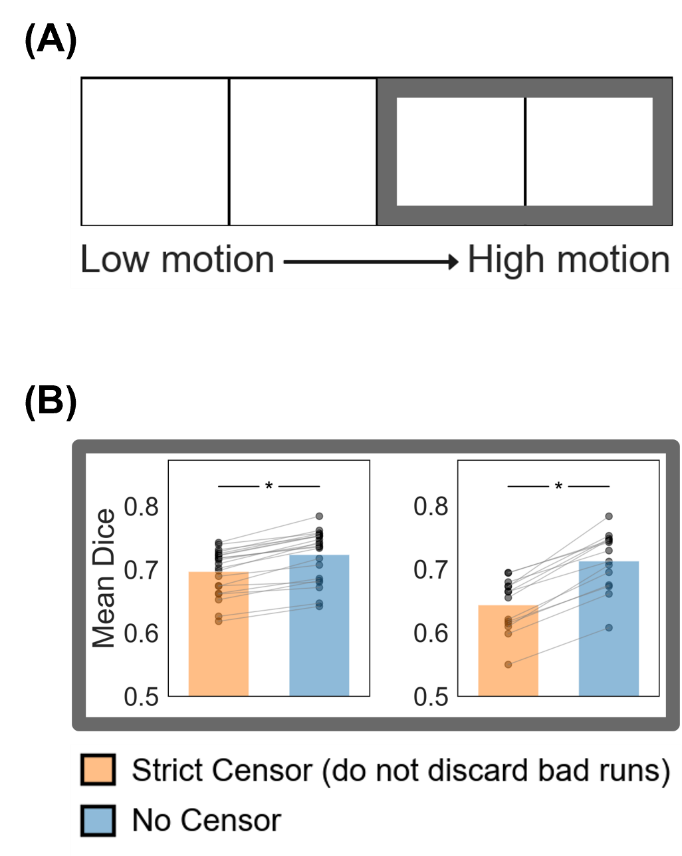

##### **Supplementary Figure 14.** Comparison of parcellation quality (Dice) between strict censoring and no censoring for the high-motion bins in the one-run FD setup. Here strict censoring refers to FD threshold of 0.08 mm (after pseudo-motion filtering) and DVARS threshold of 50. High-motion runs were not discarded for strict censoring, so a comparison could be made with no censoring (A) Schematic highlighting the two high-motion bins with thick gray outlines. (B) Parcellation quality (Dice) for the two censoring approaches across the two high-motion bins. Each cell indicates a motion bin. Each dot represents the mean Dice coefficient across 50 simulated sessions for a participant in that motion bin. P-values were computed using the two-sided paired-sampled t-test. "*" indicates statistical significance after multiple comparisons correction with FDR q < 0.05.

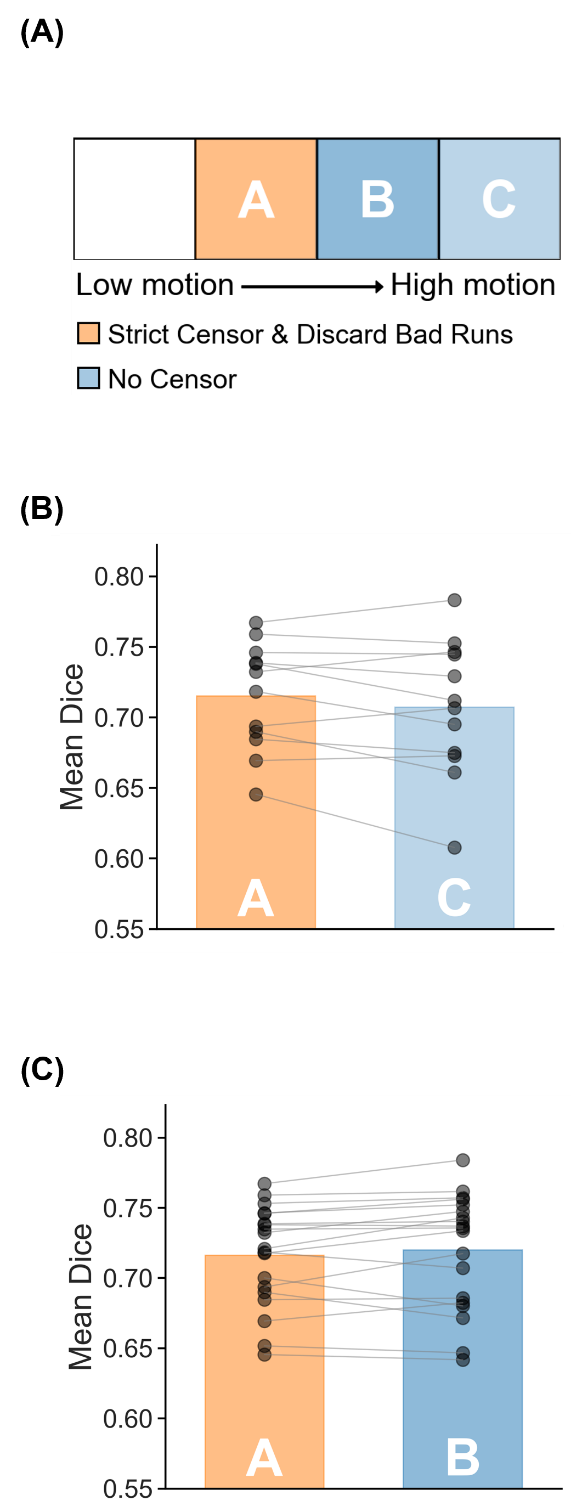

##### **Supplementary Figure 15.** Parcellation quality (Dice) for high-motion run with no censoring vs low-motion run with strict censoring. Strict censoring refers to FD threshold of 0.08 mm (after pseudo-motion filtering) and DVARS threshold of 50. Higher Dice indicates better quality. (A) Schematic illustrating the comparison. (B) Comparison of motion bins A vs C. (C) Comparison of motion bins A vs B. For both panels (B) and (C), each dot represents the mean Dice coefficient across 50 simulated sessions for a participant in that motion bin. P-values were computed using the two-sided paired-sampled t-test. Each statistical test was performed using only participants common to both motion bins. "*" indicates statistical significance after multiple comparisons correction with FDR q < 0.05.

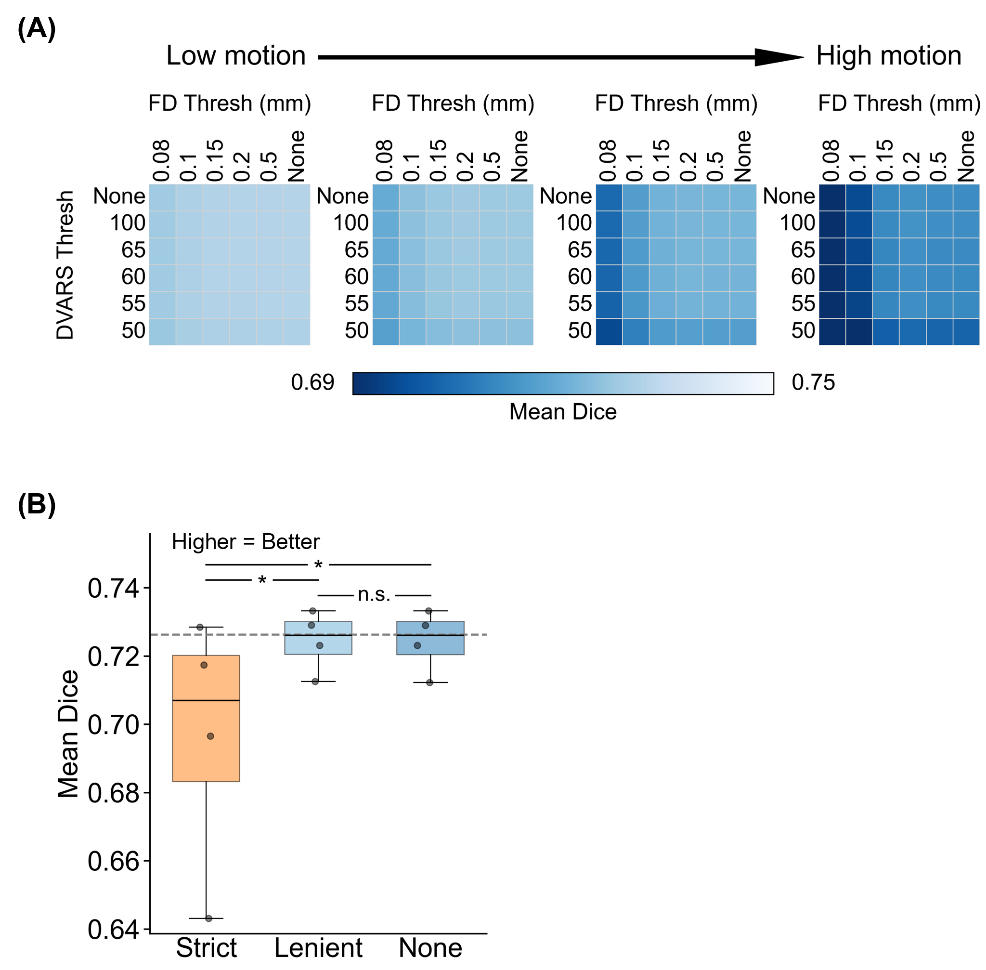

##### **Supplementary Figure 16.** Comparison of parcellation quality (Dice) across varying FD and DVARS thresholds for the one-run FD setup. For this analysis, high-motion runs were not discarded. Higher Dice indicates better quality. (A) Parcellation quality for different combinations of FD and DVARS thresholds for 4 motion bins. Each cell indicates a motion bin. For each motion bin, there are 36 small cells corresponding to different FD and DVARS thresholds. (B) Parcellation quality of strict censoring, lenient censoring and no censoring. Here strict censoring refers to FD threshold of 0.08 mm (after pseudo-motion filtering) and DVARS threshold of 50. Lenient censoring refers to FD threshold of 0.5 mm and DVARS threshold of 100. Each dot represents the Dice coefficient for a motion bin. The gray dashed line is the “noise ceiling”, achieved by picking the best combination of FD and DVARS threshold for each motion bin, and computing the median across motion bins. P-values were computed using a linear mixed effects model (p = 1.4e-25 for strict vs. none, p = 1.1e-25 for strict vs. lenient, p = 0.98 for lenient vs. none). "*" indicates statistical significance after multiple comparisons correction with FDR q < 0.05 and "n.s." indicates not significant after FDR correction. For each box plot, the horizontal line indicates the median across all motion bins. The bottom and top edges of the box indicate the 25th and 75th percentiles, respectively. The outliers are defined as data points beyond 1.5 times the interquartile range. The whiskers extend to the most extreme data points not considered outliers.

#### **Two runs FD for tree-based (Kong2026) TMS depression targets**

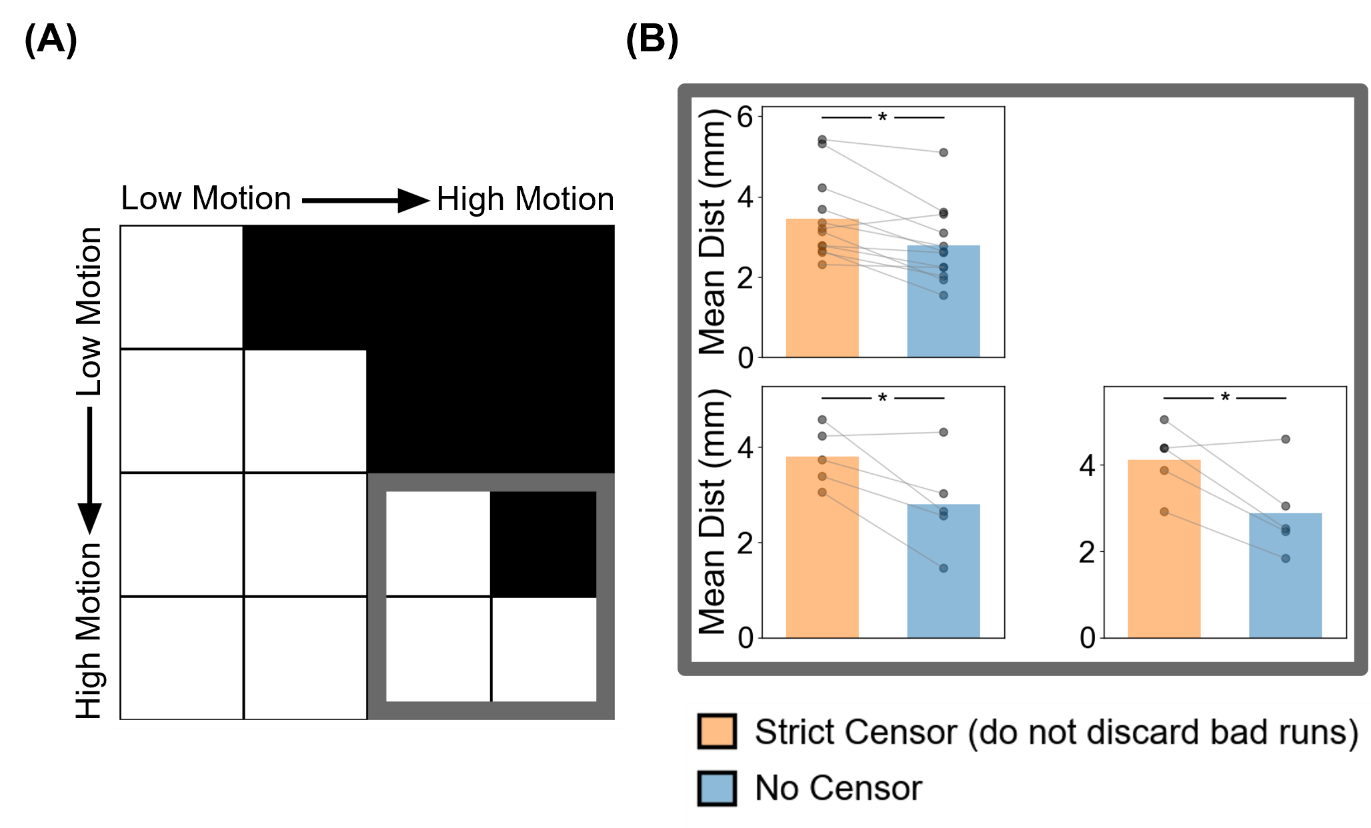

##### **Supplementary Figure 17.** Comparison of TMS depression targeting quality (Euclidean distance from ground-truth targets) between strict censoring and no censoring for the high-motion bins in the two-run FD setup. Here strict censoring refers to FD threshold of 0.08 mm (after pseudo-motion filtering) and DVARS threshold of 50. High-motion runs were not discarded for strict censoring, so a comparison could be made with no censoring. Smaller Euclidean distance indicates better quality. TMS depression targets were estimated using the tree-algorithm (Kong et al., 2026a). (A) Schematic highlighting the three high-motion bins with thick gray outlines. (B) TMS targeting quality (Euclidean distance) for the two censoring approaches across the 3 high-motion bins. Each cell in the grid indicates a motion bin. Each dot represents the mean Euclidean distance across 50 simulated sessions for a participant in that motion bin. P-values were computed using the two-sided paired-sampled t-test. "*" indicates statistical significance after multiple comparisons correction with FDR q < 0.05.

#### **One run FD for tree-based (Kong2026) TMS depression targets**

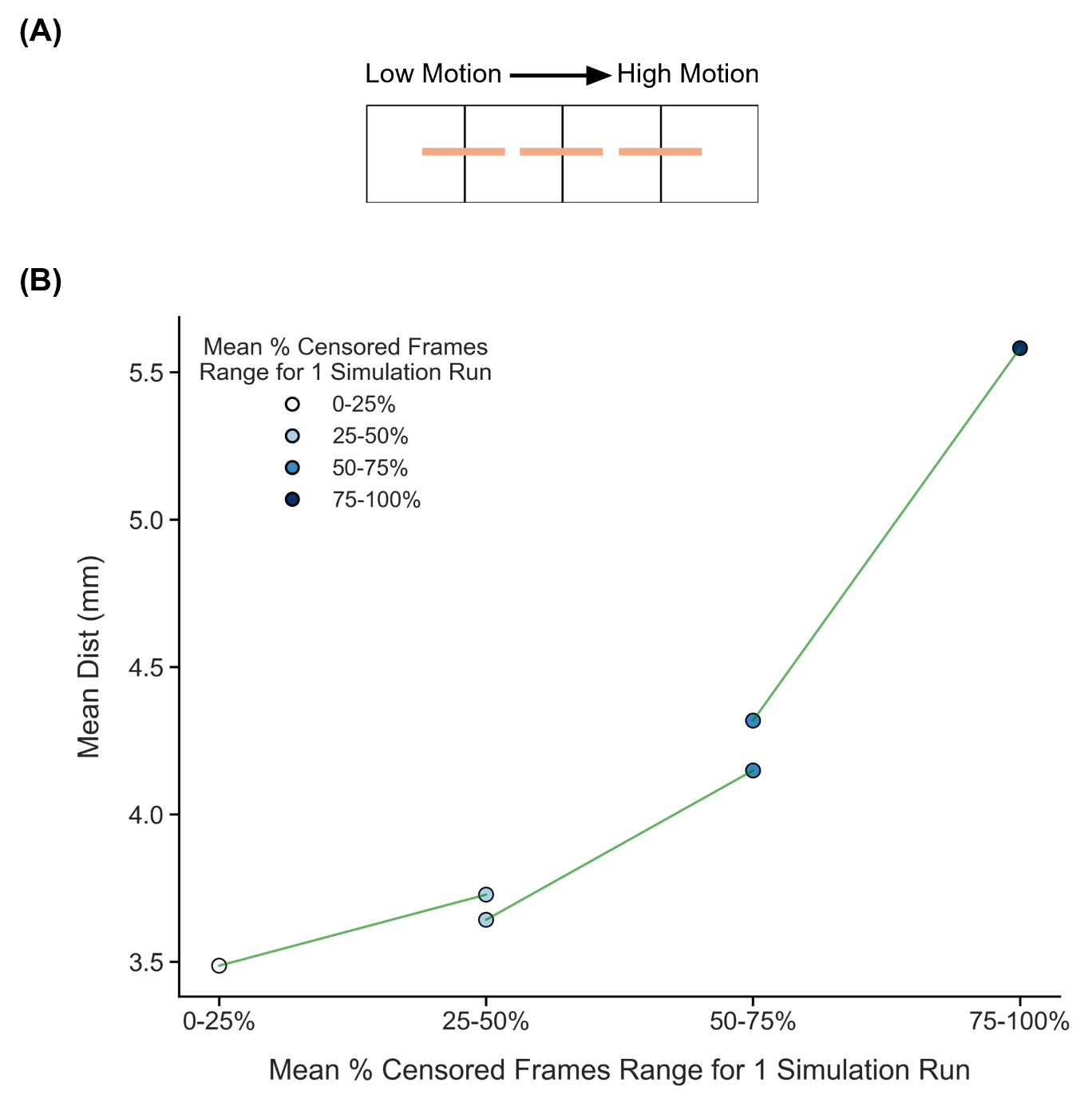

##### **Supplementary Figure 18.** Comparison of TMS depression target quality (Euclidean distance from ground-truth targets) across adjacent motion bins under strict censoring, for the one-run FD setup. Here strict censoring refers to FD threshold of 0.08 mm (after pseudo-motion filtering) and DVARS threshold of 50. High-motion runs were not discarded to allow comparison with low-motion runs. Smaller Euclidean distance to ground-truth target indicates better quality. TMS depression targets were estimated using the tree-algorithm (Kong et al., 2026a). (A) Schematic of comparisons across adjacent motion bins. Each orange line indicates a single comparison. (B) Relationship between TMS depression target quality (Euclidean distance) and motion. Each circle indicates the mean Euclidean distance across simulation runs of all participants in a motion bin. Each line indicates a single comparison between two adjacent motion bins. P-values were computed using the two-sided paired-sampled t-test using only participants common to both motion bins. An overall statistical test combining all comparisons was conducted using a linear mixed effects model (p = 7e-6).

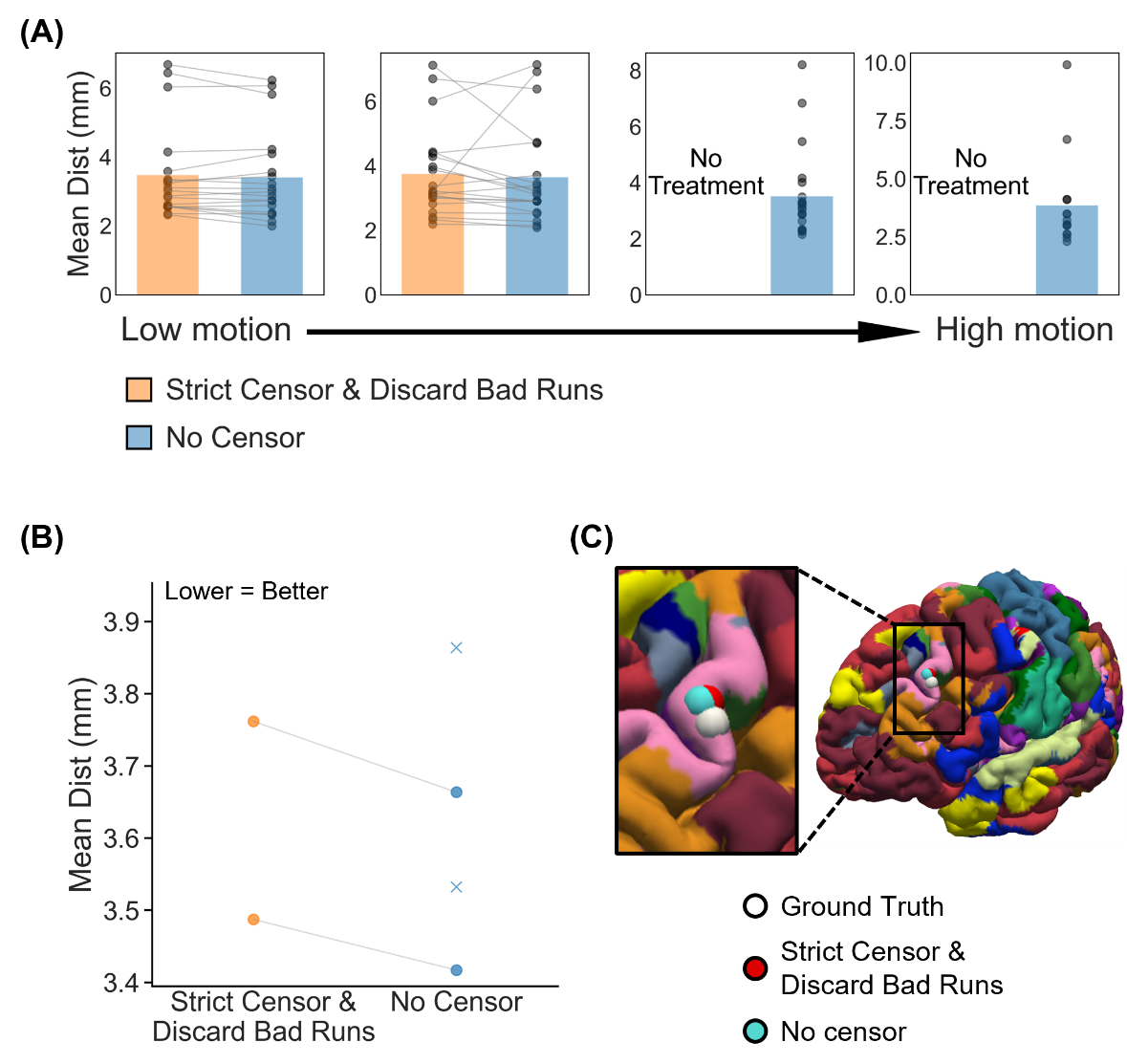

##### **Supplementary Figure 19.** Comparison of TMS depression targeting quality (Euclidean distance to ground-truth depression targets) between strict censoring and no censoring, for the one-run FD setup. Here strict censoring refers to FD threshold of 0.08 mm (after pseudo-motion filtering) and DVARS threshold of 50. High-motion runs were discarded. Smaller Euclidean distance indicates better quality. TMS depression targets were estimated using the tree-based algorithm (Kong et al., 2026a). (A) TMS depression targeting quality (Euclidean distance) for the two censoring approaches across 4 motion bins. Each cell indicates a motion bin. Each dot represents the mean Euclidean distance across 50 simulated sessions of a participant in that motion bin. P-values were computed using the two-sided paired-sampled t-test. "*" indicates statistical significance after multiple comparisons correction with FDR q < 0.05. Under strict censoring, no runs remained for the high-motion bins because all fMRI runs were discarded under this criterion. (B) TMS depression targeting quality across all motion bins for both censoring strategies. There are two pairs of dots, corresponding to the two motion bins where comparisons could be made between strict censoring and no censoring. P-value was computed using a linear mixed effects model (p = 0.52). "*" indicates statistical significance after multiple comparisons correction with FDR q < 0.05. There are two “×” corresponding to the mean Euclidean distance of the high-motion bins under the no censoring condition. The × were not included in the boxplots. For each box plot, the horizontal line indicates the median. The bottom and top edges of the box indicate the 25th and 75th percentiles, respectively. The outliers are defined as data points beyond 1.5 times the interquartile range. The whiskers extend to the most extreme data points not considered outliers. (C) Visualization of TMS depression targets in a representative participant from the highest motion bin where the strict censoring had results.

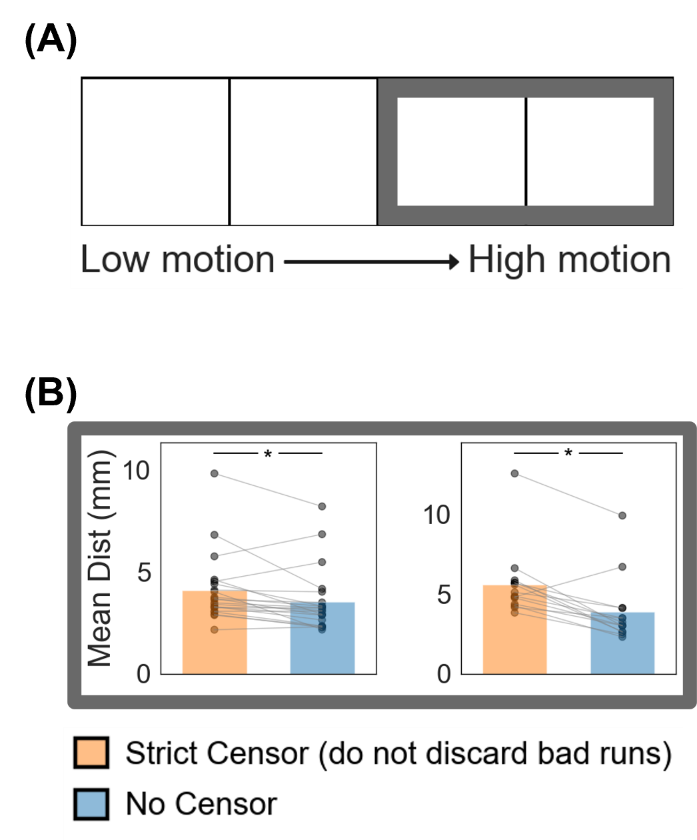

##### **Supplementary Figure 20.** Comparison of TMS depression targeting quality (Euclidean distance from ground-truth targets) between strict censoring and no censoring for the high-motion bins in the one-run FD setup. Here strict censoring refers to FD threshold of 0.08 mm (after pseudo-motion filtering) and DVARS threshold of 50. High-motion runs were not discarded for strict censoring, so a comparison could be made with no censoring. Smaller Euclidean distance indicates better quality. TMS depression targets were estimated using the tree-algorithm (Kong et al., 2026a). (A) Schematic highlighting the two high-motion bins with thick gray outlines. (B) TMS targeting quality (Euclidean distance) for the two censoring approaches across the two high-motion bins. Each cell indicates a motion bin. Each dot represents the mean Euclidean distance across 50 simulated sessions for a participant in that motion bin. P-values were computed using the two-sided paired-sampled t-test. "*" indicates statistical significance after multiple comparisons correction with FDR q < 0.05.

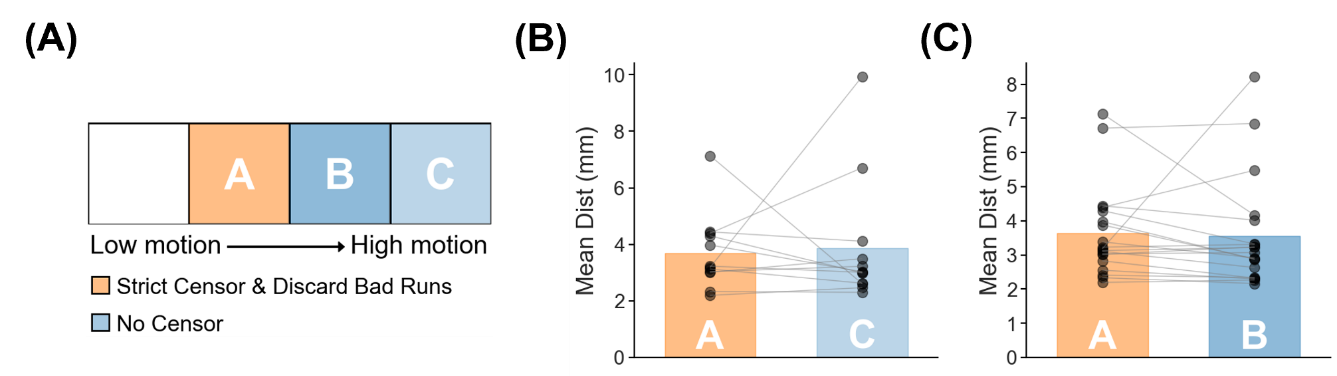

##### **Supplementary Figure 21.** TMS depression target quality (Euclidean distance to ground-truth targets) for one high-motion run with no censoring vs one low-motion run with strict censoring. Strict censoring refers to FD threshold of 0.08 mm (after pseudo-motion filtering) and DVARS threshold of 50. Lower Euclidean distance indicates better quality. TMS depression targets were estimated using the tree-algorithm (Kong et al., 2026a). (A) Schematic illustrating the comparison. (B) Comparison of motion bins A vs C. (C) Comparison of motion bins A vs B. For both panels (B) and (C), each dot represents the mean Euclidean distance across 50 simulated sessions for a participant in that motion bin. P-values were computed using the two-sided paired-sampled t-test. Each statistical test was performed using only participants common to both motion bins. "*" indicates statistical significance after multiple comparisons correction with FDR q < 0.05.

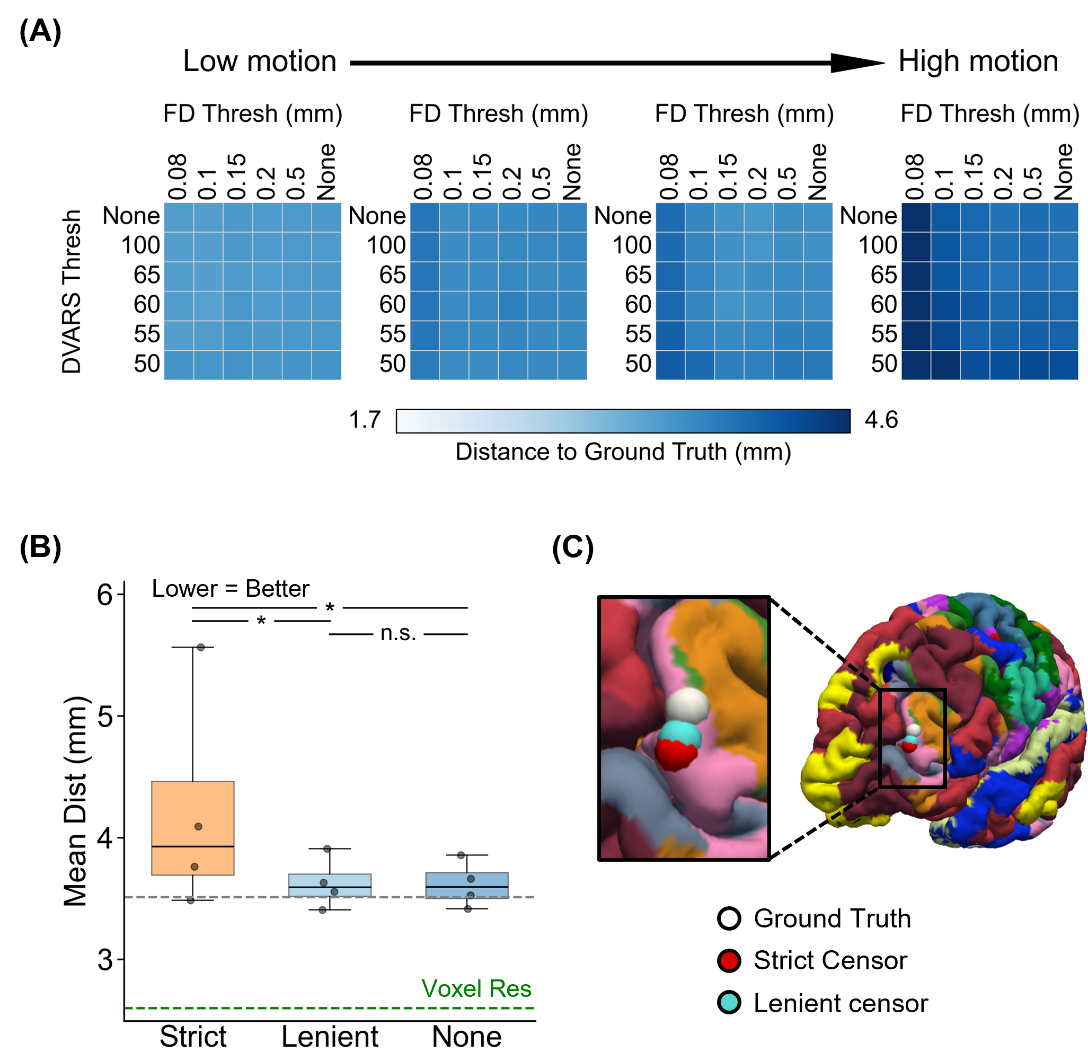

##### **Supplementary Figure 22.** Comparison of TMS depression target quality (Euclidean distance from ground-truth targets) across varying FD and DVARS thresholds for the one-run FD setup. For this analysis, high-motion runs were not discarded. Lower Euclidean distance indicates better quality. TMS depression targets were estimated using the tree-algorithm (Kong et al., 2026a). (A) TMS depression target quality (Euclidean distance) for different combinations of FD and DVARS thresholds for 4 motion bins. Each cell indicates a motion bin. For each motion bin, there are 36 small cells corresponding to different FD and DVARS thresholds. (B) TMS depression target quality for strict censoring, lenient censoring and no censoring. Here strict censoring refers to FD threshold of 0.08 mm (after pseudo-motion filtering) and DVARS threshold of 50. Lenient censoring refers to FD threshold of 0.5 mm and DVARS threshold of 100. Each dot represents the mean Euclidean distance for a motion bin. The gray dashed line is the “noise ceiling”, achieved by picking the best combination of FD and DVARS threshold for each motion bin, and computing the median across motion bins. P-values were computed using a linear mixed effects model (p = 9.3e-7 for strict vs. none, p = 1.3e-6 for strict vs. lenient, p = 0.95 for lenient vs. none). "*" indicates statistical significance after multiple comparisons correction with FDR q < 0.05 and "n.s." indicates not significant after FDR correction. For each box plot, the horizontal line indicates the median across all motion bins. The bottom and top edges of the box indicate the 25th and 75th percentiles, respectively. The outliers are defined as data points beyond 1.5 times the interquartile range. The whiskers extend to the most extreme data points not considered outliers. (C) Visualization of TMS depression targets in a representative participant from the highest motion bin.

#### **Cone (Fox2013) TMS depression targets (two runs FD)**

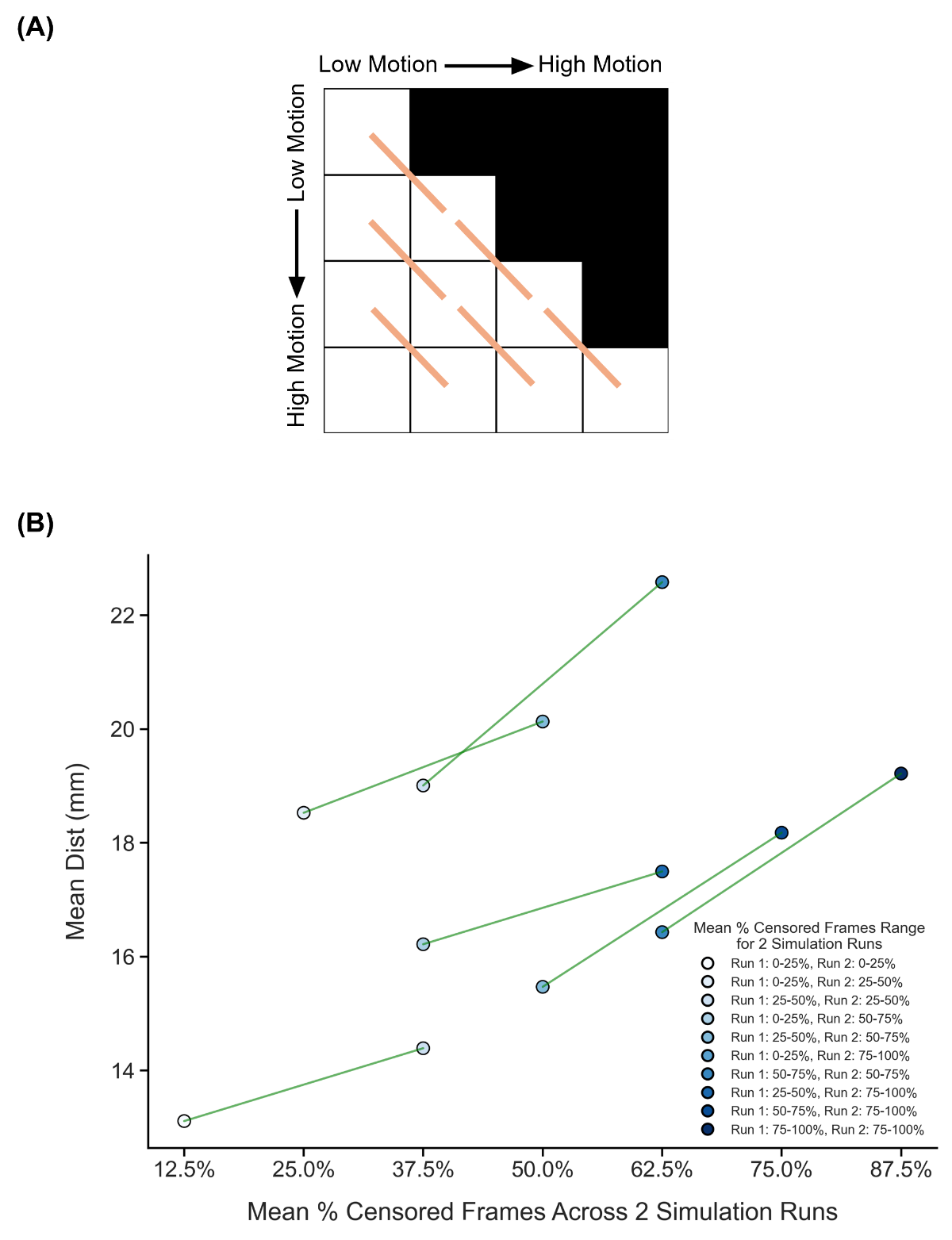

##### **Supplementary Figure 23.** Comparison of TMS depression target quality (Euclidean distance from ground-truth targets) across diagonally adjacent motion bins under strict censoring, for the two-run FD setup. Here strict censoring refers to FD threshold of 0.08 mm (after pseudo-motion filtering) and DVARS threshold of 50. High-motion runs were not discarded to allow comparison with low-motion runs. Smaller Euclidean distance to ground-truth target indicates better quality. TMS depression targets were estimated using the cone approach (Fox et al., 2013). (A) Schematic of comparisons across diagonally adjacent motion bins. Each orange line indicates a single comparison. (B) Relationship between TMS depression target quality (Euclidean distance) and motion. Each circle indicates the mean Euclidean distance across simulation runs of all participants in a motion bin. Each line indicates a single comparison between two adjacent motion bins. P-values were computed using the two-sided paired-sampled t-test using only participants common to both motion bins. An overall statistical test combining all comparisons was conducted using a linear mixed effects model (p = 1e-5).

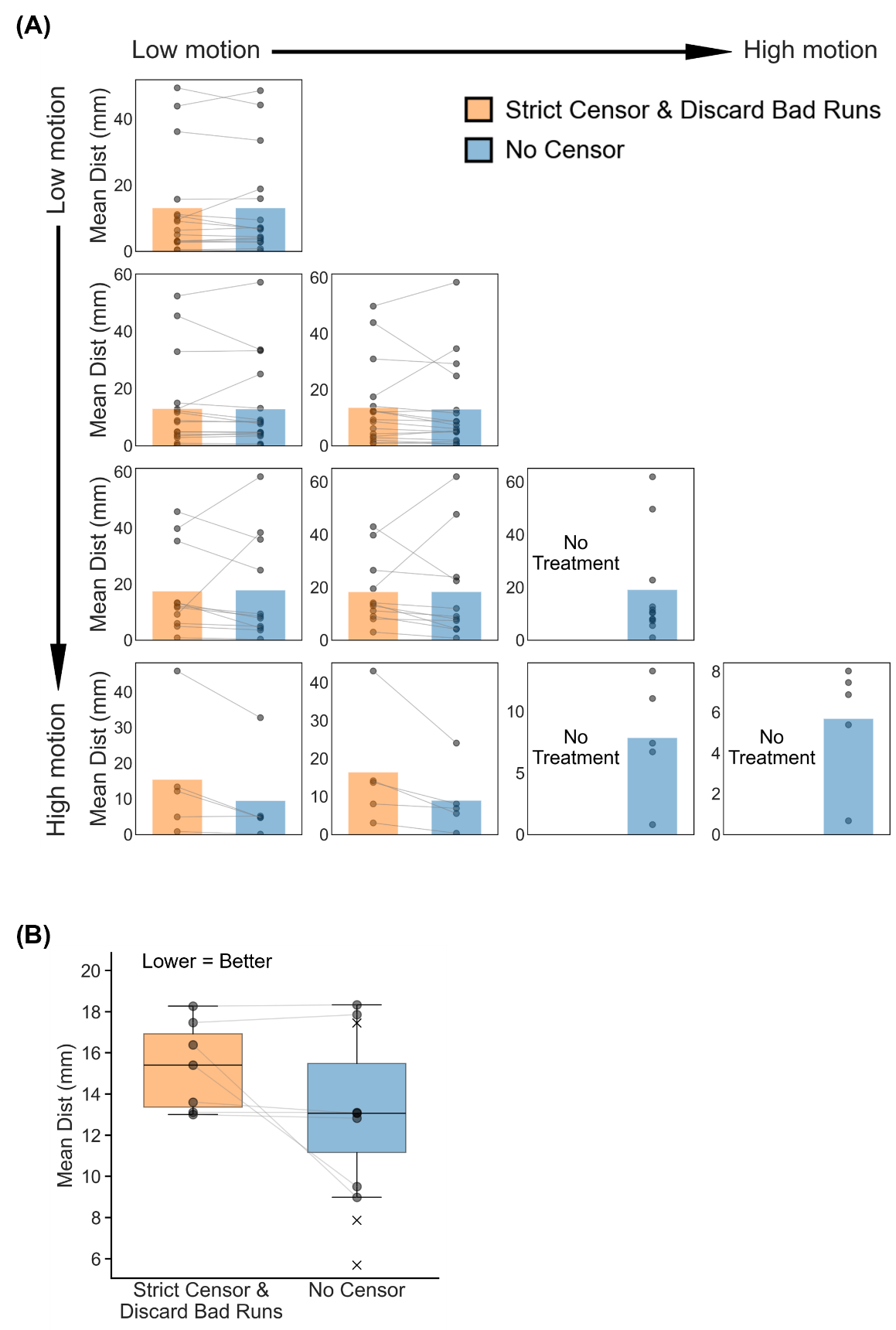

##### **Supplementary Figure 24.** Comparison of TMS depression targeting quality (Euclidean distance to ground-truth depression targets) between strict censoring and no censoring, for the two-run FD setup. Here strict censoring refers to FD threshold of 0.08 mm (after pseudo-motion filtering) and DVARS threshold of 50. High-motion runs were discarded. Smaller Euclidean distance indicates better quality. TMS depression targets were estimated using the cone approach (Fox et al., 2013). (A) TMS depression targeting quality (Euclidean distance) for the two censoring approaches across 10 motion bins. Each cell in the grid indicates a motion bin. Each dot represents the mean Euclidean distance across 50 simulated sessions of a participant in that motion bin. P-values were computed using the two-sided paired-sampled t-test. "*" indicates statistical significance after multiple comparisons correction with FDR q < 0.05. Under strict censoring, no runs remained for the bottom-right three motion bins because all fMRI runs were discarded under this criterion. (B) TMS depression targeting quality across all motion bins for both censoring strategies. There are seven pairs of dots, corresponding to the seven motion bins where comparisons could be made between strict censoring and no censoring. P-value was computed using a linear mixed effects model (p = 0.8). "*" indicates statistical significance after multiple comparisons correction with FDR q < 0.05. There are three “×” corresponding to the mean Euclidean distance of the 3 bottom-right bins under the no censoring condition. The × were not included in the boxplots. For each box plot, the horizontal line indicates the median. The bottom and top edges of the box indicate the 25th and 75th percentiles, respectively. The outliers are defined as data points beyond 1.5 times the interquartile range. The whiskers extend to the most extreme data points not considered outliers.

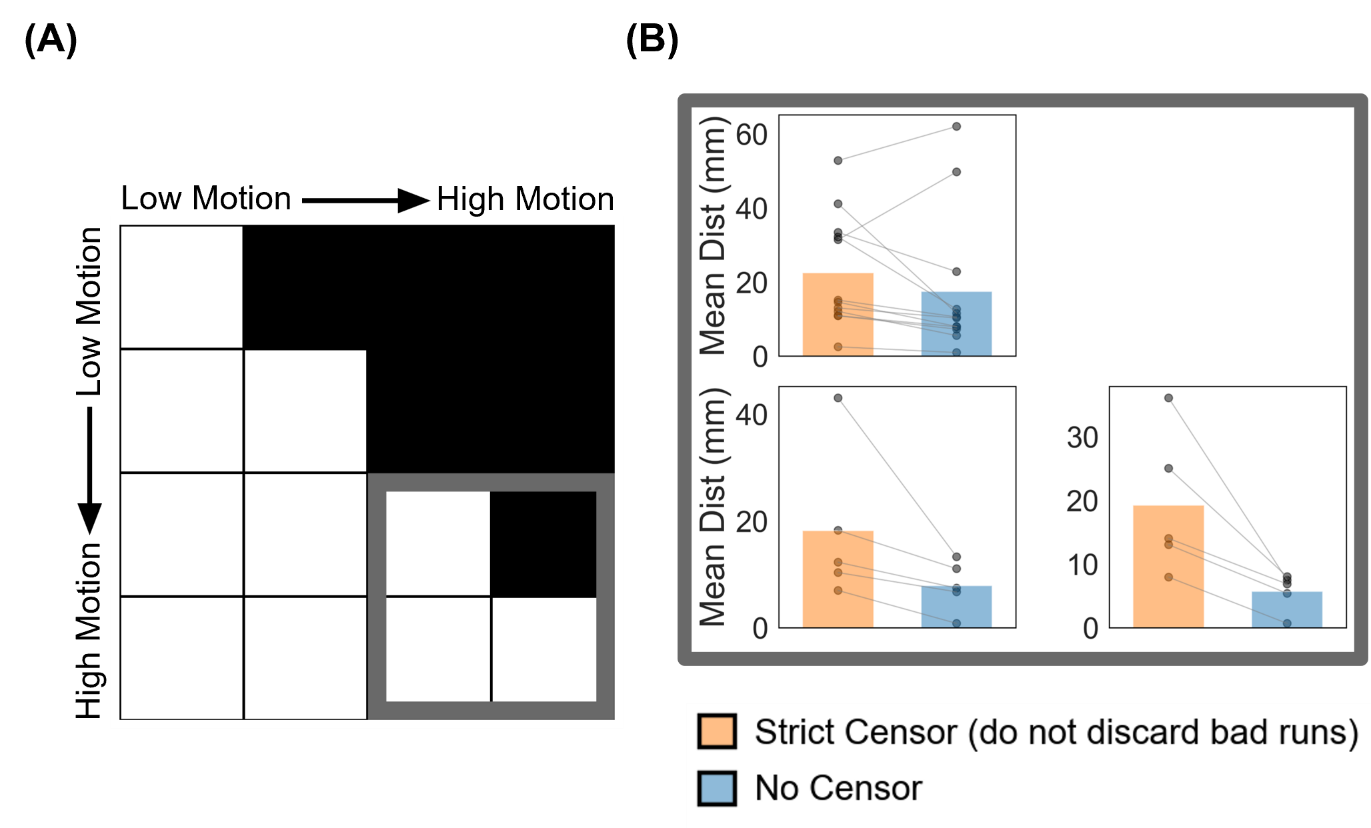

##### **Supplementary Figure 25.** Comparison of TMS depression targeting quality (Euclidean distance from ground-truth targets) between strict censoring and no censoring for the high-motion bins in the two-run FD setup. Here strict censoring refers to FD threshold of 0.08 mm (after pseudo-motion filtering) and DVARS threshold of 50. High-motion runs were not discarded for strict censoring, so a comparison could be made with no censoring. Smaller Euclidean distance indicates better quality. TMS depression targets were estimated using the cone approach (Fox et al., 2013). (A) Schematic highlighting the three high-motion bins with thick gray outlines. (B) TMS targeting quality (Euclidean distance) for the two censoring approaches across the 3 high-motion bins. Each cell in the grid indicates a motion bin. Each dot represents the mean Euclidean distance across 50 simulated sessions for a participant in that motion bin. P-values were computed using the two-sided paired-sampled t-test. "*" indicates statistical significance after multiple comparisons correction with FDR q < 0.05.

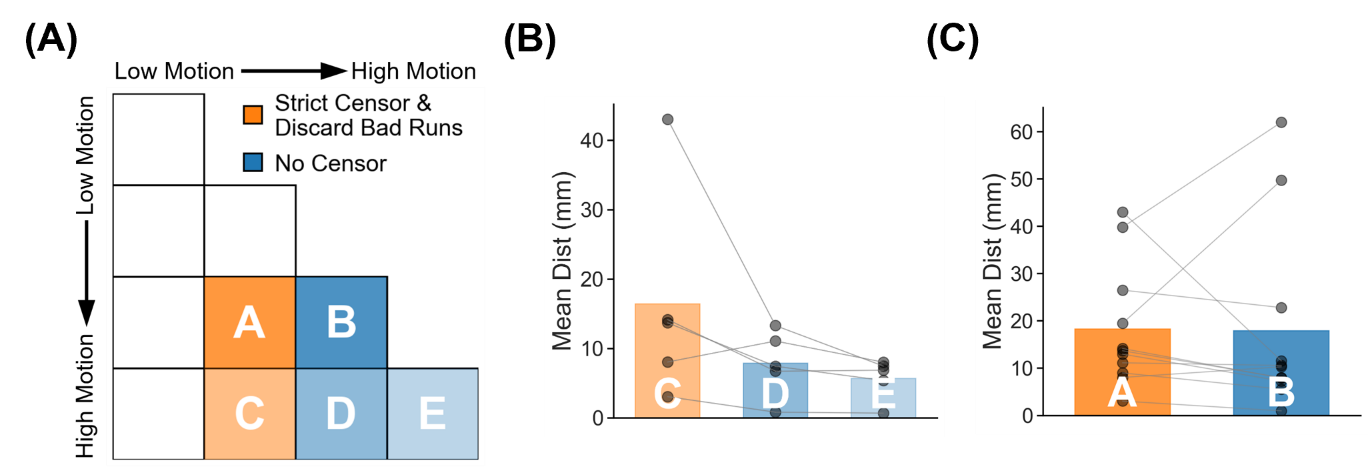

##### **Supplementary Figure 26.** TMS depression target quality (Euclidean distance to ground-truth targets) for two high-motion runs with no censoring vs mixed-motion runs with strict censoring. Here mixed-motion runs refer to sessions with one high-motion and one low-motion runs, so the high-motion run will be discarded under strict censoring. Strict censoring refers to FD threshold of 0.08 mm (after pseudo-motion filtering) and DVARS threshold of 50. Lower Euclidean distance indicates better quality. TMS depression targets were estimated using the cone approach (Fox et al., 2013). (A) Schematic illustrating the comparison. Motion bins A and B were compared, while motion bins C, D and E were compared. (B) Comparison of motion bins C vs D, and C vs E. (C) Comparison of motion bins A vs B. For both panels (B) and (C), each dot represents the mean Euclidean distance across 50 simulated sessions for a participant in that motion bin. P-values were computed using the two-sided paired-sampled t-test. Each statistical test was performed using only participants common to both motion bins. "*" indicates statistical significance after multiple comparisons correction with FDR q < 0.05.

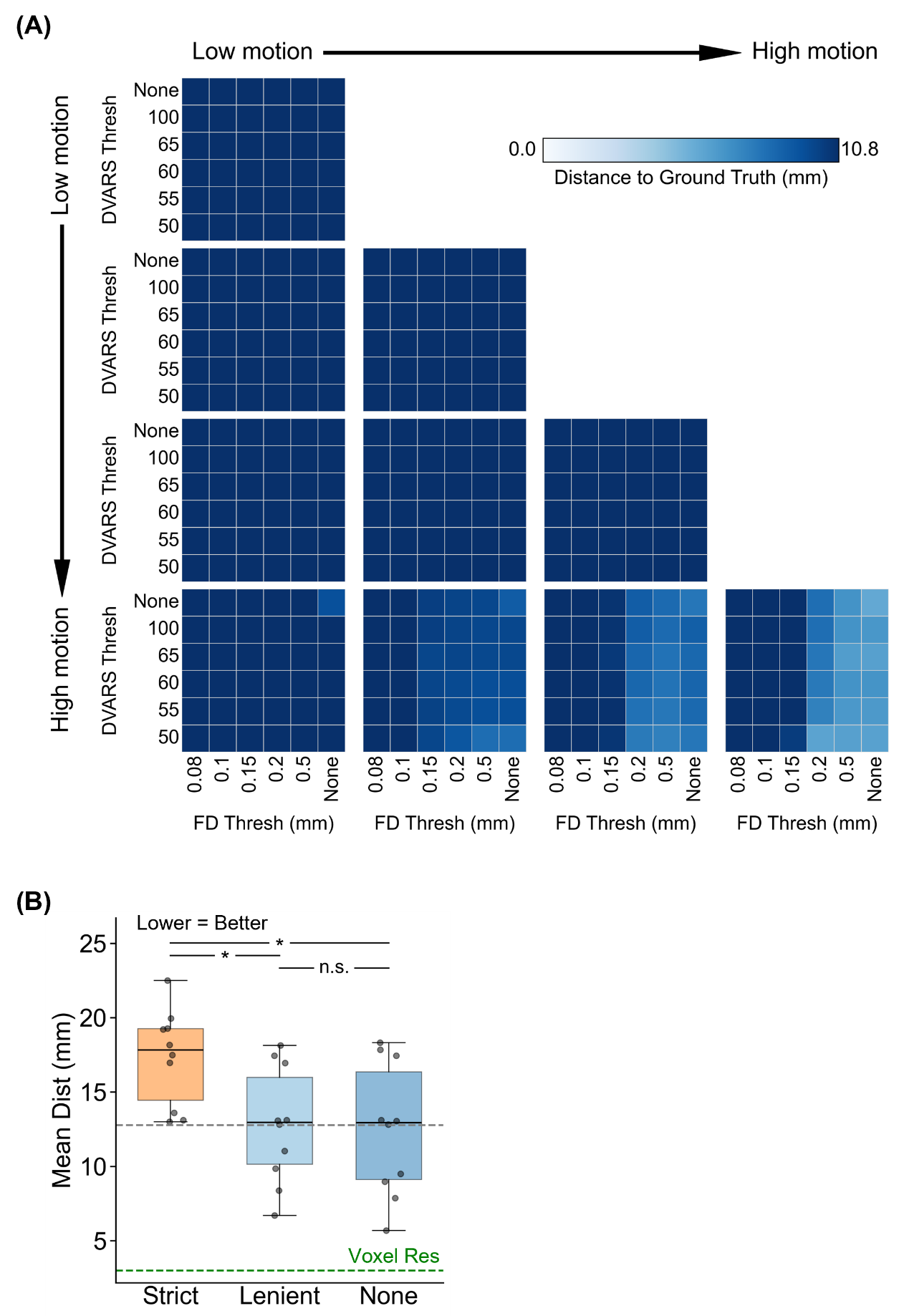

##### **Supplementary Figure 27.** Comparison of TMS depression target quality (Euclidean distance from ground-truth targets) across varying FD and DVARS thresholds for the two-run FD setup. For this analysis, high-motion runs were not discarded. Lower Euclidean distance indicates better quality. TMS depression targets were estimated using the cone approach (Fox et al., 2013). (A) TMS depression target quality (Euclidean distance) for different combinations of FD and DVARS thresholds for 10 motion bins. Each cell in the grid indicates a motion bin. For each motion bin, there are 36 small cells corresponding to different FD and DVARS thresholds. (B) TMS depression target quality for strict censoring, lenient censoring and no censoring. Here strict censoring refers to FD threshold of 0.08 mm (after pseudo-motion filtering) and DVARS threshold of 50. Lenient censoring refers to FD threshold of 0.5 mm and DVARS threshold of 100. Each dot represents the mean Euclidean distance for a motion bin. The gray dashed line is the “noise ceiling”, achieved by picking the best combination of FD and DVARS threshold for each motion bin, and computing the median across motion bins. P-values were computed using a linear mixed effects model (p = 2.7e-5 for strict vs. none, p = 4.4e-5 for strict vs. lenient, p = 0.91 for lenient vs. none). "*" indicates statistical significance after multiple comparisons correction with FDR q < 0.05 and "n.s." indicates not significant after FDR correction. For each box plot, the horizontal line indicates the median across all motion bins. The bottom and top edges of the box indicate the 25th and 75th percentiles, respectively. The outliers are defined as data points beyond 1.5 times the interquartile range. The whiskers extend to the most extreme data points not considered outliers.

#### **Cluster (Cash2021) TMS depression targets (two runs FD)**

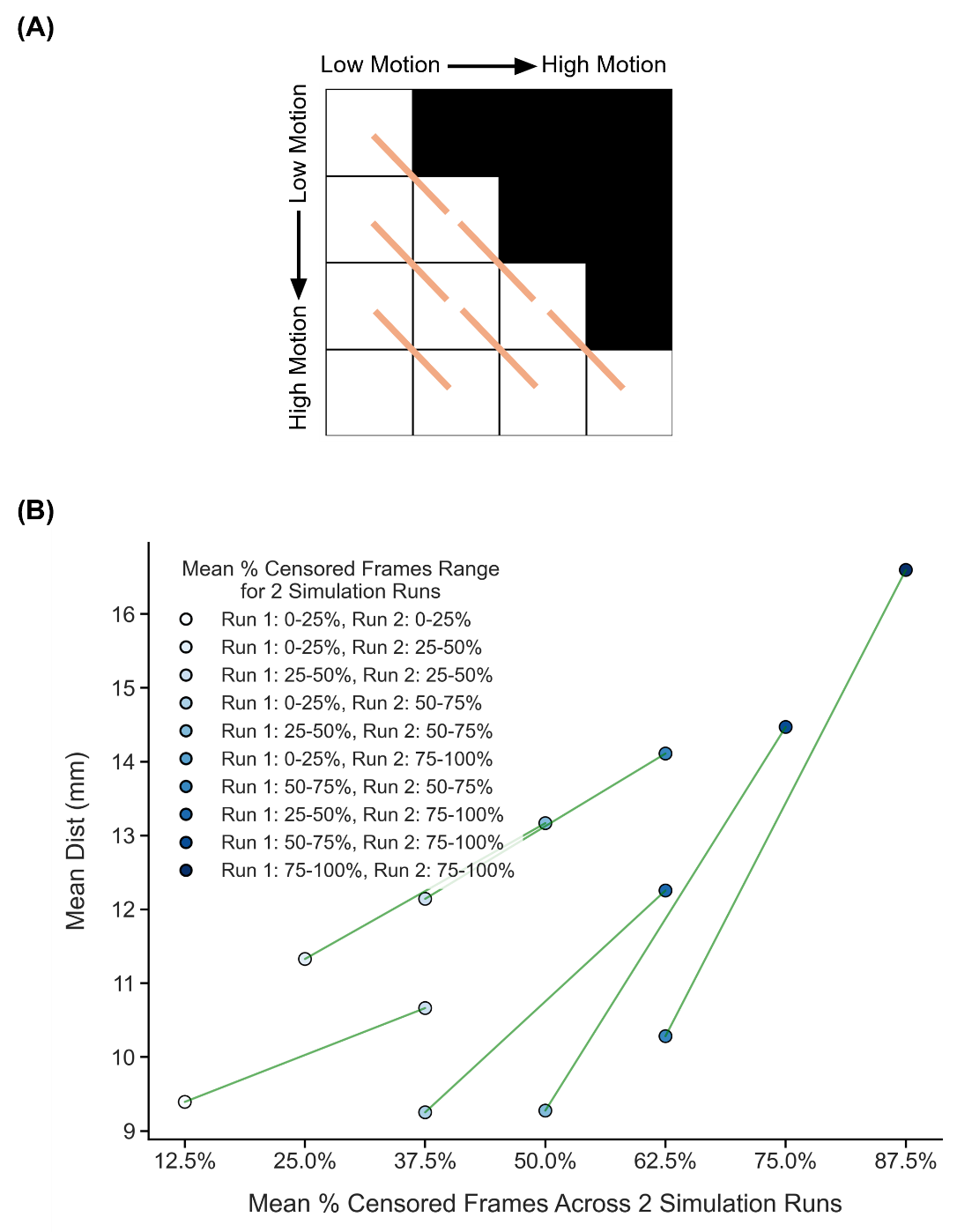

##### **Supplementary Figure 28.** Comparison of TMS depression target quality (Euclidean distance from ground-truth targets) across diagonally adjacent motion bins under strict censoring, for the two-run FD setup. Here strict censoring refers to FD threshold of 0.08 mm (after pseudo-motion filtering) and DVARS threshold of 50. High-motion runs were not discarded to allow comparison with low-motion runs. Smaller Euclidean distance to ground-truth target indicates better quality. TMS depression targets were estimated using the Cluster approach (Cash et al., 2021b). (A) Schematic of comparisons across diagonally adjacent motion bins. Each orange line indicates a single comparison. (B) Relationship between TMS depression target quality (Euclidean distance) and motion. Each circle indicates the mean Euclidean distance across simulation runs of all participants in a motion bin. Each line indicates a single comparison between two adjacent motion bins. P-values were computed using the two-sided paired-sampled t-test using only participants common to both motion bins. An overall statistical test combining all comparisons was conducted using a linear mixed effects model (p = 1.7e-7).

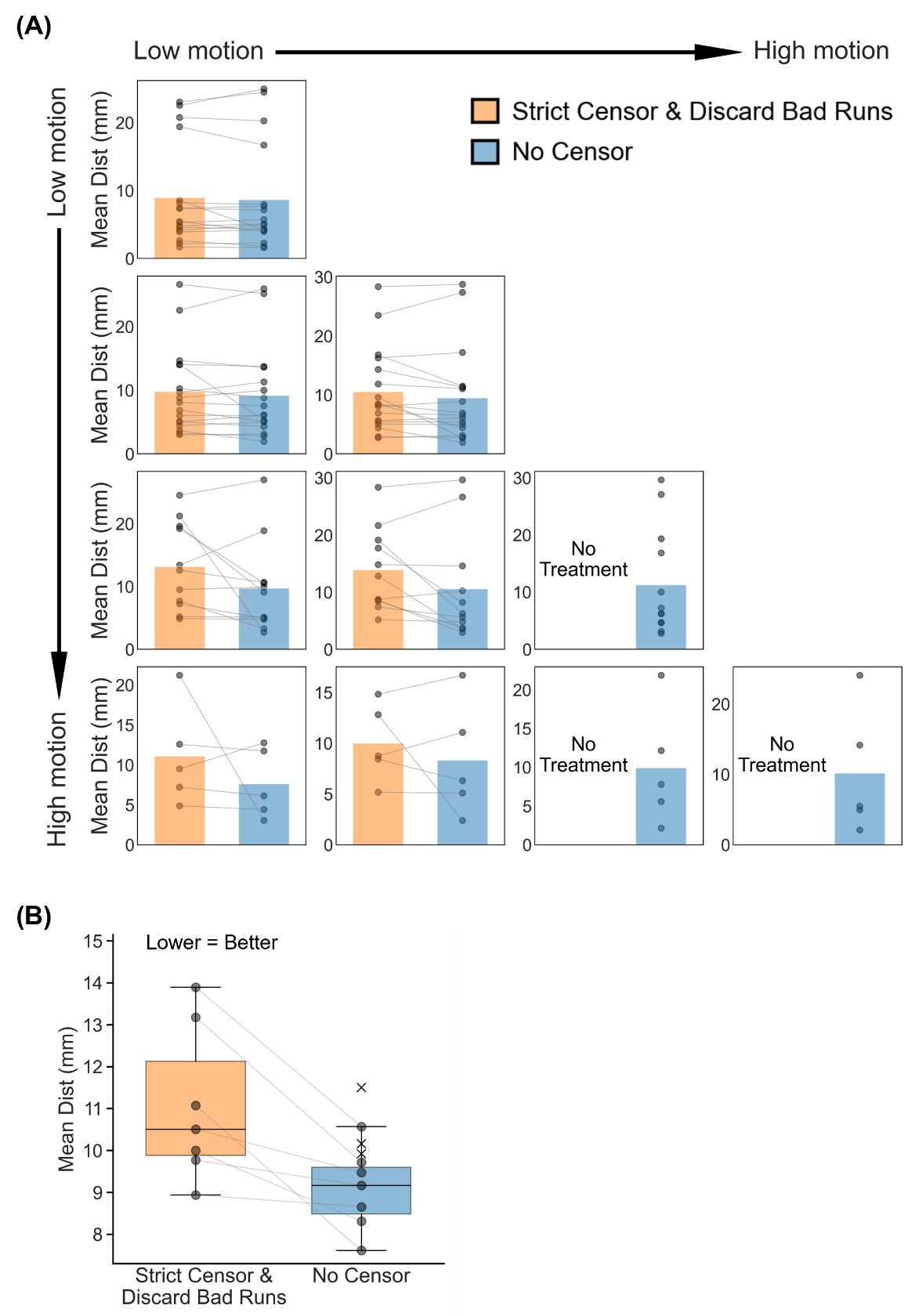

##### **Supplementary Figure 29.** Comparison of TMS depression targeting quality (Euclidean distance to ground-truth depression targets) between strict censoring and no censoring, for the two-run FD setup. Here strict censoring refers to FD threshold of 0.08 mm (after pseudo-motion filtering) and DVARS threshold of 50. High-motion runs were discarded. Smaller Euclidean distance indicates better quality. TMS depression targets were estimated using the Cluster approach (Cash et al., 2021b). (A) TMS depression targeting quality (Euclidean distance) for the two censoring approaches across 10 motion bins. Each cell in the grid indicates a motion bin. Each dot represents the mean Euclidean distance across 50 simulated sessions of a participant in that motion bin. P-values were computed using the two-sided paired-sampled t-test. "*" indicates statistical significance after multiple comparisons correction with FDR q < 0.05. Under strict censoring, no runs remained for the bottom-right three motion bins because all fMRI runs were discarded under this criterion. (B) TMS depression targeting quality across all motion bins for both censoring strategies. There are seven pairs of dots, corresponding to the seven motion bins where comparisons could be made between strict censoring and no censoring. P-value was computed using a linear mixed effects model (p = 0.03). "*" indicates statistical significance after multiple comparisons correction with FDR q < 0.05. There are three “×” corresponding to the mean Euclidean distance of the 3 bottom-right bins under the no censoring condition. The × were not included in the boxplots. For each box plot, the horizontal line indicates the median. The bottom and top edges of the box indicate the 25th and 75th percentiles, respectively. The outliers are defined as data points beyond 1.5 times the interquartile range. The whiskers extend to the most extreme data points not considered outliers.

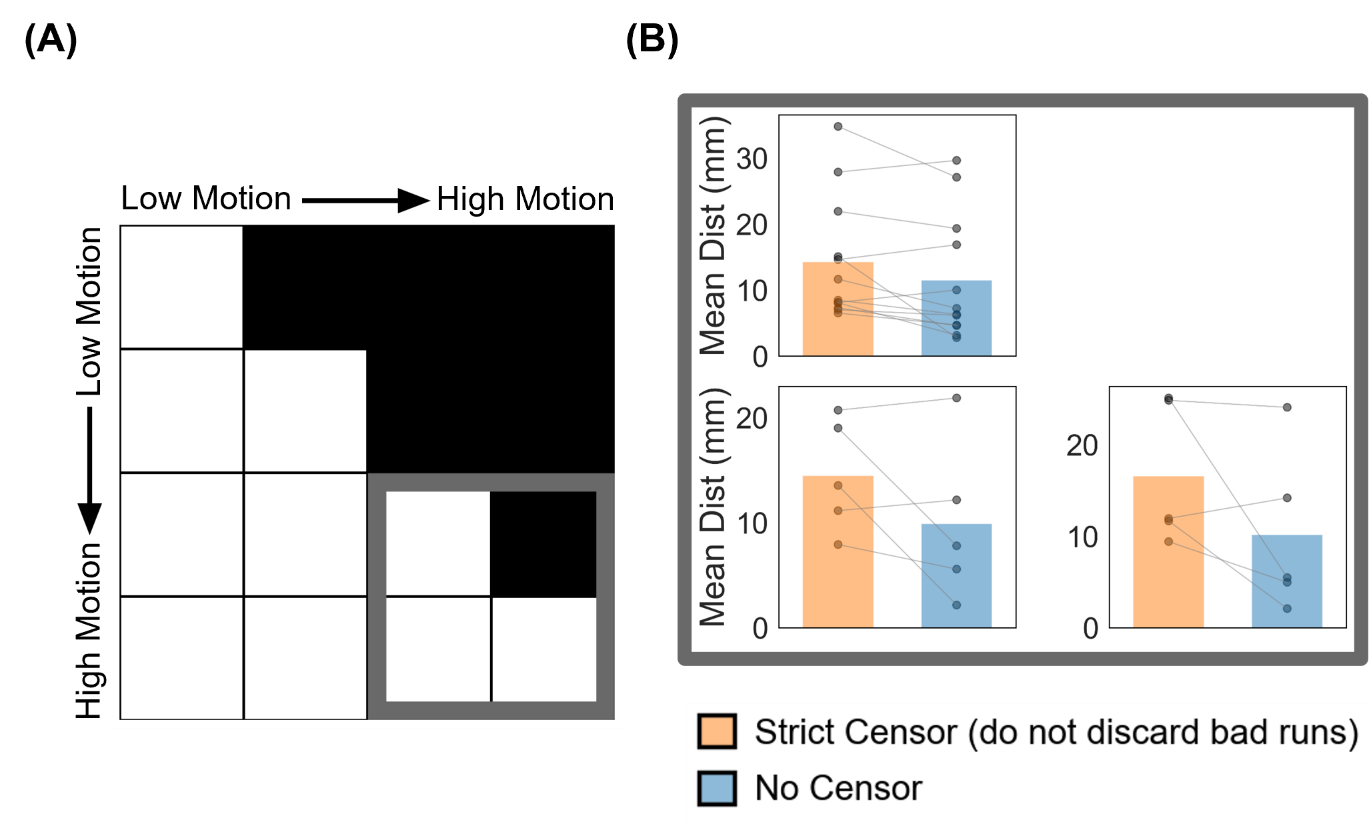

##### **Supplementary Figure 30.** Comparison of TMS depression targeting quality (Euclidean distance from ground-truth targets) between strict censoring and no censoring for the high-motion bins in the two-run FD setup. Here strict censoring refers to FD threshold of 0.08 mm (after pseudo-motion filtering) and DVARS threshold of 50. High-motion runs were not discarded for strict censoring, so a comparison could be made with no censoring. Smaller Euclidean distance indicates better quality. TMS depression targets were estimated using the Cluster approach (Cash et al., 2021b). (A) Schematic highlighting the three high-motion bins with thick gray outlines. (B) TMS targeting quality (Euclidean distance) for the two censoring approaches across the 3 high-motion bins. Each cell in the grid indicates a motion bin. Each dot represents the mean Euclidean distance across 50 simulated sessions for a participant in that motion bin. P-values were computed using the two-sided paired-sampled t-test. "*" indicates statistical significance after multiple comparisons correction with FDR q < 0.05.

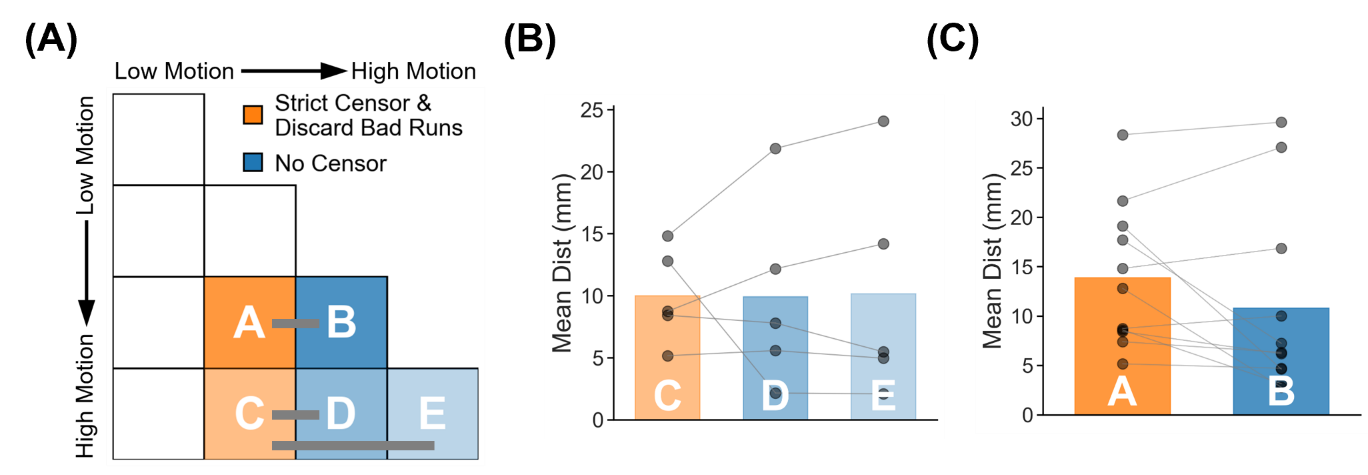

##### **Supplementary Figure 31.** TMS depression target quality (Euclidean distance to ground-truth targets) for two high-motion runs with no censoring vs mixed-motion runs with strict censoring. Here mixed-motion runs refer to sessions with one high-motion and one low-motion runs, so the high-motion run will be discarded under strict censoring. Strict censoring refers to FD threshold of 0.08 mm (after pseudo-motion filtering) and DVARS threshold of 50. Lower Euclidean distance indicates better quality. TMS depression targets were estimated using the Cluster approach (Cash et al., 2021b). (A) Schematic illustrating the comparison. Motion bins A and B were compared, while motion bins C, D and E were compared. (B) Comparison of motion bins C vs D, and C vs E. (C) Comparison of motion bins A vs B. For both panels (B) and (C), each dot represents the mean Euclidean distance across 50 simulated sessions for a participant in that motion bin. P-values were computed using the two-sided paired-sampled t-test. Each statistical test was performed using only participants common to both motion bins. "*" indicates statistical significance after multiple comparisons correction with FDR q < 0.05.

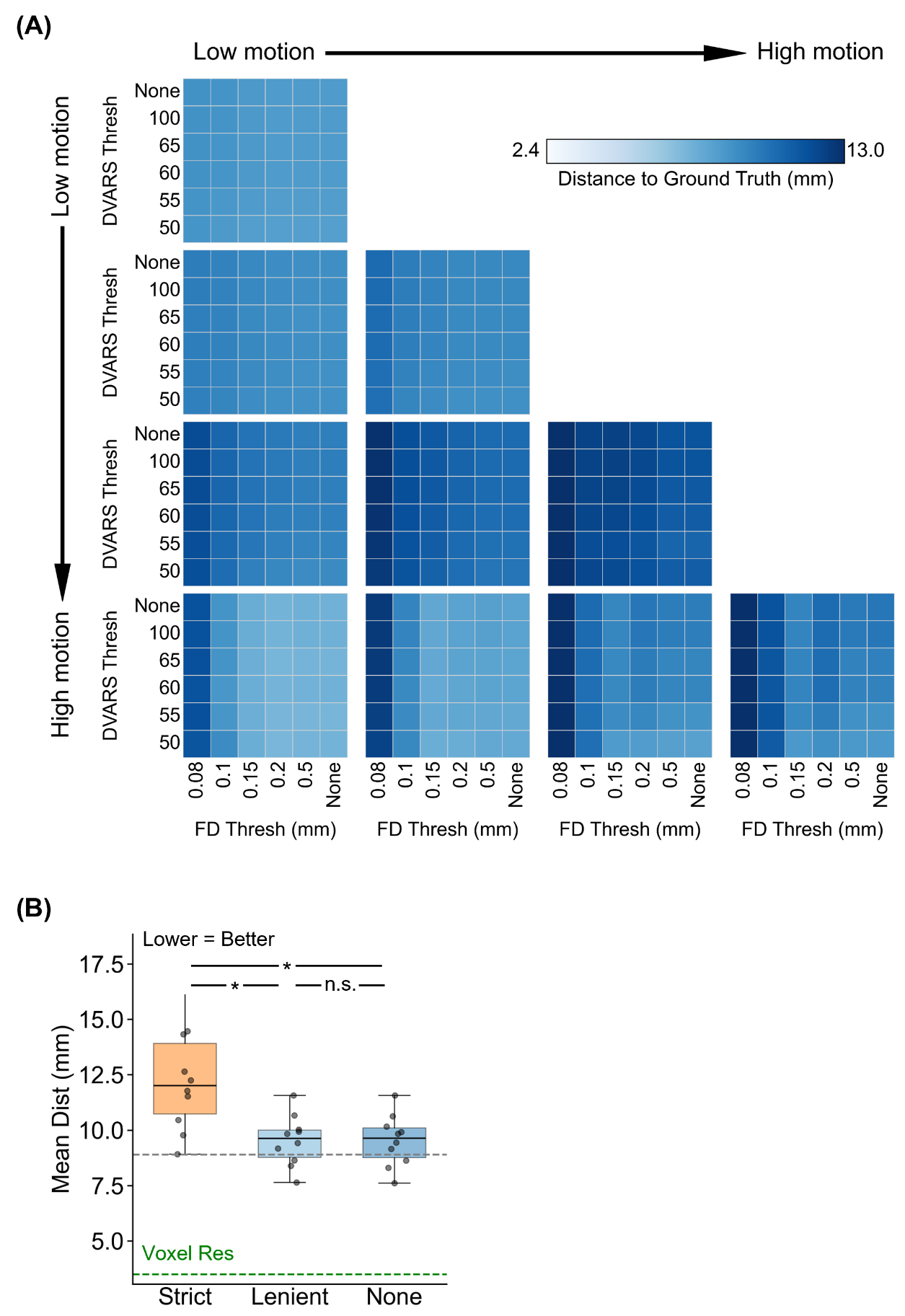

##### **Supplementary Figure 32.** Comparison of TMS depression target quality (Euclidean distance from ground-truth targets) across varying FD and DVARS thresholds for the two-run FD setup. For this analysis, high-motion runs were not discarded. Lower Euclidean distance indicates better quality. TMS depression targets were estimated using the Cluster approach (Cash et al., 2021b). (A) TMS depression target quality (Euclidean distance) for different combinations of FD and DVARS thresholds for 10 motion bins. Each cell in the grid indicates a motion bin. For each motion bin, there are 36 small cells corresponding to different FD and DVARS thresholds. (B) TMS depression target quality for strict censoring, lenient censoring and no censoring. Here strict censoring refers to FD threshold of 0.08 mm (after pseudo-motion filtering) and DVARS threshold of 50. Lenient censoring refers to FD threshold of 0.5 mm and DVARS threshold of 100. Each dot represents the mean Euclidean distance for a motion bin. The gray dashed line is the “noise ceiling”, achieved by picking the best combination of FD and DVARS threshold for each motion bin, and computing the median across motion bins. P-values were computed using a linear mixed effects model (p = 1.0e-9 for strict vs. none, p = 1.1e-9 for strict vs. lenient, p = 0.99 for lenient vs. none). "*" indicates statistical significance after multiple comparisons correction with FDR q < 0.05 and "n.s." indicates not significant after FDR correction. For each box plot, the horizontal line indicates the median across all motion bins. The bottom and top edges of the box indicate the 25th and 75th percentiles, respectively. The outliers are defined as data points beyond 1.5 times the interquartile range. The whiskers extend to the most extreme data points not considered outliers.

#### **Two runs FDrms for tree-based (Kong2026) TMS depression targets**

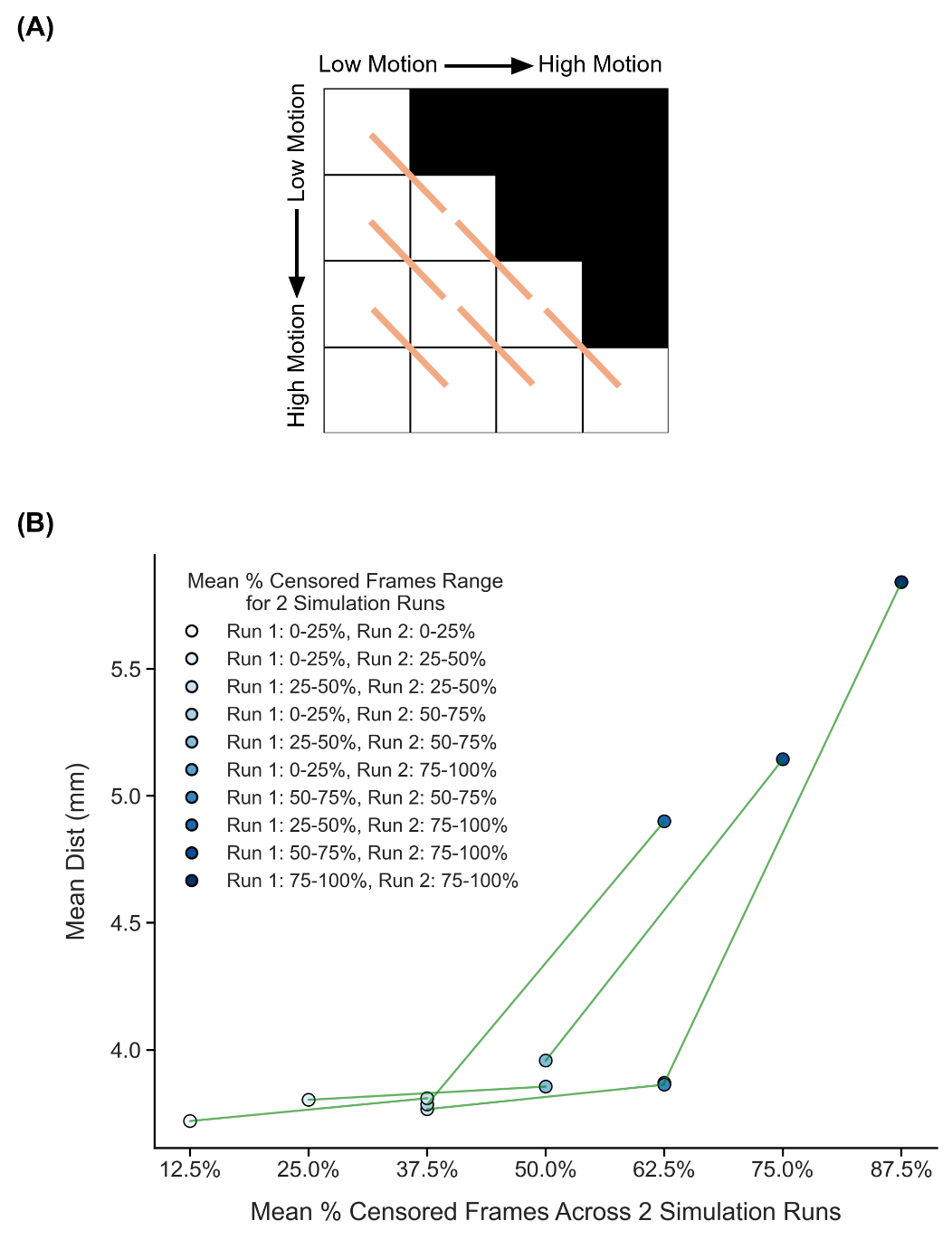

##### **Supplementary Figure 33.** Comparison of TMS depression target quality (Euclidean distance from ground-truth targets) across diagonally adjacent motion bins under strict censoring, for the two-run FDrms setup. Here strict censoring refers to FDrms threshold of 0.2 mm (without pseudo-motion filtering) and DVARS threshold of 50. High-motion runs were not discarded to allow comparison with low-motion runs. Smaller Euclidean distance to ground-truth target indicates better quality. TMS depression targets were estimated using the tree-algorithm (Kong et al., 2026a). (A) Schematic of comparisons across diagonally adjacent motion bins. Each orange line indicates a single comparison. (B) Relationship between TMS depression target quality (Euclidean distance) and motion. Each circle indicates the mean Euclidean distance across simulation runs of all participants in a motion bin. Each line indicates a single comparison between two adjacent motion bins. P-values were computed using the two-sided paired-sampled t-test using only participants common to both motion bins. An overall statistical test combining all comparisons was conducted using a linear mixed effects model (p = 8.7e-6).

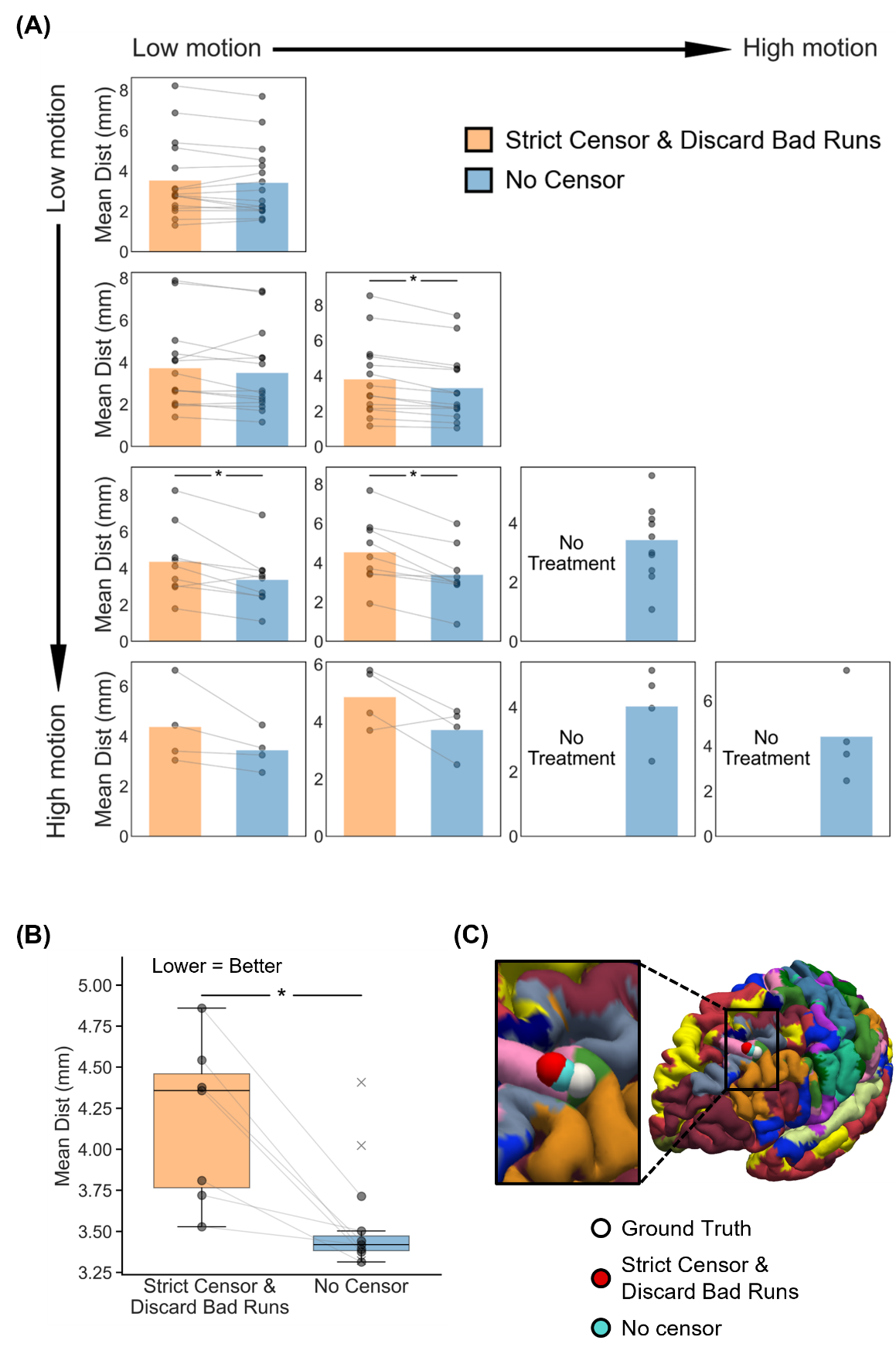

##### **Supplementary Figure 34.** Comparison of TMS depression targeting quality (Euclidean distance to ground-truth depression targets) between strict censoring and no censoring, for the two-run FDrms setup. Here strict censoring refers to FDrms threshold of 0.2 mm (without pseudo-motion filtering) and DVARS threshold of 50. High-motion runs were discarded. Smaller Euclidean distance indicates better quality. TMS depression targets were estimated using the tree-based algorithm (Kong et al., 2026a). (A) TMS depression targeting quality (Euclidean distance) for the two censoring approaches across 10 motion bins. Each cell in the grid indicates a motion bin. Each dot represents the mean Euclidean distance across 50 simulated sessions of a participant in that motion bin. P-values were computed using the two-sided paired-sampled t-test. "*" indicates statistical significance after multiple comparisons correction with FDR q < 0.05. Under strict censoring, no runs remained for the bottom-right three motion bins because all fMRI runs were discarded under this criterion. (B) TMS depression targeting quality across all motion bins for both censoring strategies. There are seven pairs of dots, corresponding to the seven motion bins where comparisons could be made between strict censoring and no censoring. P-value was computed using a linear mixed effects model (p = 8e-5). "*" indicates statistical significance after multiple comparisons correction with FDR q < 0.05. There are three “×” corresponding to the mean Euclidean distance of the 3 bottom-right bins under the no censoring condition. The × were not included in the boxplots. For each box plot, the horizontal line indicates the median. The bottom and top edges of the box indicate the 25th and 75th percentiles, respectively. The outliers are defined as data points beyond 1.5 times the interquartile range. The whiskers extend to the most extreme data points not considered outliers. (C) Visualization of TMS depression targets in a representative participant from the highest motion bin where the strict censoring had results.

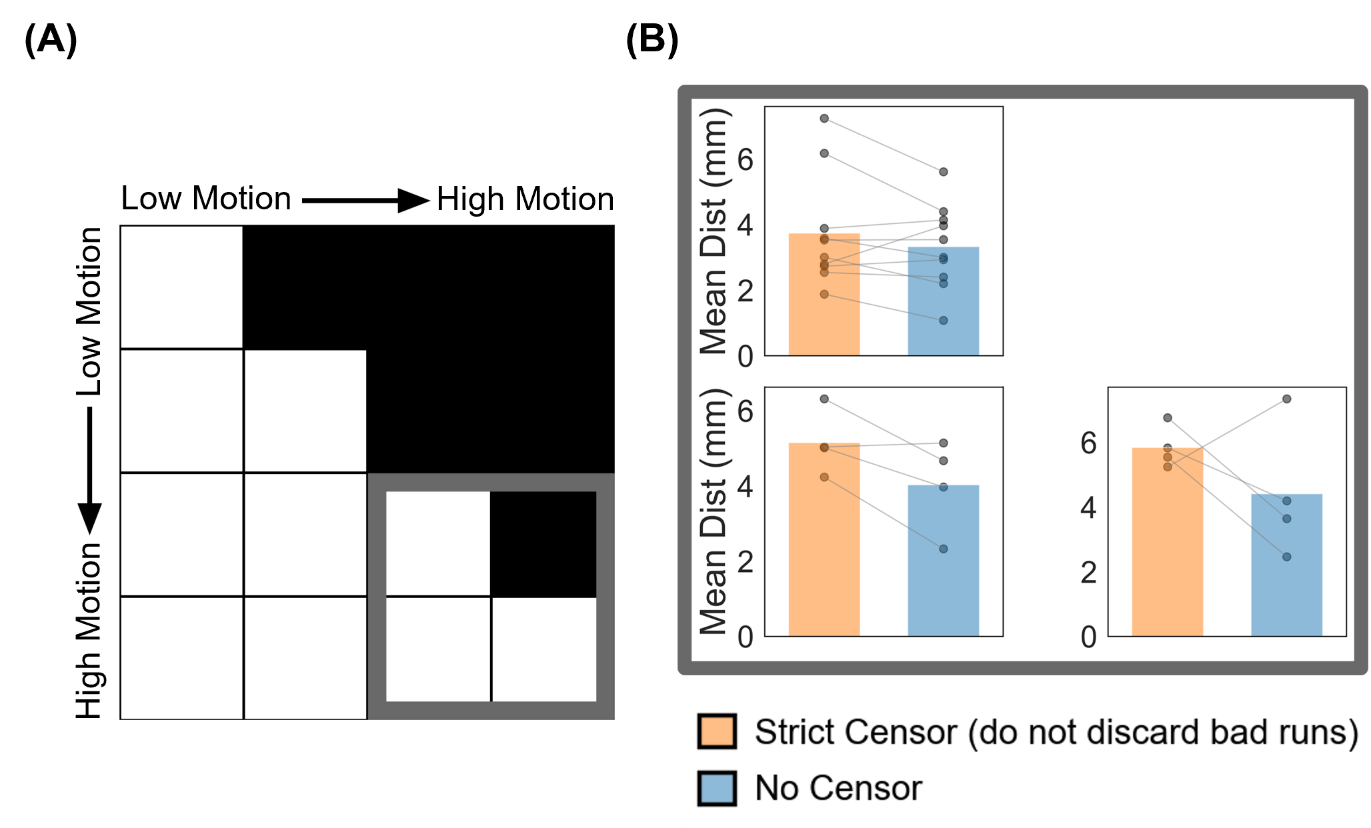

##### **Supplementary Figure 35.** Comparison of TMS depression targeting quality (Euclidean distance from ground-truth targets) between strict censoring and no censoring for the high-motion bins in the two-run FDrms setup. Here strict censoring refers to FDrms threshold of 0.2 mm (without pseudo-motion filtering) and DVARS threshold of 50. High-motion runs were not discarded for strict censoring, so a comparison could be made with no censoring. Smaller Euclidean distance indicates better quality. TMS depression targets were estimated using the tree-algorithm (Kong et al., 2026a). (A) Schematic highlighting the three high-motion bins with thick gray outlines. (B) TMS targeting quality (Euclidean distance) for the two censoring approaches across the 3 high-motion bins. Each cell in the grid indicates a motion bin. Each dot represents the mean Euclidean distance across 50 simulated sessions for a participant in that motion bin. P-values were computed using the two-sided paired-sampled t-test. "*" indicates statistical significance after multiple comparisons correction with FDR q < 0.05.

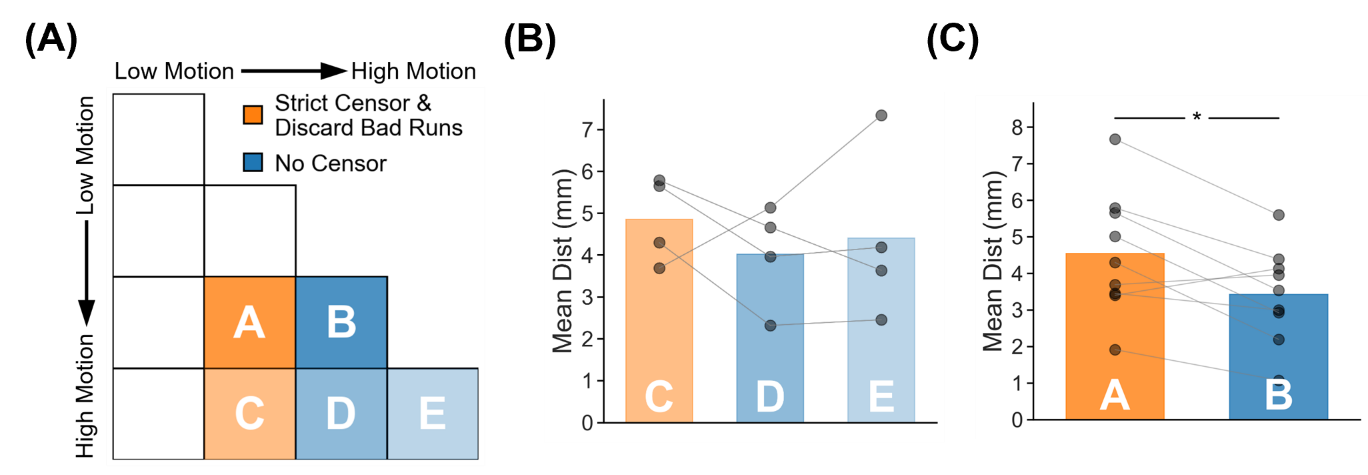

##### **Supplementary Figure 36.** TMS depression target quality (Euclidean distance to ground-truth targets) for two high-motion runs with no censoring vs mixed-motion runs with strict censoring. Here mixed-motion runs refer to sessions with one high-motion and one low-motion runs, so the high-motion run will be discarded under strict censoring. Strict censoring refers to FDrms threshold of 0.2 mm (without pseudo-motion filtering) and DVARS threshold of 50. Lower Euclidean distance indicates better quality. TMS depression targets were estimated using the tree-algorithm (Kong et al., 2026a). (A) Schematic illustrating the comparison. Motion bins A and B were compared, while motion bins C, D and E were compared. (B) Comparison of motion bins C vs D, and C vs E. (C) Comparison of motion bins A vs B. For both panels (B) and (C), each dot represents the mean Euclidean distance across 50 simulated sessions for a participant in that motion bin. P-values were computed using the two-sided paired-sampled t-test. Each statistical test was performed using only participants common to both motion bins. "*" indicates statistical significance after multiple comparisons correction with FDR q < 0.05.

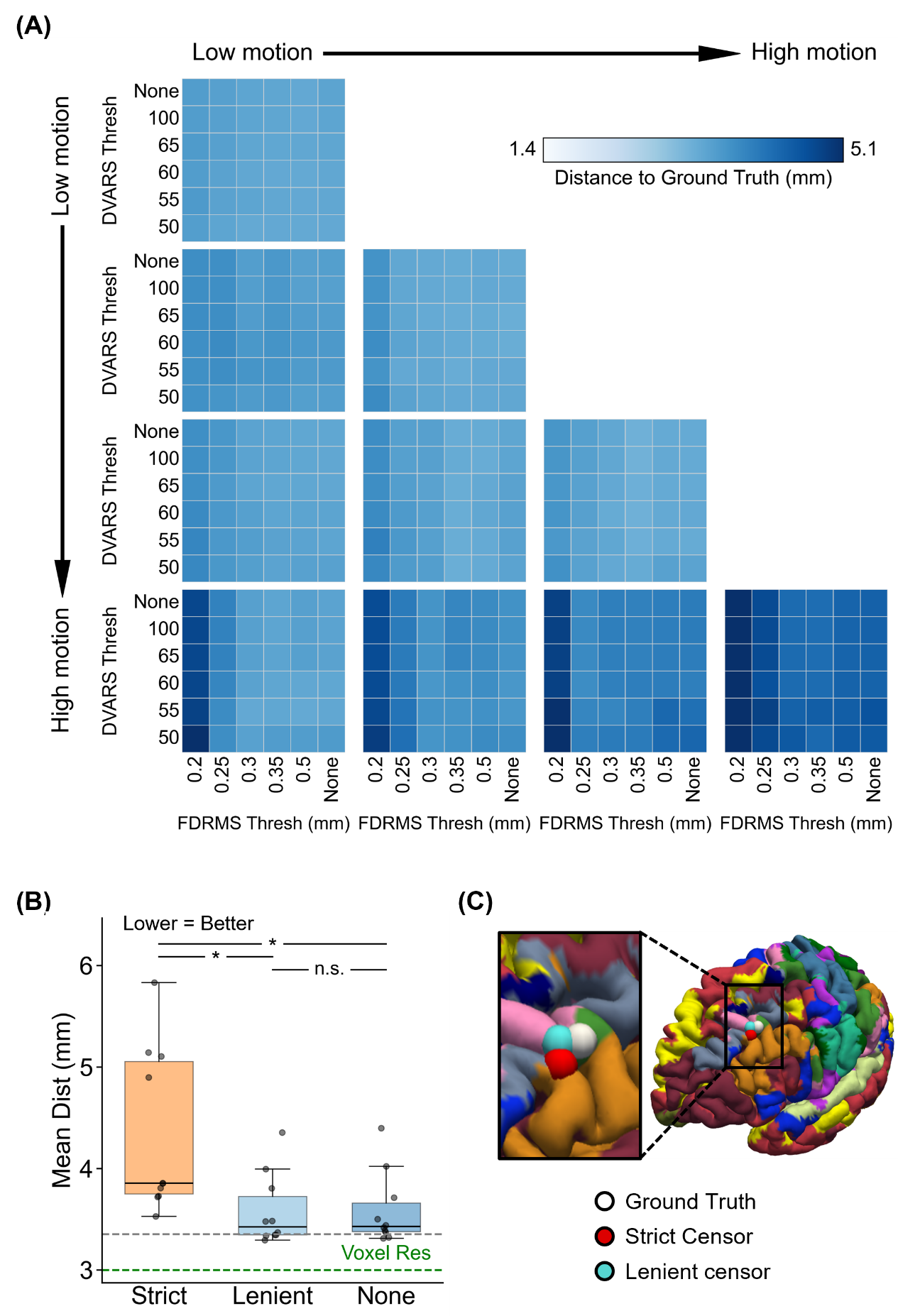

##### **Supplementary Figure 37.** Comparison of TMS depression target quality (Euclidean distance from ground-truth targets) across varying FDrms and DVARS thresholds for the two-run FDrms setup. For this analysis, high-motion runs were not discarded. Lower Euclidean distance indicates better quality. TMS depression targets were estimated using the tree-algorithm (Kong et al., 2026a). (A) TMS depression target quality (Euclidean distance) for different combinations of FDrms and DVARS thresholds for 10 motion bins. Each cell in the grid indicates a motion bin. For each motion bin, there are 36 small cells corresponding to different FDrms and DVARS thresholds. (B) TMS depression target quality for strict censoring, lenient censoring and no censoring. Here strict censoring refers to FDrms threshold of 0.2 mm (without pseudo-motion filtering) and DVARS threshold of 50. Lenient censoring refers to FDrms threshold of 0.5 mm and DVARS threshold of 100. Each dot represents the mean Euclidean distance for a motion bin. The gray dashed line is the “noise ceiling”, achieved by picking the best combination of FDrms and DVARS threshold for each motion bin, and computing the median across motion bins. P-values were computed using a linear mixed effects model (p = 5.7e-14 for strict vs. none, p = 9e-15 for strict vs. lenient, p = 0.8 for lenient vs. none). "*" indicates statistical significance after multiple comparisons correction with FDR q < 0.05 and "n.s." indicates not significant after FDR correction. For each box plot, the horizontal line indicates the median across all motion bins. The bottom and top edges of the box indicate the 25th and 75th percentiles, respectively. The outliers are defined as data points beyond 1.5 times the interquartile range. The whiskers extend to the most extreme data points not considered outliers. (C) Visualization of TMS depression targets in a representative participant from the highest motion bin.

#### **One run FDrms for tree-based (Kong2026) TMS depression targets**

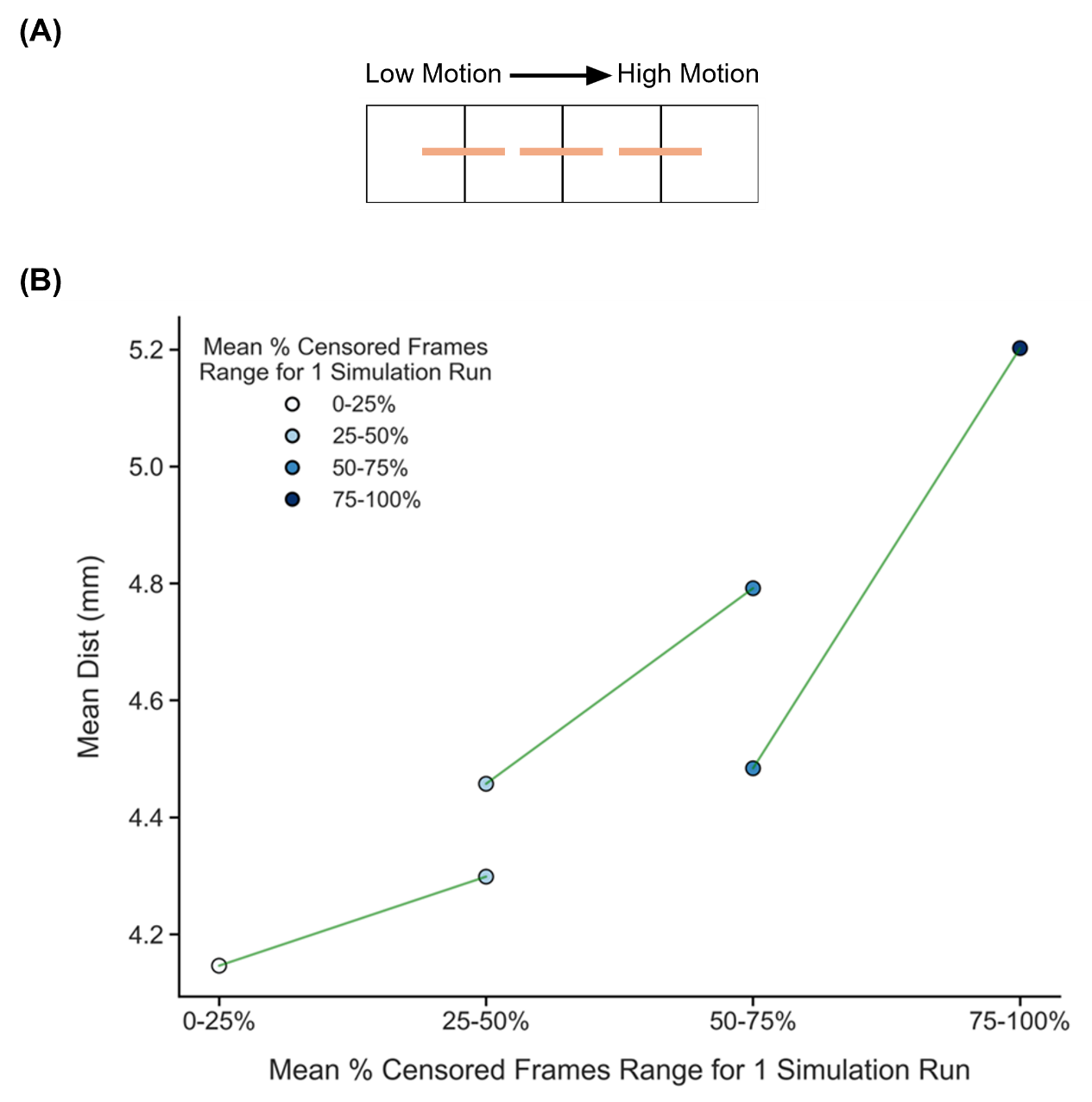

##### **Supplementary Figure 38.** Comparison of TMS depression target quality (Euclidean distance from ground-truth targets) across adjacent motion bins under strict censoring, for the one-run FDrms setup. Here strict censoring refers to FDrms threshold of 0.2 mm (without pseudo-motion filtering) and DVARS threshold of 50. High-motion runs were not discarded to allow comparison with low-motion runs. Smaller Euclidean distance to ground-truth target indicates better quality. TMS depression targets were estimated using the tree-algorithm (Kong et al., 2026a). (A) Schematic of comparisons across adjacent motion bins. Each orange line indicates a single comparison. (B) Relationship between TMS depression target quality (Euclidean distance) and motion. Each circle indicates the mean Euclidean distance across simulation runs of all participants in a motion bin. Each line indicates a single comparison between two adjacent motion bins. P-values were computed using the two-sided paired-sampled t-test using only participants common to both motion bins. An overall statistical test combining all comparisons was conducted using a linear mixed effects model (p = 3.6e-3).

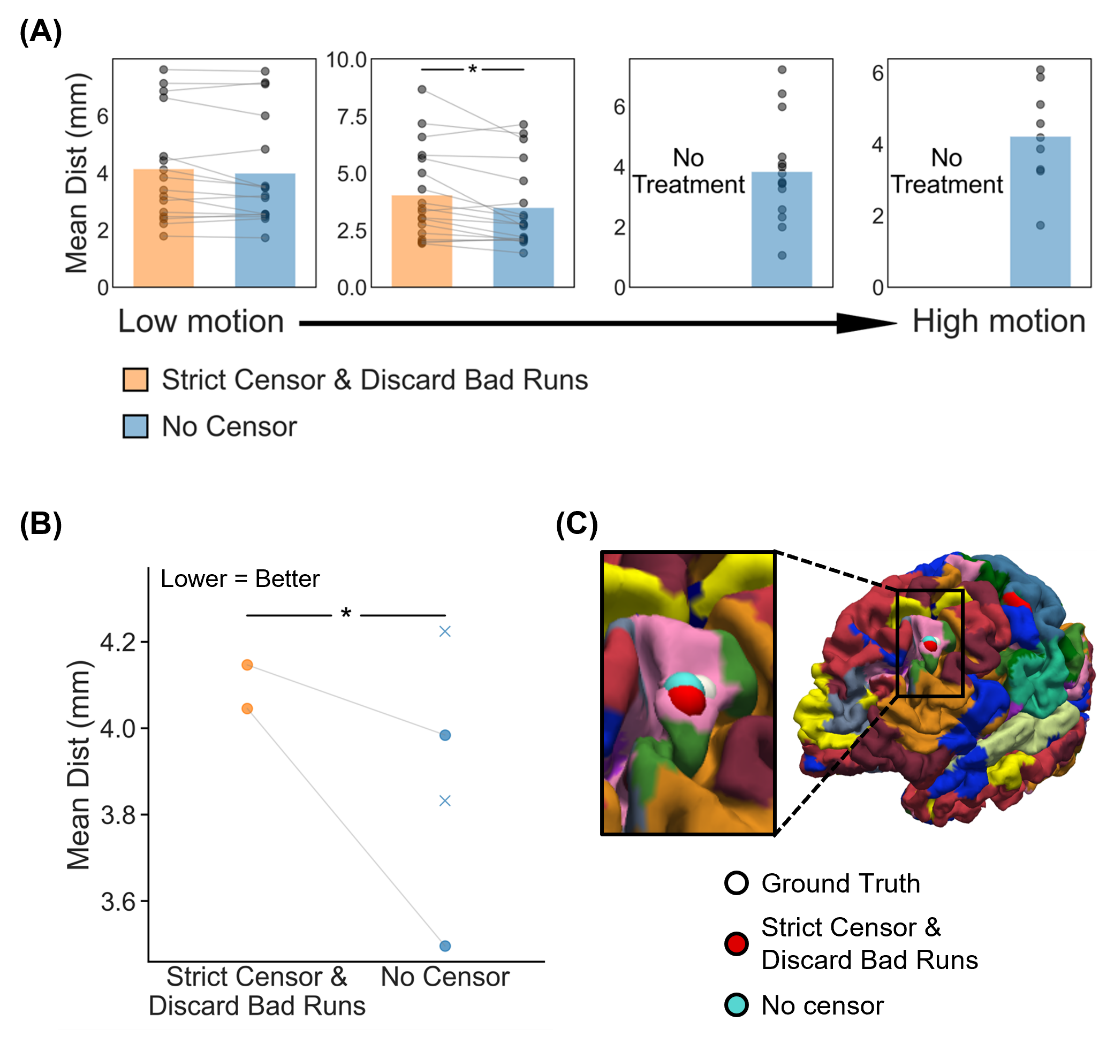

##### **Supplementary Figure 39.** Comparison of TMS depression targeting quality (Euclidean distance to ground-truth depression targets) between strict censoring and no censoring, for the one-run FDrms setup. Here strict censoring refers to FDrms threshold of 0.2 mm (without pseudo-motion filtering) and DVARS threshold of 50. High-motion runs were discarded. Smaller Euclidean distance indicates better quality. TMS depression targets were estimated using the tree-based algorithm (Kong et al., 2026a). (A) TMS depression targeting quality (Euclidean distance) for the two censoring approaches across 4 motion bins. Each cell indicates a motion bin. Each dot represents the mean Euclidean distance across 50 simulated sessions of a participant in that motion bin. P-values were computed using the two-sided paired-sampled t-test. "*" indicates statistical significance after multiple comparisons correction with FDR q < 0.05. Under strict censoring, no runs remained for the high-motion bins because all fMRI runs were discarded under this criterion. (B) TMS depression targeting quality across all motion bins for both censoring strategies. There are two pairs of dots, corresponding to the two motion bins where comparisons could be made between strict censoring and no censoring. P-value was computed using a linear mixed effects model (p = 2.2e-3). "*" indicates statistical significance after multiple comparisons correction with FDR q < 0.05. There are two “×” corresponding to the mean Euclidean distance of the high-motion bins under the no censoring condition. The × were not included in the boxplots. For each box plot, the horizontal line indicates the median. The bottom and top edges of the box indicate the 25th and 75th percentiles, respectively. The outliers are defined as data points beyond 1.5 times the interquartile range. The whiskers extend to the most extreme data points not considered outliers. (C) Visualization of TMS depression targets in a representative participant from the highest motion bin where the strict censoring had results.

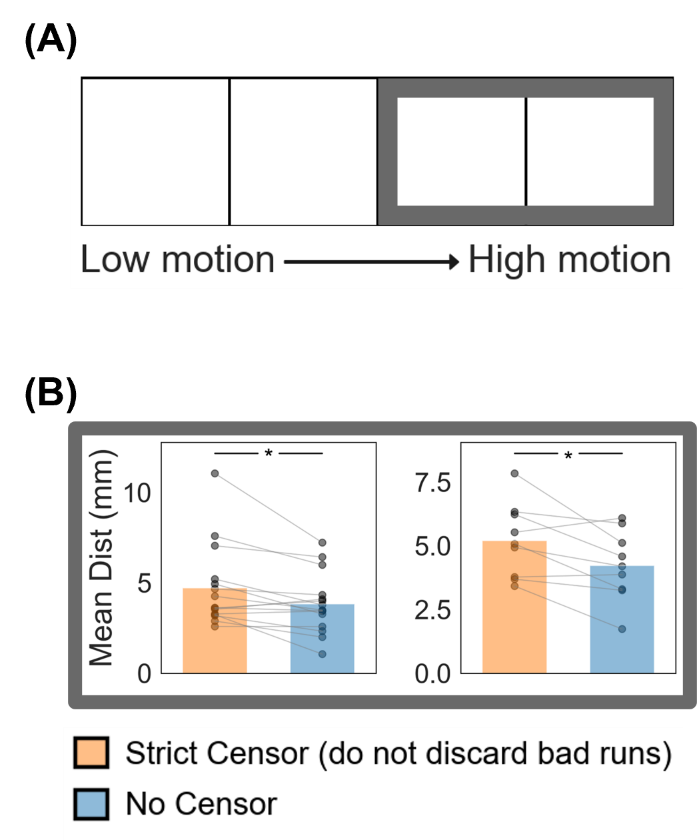

##### **Supplementary Figure 40.** Comparison of TMS depression targeting quality (Euclidean distance from ground-truth targets) between strict censoring and no censoring for the high-motion bins in the one-run FDrms setup. Here strict censoring refers to FDrms threshold of 0.2 mm (without pseudo-motion filtering) and DVARS threshold of 50. High-motion runs were not discarded for strict censoring, so a comparison could be made with no censoring. Smaller Euclidean distance indicates better quality. TMS depression targets were estimated using the tree-algorithm (Kong et al., 2026a). (A) Schematic highlighting the two high-motion bins with thick gray outlines. (B) TMS targeting quality (Euclidean distance) for the two censoring approaches across the two high-motion bins. Each cell indicates a motion bin. Each dot represents the mean Euclidean distance across 50 simulated sessions for a participant in that motion bin. P-values were computed using the two-sided paired-sampled t-test. "*" indicates statistical significance after multiple comparisons correction with FDR q < 0.05.

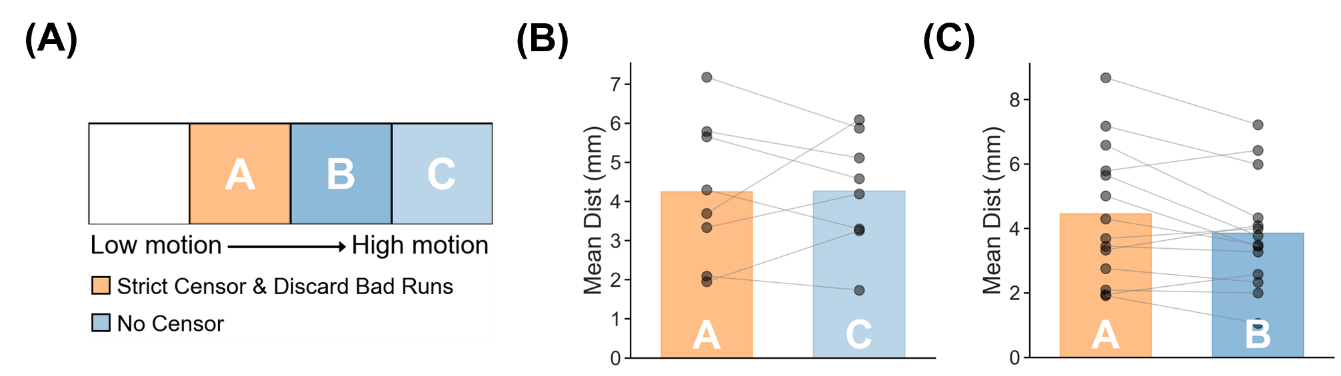

##### **Supplementary Figure 41.** TMS depression target quality (Euclidean distance to ground-truth targets) for one high-motion run with no censoring vs one low-motion run with strict censoring. Strict censoring refers to FDrms threshold of 0.2 mm (without pseudo-motion filtering) and DVARS threshold of 50. Lower Euclidean distance indicates better quality. TMS depression targets were estimated using the tree-algorithm (Kong et al., 2026a). (A) Schematic illustrating the comparison. (B) Comparison of motion bins A vs C. (C) Comparison of motion bins A vs B. For both panels (B) and (C), each dot represents the mean Euclidean distance across 50 simulated sessions for a participant in that motion bin. P-values were computed using the two-sided paired-sampled t-test. Each statistical test was performed using only participants common to both motion bins. "*" indicates statistical significance after multiple comparisons correction with FDR q < 0.05.

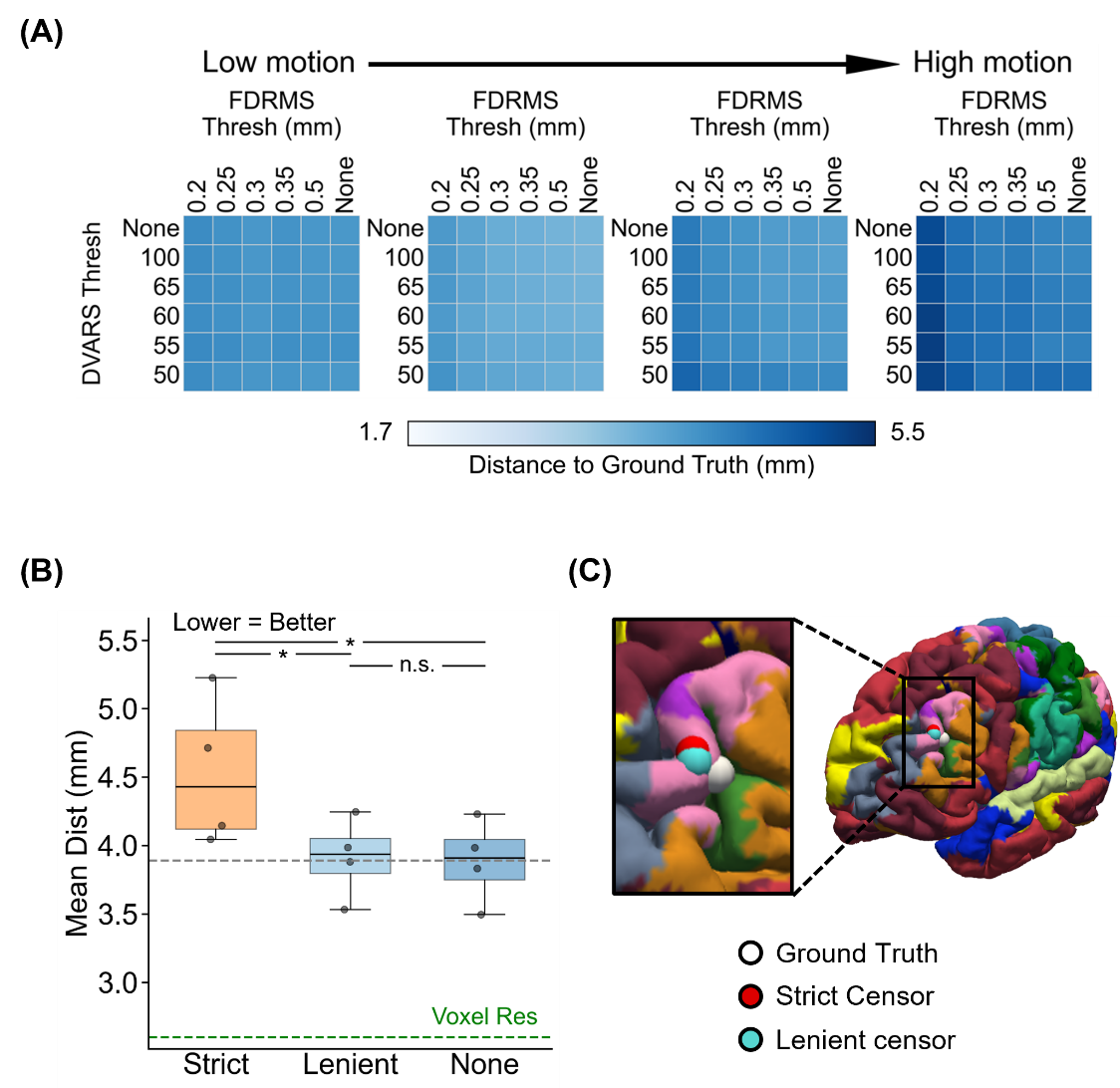

##### **Supplementary Figure 42.** Comparison of TMS depression target quality (Euclidean distance from ground-truth targets) across varying FDrms and DVARS thresholds for the one-run FDrms setup. For this analysis, high-motion runs were not discarded. Lower Euclidean distance indicates better quality. TMS depression targets were estimated using the tree-algorithm (Kong et al., 2026a). (A) TMS depression target quality (Euclidean distance) for different combinations of FDrms and DVARS thresholds for 4 motion bins. Each cell indicates a motion bin. For each motion bin, there are 36 small cells corresponding to different FDrms and DVARS thresholds. (B) TMS depression target quality for strict censoring, lenient censoring and no censoring. Here strict censoring refers to FDrms threshold of 0.2 mm (without pseudo-motion filtering) and DVARS threshold of 50. Lenient censoring refers to FDrms threshold of 0.5 mm and DVARS threshold of 100. Each dot represents the mean Euclidean distance for a motion bin. The gray dashed line is the “noise ceiling”, achieved by picking the best combination of FDrms and DVARS threshold for each motion bin, and computing the median across motion bins. P-values were computed using a linear mixed effects model (p = 1.6e-10 for strict vs. none, p = 1.1e-9 for strict vs. lenient, p = 0.77 for lenient vs. none). "*" indicates statistical significance after multiple comparisons correction with FDR q < 0.05 and "n.s." indicates not significant after FDR correction. For each box plot, the horizontal line indicates the median across all motion bins. The bottom and top edges of the box indicate the 25th and 75th percentiles, respectively. The outliers are defined as data points beyond 1.5 times the interquartile range. The whiskers extend to the most extreme data points not considered outliers. (C) Visualization of TMS depression targets in a representative participant from the highest motion bin.

#### **Simultaneous regression (two runs FD) for parcellations**

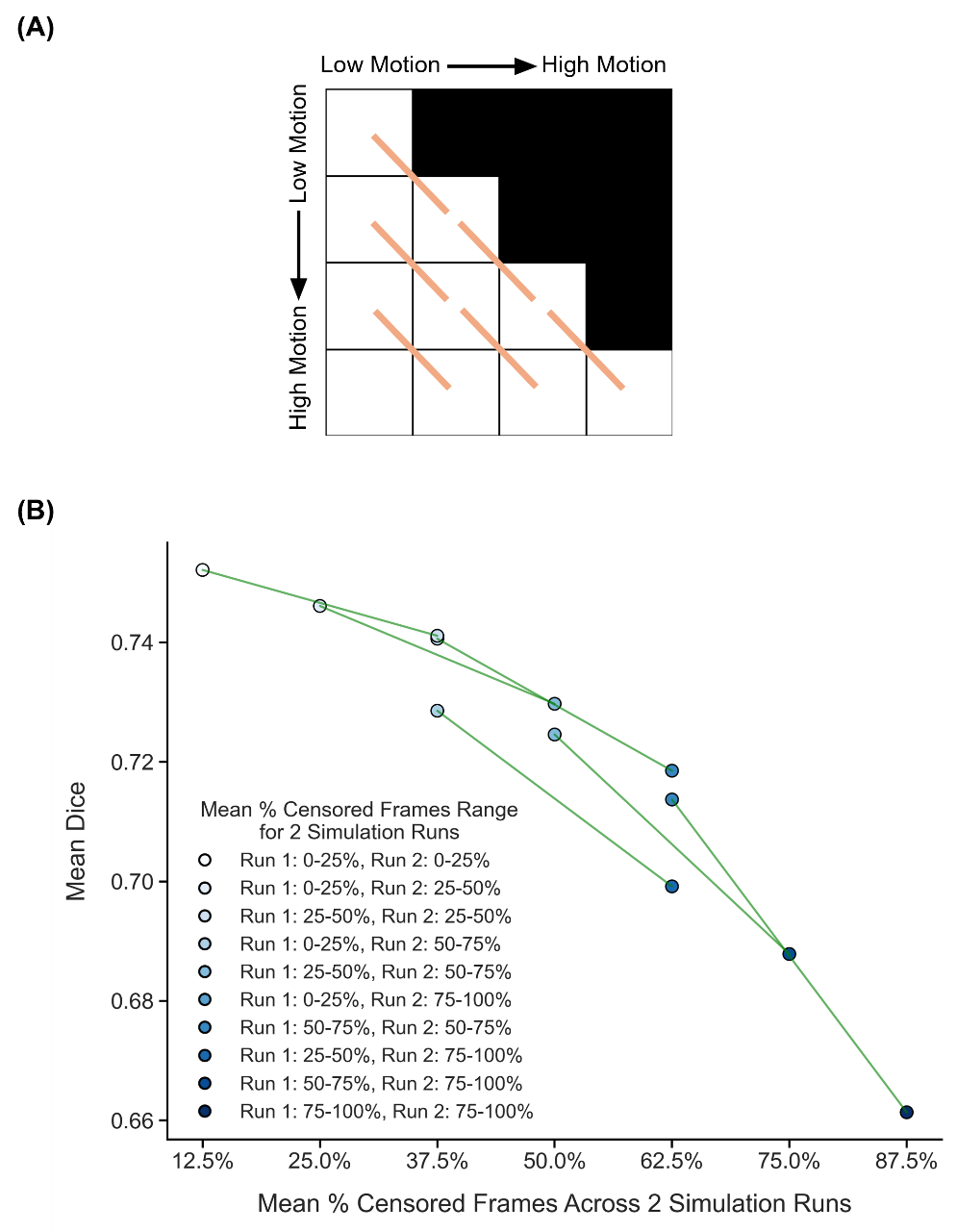

##### **Supplementary Figure 43.** Comparison of parcellation quality (Dice) across diagonally adjacent motion bins under strict censoring for the two-run FD setup, using simultaneous regression of bandpass filters together with nuisance regressors. Here strict censoring refers to FD threshold of 0.08 mm (after pseudo-motion filtering) and DVARS threshold of 50. High-motion runs were not discarded to allow comparison with low-motion runs. Higher Dice indicates better quality. (A) Schematic of comparisons across diagonally adjacent motion bins. Each orange line indicates a single comparison. (B) Relationship between parcellation quality (Dice) and motion. Each circle indicates the mean Dice coefficient across simulation runs of all participants in a motion bin. Each line indicates a single comparison between two adjacent motion bins. P-values were computed using the two-sided paired-sampled t-test using only participants common to both motion bins. An overall statistical test combining all comparisons was conducted using a linear mixed effects model (p = 4.1e-34).

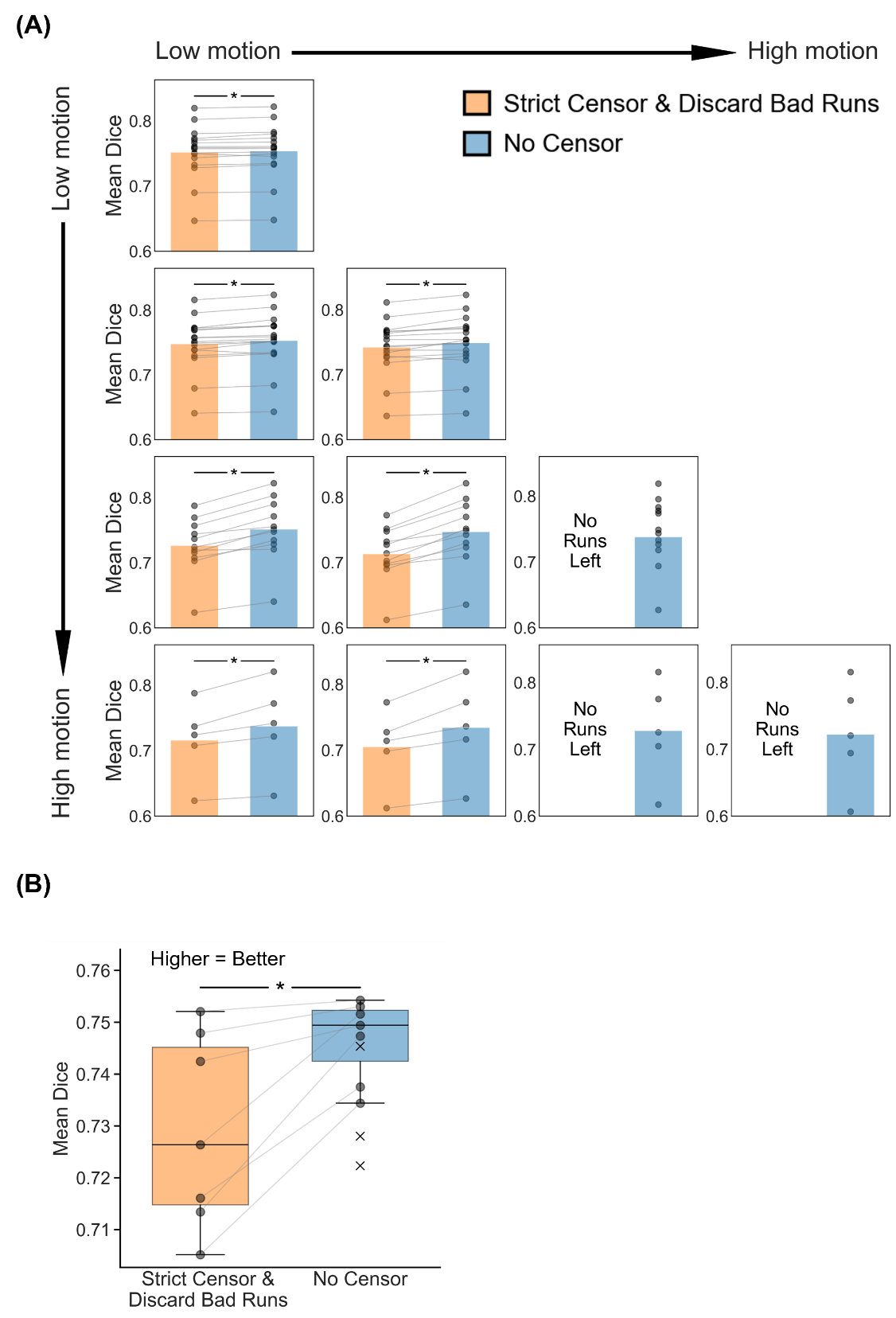

##### **Supplementary Figure 44.** Comparison of parcellation quality (Dice) between strict censoring and no censoring for the two-run FD setup, using simultaneous regression of bandpass filters together with nuisance regressors. Here, strict censoring refers to a FD threshold of 0.08 mm (after pseudo-motion filtering) and a DVARS threshold of 50. High-motion runs were discarded. Higher Dice indicates better quality. (A) Parcellation quality for the two censoring approaches across 10 motion bins. Each cell in the grid indicates a motion bin. Each dot represents the mean Dice coefficient across 50 simulated sessions of a participant in that motion bin. P-values were computed using the two-sided paired-sampled t-test. "*" indicates statistical significance after multiple comparisons correction with FDR q < 0.05. Under strict censoring, no runs remained for the bottom-right three motion bins because all fMRI runs were discarded under this criterion. (B) Parcellation quality across all motion bins for both censoring strategies. There are seven pairs of dots, corresponding to the seven motion bins where comparisons could be made between strict censoring and no censoring. P-value was computed using a linear mixed effects model (p = 1.4e-9). "*" indicates statistical significance after multiple comparisons correction with FDR q < 0.05. There are three “×” corresponding to the Dice coefficient of the three bottom-right bins under the no censoring condition. The × were not included in the boxplots or statistical comparisons between the two censoring strategies. For each box plot, the horizontal line indicates the median. The bottom and top edges of the box indicate the 25th and 75th percentiles, respectively. The outliers are defined as data points beyond 1.5 times the interquartile range. The whiskers extend to the most extreme data points not considered outliers.

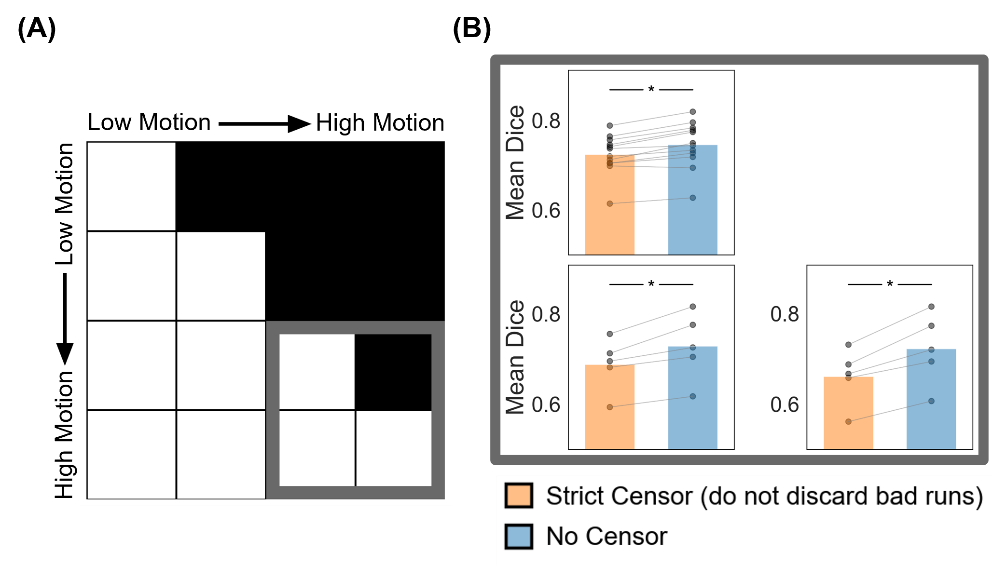

##### **Supplementary Figure 45.** Comparison of parcellation quality (Dice) between strict censoring and no censoring for the high-motion bins in the two-run FD setup, using simultaneous regression of bandpass filters together with nuisance regressors. Here strict censoring refers to FD threshold of 0.08 mm (after pseudo-motion filtering) and DVARS threshold of 50. High-motion runs were not discarded for strict censoring, so a comparison could be made with no censoring (A) Schematic highlighting the three high-motion bins with thick gray outlines. (B) Parcellation quality (Dice) for the two censoring approaches across the 3 high-motion bins. Each cell in the grid indicates a motion bin. Each dot represents the mean Dice coefficient across 50 simulated sessions for a participant in that motion bin. P-values were computed using the two-sided paired-sampled t-test. "*" indicates statistical significance after multiple comparisons correction with FDR q < 0.05.

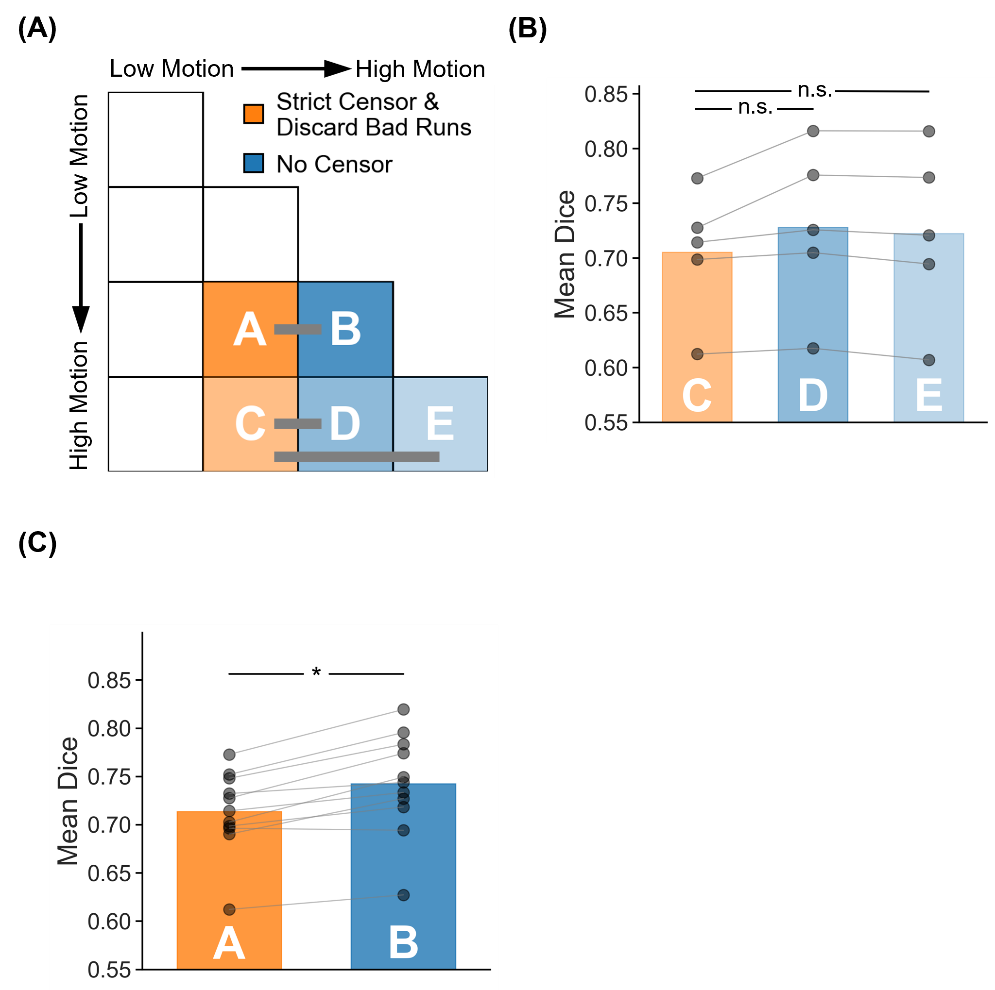

##### **Supplementary Figure 46.** Parcellation quality (Dice) for two high-motion runs with no censoring vs mixed-motion runs with strict censoring, using simultaneous regression of bandpass filters together with nuisance regressors. Here mixed-motion runs refer to sessions with one high-motion and one low-motion run, so the high-motion run will be discarded under strict censoring. Strict censoring refers to FD threshold of 0.08 mm (after pseudo-motion filtering) and DVARS threshold of 50. Higher Dice indicates better quality. (A) Schematic illustrating the comparison. Motion bins A and B were compared, while motion bins C, D and E were compared. (B) Comparison of motion bins C vs D, and C vs E. (C) Comparison of motion bins A vs B. For both panels (B) and (C), each dot represents the mean Dice coefficient across 50 simulated sessions for a participant in that motion bin. P-values were computed using the two-sided paired-sampled t-test. Each statistical test was performed using only participants common to both motion bins. "*" indicates statistical significance after multiple comparisons correction with FDR q < 0.05.

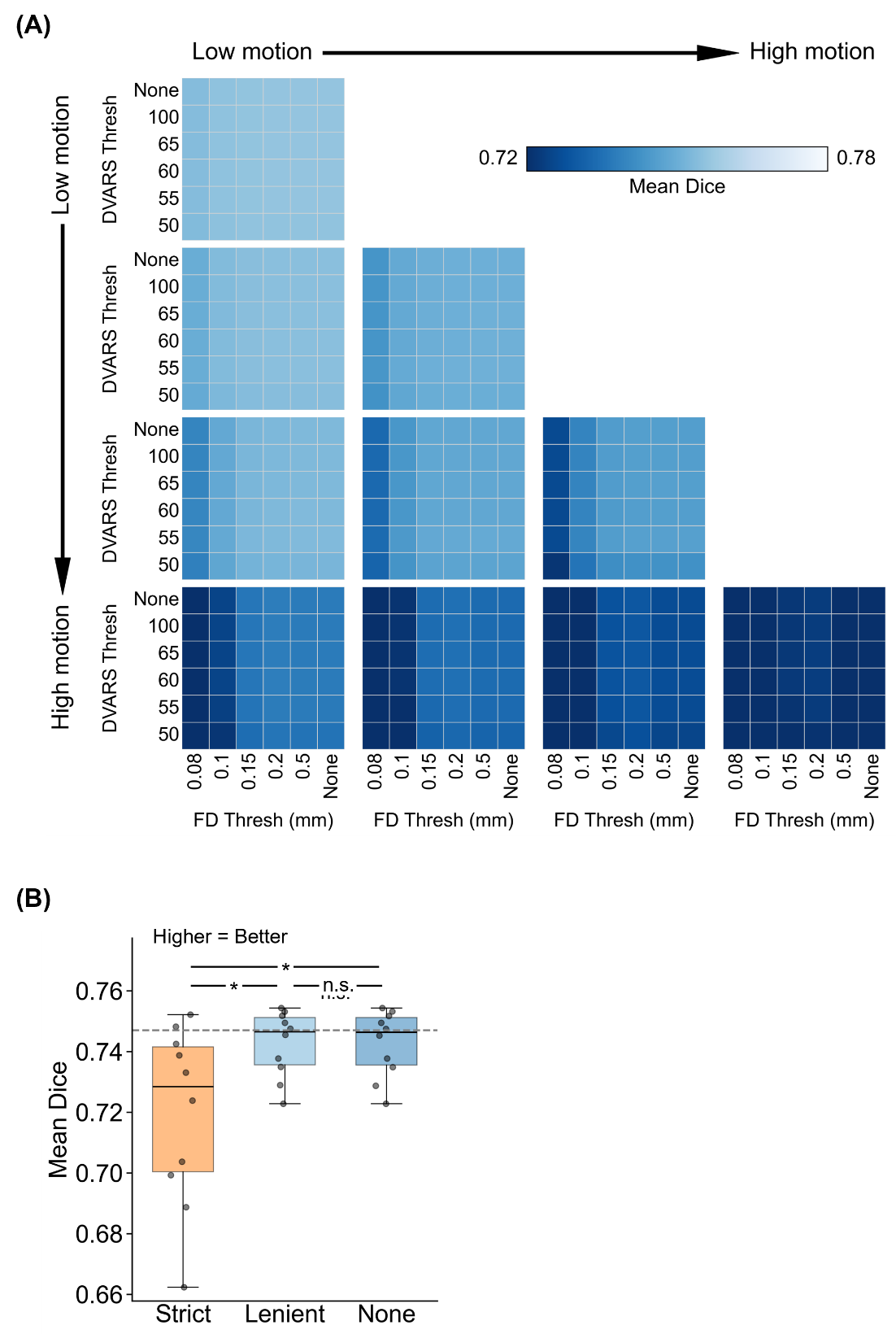

##### **Supplementary Figure 47.** Comparison of parcellation quality (Dice) across varying FD and DVARS thresholds for the two-run FD setup, using simultaneous regression of bandpass filters together with nuisance regressors. For this analysis, high-motion runs were not discarded. Higher Dice indicates better quality. (A) Parcellation quality for different combinations of FD and DVARS thresholds for 10 motion bins. Each cell in the grid indicates a motion bin. For each motion bin, there are 36 small cells corresponding to different FD and DVARS thresholds. (B) Parcellation quality of strict censoring, lenient censoring and no censoring. Here strict censoring refers to FD threshold of 0.08 mm (after pseudo-motion filtering) and DVARS threshold of 50. Lenient censoring refers to FD threshold of 0.5 mm and DVARS threshold of 100. Each dot represents the Dice coefficient for a motion bin. The gray dashed line is the “noise ceiling”, achieved by picking the best combination of FD and DVARS threshold for each motion bin, and computing the median across motion bins. P-values were computed using a linear mixed effects model (p = 3.4e-29 for strict vs. none, p = 2.5e-29 for strict vs. lenient, p = 0.98 for lenient vs. none). "*" indicates statistical significance after multiple comparisons correction with FDR q < 0.05 and "n.s." indicates not significant after FDR correction. For each box plot, the horizontal line indicates the median across all motion bins. The bottom and top edges of the box indicate the 25th and 75th percentiles, respectively. The outliers are defined as data points beyond 1.5 times the interquartile range. The whiskers extend to the most extreme data points not considered outliers.

#### **Simultaneous regression (two runs FD) for tree-based (Kong2026) TMS depression targets**

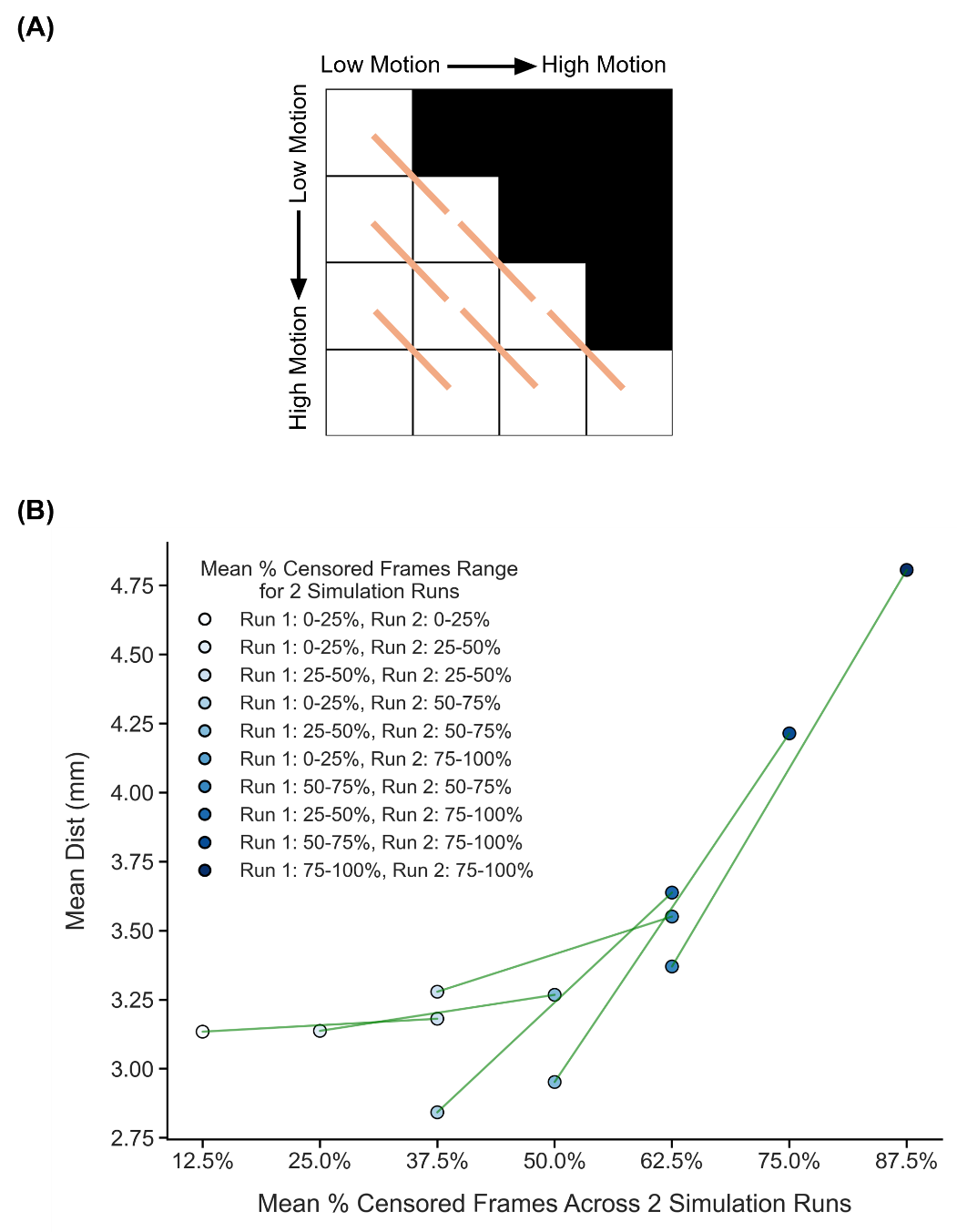

##### **Supplementary Figure 48.** Comparison of TMS depression target quality (Euclidean distance from ground-truth targets) across diagonally adjacent motion bins under strict censoring for the two-run FD setup, using simultaneous regression of bandpass filters together with nuisance regressors. Here strict censoring refers to FD threshold of 0.08 mm (after pseudo-motion filtering) and DVARS threshold of 50. High-motion runs were not discarded to allow comparison with low-motion runs. Smaller Euclidean distance to ground-truth target indicates better quality. TMS depression targets were estimated using the tree-algorithm (Kong et al., 2026a). (A) Schematic of comparisons across diagonally adjacent motion bins. Each orange line indicates a single comparison. (B) Relationship between TMS depression target quality (Euclidean distance) and motion. Each circle indicates the mean Euclidean distance across simulation runs of all participants in a motion bin. Each line indicates a single comparison between two adjacent motion bins. P-values were computed using the two-sided paired-sampled t-test using only participants common to both motion bins. An overall statistical test combining all comparisons was conducted using a linear mixed effects model (p = 6.4e-9).

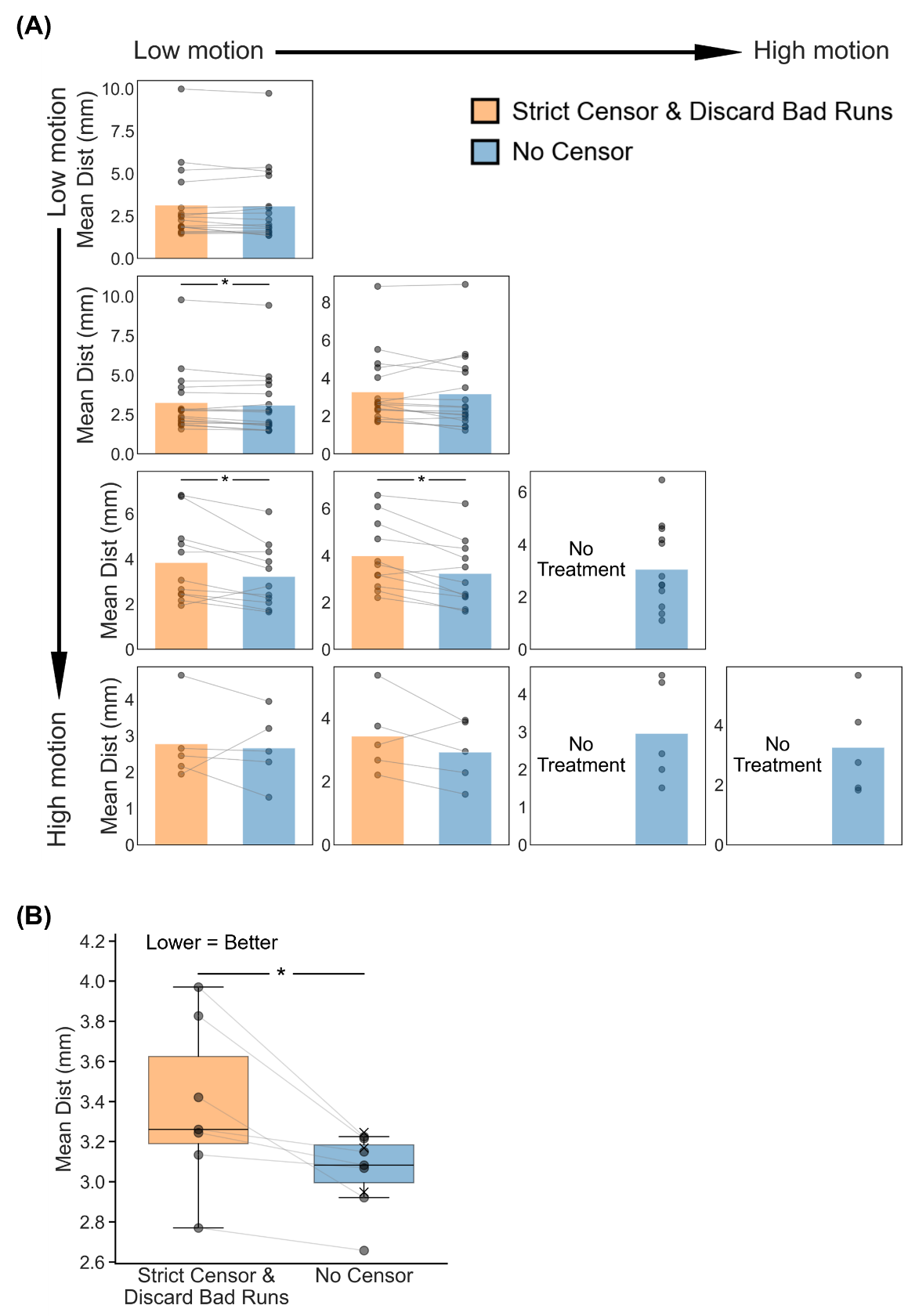

##### **Supplementary Figure 49.** Comparison of TMS depression targeting quality (Euclidean distance to ground-truth depression targets) between strict censoring and no censoring for the two-run FD setup, using simultaneous regression of bandpass filters together with nuisance regressors. Here strict censoring refers to FD threshold of 0.08 mm (after pseudo-motion filtering) and DVARS threshold of 50. High-motion runs were discarded. Smaller Euclidean distance indicates better quality. TMS depression targets were estimated using the tree-based algorithm (Kong et al., 2026a). (A) TMS depression targeting quality (Euclidean distance) for the two censoring approaches across 10 motion bins. Each cell in the grid indicates a motion bin. Each dot represents the mean Euclidean distance across 50 simulated sessions of a participant in that motion bin. P-values were computed using the two-sided paired-sampled t-test. "*" indicates statistical significance after multiple comparisons correction with FDR q < 0.05. Under strict censoring, no runs remained for the bottom-right three motion bins because all fMRI runs were discarded under this criterion. (B) TMS depression targeting quality across all motion bins for both censoring strategies. There are seven pairs of dots, corresponding to the seven motion bins where comparisons could be made between strict censoring and no censoring. P-value was computed using a linear mixed effects model (p = 0.02). "*" indicates statistical significance after multiple comparisons correction with FDR q < 0.05. There are three “×” corresponding to the mean Euclidean distance of the 3 bottom-right bins under the no censoring condition. The × were not included in the boxplots or statistical comparisons between the two censoring strategies. For each box plot, the horizontal line indicates the median. The bottom and top edges of the box indicate the 25th and 75th percentiles, respectively. The outliers are defined as data points beyond 1.5 times the interquartile range. The whiskers extend to the most extreme data points not considered outliers.

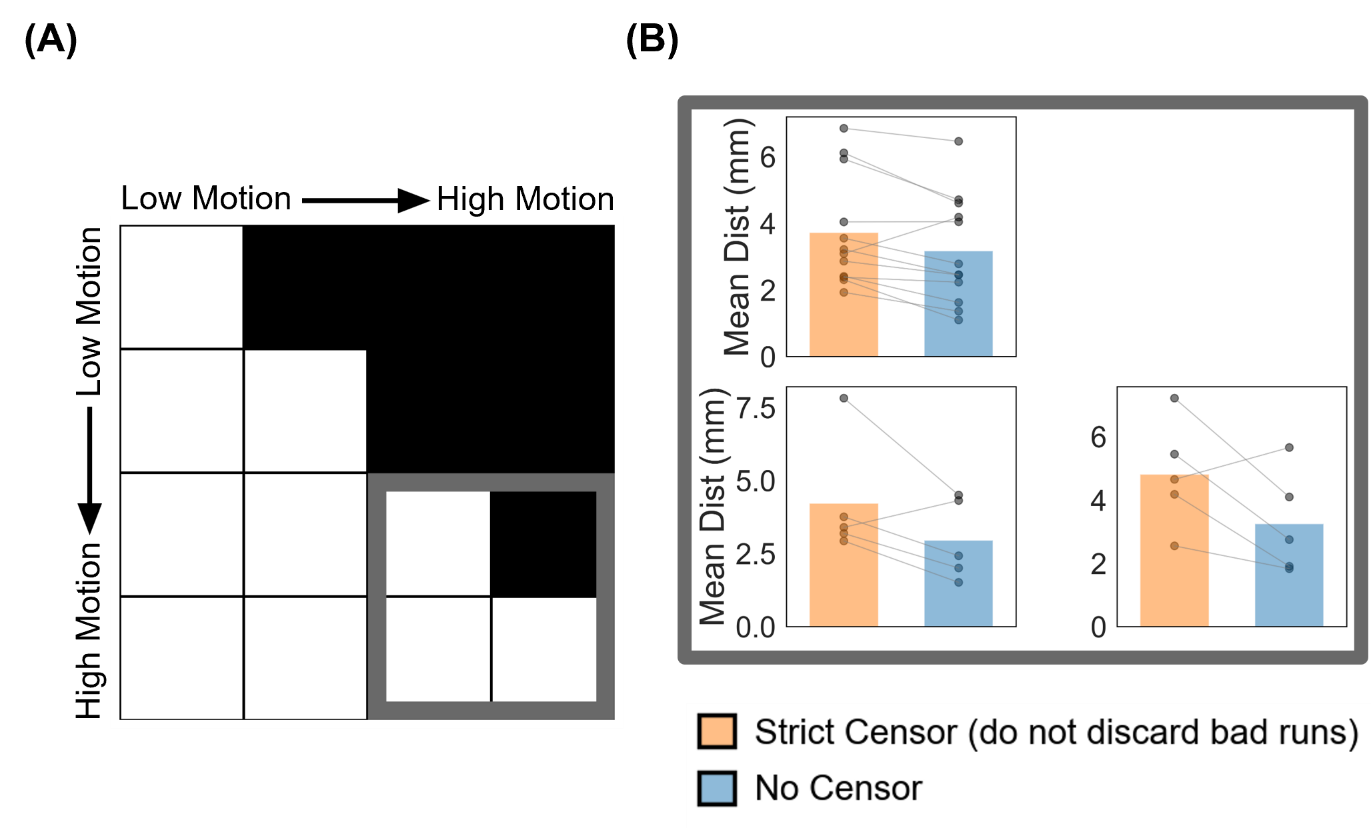

##### **Supplementary Figure 50.** Comparison of TMS depression targeting quality (Euclidean distance from ground-truth targets) between strict censoring and no censoring for the high-motion bins in the two-run FD setup, using simultaneous regression of bandpass filters together with nuisance regressors. Here strict censoring refers to FD threshold of 0.08 mm (after pseudo-motion filtering) and DVARS threshold of 50. High-motion runs were not discarded for strict censoring, so a comparison could be made with no censoring. Smaller Euclidean distance indicates better quality. TMS depression targets were estimated using the tree-algorithm (Kong et al., 2026a). (A) Schematic highlighting the three high-motion bins with thick gray outlines. (B) TMS targeting quality (Euclidean distance) for the two censoring approaches across the 3 high-motion bins. Each cell in the grid indicates a motion bin. Each dot represents the mean Euclidean distance across 50 simulated sessions for a participant in that motion bin. P-values were computed using the two-sided paired-sampled t-test. "*" indicates statistical significance after multiple comparisons correction with FDR q < 0.05.

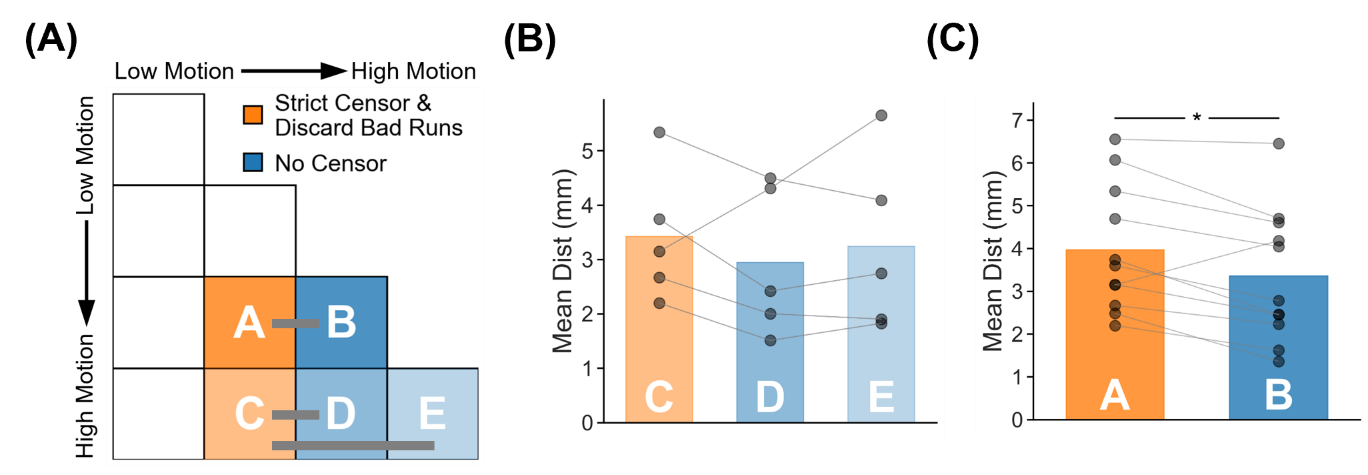

##### **Supplementary Figure 51.** TMS depression target quality (Euclidean distance to ground-truth targets) for two high-motion runs with no censoring vs mixed-motion runs with strict censoring, using simultaneous regression of bandpass filters together with nuisance regressors. Here mixed-motion runs refer to sessions with one high-motion and one low-motion run, so the high-motion run will be discarded under strict censoring. Strict censoring refers to FD threshold of 0.08 mm (after pseudo-motion filtering) and DVARS threshold of 50. Lower Euclidean distance indicates better quality. TMS depression targets were estimated using the tree-algorithm (Kong et al., 2026a). (A) Schematic illustrating the comparison. Motion bins A and B were compared, while motion bins C, D and E were compared. (B) Comparison of motion bins C vs D, and C vs E. (C) Comparison of motion bins A vs B. For both panels (B) and (C), each dot represents the mean Euclidean distance across 50 simulated sessions for a participant in that motion bin. P-values were computed using the two-sided paired-sampled t-test. Each statistical test was performed using only participants common to both motion bins. "*" indicates statistical significance after multiple comparisons correction with FDR q < 0.05.

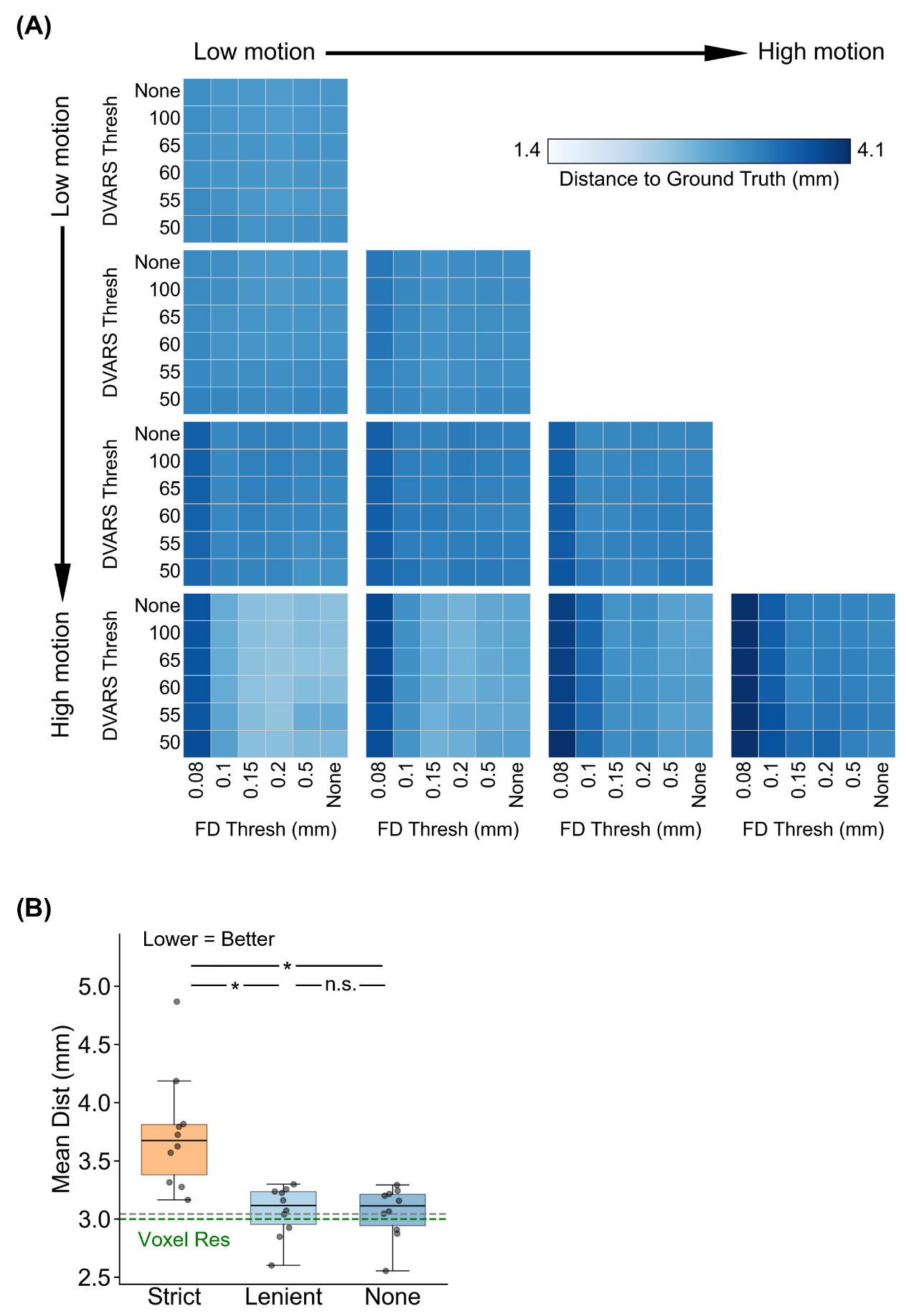

##### **Supplementary Figure 52.** Comparison of TMS depression target quality (Euclidean distance from ground-truth targets) across varying FD and DVARS thresholds for the two-run FD setup, using simultaneous regression of bandpass filters together with nuisance regressors. For this analysis, high-motion runs were not discarded. Lower Euclidean distance indicates better quality. TMS depression targets were estimated using the tree-algorithm (Kong et al., 2026a). (A) TMS depression target quality (Euclidean distance) for different combinations of FD and DVARS thresholds for 10 motion bins. Each cell in the grid indicates a motion bin. For each motion bin, there are 36 small cells corresponding to different FD and DVARS thresholds. (B) TMS depression target quality for strict censoring, lenient censoring and no censoring. Here strict censoring refers to FD threshold of 0.08 mm (after pseudo-motion filtering) and DVARS threshold of 50. Lenient censoring refers to FD threshold of 0.5 mm and DVARS threshold of 100. Each dot represents the mean Euclidean distance for a motion bin. The gray dashed line is the “noise ceiling”, achieved by picking the best combination of FD and DVARS threshold for each motion bin, and computing the median across motion bins. P-values were computed using a linear mixed effects model (p = 1.1e-12 for strict vs. none, p = 2.2e-12 for strict vs. lenient, p = 0.93 for lenient vs. none). "*" indicates statistical significance after multiple comparisons correction with FDR q < 0.05 and "n.s." indicates not significant after FDR correction. For each box plot, the horizontal line indicates the median across all motion bins. The bottom and top edges of the box indicate the 25th and 75th percentiles, respectively. The outliers are defined as data points beyond 1.5 times the interquartile range. The whiskers extend to the most extreme data points not considered outliers.

#### **Logit-transformed Dice (two runs FD) for parcellations**

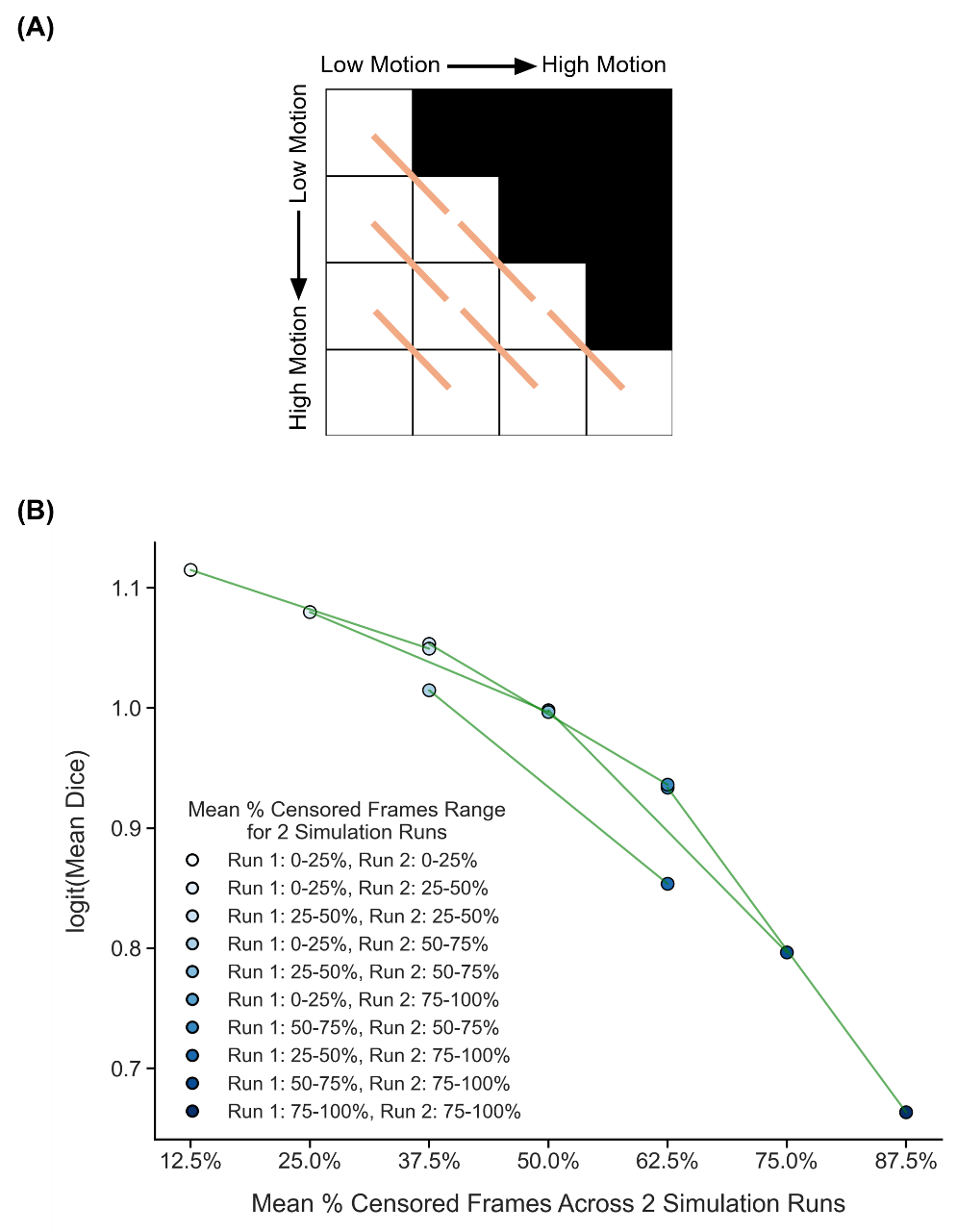

##### **Supplementary Figure 53.** Comparison of parcellation quality (logit-transformed Dice) across diagonally adjacent motion bins under strict censoring, for the two-run FD setup. Here strict censoring refers to FD threshold of 0.08 mm (after pseudo-motion filtering) and DVARS threshold of 50. High-motion runs were not discarded to allow comparison with low-motion runs. Higher logit-transformed Dice indicates better quality. (A) Schematic of comparisons across diagonally adjacent motion bins. Each orange line indicates a single comparison. (B) Relationship between parcellation quality (logit-transformed Dice) and motion. Each circle indicates the mean logit-transformed Dice coefficient across simulation runs of all participants in a motion bin. Each line indicates a single comparison between two adjacent motion bins. P-values were computed using the two-sided paired-sampled t-test using only participants common to both motion bins. An overall statistical test combining all comparisons was conducted using a linear mixed effects model (p = 3.2e-32).

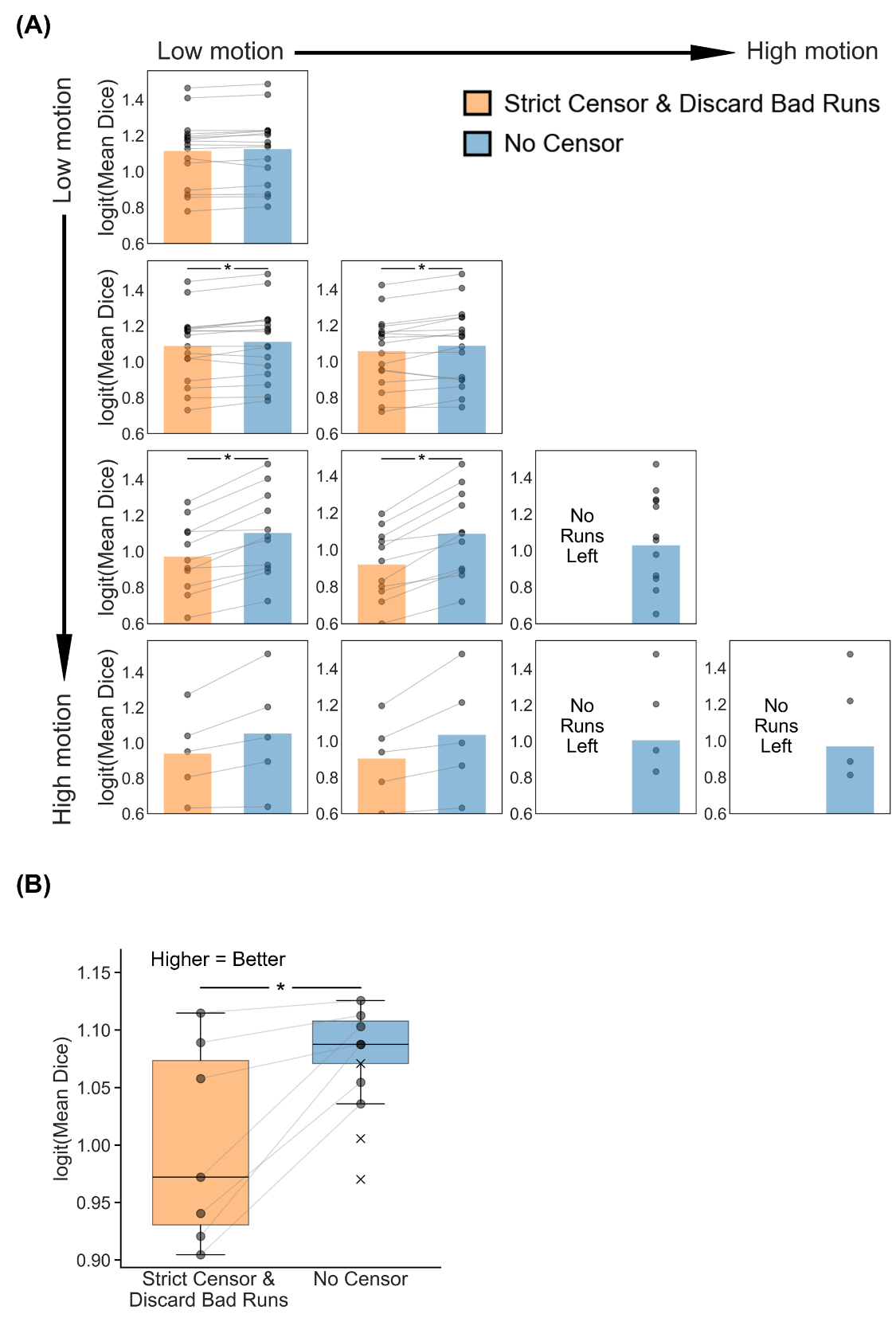

##### **Supplementary Figure 54.** Comparison of parcellation quality (logit-transformed Dice) between strict censoring and no censoring, for the two-run FD setup. Here, strict censoring refers to a FD threshold of 0.08 mm (after pseudo-motion filtering) and a DVARS threshold of 50. High-motion runs were discarded. Higher logit-transformed Dice indicates better quality. (A) Parcellation quality for the two censoring approaches across 10 motion bins. Each cell in the grid indicates a motion bin. Each dot represents the mean logit-transformed Dice coefficient across 50 simulated sessions of a participant in that motion bin. P-values were computed using the two-sided paired-sampled t-test. "*" indicates statistical significance after multiple comparisons correction with FDR q < 0.05. Under strict censoring, no runs remained for the bottom-right three motion bins because all fMRI runs were discarded under this criterion. (B) Parcellation quality across all motion bins for both censoring strategies. There are seven pairs of dots, corresponding to the seven motion bins where comparisons could be made between strict censoring and no censoring. P-value was computed using a linear mixed effects model (p = 4.8e-6). "*" indicates statistical significance after multiple comparisons correction with FDR q < 0.05. There are three “×” corresponding to the logit-transformed Dice coefficient of the three bottom-right bins under the no censoring condition. The × were not included in the boxplots or statistical comparisons between the two censoring strategies. For each box plot, the horizontal line indicates the median. The bottom and top edges of the box indicate the 25th and 75th percentiles, respectively. The outliers are defined as data points beyond 1.5 times the interquartile range. The whiskers extend to the most extreme data points not considered outliers.

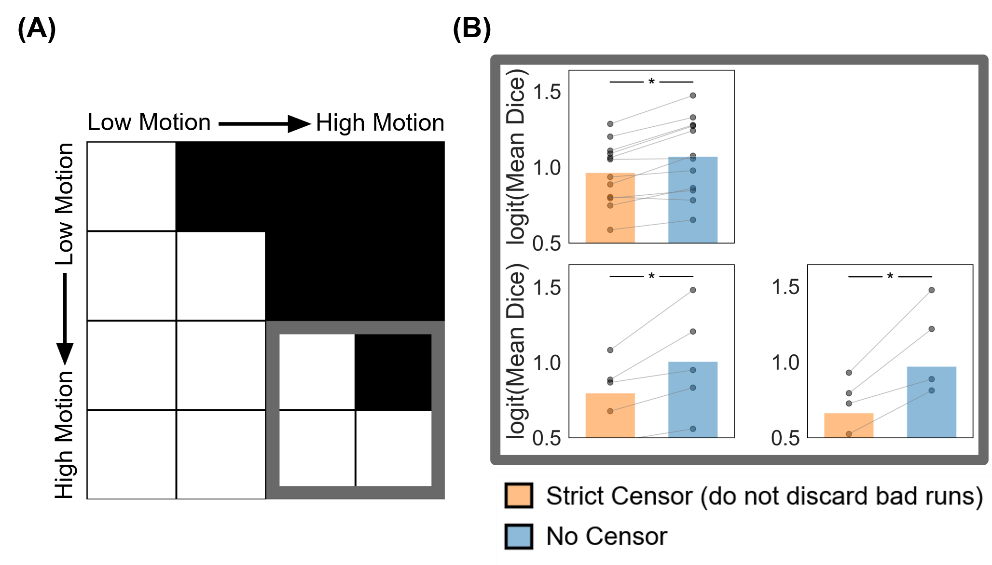

##### **Supplementary Figure 55.** Comparison of parcellation quality (logit-transformed Dice) between strict censoring and no censoring for the high-motion bins in the two-run FD setup. Here strict censoring refers to FD threshold of 0.08 mm (after pseudo-motion filtering) and DVARS threshold of 50. High-motion runs were not discarded for strict censoring, so a comparison could be made with no censoring (A) Schematic highlighting the three high-motion bins with thick gray outlines. (B) Parcellation quality (logit-transformed Dice) for the two censoring approaches across the 3 high-motion bins. Each cell in the grid indicates a motion bin. Each dot represents the mean logit-transformed Dice coefficient across 50 simulated sessions for a participant in that motion bin. P-values were computed using the two-sided paired-sampled t-test. "*" indicates statistical significance after multiple comparisons correction with FDR q < 0.05.

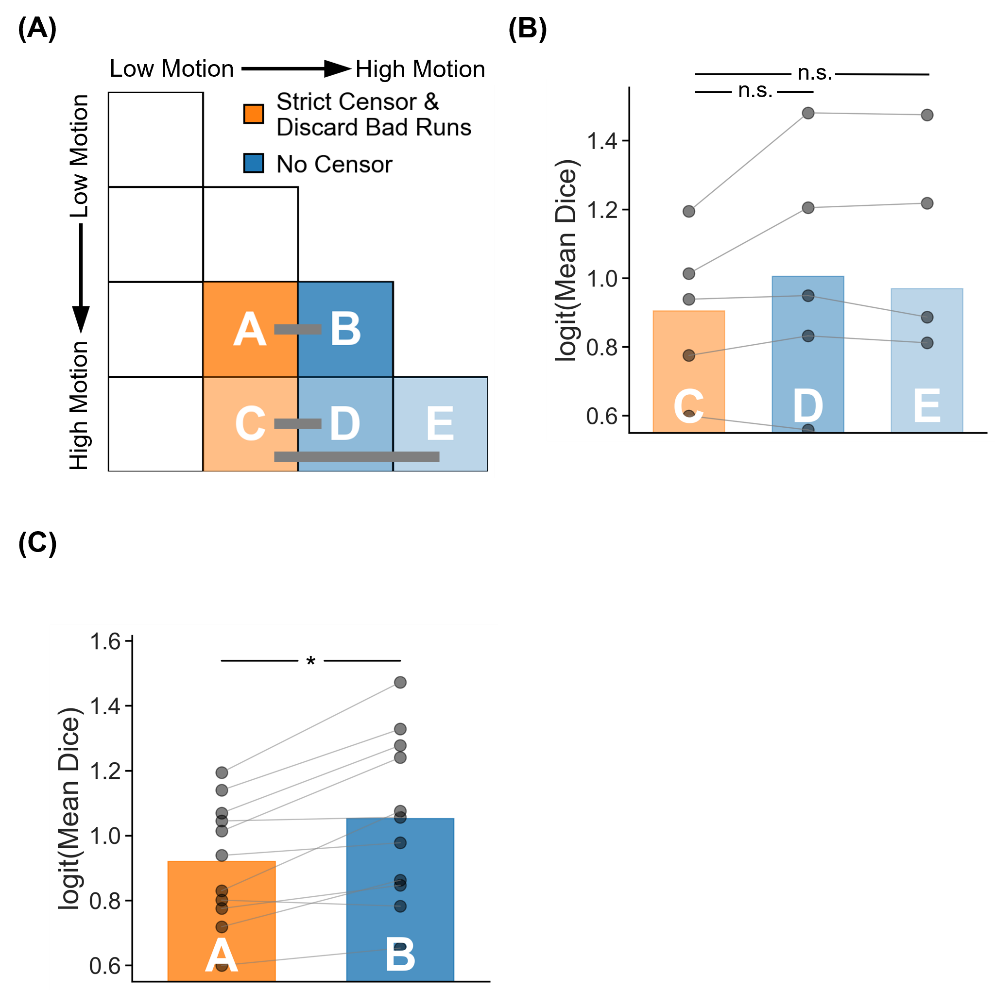

##### **Supplementary Figure 56.** Parcellation quality (logit-transformed Dice) for two high-motion runs with no censoring vs mixed-motion runs with strict censoring. Here mixed-motion runs refer to sessions with one high-motion and one low-motion run, so the high-motion run will be discarded under strict censoring. Strict censoring refers to FD threshold of 0.08 mm (after pseudo-motion filtering) and DVARS threshold of 50. Higher logit-transformed Dice indicates better quality. (A) Schematic illustrating the comparison. Motion bins A and B were compared, while motion bins C, D and E were compared. (B) Comparison of motion bins C vs D, and C vs E. (C) Comparison of motion bins A vs B. For both panels (B) and (C), each dot represents the mean logit-transformed Dice coefficient across 50 simulated sessions for a participant in that motion bin. P-values were computed using the two-sided paired-sampled t-test. Each statistical test was performed using only participants common to both motion bins. "*" indicates statistical significance after multiple comparisons correction with FDR q < 0.05. (D) Discrepancy with respect to the ground truth parcellation in a representative participant from motion bins A (with strict censoring) and B (with no censoring). Blue regions indicate agreement with ground-truth parcellation, while red regions indicate differences. (E) Parcellations of the representative participant from panel D: ground-truth parcellation (left), simulated parcellation under strict censoring for motion bin A (middle), and simulated parcellation under no censoring for motion bin B (right). Parcellations from two high-motion runs without censoring were equivalent (panel B) or superior (panel C) to those from strictly censored mixed-motion runs.

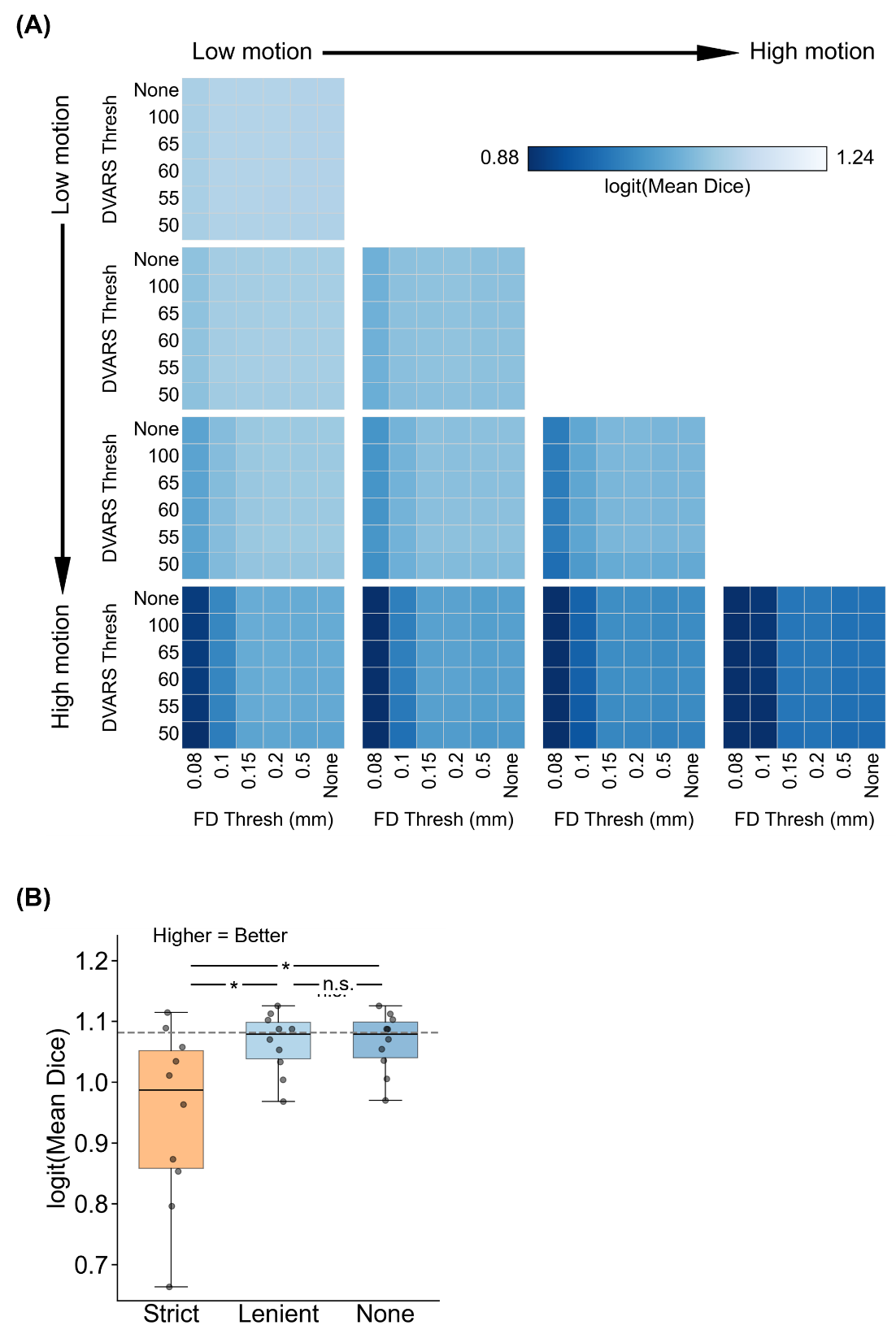

##### **Supplementary Figure 57.** Comparison of parcellation quality (logit-transformed Dice) across varying FD and DVARS thresholds for the two-run FD setup. For this analysis, high-motion runs were not discarded. Higher logit-transformed Dice indicates better quality. (A) Parcellation quality for different combinations of FD and DVARS thresholds for 10 motion bins. Each cell in the grid indicates a motion bin. For each motion bin, there are 36 small cells corresponding to different FD and DVARS thresholds. (B) Parcellation quality of strict censoring, lenient censoring and no censoring. Here strict censoring refers to FD threshold of 0.08 mm (after pseudo-motion filtering) and DVARS threshold of 50. Lenient censoring refers to FD threshold of 0.5 mm and DVARS threshold of 100. Each dot represents the logit-transformed Dice coefficient for a motion bin. The gray dashed line is the “noise ceiling”, achieved by picking the best combination of FD and DVARS threshold for each motion bin, and computing the median across motion bins. P-values were computed using a linear mixed effects model (p = 1.2e-22 for strict vs. none, p = 1.8e-22 for strict vs. lenient, p = 0.97 for lenient vs. none). "*" indicates statistical significance after multiple comparisons correction with FDR q < 0.05 and "n.s." indicates not significant after FDR correction. For each box plot, the horizontal line indicates the median across all motion bins. The bottom and top edges of the box indicate the 25th and 75th percentiles, respectively. The outliers are defined as data points beyond 1.5 times the interquartile range. The whiskers extend to the most extreme data points not considered outliers.

#### **Logarithm-transformed Euclidean distance (two runs FD) for tree-based (Kong2026) TMS depression targets**

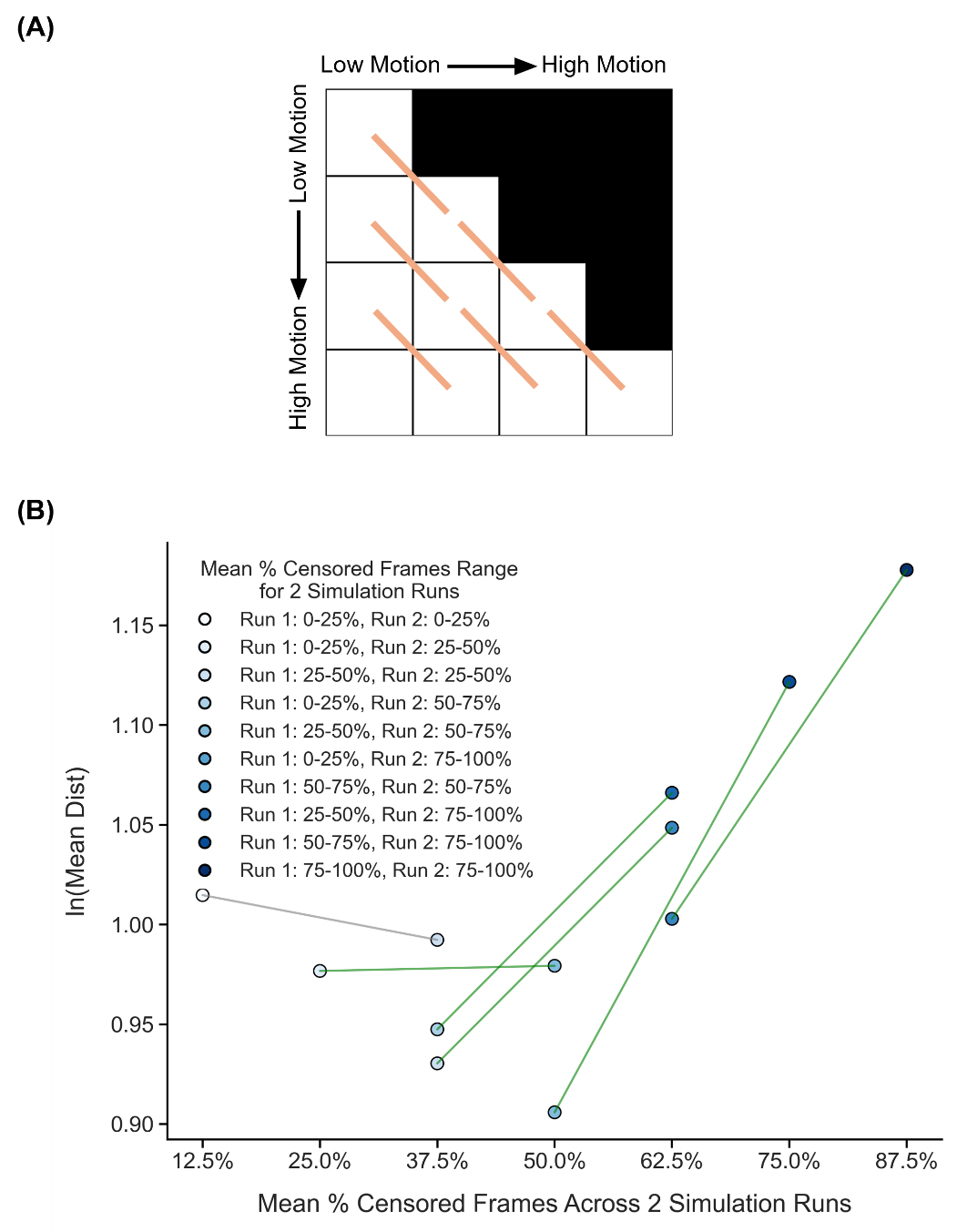

##### **Supplementary Figure 58.** Comparison of TMS depression target quality (logarithm-transformed Euclidean distance from ground-truth targets) across diagonally adjacent motion bins under strict censoring, for the two-run FD setup. Here strict censoring refers to FD threshold of 0.08 mm (after pseudo-motion filtering) and DVARS threshold of 50. High-motion runs were not discarded to allow comparison with low-motion runs. Smaller logarithm-transformed Euclidean distance to ground-truth target indicates better quality. TMS depression targets were estimated using the tree-algorithm (Kong et al., 2026a). (A) Schematic of comparisons across diagonally adjacent motion bins. Each orange line indicates a single comparison. (B) Relationship between TMS depression target quality (logarithm-transformed Euclidean distance) and motion. Each circle indicates the mean logarithm-transformed Euclidean distance across simulation runs of all participants in a motion bin. Each line indicates a single comparison between two adjacent motion bins. P-values were computed using the two-sided paired-sampled t-test using only participants common to both motion bins. An overall statistical test combining all comparisons was conducted using a linear mixed effects model (p = 2.5e-5).

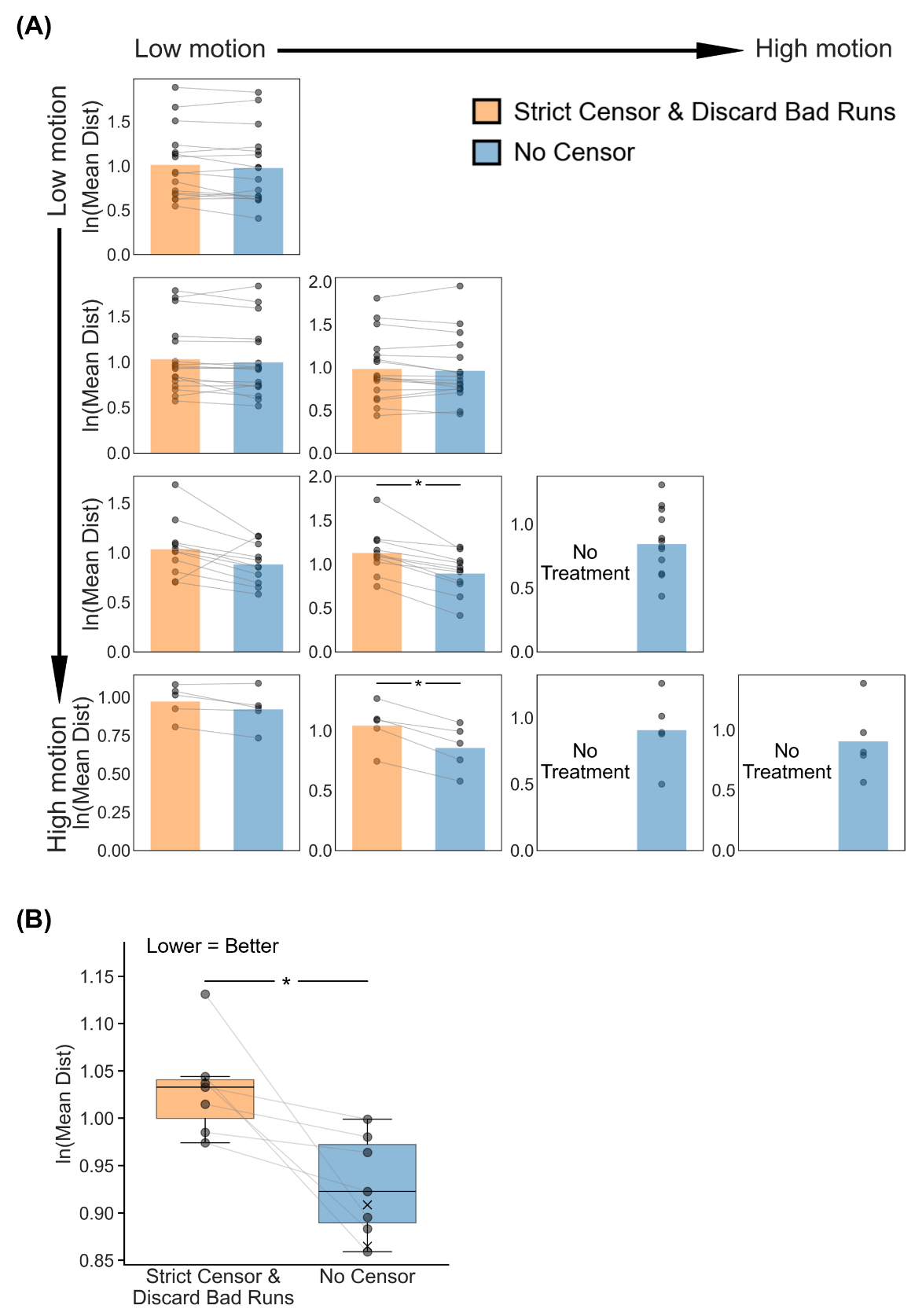

##### **Supplementary Figure 59.** Comparison of TMS depression targeting quality (logarithm-transformed Euclidean distance to ground-truth depression targets) between strict censoring and no censoring, for the two-run FD setup. Here strict censoring refers to FD threshold of 0.08 mm (after pseudo-motion filtering) and DVARS threshold of 50. High-motion runs were discarded. Smaller logarithm-transformed Euclidean distance indicates better quality. TMS depression targets were estimated using the tree-based algorithm (Kong et al., 2026a). (A) TMS depression targeting quality (logarithm-transformed Euclidean distance) for the two censoring approaches across 10 motion bins. Each cell in the grid indicates a motion bin. Each dot represents the mean logarithm-transformed Euclidean distance across 50 simulated sessions of a participant in that motion bin. P-values were computed using the two-sided paired-sampled t-test. "*" indicates statistical significance after multiple comparisons correction with FDR q < 0.05. Under strict censoring, no runs remained for the bottom-right three motion bins because all fMRI runs were discarded under this criterion. (B) TMS depression targeting quality across all motion bins for both censoring strategies. There are seven pairs of dots, corresponding to the seven motion bins where comparisons could be made between strict censoring and no censoring. P-value was computed using a linear mixed effects model (p = 8.1e-3). "*" indicates statistical significance after multiple comparisons correction with FDR q < 0.05. There are three “×” corresponding to the mean logarithm-transformed Euclidean distance of the 3 bottom-right bins under the no censoring condition. The × were not included in the boxplots or statistical comparisons between the two censoring strategies. For each box plot, the horizontal line indicates the median. The bottom and top edges of the box indicate the 25th and 75th percentiles, respectively. The outliers are defined as data points beyond 1.5 times the interquartile range. The whiskers extend to the most extreme data points not considered outliers.

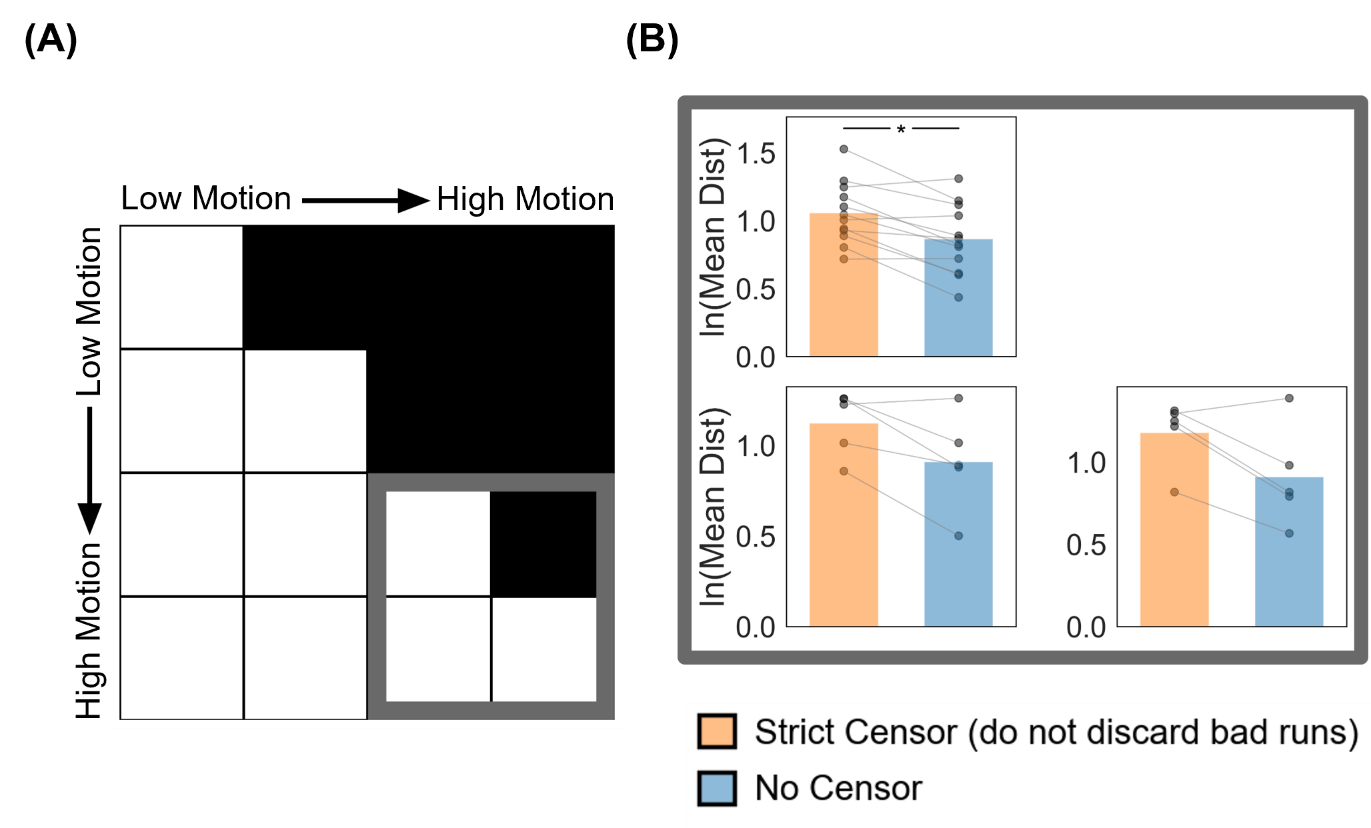

##### **Supplementary Figure 60.** Comparison of TMS depression targeting quality (logarithm-transformed Euclidean distance from ground-truth targets) between strict censoring and no censoring for the high-motion bins in the two-run FD setup. Here strict censoring refers to FD threshold of 0.08 mm (after pseudo-motion filtering) and DVARS threshold of 50. High-motion runs were not discarded for strict censoring, so a comparison could be made with no censoring. Smaller logarithm-transformed Euclidean distance indicates better quality. TMS depression targets were estimated using the tree-algorithm (Kong et al., 2026a). (A) Schematic highlighting the three high-motion bins with thick gray outlines. (B) TMS targeting quality (logarithm-transformed Euclidean distance) for the two censoring approaches across the 3 high-motion bins. Each cell in the grid indicates a motion bin. Each dot represents the mean logarithm-transformed Euclidean distance across 50 simulated sessions for a participant in that motion bin. P-values were computed using the two-sided paired-sampled t-test. "*" indicates statistical significance after multiple comparisons correction with FDR q < 0.05.

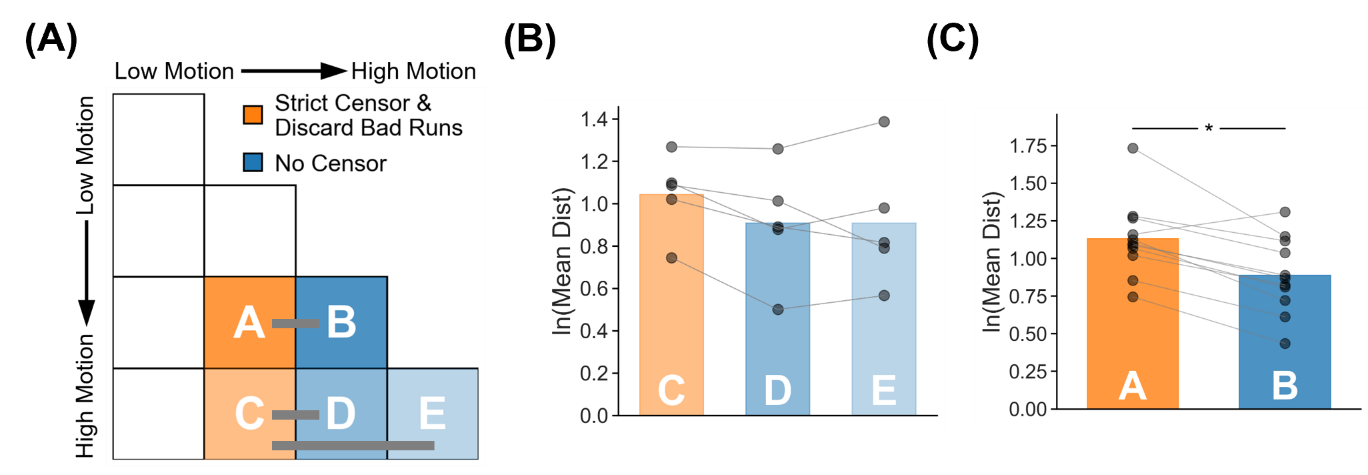

##### **Supplementary Figure 61.** TMS depression target quality (logarithm-transformed Euclidean distance to ground-truth targets) for two high-motion runs with no censoring vs mixed-motion runs with strict censoring. Here mixed-motion runs refer to sessions with one high-motion and one low-motion run, so the high-motion run will be discarded under strict censoring. Strict censoring refers to FD threshold of 0.08 mm (after pseudo-motion filtering) and DVARS threshold of 50. Lower logarithm-transformed Euclidean distance indicates better quality. TMS depression targets were estimated using the tree-algorithm (Kong et al., 2026a). (A) Schematic illustrating the comparison. Motion bins A and B were compared, while motion bins C, D and E were compared. (B) Comparison of motion bins C vs D, and C vs E. (C) Comparison of motion bins A vs B. For both panels (B) and (C), each dot represents the mean logarithm-transformed Euclidean distance across 50 simulated sessions for a participant in that motion bin. P-values were computed using the two-sided paired-sampled t-test. Each statistical test was performed using only participants common to both motion bins. "*" indicates statistical significance after multiple comparisons correction with FDR q < 0.05.

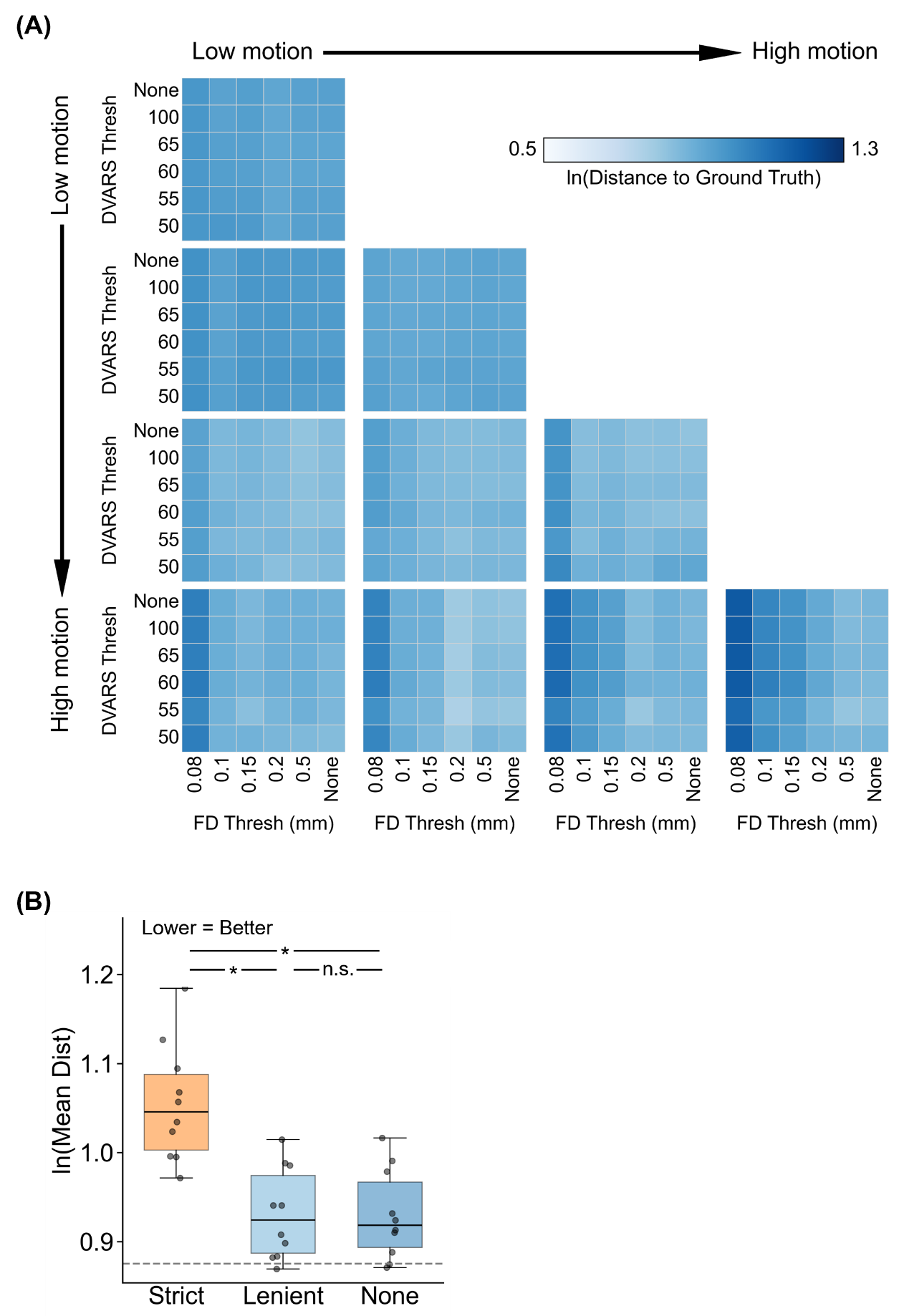

##### **Supplementary Figure 62.** Comparison of TMS depression target quality (logarithm-transformed Euclidean distance from ground-truth targets) across varying FD and DVARS thresholds for the two-run FD setup. For this analysis, high-motion runs were not discarded. Lower logarithm-transformed Euclidean distance indicates better quality. TMS depression targets were estimated using the tree-algorithm (Kong et al., 2026a). (A) TMS depression target quality (logarithm-transformed Euclidean distance) for different combinations of FD and DVARS thresholds for 10 motion bins. Each cell in the grid indicates a motion bin. For each motion bin, there are 36 small cells corresponding to different FD and DVARS thresholds. (B) TMS depression target quality for strict censoring, lenient censoring and no censoring. Here strict censoring refers to FD threshold of 0.08 mm (after pseudo-motion filtering) and DVARS threshold of 50. Lenient censoring refers to FD threshold of 0.5 mm and DVARS threshold of 100. Each dot represents the mean logarithm-transformed Euclidean distance for a motion bin. The gray dashed line is the “noise ceiling”, achieved by picking the best combination of FD and DVARS threshold for each motion bin, and computing the median across motion bins. P-values were computed using a linear mixed effects model (p = 2.6e-17 for strict vs. none, p = 1.9e-17 for strict vs. lenient, p = 0.97 for lenient vs. none). "*" indicates statistical significance after multiple comparisons correction with FDR q < 0.05 and "n.s." indicates not significant after FDR correction. For each box plot, the horizontal line indicates the median across all motion bins. The bottom and top edges of the box indicate the 25th and 75th percentiles, respectively. The outliers are defined as data points beyond 1.5 times the interquartile range. The whiskers extend to the most extreme data points not considered outliers.

#### **Hausdorff distance (two runs FD) for parcellations**

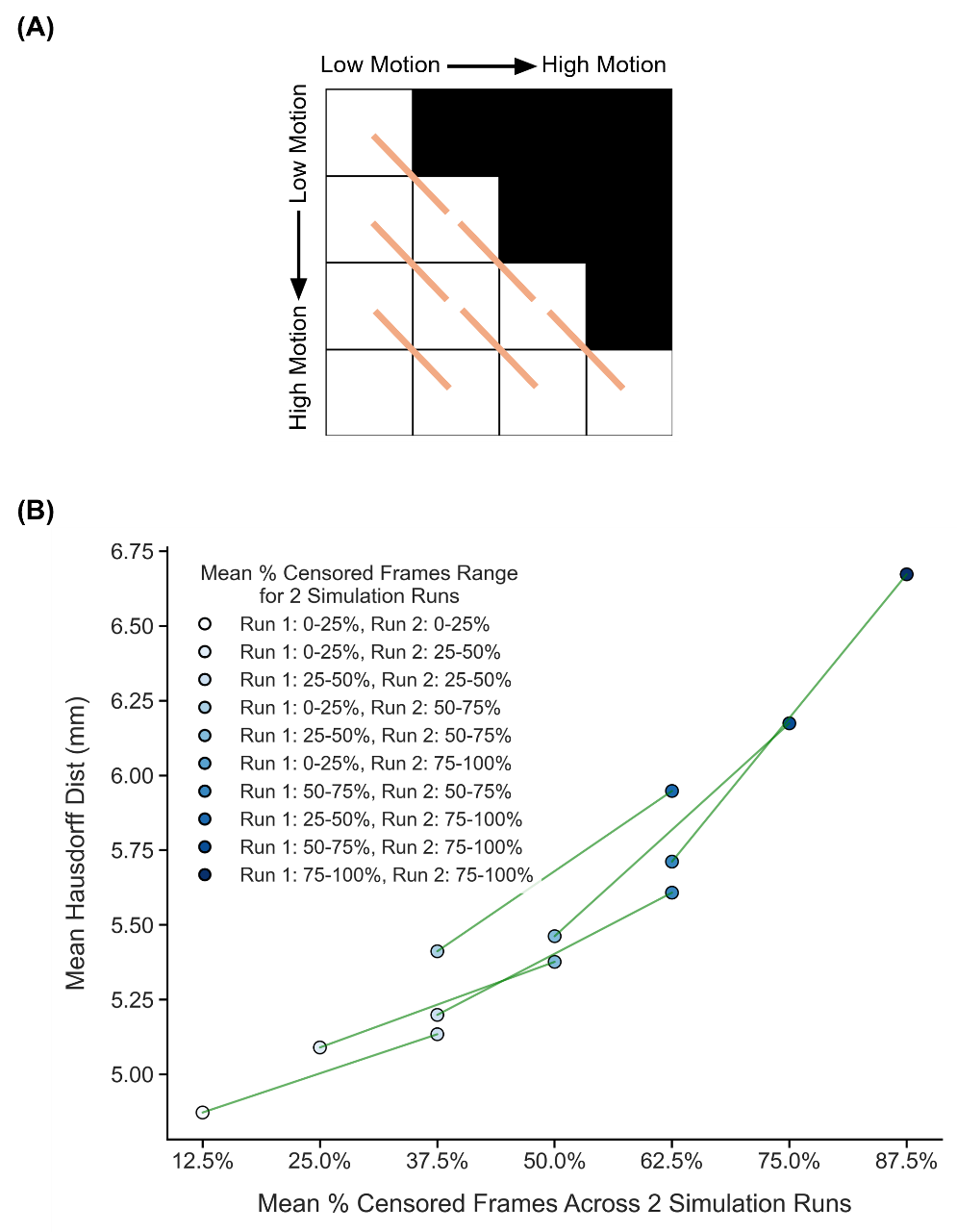

##### **Supplementary Figure 63.** Comparison of parcellation quality (mean Hausdorff distance) across diagonally adjacent motion bins under strict censoring, for the two-run FD setup. Here strict censoring refers to FD threshold of 0.08 mm (after pseudo-motion filtering) and DVARS threshold of 50. High-motion runs were not discarded to allow comparison with low-motion runs. Lower Hausdorff distance indicates better quality. (A) Schematic of comparisons across diagonally adjacent motion bins. Each orange line indicates a single comparison. (B) Relationship between parcellation quality (mean Hausdorff distance) and motion. Each circle indicates the mean Hausdorff distance across simulation runs of all participants in a motion bin. Each line indicates a single comparison between two adjacent motion bins. P-values were computed using the two-sided paired-sampled t-test using only participants common to both motion bins. An overall statistical test combining all comparisons was conducted using a linear mixed effects model (p = 3.7e-27).

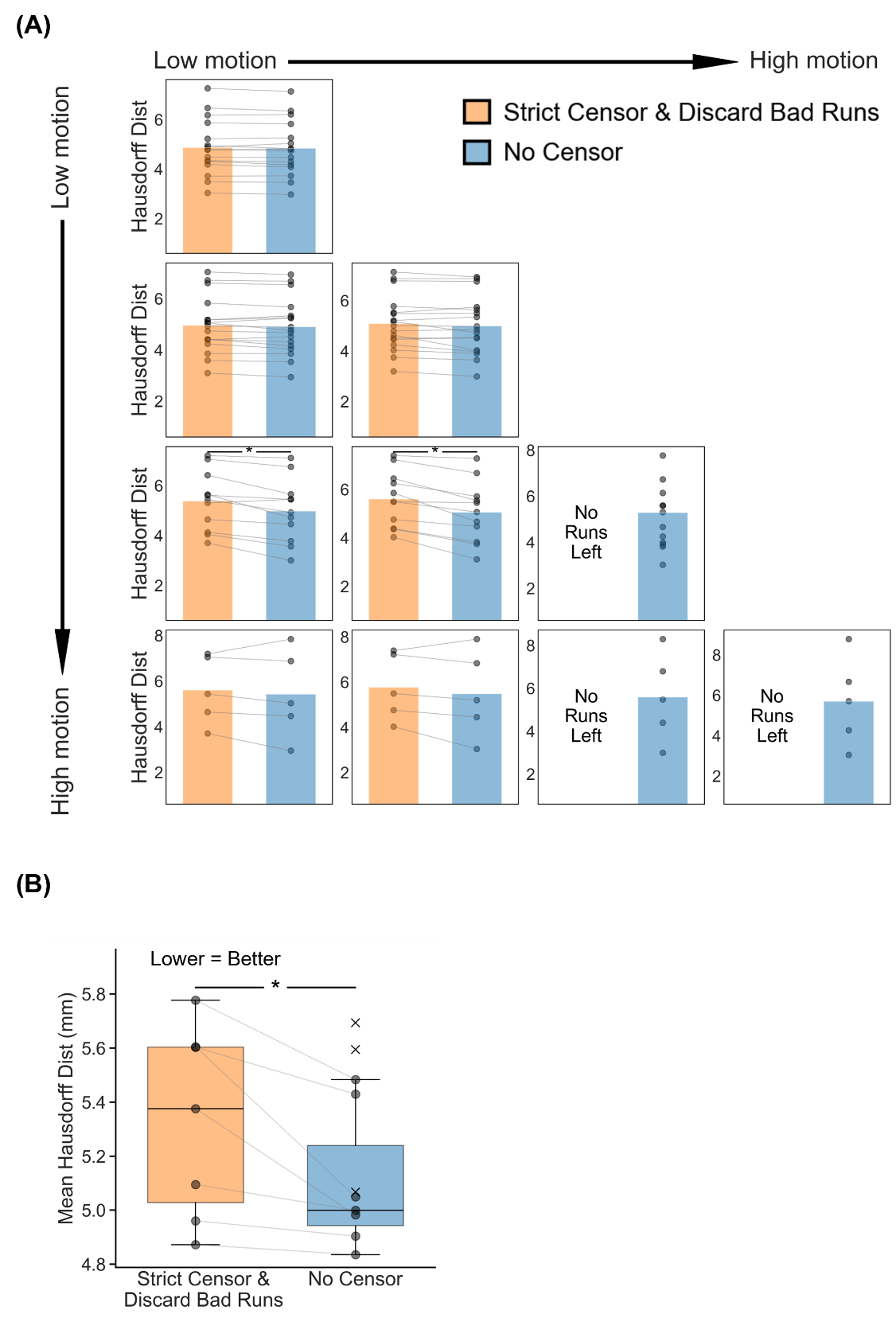

##### **Supplementary Figure 64.** Comparison of parcellation quality (mean Hausdorff distance) between strict censoring and no censoring, for the two-run FD setup. Here, strict censoring refers to a FD threshold of 0.08 mm (after pseudo-motion filtering) and a DVARS threshold of 50. High-motion runs were discarded. Lower mean Hausdorff distance indicates better quality. (A) Parcellation quality for the two censoring approaches across 10 motion bins. Each cell in the grid indicates a motion bin. Each dot represents the mean Hausdorff distance across 50 simulated sessions of a participant in that motion bin. P-values were computed using the two-sided paired-sampled t-test. "*" indicates statistical significance after multiple comparisons correction with FDR q < 0.05. Under strict censoring, no runs remained for the bottom-right three motion bins because all fMRI runs were discarded under this criterion. (B) Parcellation quality across all motion bins for both censoring strategies. There are seven pairs of dots, corresponding to the seven motion bins where comparisons could be made between strict censoring and no censoring. P-value was computed using a linear mixed effects model (p = 1.1e-3). "*" indicates statistical significance after multiple comparisons correction with FDR q < 0.05. There are three “×” corresponding to the Hausdorff distance of the three bottom-right bins under the no censoring condition. The × were not included in the boxplots or statistical comparisons between the two censoring strategies. For each box plot, the horizontal line indicates the median. The bottom and top edges of the box indicate the 25th and 75th percentiles, respectively. The outliers are defined as data points beyond 1.5 times the interquartile range. The whiskers extend to the most extreme data points not considered outliers.

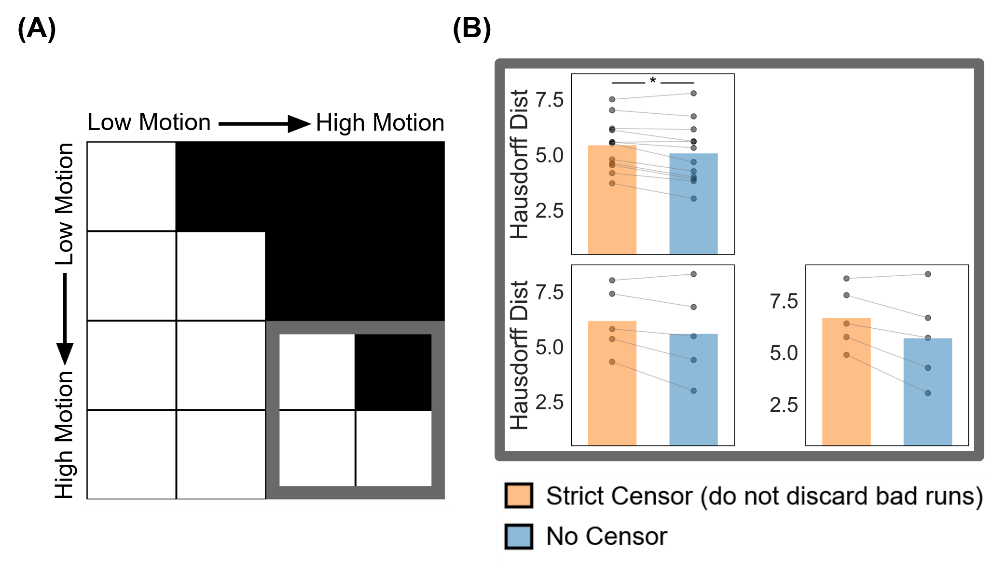

##### **Supplementary Figure 65.** Comparison of parcellation quality (Hausdorff distance) between strict censoring and no censoring for the high-motion bins in the two-run FD setup. Here strict censoring refers to FD threshold of 0.08 mm (after pseudo-motion filtering) and DVARS threshold of 50. High-motion runs were not discarded for strict censoring, so a comparison could be made with no censoring (A) Schematic highlighting the three high-motion bins with thick gray outlines. (B) Parcellation quality (Hausdorff distance) for the two censoring approaches across the 3 high-motion bins. Each cell in the grid indicates a motion bin. Each dot represents the mean Hausdorff distance across 50 simulated sessions for a participant in that motion bin. P-values were computed using the two-sided paired-sampled t-test. "*" indicates statistical significance after multiple comparisons correction with FDR q < 0.05.

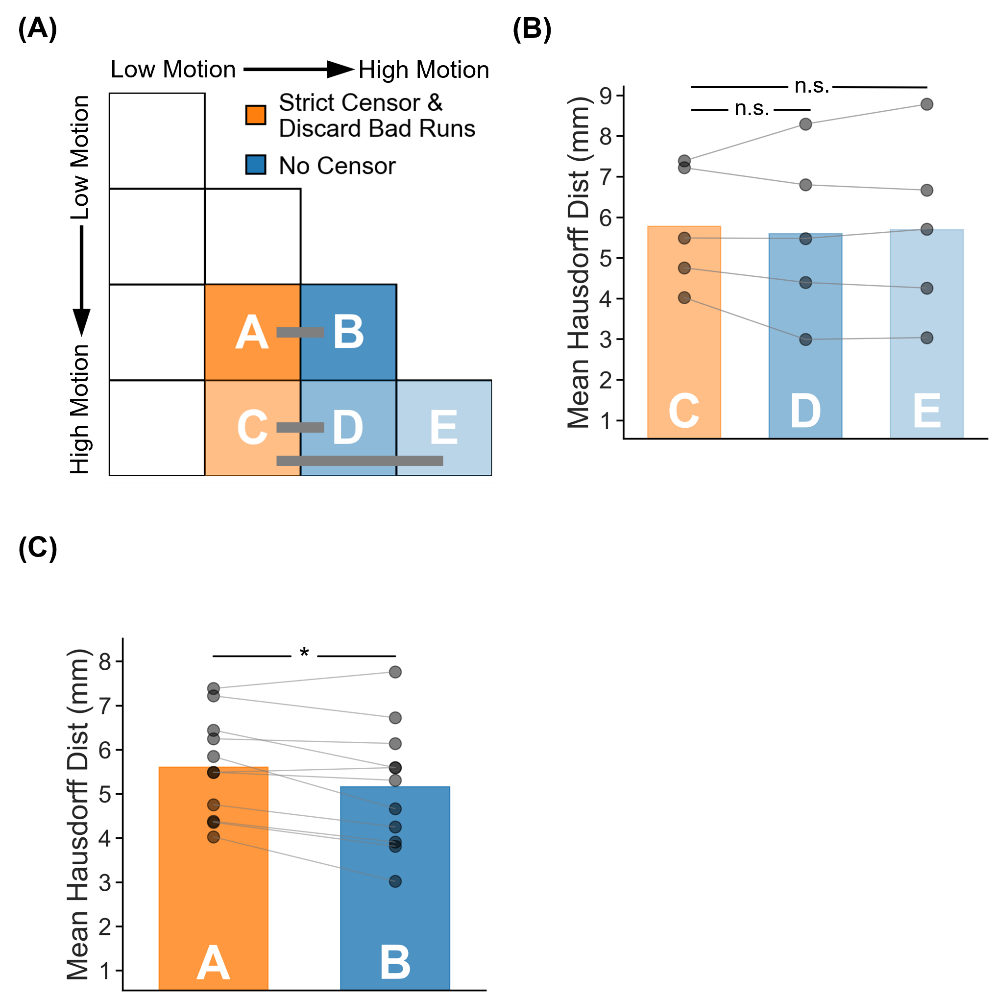

##### **Supplementary Figure 66.** Parcellation quality (Hausdorff distance) for two high-motion runs with no censoring vs mixed-motion runs with strict censoring. Here mixed-motion runs refer to sessions with one high-motion and one low-motion run, so the high-motion run will be discarded under strict censoring. Strict censoring refers to FD threshold of 0.08 mm (after pseudo-motion filtering) and DVARS threshold of 50. Lower Hausdorff distance indicates better quality. (A) Schematic illustrating the comparison. Motion bins A and B were compared, while motion bins C, D and E were compared. (B) Comparison of motion bins C vs D, and C vs E. (C) Comparison of motion bins A vs B. For both panels (B) and (C), each dot represents the mean Hausdorff distance across 50 simulated sessions for a participant in that motion bin. P-values were computed using the two-sided paired-sampled t-test. Each statistical test was performed using only participants common to both motion bins. "*" indicates statistical significance after multiple comparisons correction with FDR q < 0.05.

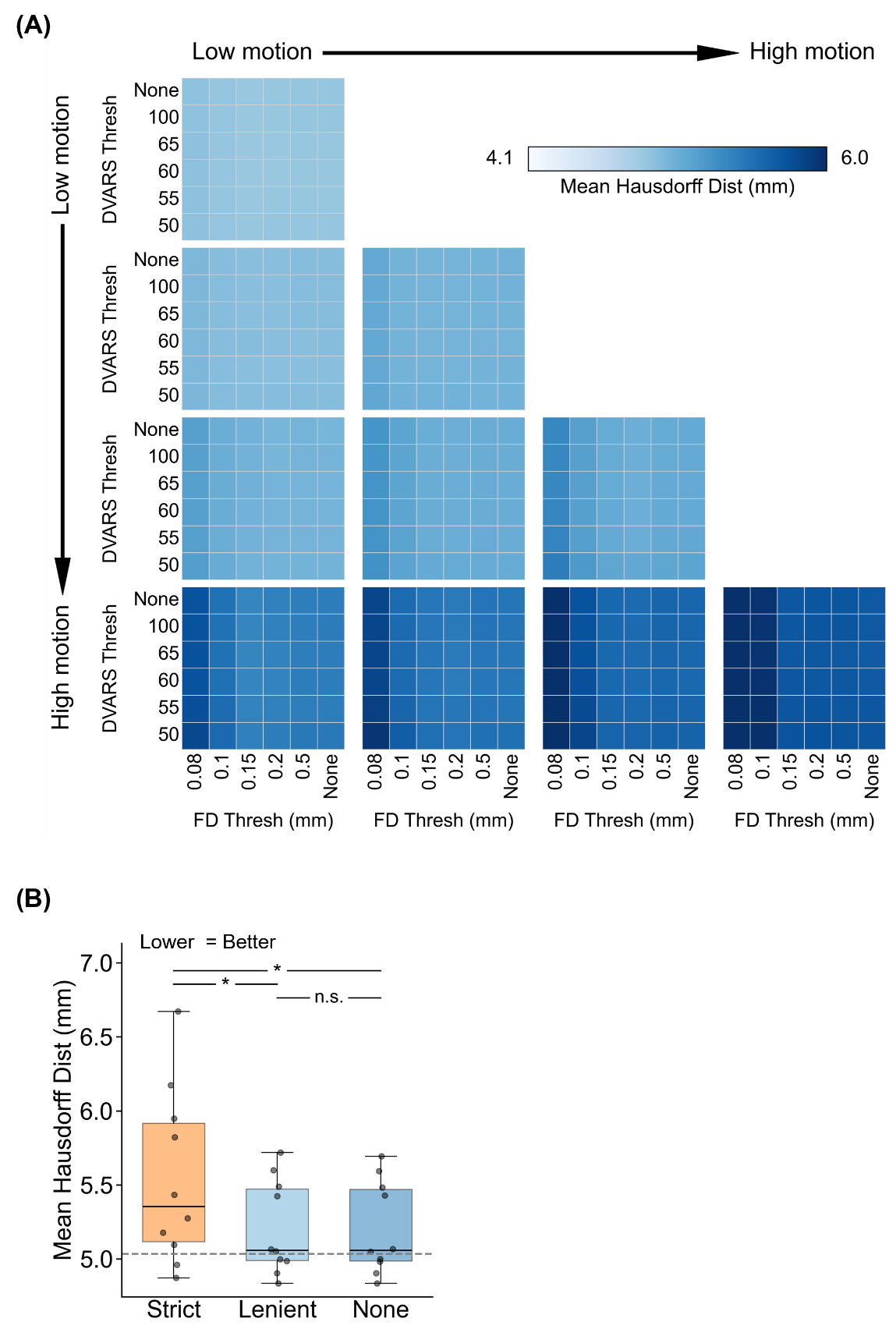

##### **Supplementary Figure 67.** Comparison of parcellation quality (Hausdorff distance) across varying FD and DVARS thresholds for the two-run FD setup. For this analysis, high-motion runs were not discarded. Lower Hausdorff distance indicates better quality. (A) Parcellation quality for different combinations of FD and DVARS thresholds for 10 motion bins. Each cell in the grid indicates a motion bin. For each motion bin, there are 36 small cells corresponding to different FD and DVARS thresholds. (B) Parcellation quality of strict censoring, lenient censoring and no censoring. Here strict censoring refers to FD threshold of 0.08 mm (after pseudo-motion filtering) and DVARS threshold of 50. Lenient censoring refers to FD threshold of 0.5 mm and DVARS threshold of 100. Each dot represents the Hausdorff distance for a motion bin. The gray dashed line is the “noise ceiling”, achieved by picking the best combination of FD and DVARS threshold for each motion bin, and computing the median across motion bins. P-values were computed using a linear mixed effects model (p = 4.3e-15 for strict vs. none, p = 6.6e-15 for strict vs. lenient, p = 0.96 for lenient vs. none). "*" indicates statistical significance after multiple comparisons correction with FDR q < 0.05 and "n.s." indicates not significant after FDR correction. For each box plot, the horizontal line indicates the median across all motion bins. The bottom and top edges of the box indicate the 25th and 75th percentiles, respectively. The outliers are defined as data points beyond 1.5 times the interquartile range. The whiskers extend to the most extreme data points not considered outliers.

#### **Continuous runs (two runs FD) for parcellations**

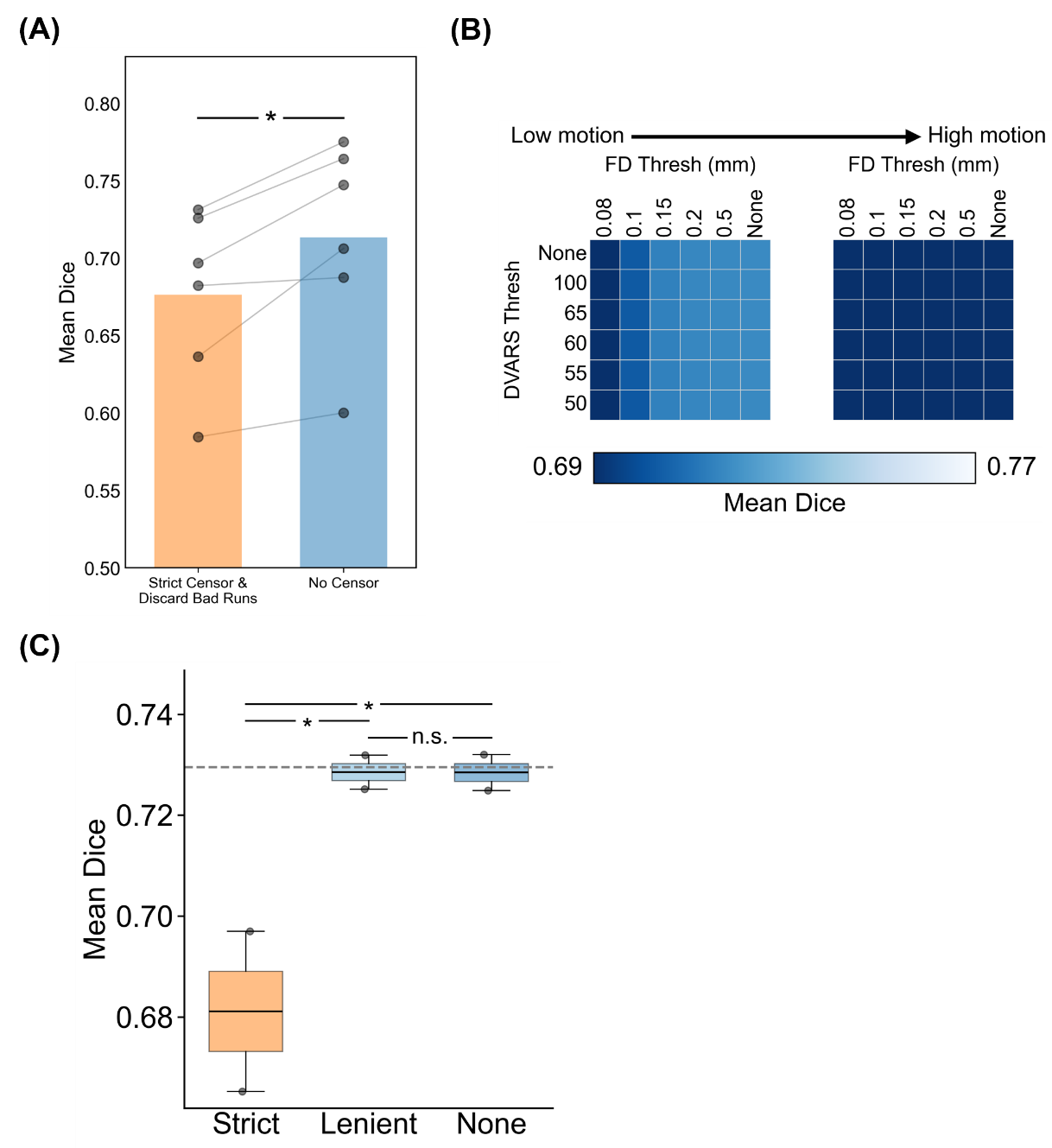

##### **Supplementary Figure 68.** Comparison of parcellation quality (Dice) using continuous real fMRI runs rather than simulated runs, for the two-run FD setup. Higher Dice indicates better quality. (A) Parcellation quality for the two censoring approaches with mixed-motion runs. Here mixed-motion runs refer to sessions with one high-motion and one low-motion run, so the high-motion run will be discarded under strict censoring. Each dot represents the mean Dice coefficient across sessions of a participant in that motion bin. P-value was computed using the two-sided paired-sampled t-test. "*" indicates statistical significance. (B, C) Comparison of parcellation quality (Dice) across varying FD and DVARS thresholds for the two-run FD setup. For this analysis, high-motion runs were not discarded. (B) Parcellation quality for different combinations of FD and DVARS thresholds for two motion bins. One motion bin has mixed-motion runs (sessions with one high-motion run and one low-motion run), and the other has two high-motion runs. Each cell in the grid indicates a motion bin. For each motion bin, there are 36 small cells corresponding to different FD and DVARS thresholds. (C) Parcellation quality of strict censoring, lenient censoring and no censoring. Here strict censoring refers to FD threshold of 0.08 mm (after pseudo-motion filtering) and DVARS threshold of 50. Lenient censoring refers to FD threshold of 0.5 mm and DVARS threshold of 100. Each dot represents the Dice coefficient for a motion bin. The gray dashed line is the “noise ceiling”, achieved by picking the best combination of FD and DVARS threshold for each motion bin, and computing the median across motion bins. P-values were computed using a linear mixed effects model (p = 1.4e-9 for strict vs. none, p = 1.6e-9 for strict vs. lenient, p = 0.98 for lenient vs. none). "*" indicates statistical significance after multiple comparisons correction with FDR q < 0.05 and "n.s." indicates not significant after FDR correction. For each box plot, the horizontal line indicates the median across all motion bins. The bottom and top edges of the box indicate the 25th and 75th percentiles, respectively. The outliers are defined as data points beyond 1.5 times the interquartile range. The whiskers extend to the most extreme data points not considered outliers.

#### **Continuous runs (two runs FD) for tree-based (Kong2026) TMS depression targets**

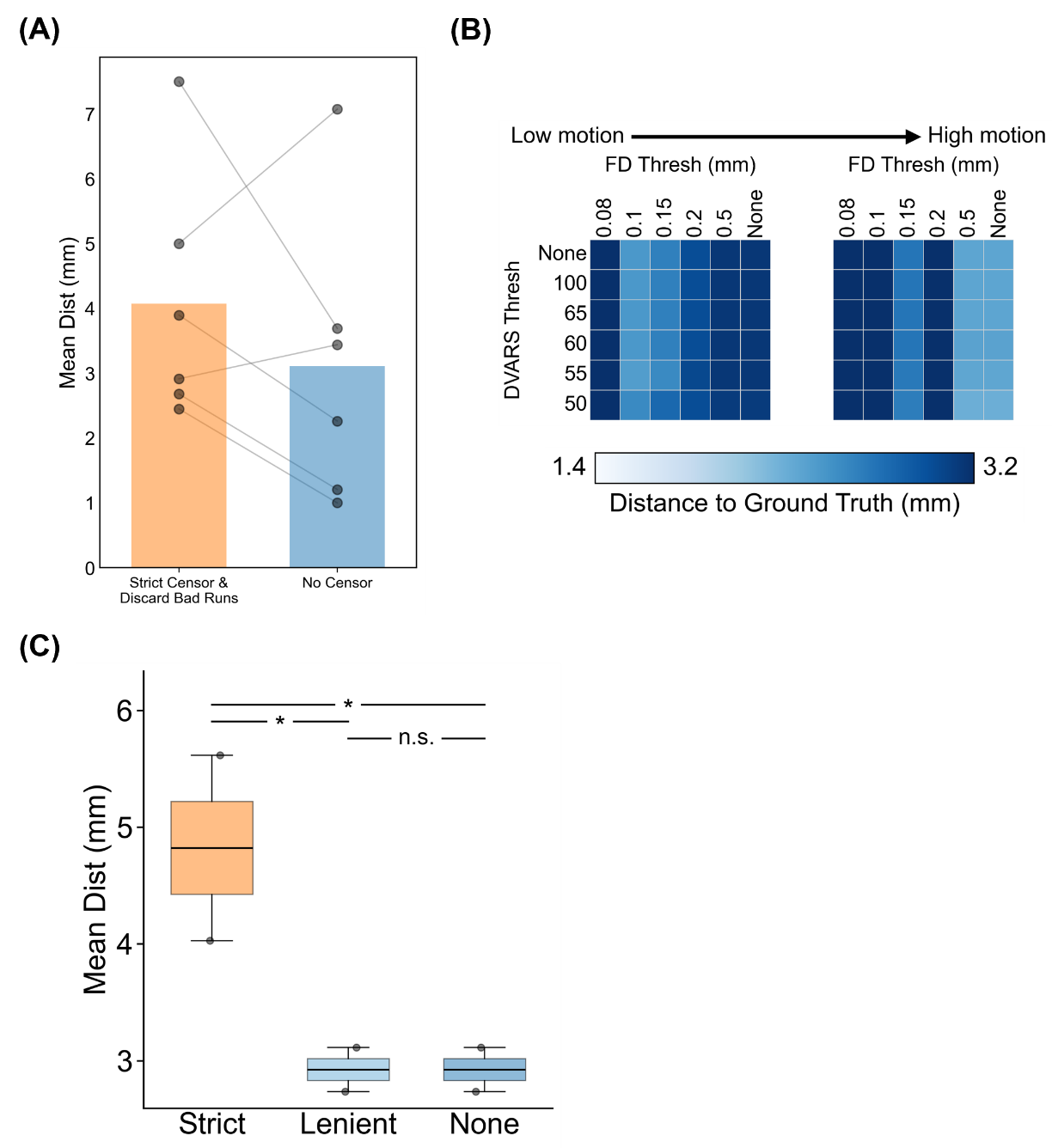

##### **Supplementary Figure 69.** Comparison of TMS depression targeting quality (Euclidean distance to ground-truth depression targets) using continuous real fMRI runs rather than simulated runs, for the two-run FD setup. Smaller Euclidean distance indicates better quality. TMS depression targets were estimated using the tree-algorithm (Kong et al., 2026a). (A) TMS depression targeting quality (Euclidean distance) for the two censoring approaches with mixed-motion runs. Here mixed-motion runs refer to sessions with one high-motion and one low-motion run, so the high-motion run will be discarded under strict censoring. Each dot represents the mean Euclidean distance across sessions of a participant in that motion bin. P-value was computed using the two-sided paired-sampled t-test. "*" indicates statistical significance. (B, C) Comparison of TMS depression targeting quality (Euclidean distance) across varying FD and DVARS thresholds for the two-run FD setup. For this analysis, high-motion runs were not discarded. (B) TMS depression targeting quality for different combinations of FD and DVARS thresholds for two motion bins. One motion bin has mixed-motion runs (sessions with one high-motion run and one low-motion run), and the other has two high-motion runs. Each cell in the grid indicates a motion bin. For each motion bin, there are 36 small cells corresponding to different FD and DVARS thresholds. (C) TMS depression targeting quality of strict censoring, lenient censoring and no censoring. Here strict censoring refers to FD threshold of 0.08 mm (after pseudo-motion filtering) and DVARS threshold of 50. Lenient censoring refers to FD threshold of 0.5 mm and DVARS threshold of 100. Each dot represents the mean Euclidean distance for a motion bin. The gray dashed line is the “noise ceiling”, achieved by picking the best combination of FD and DVARS threshold for each motion bin, and computing the median across motion bins. P-values were computed using a linear mixed effects model (p = 0.01 for strict vs. none, p = 0.01 for strict vs. lenient, p = 1.0 for lenient vs. none). "*" indicates statistical significance after multiple comparisons correction with FDR q < 0.05 and "n.s." indicates not significant after FDR correction. For each box plot, the horizontal line indicates the median across all motion bins. The bottom and top edges of the box indicate the 25th and 75th percentiles, respectively. The outliers are defined as data points beyond 1.5 times the interquartile range. The whiskers extend to the most extreme data points not considered outliers.

#### **Anxiety targeting (two runs FD) for Tree-based (Kong2026) TMS targets**

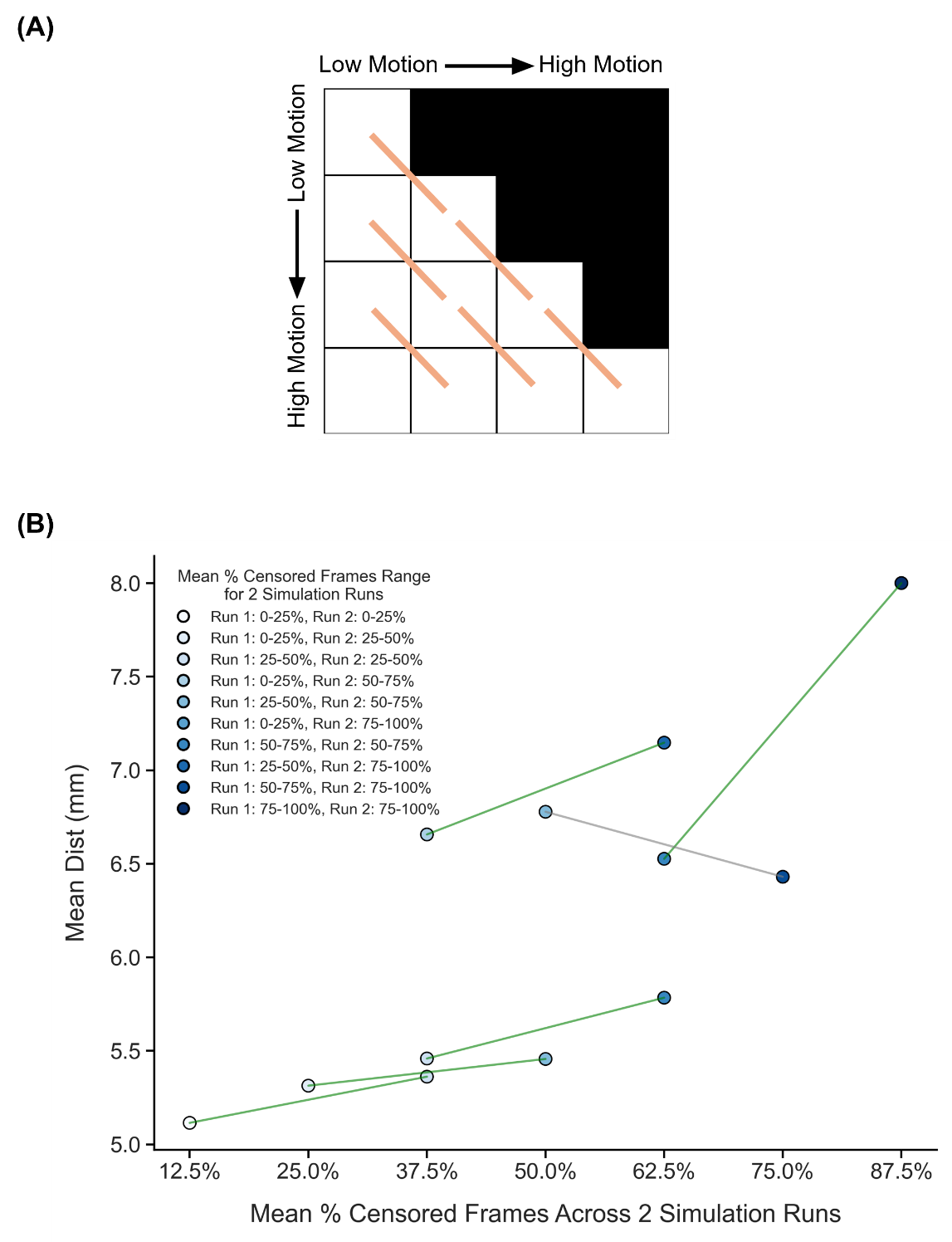

##### **Supplementary Figure 70.** Comparison of TMS anxiety target quality (Euclidean distance from ground-truth targets) across diagonally adjacent motion bins under strict censoring, for the two-run FD setup. Here strict censoring refers to FD threshold of 0.08 mm (after pseudo-motion filtering) and DVARS threshold of 50. High-motion runs were not discarded to allow comparison with low-motion runs. Smaller Euclidean distance to ground-truth target indicates better quality. TMS anxiety targets were estimated using the tree-algorithm (Kong et al., 2026a). (A) Schematic of comparisons across diagonally adjacent motion bins. Each orange line indicates a single comparison. (B) Relationship between TMS anxiety target quality (Euclidean distance) and motion. Each circle indicates the mean Euclidean distance across simulation runs of all participants in a motion bin. Each line indicates a single comparison between two adjacent motion bins. P-values were computed using the two-sided paired-sampled t-test using only participants common to both motion bins. An overall statistical test combining all comparisons was conducted using a linear mixed effects model (p = 0.03), but the effect was not significant after FDR correction.

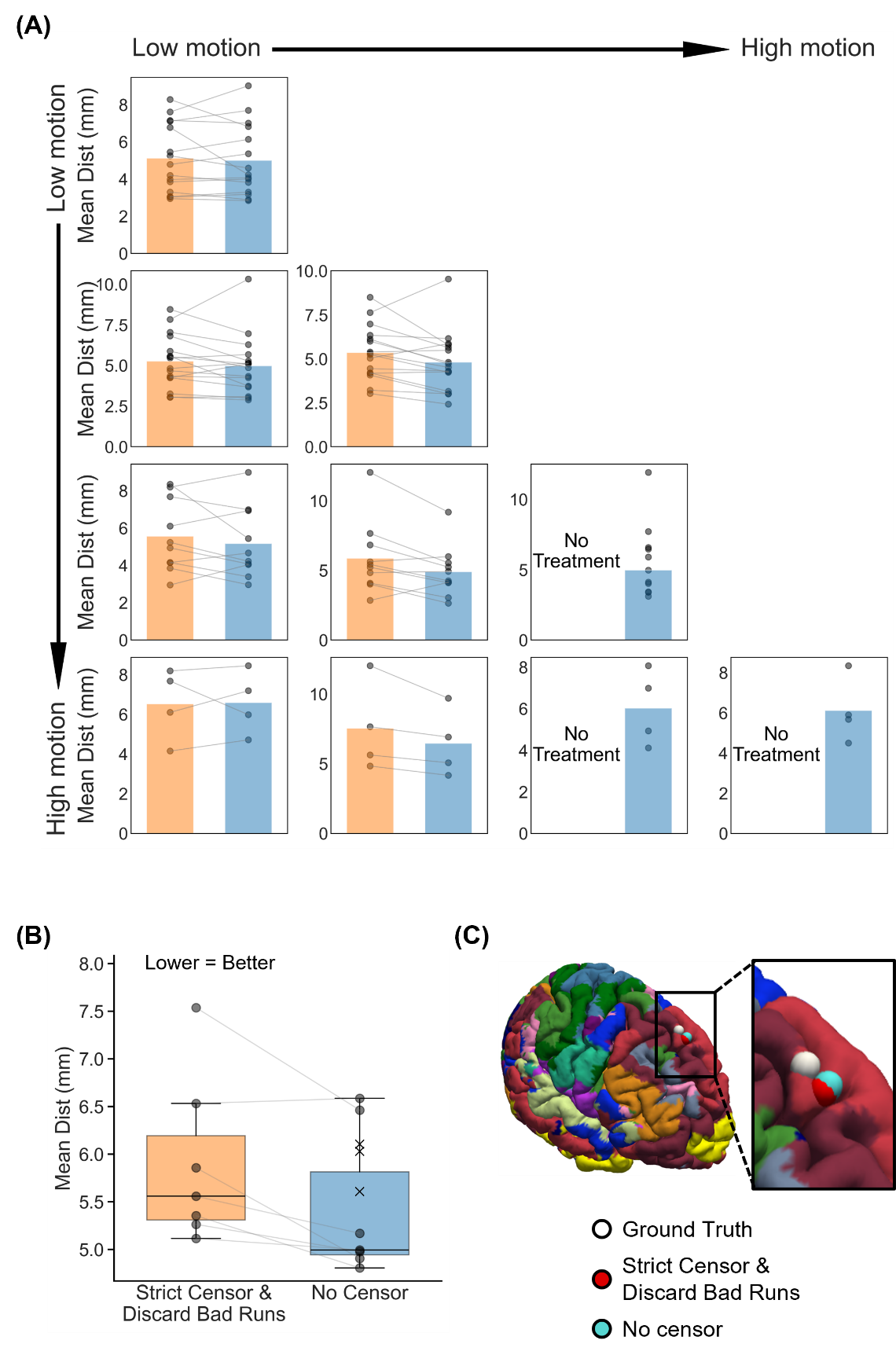

##### **Supplementary Figure 71.** Comparison of TMS anxiety targeting quality (Euclidean distance to ground-truth anxiety targets) between strict censoring and no censoring, for the two-run FD setup. Here strict censoring refers to FD threshold of 0.08 mm (after pseudo-motion filtering) and DVARS threshold of 50. High-motion runs were discarded. Smaller Euclidean distance indicates better quality. TMS anxiety targets were estimated using the tree-based algorithm (Kong et al., 2026a). (A) TMS anxiety targeting quality (Euclidean distance) for the two censoring approaches across 10 motion bins. Each cell in the grid indicates a motion bin. Each dot represents the mean Euclidean distance across 50 simulated sessions of a participant in that motion bin. P-values were computed using the two-sided paired-sampled t-test. "*" indicates statistical significance after multiple comparisons correction with FDR q < 0.05. Under strict censoring, no runs remained for the bottom-right three motion bins because all fMRI runs were discarded under this criterion. (B) TMS anxiety targeting quality across all motion bins for both censoring strategies. There are seven pairs of dots, corresponding to the seven motion bins where comparisons could be made between strict censoring and no censoring. P-value was computed using a linear mixed effects model (p = 0.02), but the effect was not significant after FDR correction. "*" indicates statistical significance after multiple comparisons correction with FDR q < 0.05. There are three “×” corresponding to the mean Euclidean distance of the 3 bottom-right bins under the no censoring condition. The × were not included in the boxplots. For each box plot, the horizontal line indicates the median. The bottom and top edges of the box indicate the 25th and 75th percentiles, respectively. The outliers are defined as data points beyond 1.5 times the interquartile range. The whiskers extend to the most extreme data points not considered outliers. (C) Visualization of TMS anxiety targets in a representative participant from the highest motion bin where the strict censoring had results.

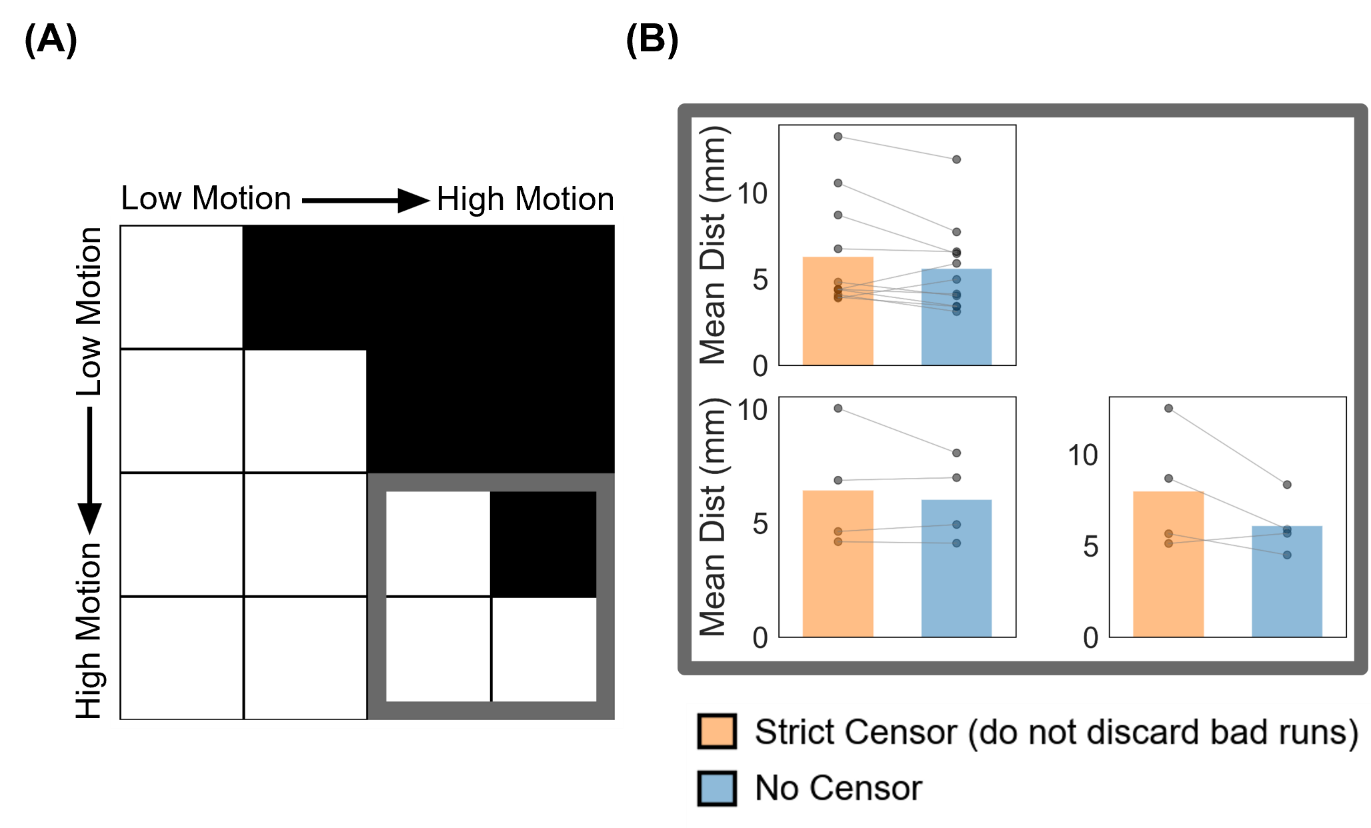

##### **Supplementary Figure 72.** Comparison of TMS anxiety targeting quality (Euclidean distance from ground-truth targets) between strict censoring and no censoring for the high-motion bins in the two-run FD setup. Here strict censoring refers to FD threshold of 0.08 mm (after pseudo-motion filtering) and DVARS threshold of 50. High-motion runs were not discarded for strict censoring, so a comparison could be made with no censoring. Smaller Euclidean distance indicates better quality. TMS anxiety targets were estimated using the tree-algorithm (Kong et al., 2026a). (A) Schematic highlighting the three high-motion bins with thick gray outlines. (B) TMS targeting quality (Euclidean distance) for the two censoring approaches across the 3 high-motion bins. Each cell in the grid indicates a motion bin. Each dot represents the mean Euclidean distance across 50 simulated sessions for a participant in that motion bin. P-values were computed using the two-sided paired-sampled t-test. "*" indicates statistical significance after multiple comparisons correction with FDR q < 0.05.

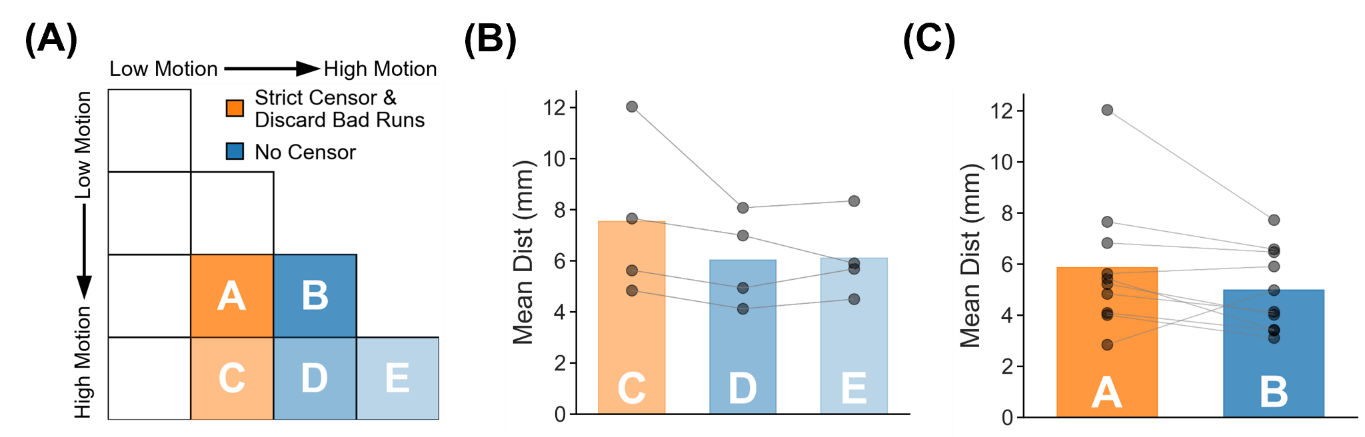

##### **Supplementary Figure 73.** TMS anxiety target quality (Euclidean distance to ground-truth targets) for two high-motion runs with no censoring vs mixed-motion runs with strict censoring. Here mixed-motion runs refer to sessions with one high-motion and one low-motion runs, so the high-motion run will be discarded under strict censoring. Strict censoring refers to FD threshold of 0.08 mm (after pseudo-motion filtering) and DVARS threshold of 50. Lower Euclidean distance indicates better quality. TMS anxiety targets were estimated using the tree-algorithm (Kong et al., 2026a). (A) Schematic illustrating the comparison. Motion bins A and B were compared, while motion bins C, D and E were compared. (B) Comparison of motion bins C vs D, and C vs E. (C) Comparison of motion bins A vs B. For both panels (B) and (C), each dot represents the mean Euclidean distance across 50 simulated sessions for a participant in that motion bin. P-values were computed using the two-sided paired-sampled t-test. Each statistical test was performed using only participants common to both motion bins. "*" indicates statistical significance after multiple comparisons correction with FDR q < 0.05.

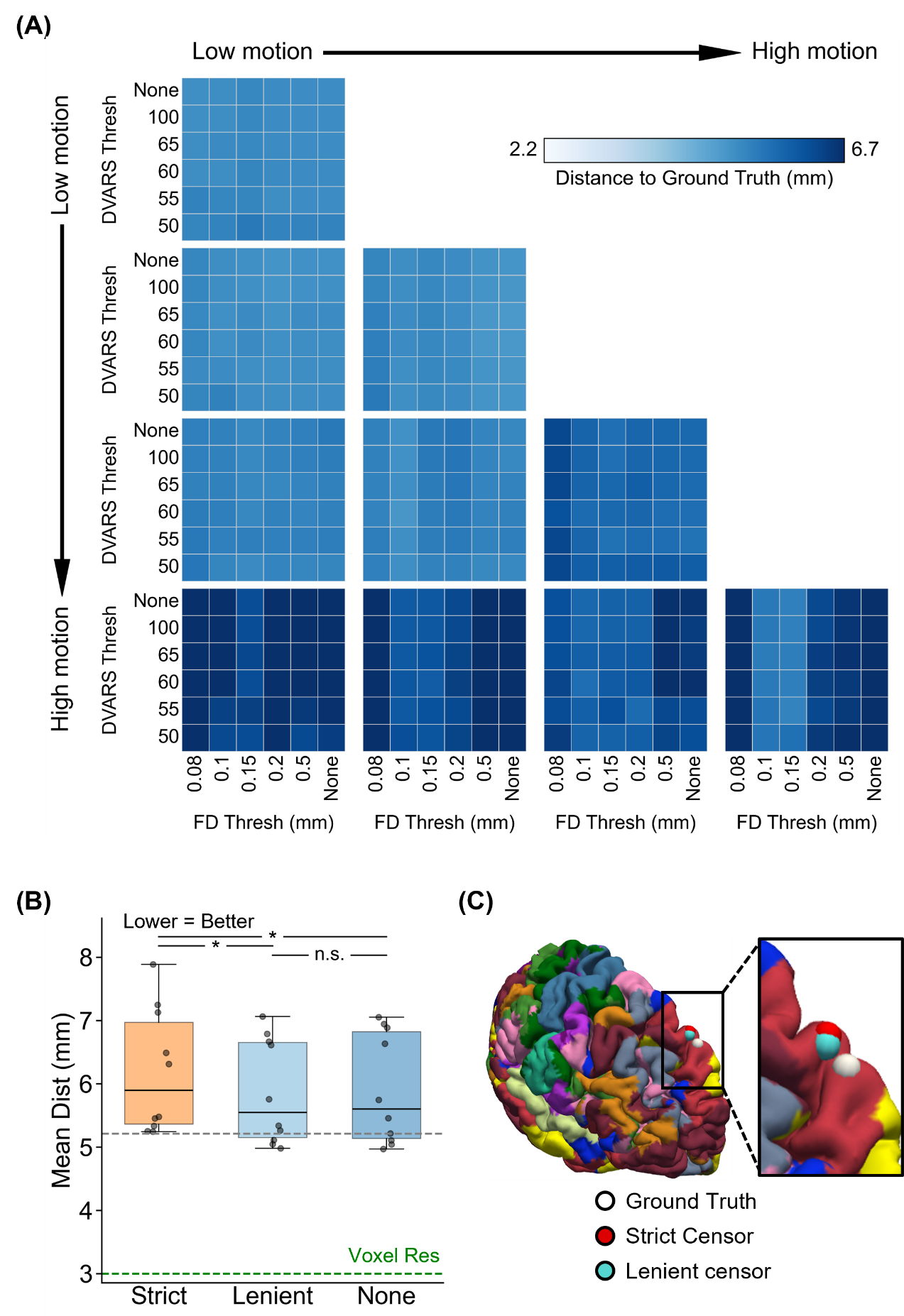

##### **Supplementary Figure 74.** Comparison of TMS anxiety target quality (Euclidean distance from ground-truth targets) across varying FD and DVARS thresholds for the two-run FD setup. For this analysis, high-motion runs were not discarded. Lower Euclidean distance indicates better quality. TMS anxiety targets were estimated using the tree-algorithm (Kong et al., 2026a). (A) TMS anxiety target quality (Euclidean distance) for different combinations of FD and DVARS thresholds for 10 motion bins. Each cell in the grid indicates a motion bin. For each motion bin, there are 36 small cells corresponding to different FD and DVARS thresholds. (B) TMS anxiety target quality for strict censoring, lenient censoring and no censoring. Here strict censoring refers to FD threshold of 0.08 mm (after pseudo-motion filtering) and DVARS threshold of 50. Lenient censoring refers to FD threshold of 0.5 mm and DVARS threshold of 100. Each dot represents the mean Euclidean distance for a motion bin. The gray dashed line is the “noise ceiling”, achieved by picking the best combination of FD and DVARS threshold for each motion bin, and computing the median across motion bins. P-values were computed using a linear mixed effects model (p = 5.1e-3 for strict vs. none, p = 2.5e-3 for strict vs. lenient, p = 0.82 for lenient vs. none). "*" indicates statistical significance after multiple comparisons correction with FDR q < 0.05 and "n.s." indicates not significant after FDR correction. For each box plot, the horizontal line indicates the median across all motion bins. The bottom and top edges of the box indicate the 25th and 75th percentiles, respectively. The outliers are defined as data points beyond 1.5 times the interquartile range. The whiskers extend to the most extreme data points not considered outliers. (C) Visualization of TMS anxiety targets in a representative participant from the highest motion bin.

#### **Anxiety targeting (two runs FDrms) for tree-based (Kong2026) TMS targets**

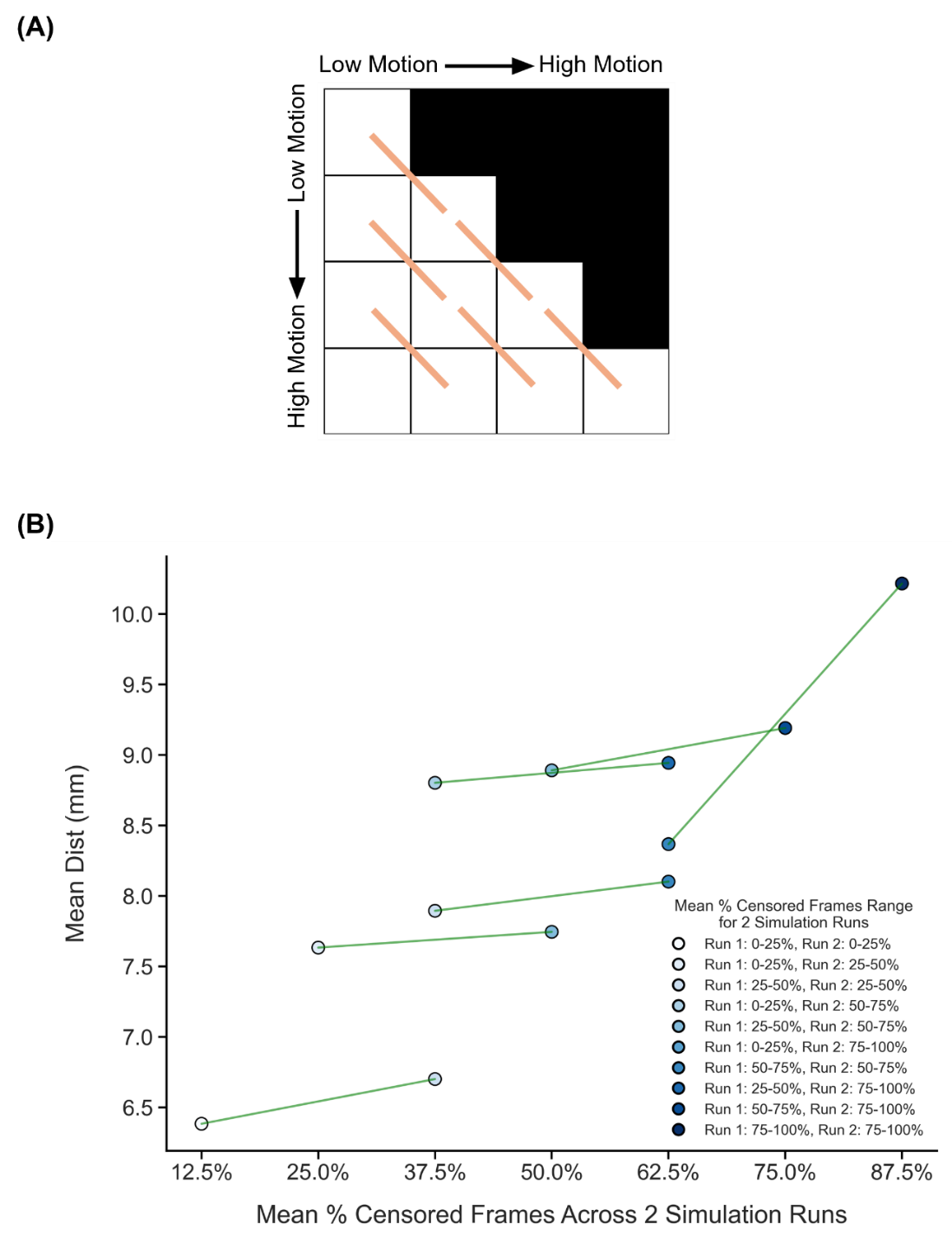

##### **Supplementary Figure 75.** Comparison of TMS anxiety target quality (Euclidean distance from ground-truth targets) across diagonally adjacent motion bins under strict censoring, for the two-run FDrms setup. Here strict censoring refers to FDrms threshold of 0.2 mm (without pseudo-motion filtering) and DVARS threshold of 50. High-motion runs were not discarded to allow comparison with low-motion runs. Smaller Euclidean distance to ground-truth target indicates better quality. TMS anxiety targets were estimated using the tree-algorithm (Kong et al., 2026a). (A) Schematic of comparisons across diagonally adjacent motion bins. Each orange line indicates a single comparison. (B) Relationship between TMS anxiety target quality (Euclidean distance) and motion. Each circle indicates the mean Euclidean distance across simulation runs of all participants in a motion bin. Each line indicates a single comparison between two adjacent motion bins. P-values were computed using the two-sided paired-sampled t-test using only participants common to both motion bins. An overall statistical test combining all comparisons was conducted using a linear mixed effects model (p = 6.3e-3).

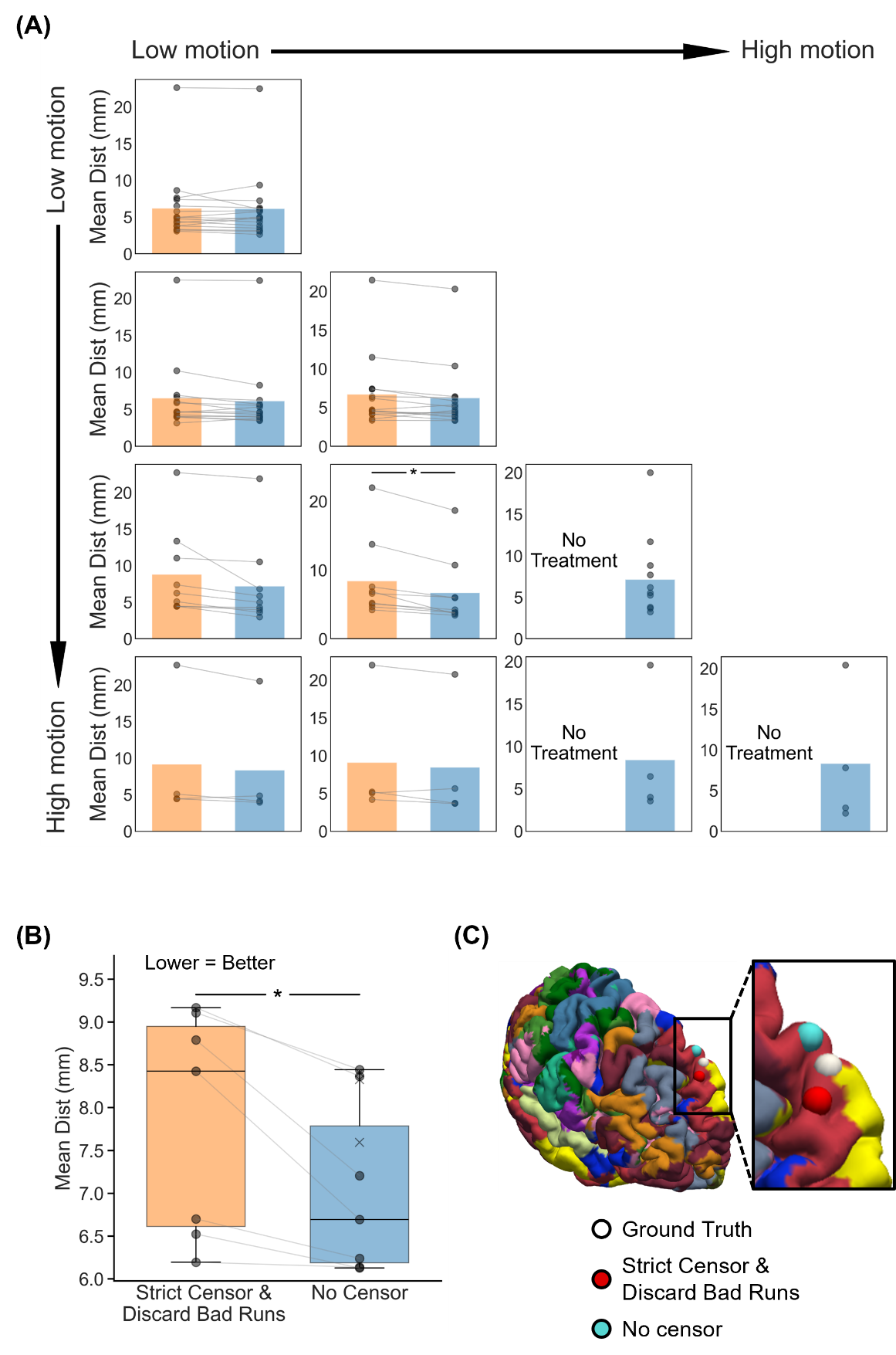

##### **Supplementary Figure 76.** Comparison of TMS anxiety targeting quality (Euclidean distance to ground-truth anxiety targets) between strict censoring and no censoring, for the two-run FDrms setup. Here strict censoring refers to FDrms threshold of 0.2 mm (without pseudo-motion filtering) and DVARS threshold of 50. High-motion runs were discarded. Smaller Euclidean distance indicates better quality. TMS anxiety targets were estimated using the tree-based algorithm (Kong et al., 2026a). (A) TMS anxiety targeting quality (Euclidean distance) for the two censoring approaches across 10 motion bins. Each cell in the grid indicates a motion bin. Each dot represents the mean Euclidean distance across 50 simulated sessions of a participant in that motion bin. P-values were computed using the two-sided paired-sampled t-test. "*" indicates statistical significance after multiple comparisons correction with FDR q < 0.05. Under strict censoring, no runs remained for the bottom-right three motion bins because all fMRI runs were discarded under this criterion. (B) TMS anxiety targeting quality across all motion bins for both censoring strategies. There are seven pairs of dots, corresponding to the seven motion bins where comparisons could be made between strict censoring and no censoring. P-value was computed using a linear mixed effects model (p = 0.01). "*" indicates statistical significance after multiple comparisons correction with FDR q < 0.05. There are three “×” corresponding to the mean Euclidean distance of the 3 bottom-right bins under the no censoring condition. The × were not included in the boxplots. For each box plot, the horizontal line indicates the median. The bottom and top edges of the box indicate the 25th and 75th percentiles, respectively. The outliers are defined as data points beyond 1.5 times the interquartile range. The whiskers extend to the most extreme data points not considered outliers. (C) Visualization of TMS anxiety targets in a representative participant from the highest motion bin where the strict censoring had results.

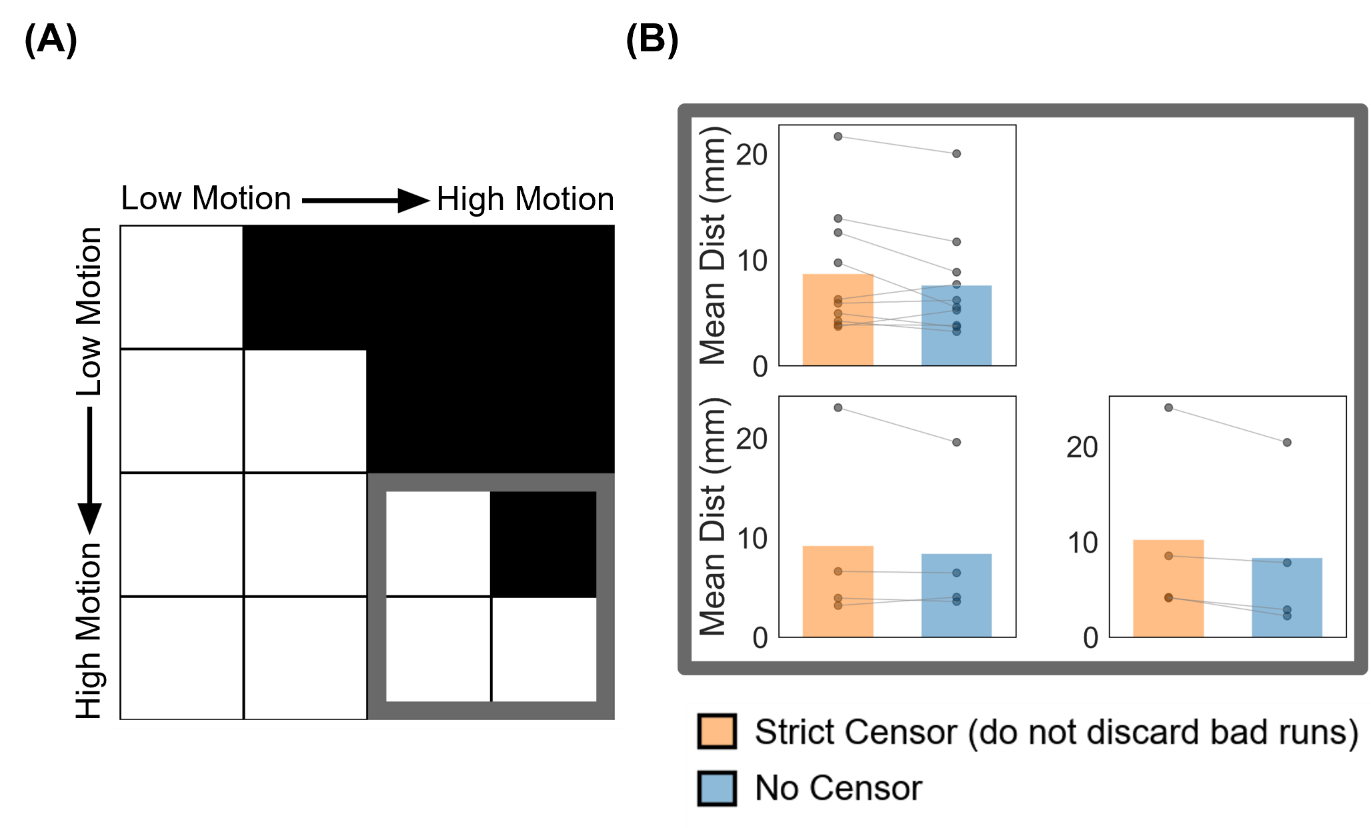

##### **Supplementary Figure 77.** Comparison of TMS anxiety targeting quality (Euclidean distance from ground-truth targets) between strict censoring and no censoring for the high-motion bins in the two-run FDrms setup. Here strict censoring refers to FDrms threshold of 0.2 mm (without pseudo-motion filtering) and DVARS threshold of 50. High-motion runs were not discarded for strict censoring, so a comparison could be made with no censoring. Smaller Euclidean distance indicates better quality. TMS anxiety targets were estimated using the tree-algorithm (Kong et al., 2026a). (A) Schematic highlighting the three high-motion bins with thick gray outlines. (B) TMS targeting quality (Euclidean distance) for the two censoring approaches across the 3 high-motion bins. Each cell in the grid indicates a motion bin. Each dot represents the mean Euclidean distance across 50 simulated sessions for a participant in that motion bin. P-values were computed using the two-sided paired-sampled t-test. "*" indicates statistical significance after multiple comparisons correction with FDR q < 0.05.

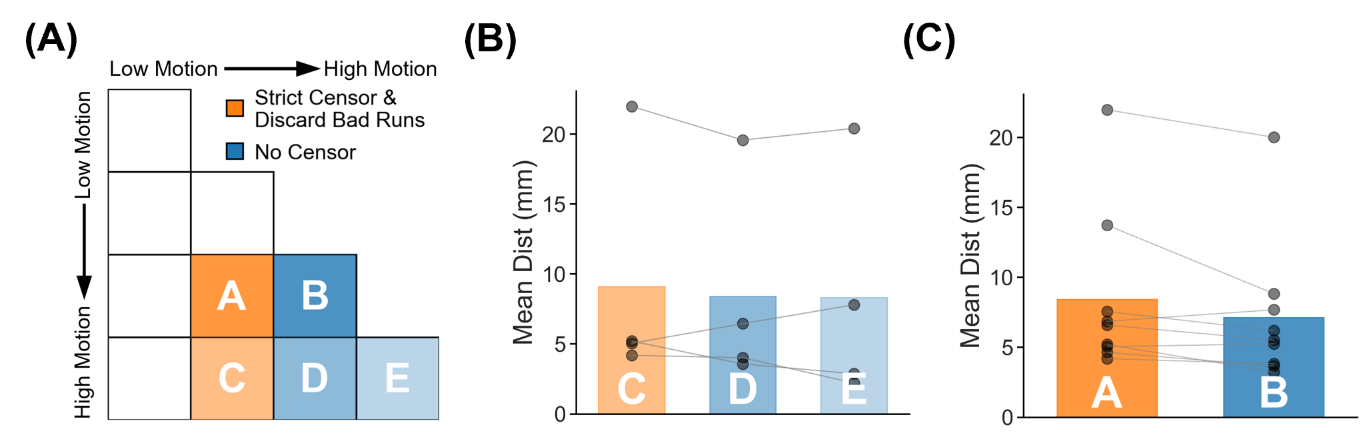

##### **Supplementary Figure 78.** TMS anxiety target quality (Euclidean distance to ground-truth targets) for two high-motion runs with no censoring vs mixed-motion runs with strict censoring. Here mixed-motion runs refer to sessions with one high-motion and one low-motion runs, so the high-motion run will be discarded under strict censoring. Strict censoring refers to FDrms threshold of 0.2 mm (without pseudo-motion filtering) and DVARS threshold of 50. Lower Euclidean distance indicates better quality. TMS anxiety targets were estimated using the tree-algorithm (Kong et al., 2026a). (A) Schematic illustrating the comparison. Motion bins A and B were compared, while motion bins C, D and E were compared. (B) Comparison of motion bins C vs D, and C vs E. (C) Comparison of motion bins A vs B. For both panels (B) and (C), each dot represents the mean Euclidean distance across 50 simulated sessions for a participant in that motion bin. P-values were computed using the two-sided paired-sampled t-test. Each statistical test was performed using only participants common to both motion bins. "*" indicates statistical significance after multiple comparisons correction with FDR q < 0.05.

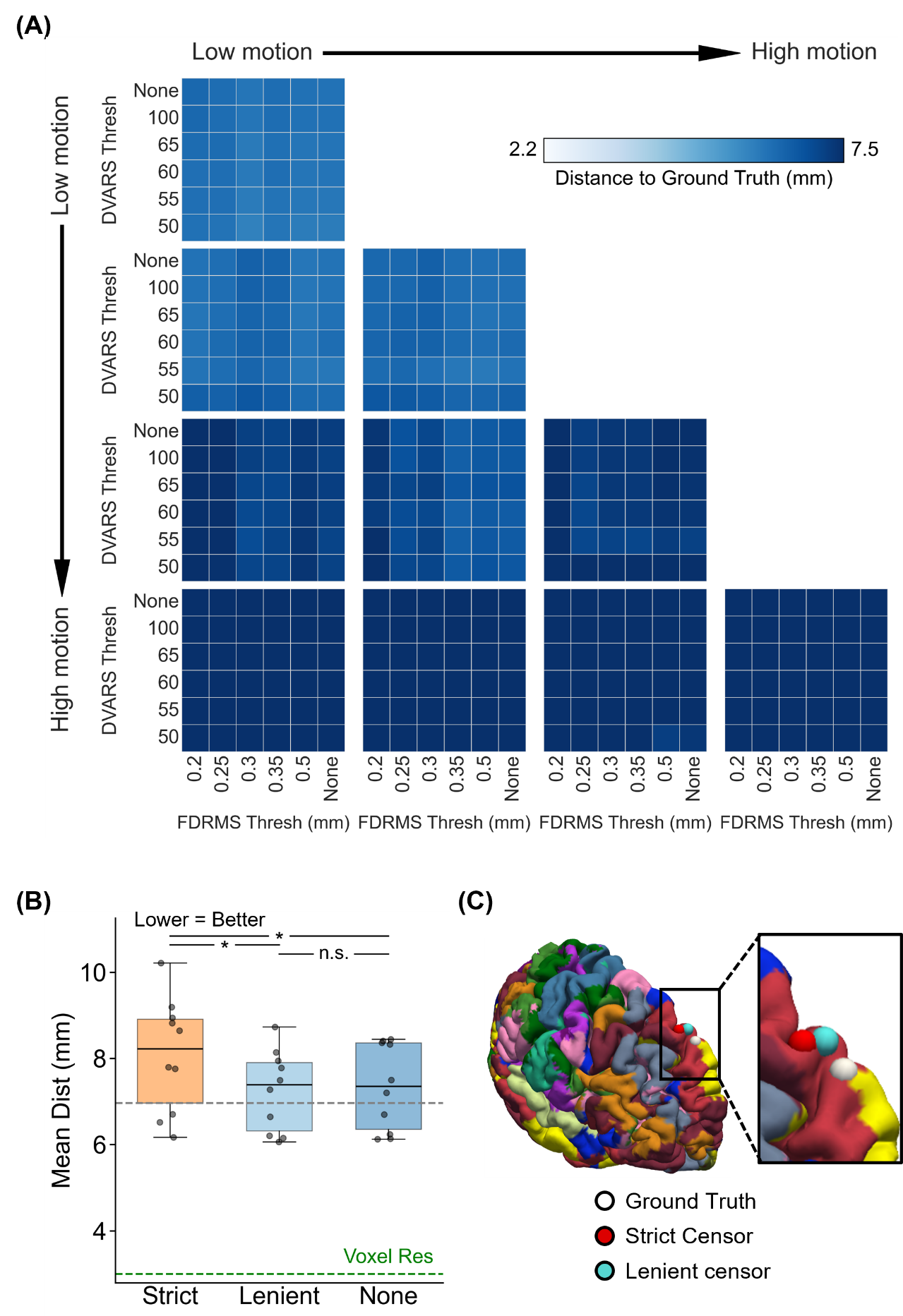

##### **Supplementary Figure 79.** Comparison of TMS anxiety target quality (Euclidean distance from ground-truth targets) across varying FDrms and DVARS thresholds for the two-run FDrms setup. For this analysis, high-motion runs were not discarded. Lower Euclidean distance indicates better quality. TMS anxiety targets were estimated using the tree-algorithm (Kong et al., 2026a). (A) TMS anxiety target quality (Euclidean distance) for different combinations of FDrms and DVARS thresholds for 10 motion bins. Each cell in the grid indicates a motion bin. For each motion bin, there are 36 small cells corresponding to different FDrms and DVARS thresholds. (B) TMS anxiety target quality for strict censoring, lenient censoring and no censoring. Here strict censoring refers to FDrms threshold of 0.2 mm (without pseudo-motion filtering) and DVARS threshold of 50. Lenient censoring refers to FDrms threshold of 0.5 mm and DVARS threshold of 100. Each dot represents the mean Euclidean distance for a motion bin. The gray dashed line is the “noise ceiling”, achieved by picking the best combination of FDrms and DVARS threshold for each motion bin, and computing the median across motion bins. P-values were computed using a linear mixed effects model (p = 4.2e-8 for strict vs. none, p = 2.4e-9 for strict vs. lenient, p = 0.63 for lenient vs. none). "*" indicates statistical significance after multiple comparisons correction with FDR q < 0.05 and "n.s." indicates not significant after FDR correction. For each box plot, the horizontal line indicates the median across all motion bins. The bottom and top edges of the box indicate the 25th and 75th percentiles, respectively. The outliers are defined as data points beyond 1.5 times the interquartile range. The whiskers extend to the most extreme data points not considered outliers. (C) Visualization of TMS anxiety targets in a representative participant from the highest motion bin.

#### **Discussion section**

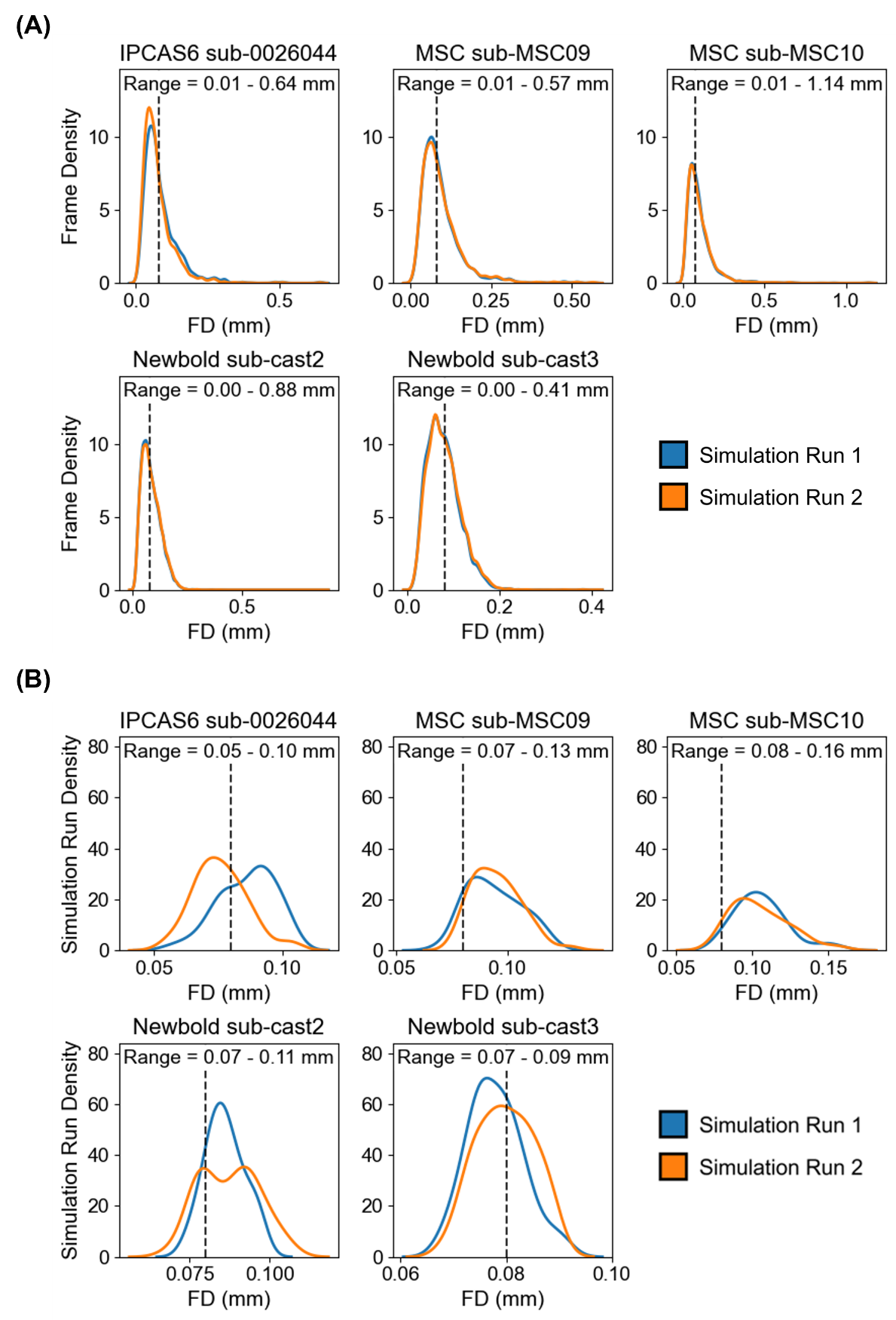

##### **Supplementary Figure 80.** FD distributions for each participant from the highest-motion bin in the two-run FD setup. Recall that the FD values were obtained after pseudo-motion filtering. (A) Each observation in the distribution corresponds to the FD of a single fMRI frame. The dotted black line indicates the FD threshold of 0.08 mm. Maximum frame-level FD was 1.14 mm, observed in sub-MSC10 from the MSC dataset. (B) Each observation in the distribution corresponds to the mean FD averaged across all frames in a simulated run. Maximum run-level FD was 0.16 mm, observed in sub-MSC10 from the MSC dataset.

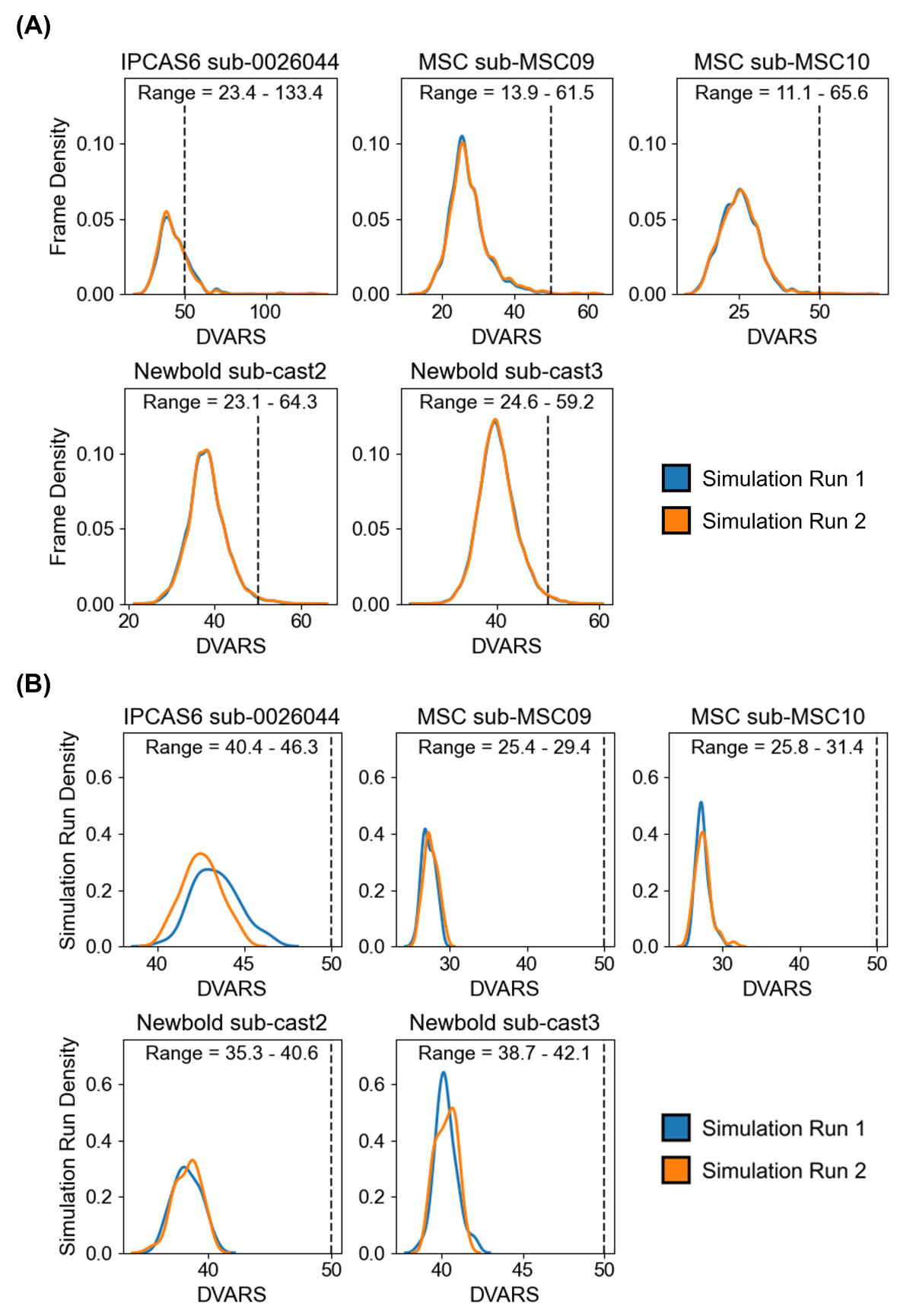

##### **Supplementary Figure 81.** DVARS distributions for each participant from the highest-motion bin in the two-run FD setup. (A) Each observation in the distribution corresponds to the DVARS of a single fMRI frame. The dotted black line indicates the DVARS threshold of 50. Maximum frame-level DVARS was 133.4, observed in sub-0026044 from the IPCAS6 dataset. (B) Each observation in the distribution corresponds to the mean DVARS averaged across all frames in a simulated run. Maximum run-level DVARS was 46.3, observed in sub-0026044 from the IPCAS6 dataset.

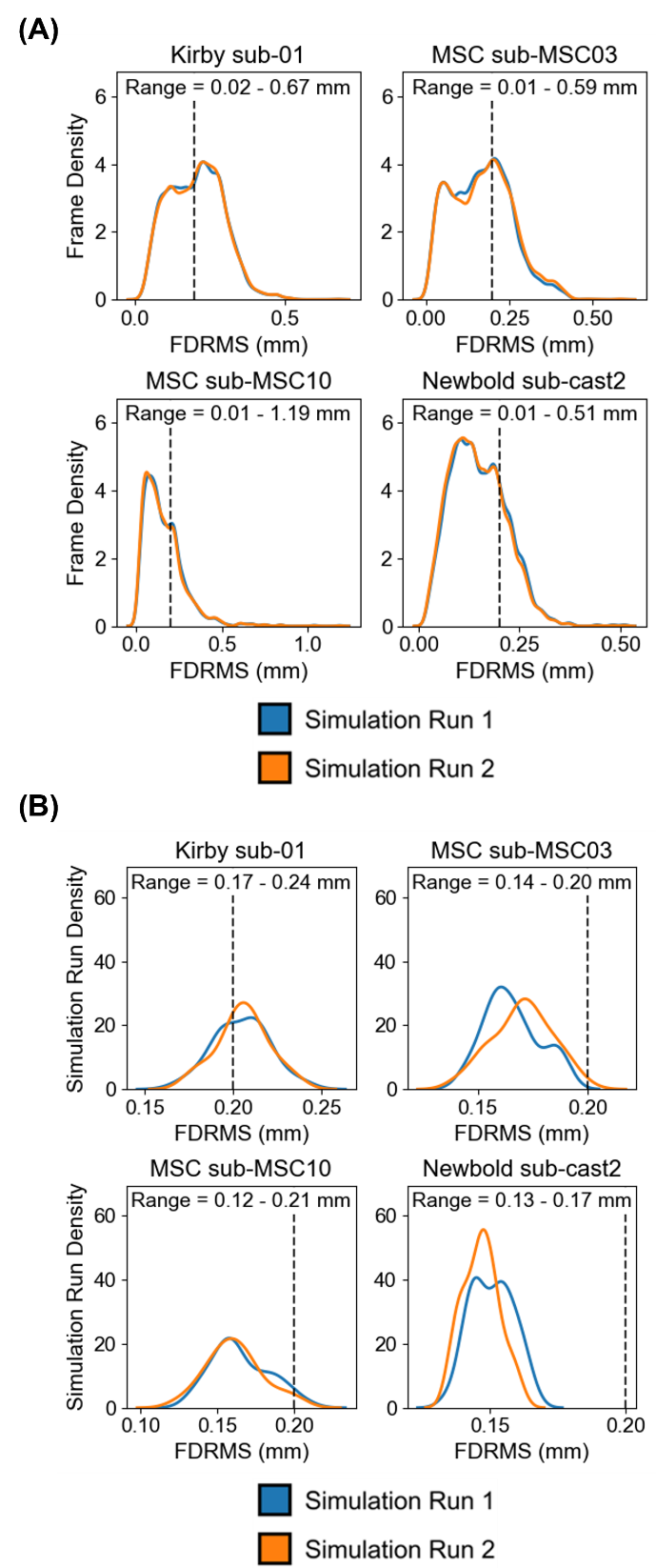

##### **Supplementary Figure 82.** FDrms distributions for each participant from the highest-motion bin in the two-run FDrms setup. Recall that the FDrms values were obtained without pseudo-motion filtering. (A) Each observation in the distribution corresponds to the FDrms of a single fMRI frame. The dotted black line indicates the FDrms threshold of 0.2 mm. Maximum frame-level FDrms was 1.19 mm, observed in sub-MSC10 from the MSC dataset. (B) Each observation in the distribution corresponds to the mean FDrms averaged across all frames in a simulated run. Maximum run-level FDrms was 0.24 mm, observed in sub-01 from the Kirby dataset.

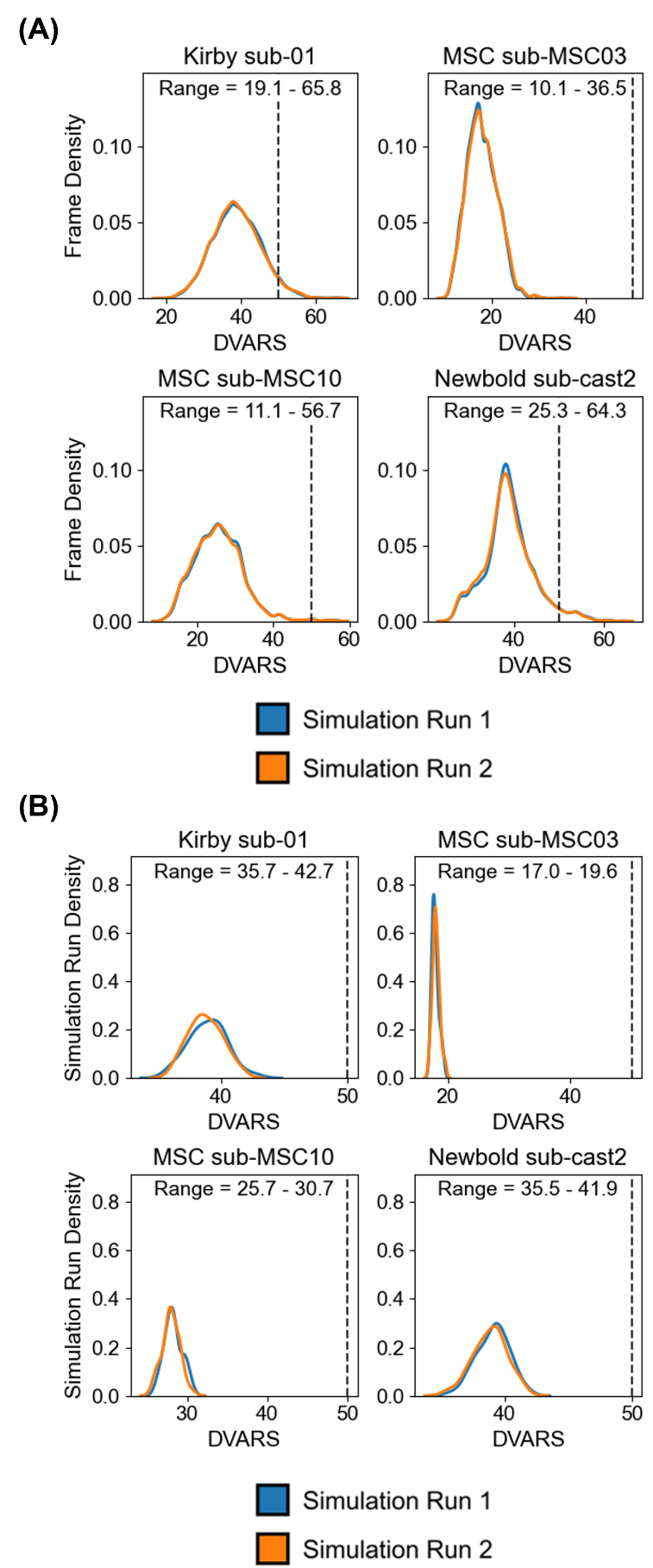

##### **Supplementary Figure 83.** DVARS distributions for each participant from the highest-motion bin in the two-run FDrms setup. (A) Each observation in the distribution corresponds to the DVARS of a single fMRI frame. The dotted black line indicates the DVARS threshold of 50. Maximum frame-level DVARS was 65.8, observed in sub-01 from the Kirby dataset. (B) Each observation in the distribution corresponds to the mean DVARS averaged across all frames in a simulated run. Maximum run-level DVARS was 42.7, observed in sub-01 from the Kirby dataset.
